# Water content, transition temperature and fragility influence protection and anhydrobiotic capacity

**DOI:** 10.1101/2023.06.30.547256

**Authors:** John F. Ramirez, U.G.V.S.S. Kumara, Navamoney Arulsamy, Thomas C. Boothby

## Abstract

Water is essential for metabolism and all life processes. Despite this, many organisms distributed across the kingdoms of life survive near-complete desiccation or anhydrobiosis (Greek for “life without water”). Increased intracellular viscosity, leading to the formation of a vitrified state is necessary, but not sufficient, for survival while dry. What properties of a vitrified system make it desiccation-tolerant or -sensitive are unknown. We have analyzed 18 different *in vitro* vitrified systems, composed of one of three protective disaccharides (trehalose, sucrose, or maltose) and varying amounts of glycerol, quantifying their enzyme-protective capacity and their material properties in a dry state. We find that protection conferred by mixtures containing maltose correlates strongly with increased water content, increased glass-transition temperature, and reduced glass former fragility, while the protection of glasses formed with sucrose correlates with increased glass transition temperature and the protection conferred by trehalose glasses correlates with reduced glass former fragility. Thus, *in vitro* different vitrified sugars confer protection through distinct material properties. Extending on this, we have examined the material properties of a dry desiccation tolerant and intolerant life stage from three different organisms. In all cases, the dried desiccation tolerant life stage of an organism had an increased glass transition temperature relative to its dried desiccation intolerant life stage, and this trend is also seen in all three organisms when considering reduced glass former fragility. These results suggest that while drying of different protective sugars *in vitro* results in vitrified systems with distinct material properties that correlate with their enzyme-protective capacity, in nature organismal desiccation tolerance relies on a combination of these properties. This study advances our understanding of how protective and non-protective glasses differ in terms of material properties that promote anhydrobiosis. This knowledge presents avenues to develop novel stabilization technologies for pharmaceuticals that currently rely on the cold-chain.

**Statement of significance:** For the past three decades the anhydrobiosis field has lived with a paradox, while vitrification is necessary for survival in the dry state, it is not sufficient. Understanding what property(s) distinguishes a desiccation tolerant from an intolerant vitrified system and how anhydrobiotic organisms survive drying is one of the enduring mysteries of organismal physiology. Here we show *in vitro* the enzyme-protective capacity of different vitrifying sugars can be correlated with distinct material properties. However, *in vivo,* diverse desiccation tolerant organisms appear to combine these material properties to promote their survival in a dry state.

**Highlights:**

- The enzyme-protective capacities of different glass forming sugars correlate with distinct material properties.
- Material properties of dried anhydrobiotic organisms differ dramatically when examined in desiccation tolerant and intolerant life stages.
- Organismal desiccation tolerance is concomitant with changes in glassy properties including increased glass transition temperature and reduced glass former fragility.

## 4.1 Introduction

Water is required for metabolism and so is often considered essential for life. However, a number of organisms, spread across every biological kingdom, are capable of surviving near-complete water loss through a process known as anhydrobiosis (Greek for “life without water”) [1]. As organisms dry, they face a number of physical and chemical changes to their cellular environment [1,2]. As water is lost, cellular constituents are concentrated, molecular crowding increases, pH and ionic concentrations change, and osmotic pressure increases [2]. These physiochemical changes lead to detrimental perturbations such as protein unfolding, aggregation, and membrane leakage [2]. Importantly, drying is not an all-or-nothing process and these changes as well as the perturbation they induce, occur along a continuum, with some perturbations occurring earlier as an organism is dehydrating while others manifest later, once more substantial amounts of water have been lost [1–4]. How organisms survive desiccation is one of the enduring mysteries of organismal physiology.

Historically, anhydrobiosis has been thought to be mediated, at least in part, through the concentration of cellular constituents until these constituents solidify into a vitrified material (a glass). In this hypothesis, known as the ‘vitrification hypothesis,’ glasses slow physical and biochemical change, making them natural promoters of desiccation tolerance. Within the anhydrobiosis field, vitrification is considered a necessary process for desiccation tolerance [5–8].

However, a major shortcoming of the vitrification hypothesis has been known for decades, namely the observation that essentially every biological, or sufficiently heterogeneous, system will vitrify when dried, regardless of whether it is desiccation-tolerant or -sensitive [1,5–7]. This observation implies that while vitrification is necessary, it is not sufficient for desiccation tolerance and that there must be some property, or properties, that distinguishes a protective from a non-protective vitrified state [5,9]. The properties distinguishing a desiccation-protective glass from a non-protective glass are not currently fully understood.

Previous studies have identified that small additions of glycerol changes the enzyme-protective capacity of trehalose [10]. However, the material properties of these mixtures and how they correspond with changes in the level of protection have not been investigated. To address this gap in knowledge, we test the hypothesis that the enzyme-protective capacity of disaccharide-glycerol mixtures during desiccation correlates with their material properties. These properties include water content, glass transition temperature, and glass former fragility.

Water content is a property of vitrified materials that has been implicated in survival during extreme desiccation [4,11,12]. Water content refers to the mass percent of water in a desiccated sample. This can be measured by taking the starting mass of a desiccated sample and dividing by the mass of the sample after heating to a temperature sufficient to evaporate residual water. While hydrated, water molecules within a cell are able to solvate and then stabilize sensitive intracellular components. By retaining more water, it has been proposed that a vitrified material could prevent damage to sensitive intracellular components by maintaining hydration shells around them [3,4]. Additionally, residual water is implicated in several other proposed mechanisms of desiccation tolerance such as water entrapment [13–17], preferential exclusion [16,18], and the anchorage hypothesis [19–22].

Glass transition temperature (*T_g_*) is the temperature at which a vitrified solid transitions from a glassy to a rubbery state [23–25]. Increases or decreases to the *T_g_* of a vitrified material occur through the inclusion of an additive [26]. Increasing the glass transition temperature of a vitrified material has been observed to increase the shelf-life of sensitive proteins in a dry state [27] and is implicated in being essential for survival during desiccation at low relative humidity [3]. At the organismal level, it has been demonstrated that many anhydrobiotic organisms survive heating up to, but not beyond, their *T_g_* [6,28]. This suggests that anhydrobiotic organisms rely on being in a vitrified state and the production of small molecules which increase *T_g_* may be an effective strategy for increasing desiccation tolerance, or at least for increasing thermal tolerance while desiccated.

Finally, glass former fragility has been implicated as a key property of vitrified materials that promotes desiccation tolerance [9]. This property distinguishes strong glass forming materials, whose viscosity increases steadily well before the liquid-to-solid transition, from fragile glass forming materials, whose viscosity increases slowly at first but then rises abruptly at the onset of vitrification [9,23,29]. It should be noted that in this context, glass fragility/strength does not refer to the brittleness of a vitrified material, but rather to how the viscosity of the material changes as it approaches the point of vitrification. It is hypothesized that strong glass forming materials confer more protection during desiccation than their more fragile counterparts [4,9,23,29]. This hypothesis relies on the logic that a fragile glass forming material will not produce a sufficiently viscous state to slow down or prevent perturbation such as protein unfolding and aggregation until it is too late. Conversely, a strong glass forming material will increase in viscosity and provide protection along the continuum of drying.

To empirically test which, if any, of these three material properties correlate with desiccation tolerance, we first used a panel of simple reductive systems, each composed of two mediators of desiccation tolerance - a disaccharide including maltose [30,31], sucrose [1,5,32], or trehalose [6,7,28,33–35] and the polyol glycerol [36–38]. Comparing the measured material properties (water content, *T_g_*, and glass former fragility) of our disaccharide-glycerol glasses with their *in vitro* enzyme-protective capacities (Fig. 1a), we find that there is not a strict pattern in terms of the correlation of material properties to protection that all disaccharides follow. Instead, it appears that each disaccharide has a particular material property that is best correlated with its enzyme-protective capacity.

**Figure 1:**
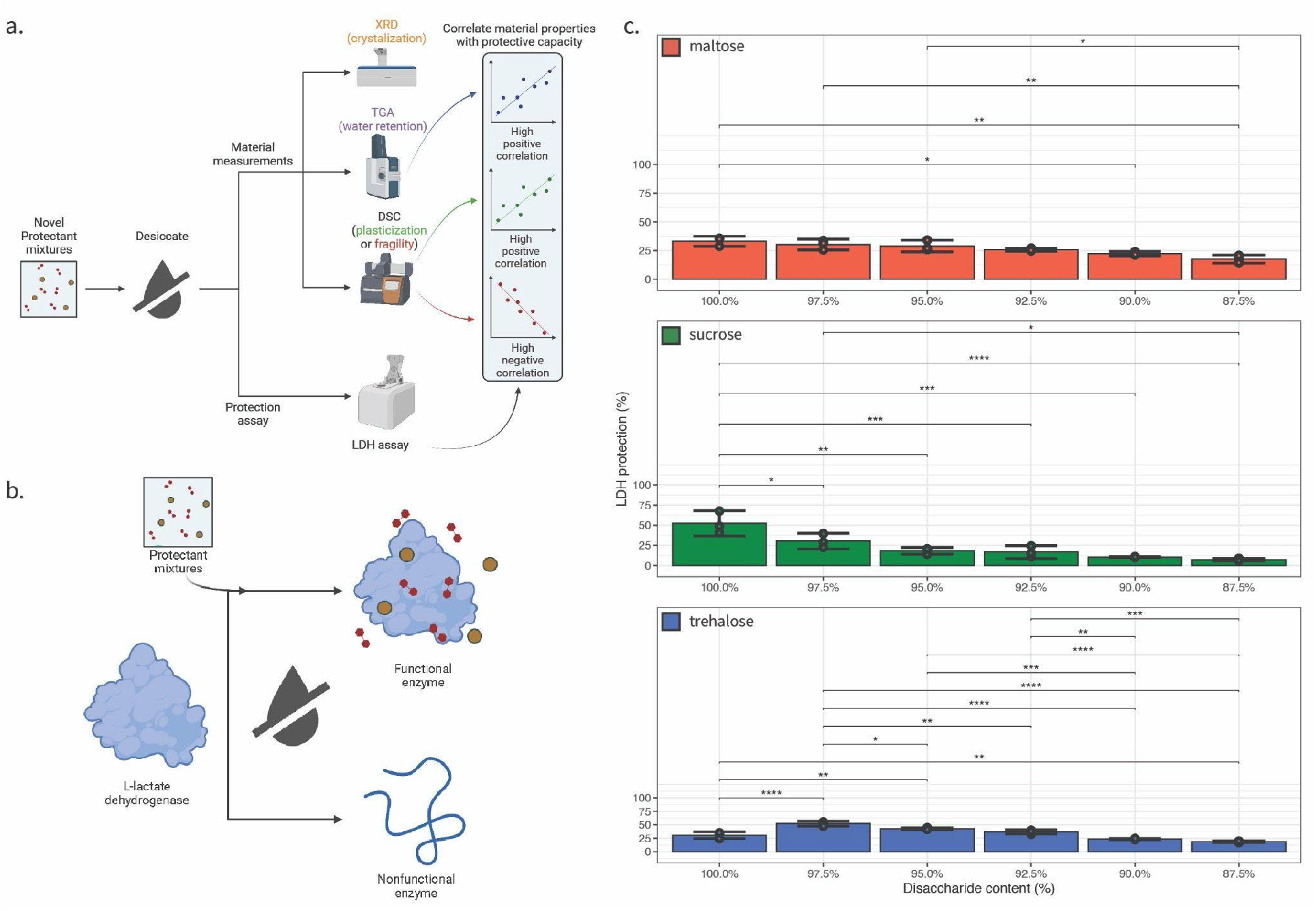
Vitrified disaccharide-glycerol mixtures display variable enzyme-protective properties. a) Schematic overview of workflow to produce correlation values between material properties and enzyme protection effect. b) Schematic of LDH assay. c) Lactate dehydrogenase (LDH) enzyme protection values for disaccharide-glycerol mixtures organized by disaccharide. Statistics calculated using a one-way ANOVA and Tukey post-hoc test: p-value ≤ 0.05 (*) p-value ≤ 0.01 (**), p-value ≤ 0.001 (***), pairwise relationships not shown are not significant, error bars represent 95% CI.

We extend this analysis into three organismal systems, each of which has a well-characterized desiccation tolerant and intolerant life stage(s). We find that while the enzyme-protective capacity of our *in vitro* systems tend to correlate with a single material property, our *in vivo* studies showed both increased glass transition temperature and reduced glass former fragility are hallmarks of desiccation tolerance. This suggests that anhydrobionts modulate multiple material properties to promote organismal desiccation tolerance.

The extension of our mechanistic understanding of the principles underlying desiccation-tolerance provides insights into how organisms can cope with changing, often extreme, environments. These findings also help to address the decades-long paradox that some glasses are protective, and others not, by providing insights into material properties that promote the protective capacity of different vitrifying mixtures. Our work identifies increased glass transition temperature and glass former fragility as major considerations in the development of technologies for the dry preservation of pharmaceuticals and the engineering of crops that are better able to withstand climate change and extreme weather.

## 5.1 Materials and Methods

### 5.2 Disaccharide-glycerol mixtures

D-Maltose monohydrate was sourced from Caisson Labs (M004-500GM). D-Sucrose was sourced from Sigma-Aldrich (S0389-500G). D-Trehalose dihydrate was sourced from VWR (VWRB3599-1KG). Glycerol was sourced from Biobasic (GB0232). Mixtures of each disaccharide and glycerol were made in 25 mM Tris at pH 7.0. Individual masses of each component were formulated (weight by weight) to additively produce mixtures of 10 g/L.

### 5.3 Sample Desiccation

Samples were desiccated using a speedvac (Savant SpeedVac SC110 with a Thermo OFP400 vacuum pump) for 16 hours. Prior to desiccation, 1 mL aliquot samples were dispensed into individual plastic weigh-boats (for aqueous samples) and at least 200 mg of organism samples were similarly loaded into individual plastic weigh-boats. The greater surface area of the weigh-boat, as opposed to Eppendorf tubes, allows for even desiccation of the entire sample, which reduces noise on the DSC. After the 16-hour desiccation, DSC samples were transferred to pre-massed pairs of DSC aluminum hermetic pan and aluminum hermetic lids (TA 900793.901 and 901684.901, respectively) while TGA samples were transferred to pre-tared platinum crucibles (TA 957207.904), and XRD samples were kept in the desiccation weigh-boats. DSC sample masses were determined after the sample was sealed within the pan and lid.

### 5.4 Single crystal X-ray diffractometry

Powder diffraction patterns for the samples were measured at 20 °C on a Bruker SMART APEX II CCD area detector system equipped with a graphite monochromator and a Mo K fine-focus sealed tube operated at 1.2 kW power. The dried samples were rolled into a ball of approximate diameter 0.5 mm, mounted on a goniometer head using a glass fiber, and centered using the APEX2 software. The detector was placed at 6.12 cm during the data collection. Four frames of data were collected at four different sets of angles with a scan width of 0.2 ° and an exposure time of 3 min per frame. The frames were integrated using the APEX2 program. The measured powder diffraction images were integrated, and the data were plotted in the 5 to 50 ° 2θ. All analysis was performed using the APEX3 Software Suite V2017.3-0, Bruker AXS Inc.: Madison, WI, 2017.

### 5.5 Lactate dehydrogenase (LDH) enzyme protection assay

LDH assays were performed using a combination of methodologies described in Goyal *et al.,* 2005 and Boothby *et al.,* 2017. Briefly, stock solutions of 25 mM Tris HCl (pH 7.0), 100 mM sodium phosphate (pH 6.0), and 2 mM pyruvate prepared in bulk and stored at room temperature prior to assay. In addition, 10 mM NADH was also prepared prior to the assay and then stored at 4 °C. L-Lactate Dehydrogenase (LDH) was sourced from Sigma (SKU #10127230001) and is supplied in ammonium sulfate at a pH of approximately 7. Prior to assay, LDH was diluted to a working concentration of 1 g/L. Experimental disaccharide-glycerol mixtures were formulated with LDH at a 1:10 (LDH:disaccharide-glycerol) ratio. Enough solution was prepared for three test excipient replicates and three control replicates each with a total volume of 50 μL. Each experimental and control mixture were aliquoted into a 1.5 mL microcentrifuge tube. The test mixtures were then vacuum desiccated for 16 hours with controls kept refrigerated at 4 °C. After vacuum desiccation, control and test excipient mixtures were brought to 250 μL total volume. Absorbance readings at 340 nm wavelength were taken every two seconds for 60 seconds with a quartz cuvette on a Thermo Scientific NanoDrop One^c^ (Thermo Scientific 840274200) spectrophotometer. A 100mM sodium phosphate and 2mM pyruvate solution was used as a blank. For control samples and experimental samples, 10 μL of control mixture or 10 μL of experimental sample mixture were combined with 10 μL of NADH and 980 μL of the 100mM sodium phosphate, 2mM pyruvate solution and then were measured.

This same procedure was used to measure the A340 of the experimental and control mixtures. A340 was plotted as a function of time and the slope of the linear portion of this plot calculated. A ratio of experimental over control slope, multiplied by one hundred, was taken to produce the percent protection of the experimental mixture. All raw LDH protection data is available in Supplemental File 8 while calculation of percent protection is available in Supplemental File 1.

### 5.6 Thermogravimetric analysis

Samples were run on a TA TGA5500 instrument in 100 μL platinum crucibles (TA 952018.906). Crucibles were tared prior to each run and prior to sample loading. Crucibles were loaded with between 5 mg and 10 mg of sample mixture. Each sample was heated from 30 °C to 220 °C at a 10 °C per minute ramp. All TGA data and thermograms are available in Supplemental File 2 and Supplemental File 3.

Determination of water loss was conducted using TA’s Trios software (TA Instruments TRIOS version #5.0.0.44608). Thermograms were used to calculate starting masses of samples and the mass of samples at the plateau that occurs after ∼100 °C but before the thermal denaturation at ∼200 °C. The Trios software “Smart Analysis” tool was used to identify the inflection point between these two mass loss events.

### 5.7 Differential scanning calorimetry

Samples were run on a TA DSC2500 instrument with Trios software (TA Instruments TRIOS version #5.0.0.44608). Analysis of DSC output was performed using Trios software. The heating run consisted of being equilibrated at -10 °C and then heated using a 10 °C per minute ramp to 220 °C. All raw DSC measurements are available in Supplemental File 4 and Supplemental File 5.

### 5.8 Fragility (m-index) calculation

Trios software (TA Instruments TRIOS version #5.0.0.44608) provided by TA Instruments was used to perform analysis of the DSC data. Calculations of glass former fragility (m-index) were performed based on equations 10 and 14 proposed by Crowley and Zografi [39]. On a thermogram with a completed heating ramp to 220 °C, the degradation peak, melt peak, and glass transition were identified. The Trios software built-in Onset and Endset analysis was used to determine the Glass transition onset and offset (endset). The software-identified glass transition onset and offset was used to calculate the m-index using Crowley and Zografi’s equations 10 where m is the alternative fragility parameter, ΔE_Tg_ is the activation enthalpy of structural relaxation at T_g_, R is the gas constant, T_g_ is the experimental glass transition temperature onset, and 14 where ΔE_η_ is the activation enthalpy for viscosity, R is the gas constant, T_g_ is the experimental glass transition onset temperature, T_g_ is the experimental glass transition offset temperature, and constant is an empirical constant of 5 [39]. A mean of each set of replicates was obtained. All fragility calculations are available in Supplemental File 6.

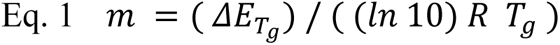

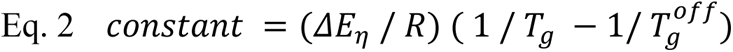

### 5.9 Selected organisms for in vivo assays

*Artemia franciscana*, *Caenorhabditis elegans,* and *Saccharomyces cerevisiae* were all selected based on availability of access to both desiccation sensitive and desiccation tolerant life stages. *Artemia franciscana* adults (#1) and cysts (#11) were acquired from the Northeast Brine Shrimp, LLC. *A. franciscana* adults were separated from culture media by tube-top filtration using a pluriSelect 200 μm pluriStrainer (#43-50200-03) prior to desiccation.

*Caenorhabditis elegans non-dauered daf-2* strains were cultured in S Medium [40] at 16 °C and fed with *E. coli* OP50. Dauered pre-conditioned daf-2 strain worms were cultured identically to non-dauered worms but when the initial aliquot of *E. coli* OP50 were consumed an additional five days of incubation at 25 °C was given to starve the culture and induce dauer arrest. Dauer worms were then placed on non-spotted agar plates in a 25 °C and 95% relative humidity atmosphere for 3 additional days to accomplish pre-conditioning. Both non-dauer and pre-conditioned dauer *C. elegans* sample sets were separated from media and residual OP50 food stock by tube-top filtration using a pluriSelect 200 μm pluriStrainer (#43-50200-03) prior to desiccation.

*Saccharomyces cerevisiae* strain BY4742 were cultured aerobically in YPD media [41] at 30 °C. After nine hours samples were taken representing the logarithmic growth stage yeast. The remaining culture was allowed to grow for an additional five days to ensure confluent growth in the stationary phase. Both logarithmic and stationary phase samples were pelleted by centrifugation, media was decanted, and a water wash of cell pellets was performed prior to desiccation.

### 5.10 Statistical methods

One-way ANOVA and Tukey post-hoc test was used for all pairwise comparisons. A p-value of less than 0.05 being one level of statistical significance (*), less than 0.01 being two levels of statistical significance (**), and less than 0.001 being three levels of statistical significance (***). All error values represent 95% confidence intervals.

Student’s T-Test was used to compare statistical differences between desiccation tolerant and desiccation sensitive life stages. A p-value of less than 0.05 being one level of statistical significance (*). All error values represent 95% confidence intervals.

### 6.1 Key resources table

| Reagent or resource | Source | Identifier |
| --- | --- | --- |
| D-Maltose monohydrate | Caisson | M004-500GM |
| D-Sucrose | Sigma-Aldrich | S0389-500G |
| D-Trehalose dihydrate | VWR | VWRB3599-1KG |
| Glycerol | Biobasic | GB0232 |
| Tris HCl | US Biological | 1185-53-1 |
| Sodium pyruvate | TCI | P0582 |
| NADH, Disodium Salt | Millipore Sigma | 481913 |
| L-Lactate Dehydrogenase | Sigma | SKU #10127230001 |
| 100 μL platinum crucibles | TA instruments | TA 952018.906 |
| DSC aluminum hermetic pan | TA instruments | TA 900793.901 |
| DSC aluminum hermetic lids | TA instruments | TA 901684.901 |
| 200 μm pluriStrainer | pluriSelect | #43-50200-03 |
| *A. franciscana* adults | Northeast Brine Shrimp LLC | #1 |
| *A. franciscana* cysts | Northeast Brine Shrimp LLC | #11 |
| *C. elegans daf-2* | Caenorhabditis Genetics Center | CB1370 |
| *E. coli* OP50 | Caenorhabditis Genetics Center | OP50 |
| *S. cerevisiae* BY4742 | ATCC | 201389 |

## 7.1 Results

### 7.2 Disaccharide-glycerol mixtures vitrify when dried

To begin to address which material properties correlate with enzyme-protective capacity in a vitreous state, we generated 18 different glass forming mixtures composed of one of three disaccharides (maltose, sucrose, or trehalose) and varying amounts of glycerol [10,42]. Previous studies have identified that small additions of glycerol changes the enzyme-protective capacity of trehalose. However, the material properties of these mixtures and how they correlate with changes in the level of protection have not been investigated. Furthermore, it is not known to what extent additions of glycerol will influence the material and enzyme-protective properties of other disaccharides or how these two properties are linked [10].

Maltose is a reducing disaccharide consisting of two glucose molecules joined by an ɑ (1→4) bond [43]. Sucrose is a non-reducing disaccharide formed by the glycosidic linkage between C1 of a glucose molecule to the C2 of a fructose molecule [44]. Trehalose is a non-reducing disaccharide formed through the (1-1) glycosidic linkage of two glucose molecules [45]. While trehalose and sucrose are better recognized mediators of desiccation tolerance, all three of these disaccharides are known to accumulate in a number of anhydrobiotic organisms during drying and in many cases they are known to be essential for surviving desiccation [6,28,30,31,33–35]. Maltose, though rarely seen as a desiccation-protectant in nature, is accumulated in some resurrection plants [6,28,30,31,33–35]. Furthermore, maltose has been shown to confer, short-but not long-term, desiccation tolerance in yeast, likely due to its propensity to be reduced [34]. Glycerol is a polyol that has been implicated in the tolerance of several stresses ranging from drying to freezing [36,38].

Mixtures containing 100%, 97.5%, 95%, 92.5%, 90%, and 87.5% of a single disaccharide (maltose, sucrose, or trehalose), combined with glycerol (weight % by weight % with glycerol) were created. Mixtures were dried overnight in a vacuum desiccator for 16 hours to produce glasses.

To ensure that these mixtures vitrified, rather than crystallized, each sample’s powder diffraction pattern was observed by powder X-ray diffraction (XRD) using Mo Kα radiation. The integrated plots of our 18 vitrifying samples contain a nearly identical broad absorption, and do not reveal any sharp peaks (Fig. S1a and S1b). The absence of sharp peaks indicates that the samples are glassy with no crystallinity [46,47]. Powder diffraction data was also measured for a desiccated sample of D-(+)-glucose, which is known to crystalize when dried. In contrast to the XRD patterns measured for the disaccharide-glycerol samples, the diffraction pattern for glucose exhibited a large number of distinct, closely spaced peaks due to the crystalline nature of this sample (Fig. S1a and S1b). These results indicate that maltose, sucrose, or trehalose by themselves or in conjunction with varying degrees of glycerol vitrify when dried under the drying regime used here (see Methods).

### 7.3 Disaccharide-glycerol mixtures have varying levels of enzyme-protection during desiccation

To address the question of which property(s) of a vitrified solid correlate with enzyme-protective capacity during desiccation, we assessed the ability of our 18 disaccharide-glycerol mixtures to protect the enzyme lactate dehydrogenase (LDH) during desiccation [7,48,49] (Fig. 1b). Previous reports have shown that LDH is sensitive to desiccation and that drying, and rehydration of this enzyme results in ∼95-99% loss in functionality [49] (Fig. 1b). Taking the ratio of the enzymatic activity of rehydrated *versus* control LDH, we observed that the protection of desiccated LDH varied significantly between different disaccharide-glycerol mixtures (Fig. 1c).

For maltose-glycerol mixtures, the highest level of protection was observed for our 100% maltose sample, while the lowest level of protection was conferred by the 87.5% maltose sample. No significant difference in protection was observed until the percentage of maltose in mixtures reached 92.5%, after which protection steadily decreased (Fig. 1c, top panel).

Similar to maltose-glycerol mixtures, sucrose-glycerol mixtures showed the highest level of protection at 100% sucrose, while the lowest level of protection was conferred by the 87.5% sucrose sample. However, unlike maltose samples, sucrose mixtures rapidly lost enzyme-protective capacity with statistically significant decreases in protection being observed upon the first (2.5%) addition of glycerol (Fig. 1b, middle panel).

Trehalose-glycerol mixtures differed from both maltose and sucrose mixtures in that the highest level of protection was achieved at 97.5% (Fig. 1c, bottom panel). Additionally, while mixing of maltose or sucrose with glycerol resulted in decreases in protection, additions of glycerol to trehalose were found to have a nonmonotonic relationship with protection (Fig. 1c, bottom panel).

These results demonstrate that different disaccharide and glycerol mixtures provide varying levels of protection to LDH during desiccation and rehydration, with trehalose responding in a non-monotonic fashion, maltose decreasing in enzyme-protective capacity in a linear fashion, and protection conferred by sucrose decreasing exponentially as a function of glycerol content.

### 7.4 Water content correlates with the enzyme-protective capacity of maltose-glycerol, but not trehalose-glycerol or sucrose-glycerol, mixtures

To begin to assess which properties of a vitrified system correlate with protection, we first assessed whether or not water content could account for these differences. To assess whether the differences in protection observed in our vitrified samples correspond to the amount of water they retain, we tested each of our mixtures using thermogravimetric analysis (TGA) (Fig. 2a and 2b). Water contents of our 18 dry mixtures ranged from 10.52 to 0.81% (Fig. 2c). Pure dry trehalose contained approximately 8% water content, while sucrose and maltose contained less and approximately the same water content (5-6%) (Fig. 2c). In maltose-glycerol mixtures the amount of retained water decreased with each addition of glycerol (Fig. 2c). In contrast, water content did not vary significantly in any sucrose-glycerol mixture. The trehalose-glycerol mixtures only demonstrate statistically significant decreases in water content after the addition of 5% glycerol (Fig. 2c).

**Figure 2:**
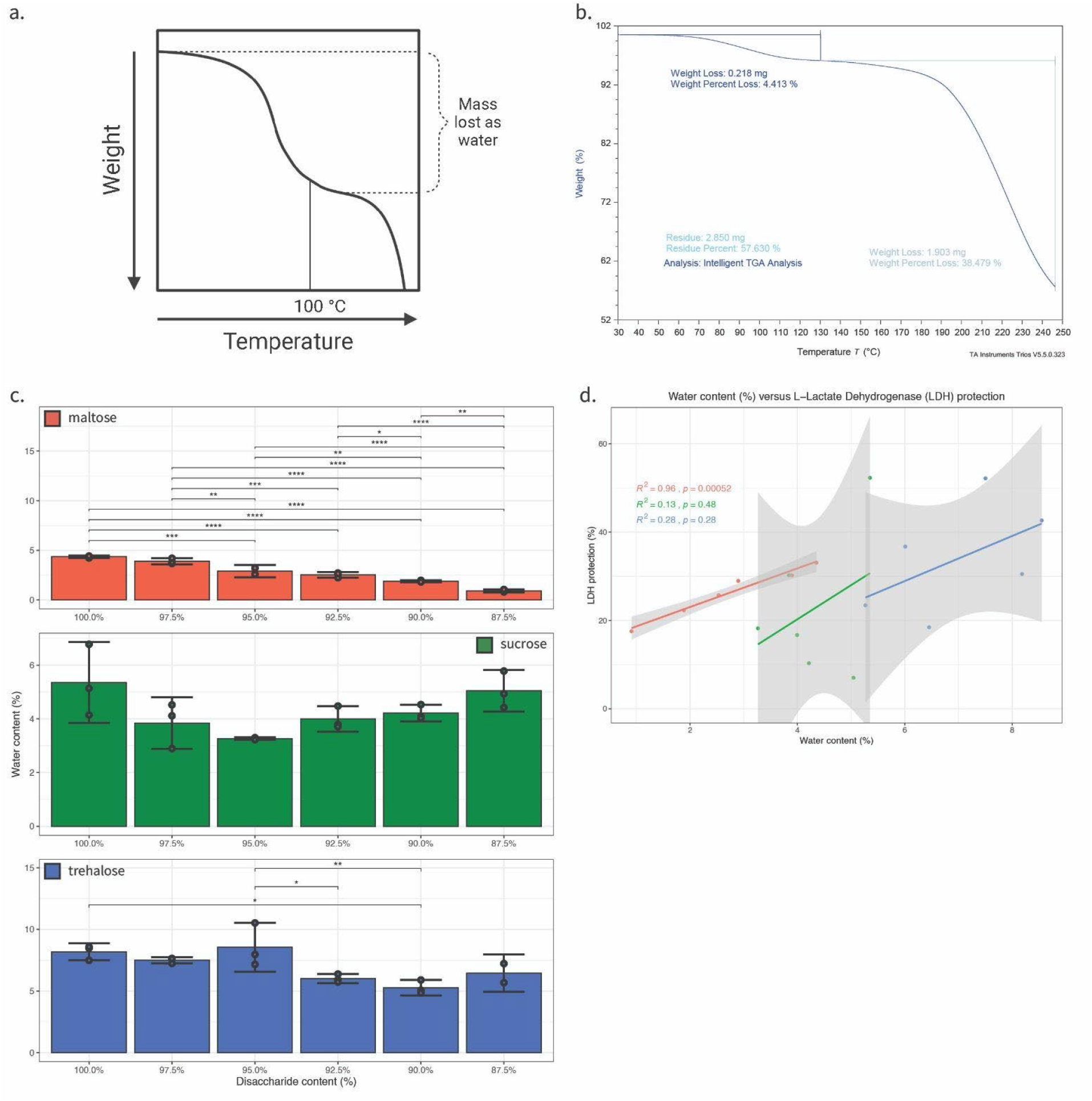
Water content correlates with the enzyme-protective capacity of maltose-glycerol but not sucrose- or trehalose-glycerol glasses. a) Schematic representation of an idealized TGA thermogram highlighting region of mass lost as water. b) Example TGA thermogram of a 100% trehalose-glycerol mixture. c) Water content values for disaccharide-glycerol mixtures organized by disaccharide. d) Correlation plot of water content versus protection for all disaccharide-glycerol mixtures (red = maltose, green = sucrose, blue = trehalose). Statistics calculated using a one-way ANOVA and Tukey post-hoc test: p-value ≤ 0.05 (*) p-value ≤ 0.01 (**), p-value ≤ 0.001 (***), pairwise relationships not shown are not significant, error bars represent 95% CI.

After observing distinct water retentive behaviors in dry disaccharide-glycerol mixtures, we assessed the relationship between water content and protection. For each sugar, there was a positive trend between enzyme-protection and increasing water content, however the correlation between these properties varied significantly between sugars. For maltose mixtures, this correlative analysis produced an R^2^ value of 0.96 (p-value = 0.00052) (Fig. 2d and S5a), an R^2^ value of 0.13 (p-value = 0.48) (for sucrose mixtures (Fig. 2d and S5b), and R^2^ value of 0.28 (p-value = 0.28) for trehalose mixtures (Fig. 2d and S5c). These results indicate that for maltose-glycerol systems the amount of retained water is a good indicator of the enzyme-protective capacity in the dry state. However, the amount of water in mixtures made of dry trehalose or sucrose and glycerol is a poor indicator of enzyme-protective capacity (Fig. 2d).

It should be acknowledged that precise determinations of water content using TGA can be complicated by an overlap between the offset of water evaporation and the onset of deterioration of a material. We found that for maltose and sucrose samples there was no such overlap, however in some cases trehalose samples showed an overlap requiring more refined methods of analysis. For example, when measuring the water content of some trehalose-glycerol mixtures, the step transitions calculated were numerous and ambiguously bordered the step transitions attributed to deterioration of a sample (Fig. S7b). In these cases, we made use of the derivative weight loss to differentiate between water loss and sample deterioration (Fig. S7a and S7b). This may result in less accurate determinations of water content for very complex thermograms. However, the behavior of most samples, especially the maltose- and sucrose-glycerol mixtures, were much more straightforward and allowed for simple differentiation of the water-loss and deterioration processes (Fig. S7a). Adding validity to our determination of water contents, even in complex samples, was the alignment of glass transition temperatures and water contents from previously published work [14].

### 7.5 An increase in the glass transition temperature correlates with enzyme protection conferred by maltose-glycerol and to a lesser extend sucrose-glycerol and trehalose-glycerol mixtures in the dry state

After observing that water retention is a poor indicator of the enzyme-protective capacity of trehalose- and sucrose-glycerol glasses, we wondered if glass transition temperature (*T_g_*) might be a property that correlates with the stabilizing effects of these sugars. Previous work has established that even small amounts of additives, such as glycerol, can lead to decreased or increased *T_g_* of a vitrifying material (Fig. 3a and 3b) [50–55]. With this in mind, we were curious if different additions of glycerol to our disaccharides would serve to increase or decrease the *T_g_* resulting glasses, and whether or not these changes in *T_g_* correlates with the enzyme-protective capacity of our mixtures.

**Figure 3:**
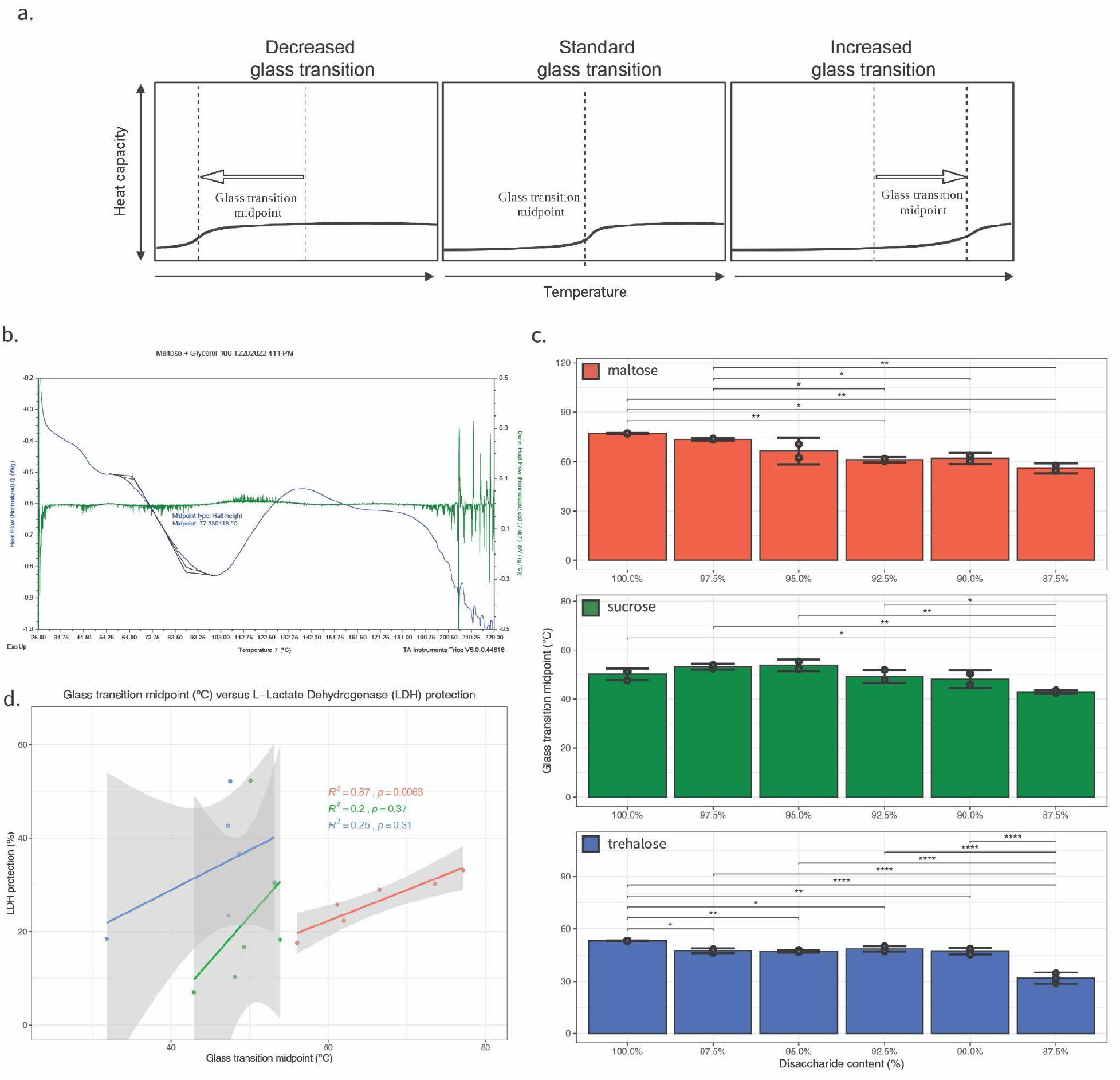
Anti-plasticization of maltose- and sucrose-, but not trehalose-glycerol glasses correlate with enzyme protection. a) Schematic representation of an idealized DSC thermogram illustrating a decrease or increase in glass transition midpoint. b) Example DSC thermogram of a 100% trehalose-glycerol mixture. c) Glass transition midpoint values for disaccharide-glycerol mixtures organized by disaccharide. d) Correlation plot of glass transition midpoint versus protection for all disaccharide-glycerol mixtures (red = maltose, green = sucrose, blue = trehalose). Statistics calculated using a one-way ANOVA and Tukey post-hoc test: p-value ≤ 0.05 (*) p-value ≤ 0.01 (**), p-value ≤ 0.001 (***), pairwise relationships not shown are not significant, error bars represent 95% CI.

The glass transition temperature onset and offset, the temperatures that the material starts and stops undergoing a change from a glassy state to a rubbery state, were assessed using differential scanning calorimetry (DSC), and the glass transition midpoint temperature calculated by taking the mean of the onset and offset temperature (Supplemental File 6). Figures 3c, S3a, and S3b show the average *T_g_* onset, offset, and midpoint for each of our mixtures. Here one can see that when considering the glass transition midpoint even small additions of glycerol, starting at 2.5%, act to decrease the *T_g_* for trehalose, while decreasing the *T_g_* of maltose began at 7.5% glycerol and a decrease in sucrose is only observed with an even larger (12.5%) addition of glycerol (Fig. 3c). For all disaccharide-glycerol mixtures, glass transition midpoint temperatures are positively correlated with enzyme protective capacity, but none of the mixtures are significantly correlated (Fig. 3d).

However, when considering the onset or offset glass transition temperature for each set of disaccharide-glycerol mixtures, we saw different behaviors appear. For example, when observing changes to the glass transition onset temperature for the maltose-glycerol mixtures, even after the addition of 12.5% glycerol there was no significant change (Fig. S3a). As opposed to maltose, sucrose saw a significant decrease in glass transition onset temperature after the addition of 12.5% glycerol (Fig. S3a). Finally, the trehalose-glycerol mixtures show statistically significant changes in the glass transition onset temperature beginning with the first addition of glycerol (Fig. S3a). The trehalose-glycerol mixtures also showed a statistically significant decrease after the addition of 12.5% glycerol (Fig. S3a).

When observing the offset of glass transition temperature for each of the disaccharide-glycerol mixtures, we observed that the maltose-glycerol and sucrose-glycerol mixtures demonstrated similar behavior to the midpoint observations. Specifically, starting at 5% for maltose and 12.5% for sucrose, glycerol acts to decrease the glass transition offset temperature (Fig. S3c). On the other hand, trehalose does not show a statistically significant decrease in the glass transition offset until after the addition of 10% glycerol (Fig. S3c).

Next, we evaluated the relationship between enzyme-protective capacity and the onset, offset, and midpoint *T_g_* for each mixture (Fig. 3d, S4, S5d, S5e, and S5f). This correlative analysis produced positive correlations with R^2^ values ranging from 0.87 to 0.81 for maltose (Fig. 3d, S4, S5d), positive correlations with R^2^ values ranging from 0.38 to 0.1 for sucrose (Fig. 3d, S4, S5e), and positive correlations for trehalose the R^2^ values ranged from 0.36 to 0.1 (Fig. 3d, S4, and S5f). We observe that the enzyme-protective capacity of the maltose-glycerol mixtures are influenced, but not significantly, by variation in their *T_g_* and that the enzyme-protective capacity of the sucrose- and trehalose-glycerol mixtures are moderately influenced by variation in their *T_g_* (Fig. 3d).

Finally, we examined the relationship between the enzyme-protective effect of our sugar mixtures and the difference between the temperature at which LDH is protected (*T_exp_* = 22 °C) their *T_g_*. We reasoned that for a mixture with a *T_g_* close to *T_exp_* protection might be lowered due to the mixture undergoing a relaxation.

Consistent with this reasoning, for all disaccharide-glycerol mixtures enzyme-protection and *T_exp_*-*T_g_* had a negative correlation (S6). However, only the maltose mixtures approached, but was not, a significant correlation between enzyme-protection and *T_exp_*-*T_g_* (S6). This indicates that at the temperature used in this study, relaxation of dry sugar mixtures due to a similarity in *T_exp_* and *T_g_* is not a significant influence on enzyme-protection.

Our results demonstrate that generally, protection positively correlates with increasing *T_g_*. However, this correlation is clearly stronger for some sugar-glasses (e.g., maltose) compared to others (e.g., sucrose).

### 7.6 Vitrified maltose-glycerol and trehalose-glycerol mixtures demonstrate a reduction in glass former fragility that is correlated with enzyme-protection

Next, we empirically determined the glass former fragility (m-index; [25,39] of our disaccharide-glycerol mixtures (Fig. 4a). Moynihan *et. al.* described the use of thermal methods to explore the relationship between the width of a glass transition and the activation enthalpy for viscosity [56]. Crowley and Zografi later applied the assumption that the activation enthalpy for viscosity is equivalent to the activation enthalpy of structural relaxation at T_g_ [39]. Utilizing this method glass former fragility (m-index) was calculated from DSC thermogram outputs (Fig. 4a and 4b) [39]. The fragility of glass forming solutions composed from pure disaccharides was lowest for maltose, followed by sucrose, and finally trehalose (Fig. 4c). For maltose samples, glass former fragility steadily increased with each addition of glycerol, but did not vary significantly (Fig. 4c). However, for both sucrose and trehalose small additions of glycerol (up to 5 and 2.5%, respectively) decreased glass former fragility (Fig. 4c). However, greater additions of glycerol to both trehalose and sucrose increased glass former fragility (Fig. 4c). For sucrose, variations in glass former fragility were not significant. On the other hand, after the addition of 10% glycerol, trehalose saw significant increases in glass former fragility (Fig. 4c).

**Figure 4:**
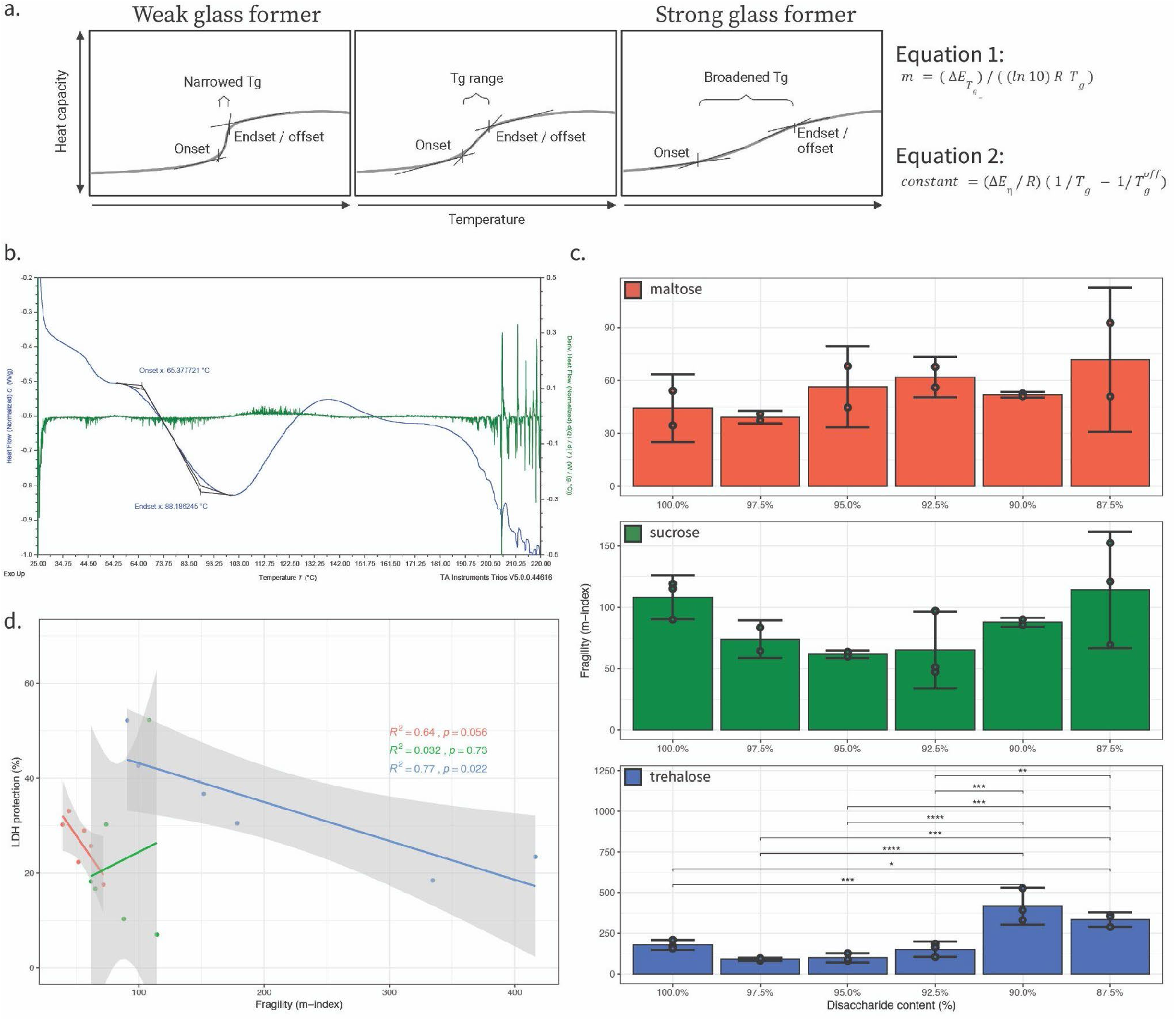
Reduced glass forming fragility of maltose- and trehalose-, but not sucrose-glycerol glasses correlates with enzyme protection. a) Schematic representation of idealized DSC thermograms illustrating a narrowing or broadening of the glass transition and the two equations used to calculate m-index. Equation 1 where m is the alternative fragility parameter, ΔE_Tg_ is the activation enthalpy of structural relaxation at T_g_, R is the gas constant, T_g_ is the experimental glass transition temperature onset, and 2 where ΔE_η_ is the activation enthalpy for viscosity, R is the gas constant, T_g_ is the experimental glass transition onset temperature, T_g_^off^ is the experimental glass transition offset temperature, and constant is an empirical constant of 5. Equation 1 and 2 in our manuscript are derived from equations 10 and 14, respectively from [39]. Please note that Equation 14 is equivalent to equation 7 from Moynihan *et. al.* [57] b.) Example thermogram of 100% trehalose with glass transition onset, glass transition offset, and 1st derivative with respect to temperature (green) shown. c) Glass former fragility (m-index) values for disaccharide-glycerol mixtures organized by disaccharide. d) Correlation plot of glass former fragility (m-index) versus protection for all disaccharide-glycerol mixtures (red = maltose, green = sucrose, blue = trehalose). Statistics calculated using a one-way ANOVA and Tukey post-hoc test: p-value ≤ 0.05 (*) p-value ≤ 0.01 (**), p-value ≤ 0.001 (***), pairwise relationships not shown are not significant, error bars represent 95% CI.

Next, we assessed the relationship between the fragility of our glass forming mixtures and protection of LDH. For maltose and trehalose, the trend between enzyme-protection and glass former fragility was negative, while for sucrose this trend was slightly positive. As observed previously, the strength of these trends varied dramatically between sugars, as this correlative analysis produced an R^2^ value of 0.64 for maltose (Fig. 4d and S5g), an R^2^ value of 0.032 for sucrose (Fig. 4d and S5h), and R^2^ value of 0.77 for trehalose (Fig. 4d and S5i). This indicates that for maltose and trehalose, observed variation in glass former fragility can explain some of the enzyme-protective capacity of the desiccated mixtures (Fig. 4d).

It was observed that the glass forming fragility of 100% sucrose was potentially behaving differently from mixtures containing glycerol additions. It was supposed that this datapoint might be ‘dragging’ the relationship between the glass former fragility of our glass forming mixtures and protection of LDH upward into a positive correlation. A correlation was calculated excluding the pure (100%) disaccharide samples from the mixtures (97.5-87.5%). This exclusion of the pure disaccharide samples greatly improved the correlation value of the sucrose mixtures, from an R^2^ value of 0.032 to an R^2^ value of 0.41 (Fig. S8c and S8d). Interestingly, the correlation for the trehalose-glycerol was also improved by excluding the 100% disaccharide samples, from an R^2^ value of 0.77 to an R^2^ value of 0.83 (Fig. S8e and S8f). However, the correlation for the maltose-glycerol mixtures was made worse, from an R^2^ value of 0.64 to an R^2^ value of 0.57 (Fig. S8a and S8b). These results suggest that while pure disaccharides may behave in substantively different ways than disaccharide-glycerol mixtures, this is not always the case.

### 7.7 Increased T_g_ and decreased glass former fragility are characteristics of anhydrobiotic life-stages

We wondered if the correlation between material properties, such as minimal glass former fragility or anti-platicization, and enzyme-protection that we observed *in vitro* appeared to carry over to organismal systems. Here we considered reduced glass former fragility and anti-plasticization, but not water retention, since water content has previously been observed to influence glass-like properties [26,58–60]. This influence is observed in our samples, where for example in maltose-glycerol mixtures, we see a very strong correlation (R^2^ = 0.989) between water content and increased *Tg* (Fig. S2a).

We measured the *T_g_* and m-indexes of three desiccation tolerant organisms when in a desiccation sensitive life stage or a desiccation tolerant life stage. The three desiccation tolerant organisms we selected for this study were: the brine shrimp *Artemia franciscana*, the nematode worm *Caenorhabditis elegans,* and yeast *Saccharomyces cerevisiae* (Fig. 5a). Each of the selected organisms is known to accumulate protective disaccharides and polyols during drying [34,36,61–63].

**Figure 5:**
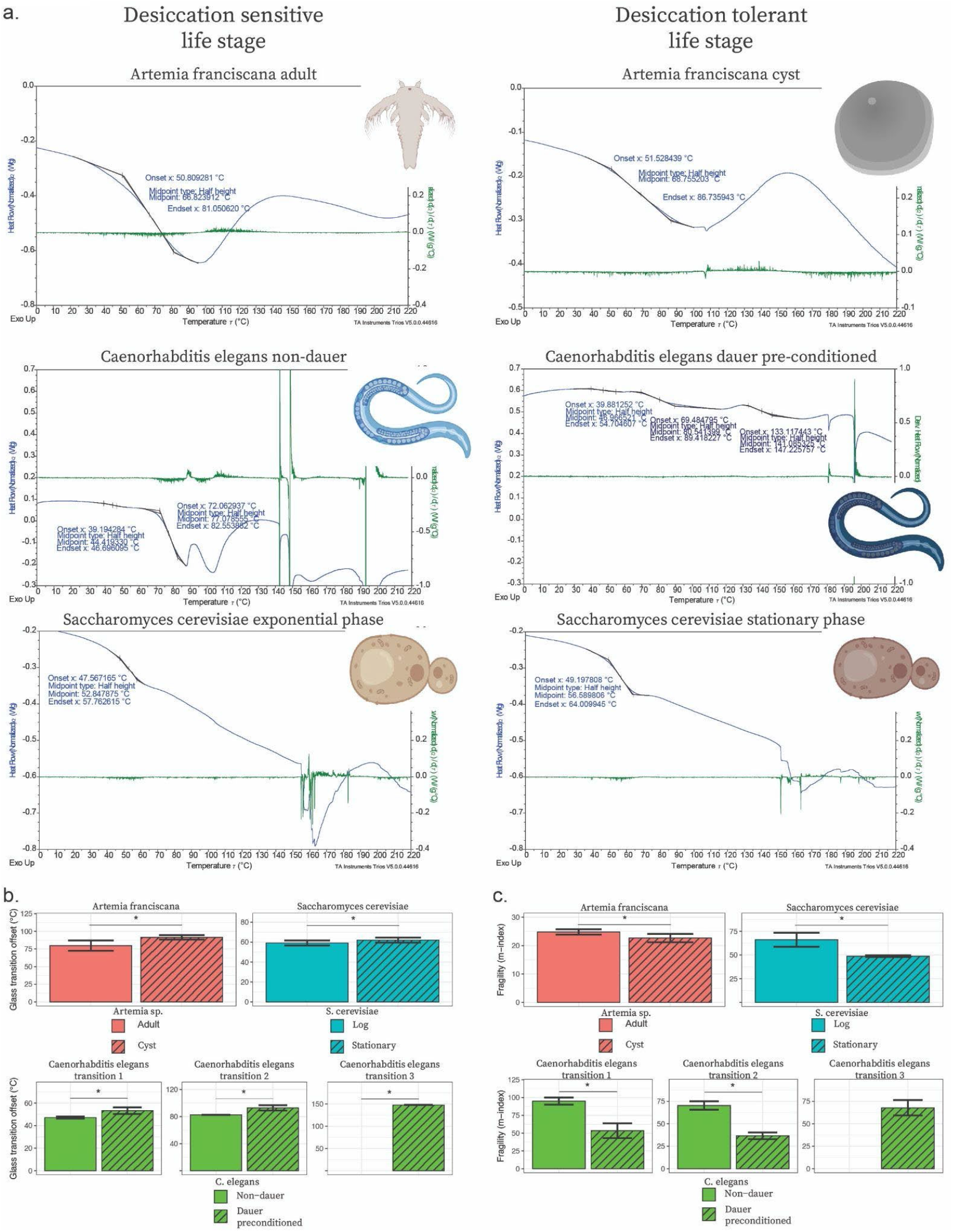
*In vivo* glass transition temperature is increased, and glass former fragility reduced in desiccation-tolerant *versus* sensitive life stages of diverse anhydrobiotic organisms. a) Selected organisms for *in vivo* assays with example DSC thermogram outputs. b) Glass transition offset values for selected desiccation tolerant organisms organized by organism and life stage. c) Glass former fragility (m-index) values for selected desiccation tolerant organisms organized by organism and life stage. Statistics calculated using T-test: p-value ≤< 0.05 (*), pairwise relationships not shown are not significant, error bars represent 95% CI.

For *A. franciscana*, there was a significant increase in *T_g_* offset between the adult (desiccation-sensitive life stage) and cyst (desiccation-tolerant life stage) (Fig. 5b). For *S. cerevisiae*, we saw a statistical increase between the exponential phase (desiccation sensitive life stage) and stationary phase (desiccation tolerant life stage) when measuring *T_g_* offset (Fig. 5b). And finally, for *C. elegans,* we observed complex thermograms containing more than one obvious glass transition. Specifically, we observed two glass transitions on the thermograms of non-dauer worms and three glass transitions on the thermograms of dauer pre-conditioned worms. The two glass transitions from non-dauer worms were paired with the first two glass transitions from the dauer pre-conditioned worms based on similarities in glass transition temperature. The trend of desiccation tolerant life stages possessing a significantly higher *T_g_* offset in comparison to the desiccation sensitive life stage continues across all three ranges of potential glass transitions for the *C. elegans* non-dauer and dauer pre-conditioned worms (Fig. 5b).

Thus, for all three of the desiccation tolerant organisms tested we observe significantly increased *T_g_* in the anhydrobiotic state, indicating that increased *T_g_* is a hallmark of some desiccation-tolerant organismal systems.

Next, we evaluated our three selected organisms’ life stages for changes in m-index (glass former fragility). For *A. franciscana,* we saw significant decrease between the adult (desiccation sensitive life stage) and cyst (desiccation tolerant life stage) when measuring m-index (Fig. 5c). For *S. cerevisiae*, we also saw a significant decrease between the exponential phase (desiccation-sensitive life stage) and stationary phase (desiccation-tolerant life stage) when measuring m-index (Fig. 5c). For *C. elegans*, we saw a significant decrease in glass former fragility between the non-dauer (desiccation sensitive life stage) and dauer pre-conditioned (desiccation tolerant life stage) across all three glass transitions.

Taken together, these results indicate that while *in vitro* simple mixtures of protectants vary widely in what properties correlate with protection in the vitrified state, *in vivo* both increased glass transition temperature and decreased glass former fragility are good indicators of survival in the dry state.

## 8.1 Discussion and conclusions

Since its conception, the vitrification hypothesis has provided a compelling possible explanation as to how anhydrobiotic organisms preserve their cells and cellular components during desiccation [1,5–7]. However, while vitrification is considered necessary for desiccation tolerance it is not sufficient [5,9]. This implies that there is some property of a glassy material that makes it more, or less, protective.

Here we have quantified the enzyme-protective capacity and material properties of 18 different vitrified systems, each composed of one of three different disaccharides (maltose, sucrose, and trehalose) formulated with varying amounts of glycerol (0-12.5%). We find that both enzyme-protective capacity and material properties of disaccharide-glycerol mixtures are modulated differently depending on the disaccharide used. Consistent with this, water retention (maltose), increased T_g_ (sucrose), and reduced glass fragility (trehalose) each correlated best with protection for a particular disaccharide. Interestingly, reduced glass former fragility and increased glass transition (anti-plasticization) is observed in desiccation-tolerant life stages of diverse organisms. Thus, individual protective properties observed for reductive enzyme systems appear to be used in combination *in vivo*.

### 8.2 Water retention, mechanisms of protection involving water, and water’s effects on glass transition temperature and glass former fragility

Our results demonstrate that different disaccharide-glycerol mixtures contain different quantities of water. Furthermore, the relationship between water content and protection also varies between disaccharide-glycerol mixtures. Maltose-glycerol mixtures display a significant positive correlation between water content and enzyme protection, while the sucrose-glycerol and trehalose-glycerol mixtures show only mildly positive correlations (R^2^ =0.13 and 0.28, respectively).

The water content of a glassy material could affect enzyme-protection through several mechanisms. Loss of water during dehydration can lead to a loss of important, stabilizing hydrogen bonds, which help to maintain protein folding. While theories, such as the water replacement hypothesis, which propose mechanisms by which this loss of this hydrogen bond next work can be dealt with, others such as the water entrapment [13–17], preferential exclusion/hydration hypothesis [16,18], or anchorage hypothesis [19–22] offer up mechanisms by which residual water can be utilized to provide protection even at low levels.

The water entrapment hypothesis posits that a protectant which has a strong affinity for water but is preferentially excluded from client molecules could help to coordinate small amounts of residual water into proximity with desiccation-sensitive material (e.g., proteins, membrane, etc.). The effect would be to entrap and increase the local concentration of water around these desiccation-sensitive molecules which could help to maintain the hydrogen bond network needed for integrity. Conversely, the preferential exclusion hypothesis posits that a protectant which preferentially interacts with itself, to the exclusion of both water and client molecules, could in effect act as a space filling molecule. In this capacity, the protectant would reduce the overall accessible volume within the cell increasing the effective concentration of water and forcing water molecules into proximity with desiccation-sensitive molecules. Finally, the anchorage hypothesis posits that client molecules interact with the water-protectant matrix, and this interaction reduces the likelihood of protein unfolded since unfolding would have to lead to a recording of the water-protectant matrix.

Beyond water serving directly in the stabilization of biomolecules via the formation of a hydrogen bond network, water can also serve as an important plasticizing agent of biological and hydrophilic materials. This means that increasing water content in a vitrified material typically leads to a decrease in *T_g_*, which is also considered to reduce protection. However, while this is generally true, there have been reports of *bona fide* anti-plasticization effects of water [59]. Here we observe that water content of maltose-, sucrose-, and trehalose-glycerol glasses has varying degrees of a strong plasticizing effect on the glass transition. The only negative correlation between water content and decreased *T_g_* was observed for the sucrose-glycerol mixtures (R^2^ = 0.44, Fig. S2c). In trehalose-glycerol mixtures, there was essentially no correlation between water content and decreased *T_g_* (R^2^ = 0.088, Fig. S2e). Finally, in maltose-glycerol mixtures, rather than seeing water correlate with decreased *T_g_*, surprisingly we observe a strong correlation between water content and increased *T_g_* (R^2^ = 0.94, Fig. S2a). This again showcases how each of the disaccharides, when in a desiccated disaccharide-glycerol mixture, displays disparate changes in material properties.

When instead considering the relationship between water content and glass former fragility, we again observe differing results by disaccharide-glycerol mixture. Here we see that the desiccated maltose-glycerol mixtures have a strong negatively correlated relationship (R^2^ = 0.74, Fig. S2b), sucrose has a strong positively correlated relationship (R^2^ = 0.88, Fig. S2d), and trehalose has a weak positively correlated relationship (R^2^ = 0.29, Fig. S2f).

These results might lead one to believe that at the water contents examined here (< 11%) water retention might be a potential predictor of enzyme protection capacity. However, water content also seems to influence other material properties of the vitrified system in a non-stereotyped fashion. For example, increasing water content in maltose-glycerol mixtures strongly correlates with reduced glass former fragility, while in sucrose and trehalose-glycerol mixtures increasing water content increases glass former fragility. Thus, we conclude that water content itself is not a good predictor of desiccation tolerance nor of other properties of a vitrified system.

### 8.3 The enzyme-protective capacity of sucrose-glycerol mixtures was most influenced by increases and decreases in glass transition temperature

Glass transition temperature (*T_g_*) is the temperature at which a hard glassy material will begin to transition into a rubbery solid [64]. Figure 3a is a schematic illustration of how increases or decreases in the *T_g_* are captured and visualized on a thermogram. The relationship of increases or decreases in *T_g_* with protection varied between different disaccharide-glycerol mixtures but was most important for the sucrose-glycerol mixtures. While we observed that only maltose-glycerol mixtures had a strong relationship between increased *T_g_* and protection (Fig. 3d), in the sucrose-glycerol mixtures we see that addition of glycerol results in an increase in the *T_g_* when added up to 5%, but then further addition of glycerol causes a significant decrease in the *T_g_*. Furthermore, the influence of glycerol on the *T_g_* of sucrose-glycerol mixtures correlates weakly with protection but is the highest correlation with respect to enzyme-protection (Fig. 2d, 3d, and 4d). Finally, in the trehalose-glycerol mixtures we see at first a significant decrease coinciding with the first addition of glycerol (2.5%) and then no significant variation in the glass transition midpoint until the addition of significantly more (12.25%) glycerol where it decreases the *T_g_*, and the small changes in *T_g_* that are observed show no correlation with protection.

Thus, while increases in *T_g_* are predictive of the protection conferred by maltose-glycerol mixtures, and to a lesser extent sucrose-glycerol, this predictive capacity does not extend to all vitrified systems.

### 8.4 The difference between the experimental temperature of enzyme protection and glass transition temperature did not correlate with protection

The closer a glassy protectant is to its *T_g_*, the more molecular motion should be introduced, which could result in a loss of protective capacity. However, we observed that the difference in the experimental temperature at which LDH assays were conducted and the *T_exp_-T_g_* of a protective glass did not correlate significantly with enzyme-protective capacity (S6). This does not mean that as a protective glass approaches its *T_g_* that it does not lose enzyme-protective capacity, since it is possible that the experimental temperatures used here were still sufficiently lower than *T_g_* to negatively impact enzyme-protection. Further studies where the experimental temperature is brought much closer to *T_g_* could be insightful in this regard.

### 8.5 The relationship between T_g_ and protection varies dramatically depending on whether onset, midpoint, or offset glass transition temperatures are considered

When referring to the glass transition or the material properties based on the glass transition, the standard measure is to use the glass transition midpoint. This value is inherently influenced by both the glass transition onset and offset temperatures (Fig. 3c, S3a, and S3b) [65]. The glass transition offset temperature is representative of the point at which the ‘glassy’ nature of a vitreous system is finally overcome (Fig. S3b). By contrast the glass transition onset is only representative of the start of the transition of a ‘glassy’ state to a rubber-like solid (Fig. S3a).

In evaluating the relationship between *T_g_* and protection, we observed a dramatic variation in this correlation depending on whether we used the onset, midpoint, or offset glass transition temperature (Fig. S4). Here we have reported midpoint *T_g_* as is convention, but also have included correlations between protection and onset/offset *T_g_* (Fig. S4).

### 8.6 Disaccharide-glycerol mixtures with similar concentrations produce fragilities of differing glass former fragility (m-index)

The fragility of different glass former mixtures varied by disaccharide. Only maltose and trehalose showed evidence of a relationship between glass former fragility and enzyme protection capacity. Interestingly, when comparing different disaccharide-glycerol systems that provide similar enzyme-protective capacity, those mixtures did not necessarily produce glasses with similar glass former fragility (m-index) measurements. For example, when comparing the 97.5% maltose, 97.5% sucrose, and 92.5% trehalose disaccharide-glycerol mixtures that provide approximately 30% enzyme protection during desiccation (30.21%, 30.26, and 36.72% respectively), those mixtures produced had a glass former fragility (m-index) of 39.15, 74.01, and 151.81 respectively (Table 1). In addition, when comparing the most enzyme-protective mixtures for each disaccharide-glycerol mixture, 100% maltose, 100% sucrose and 97.5% trehalose, only trehalose produced the lowest glass former fragility (m-index) measurement. This indicates that each disaccharide-glycerol system produced glass former fragility patterns that are only comparable within that system and not between different disaccharide-glycerol systems. Again, just as with water retention and shifts in the *T_g_*, we see that the addition of glycerol induces different degrees of glass former fragility and that this property is a good indicator of protection for some sugar glasses, but not for others.

**Table 1:**
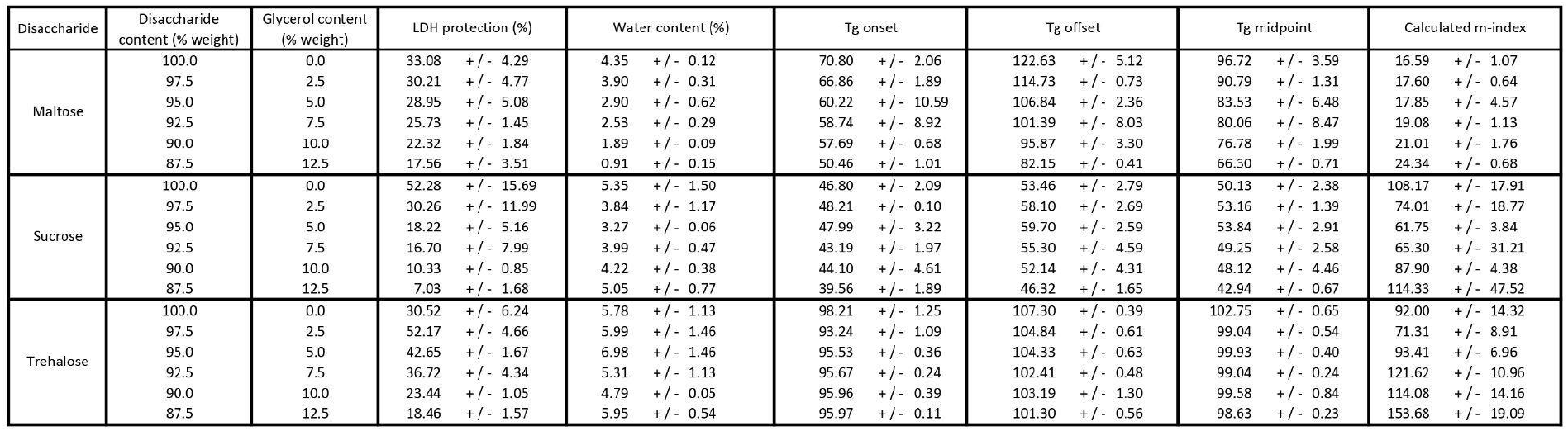
*In vitro* measurement results organized by disaccharide-glycerol content and by material property, error values represent 95% CI.

### 8.7 Both change in T_g_ and glass former fragility are hallmarks of organismal desiccation tolerance

The results from our *in vitro* experiments, while novel on their own, beg the question of whether these findings apply to whole desiccation tolerant organisms. Examining three different organisms in both a desiccation-tolerant and -sensitive life stage we observe that both increased *T_g_* and reduced m-index (glass former fragility) are hallmarks of successful anhydrobiosis. Specifically, when considering changes in *T_g_*, we see that for each organism, *A. franciscana*, *C. elegans,* and *S. cerevisiae,* there is a statistically significant increase in *T_g_* when comparing the desiccation-sensitive life stage to the desiccation-tolerant life stage. When considering the impact of glass former fragility, we yet again see that for each organism, *A. franciscana*, *C. elegans,* and *S. cerevisiae,* there is a statistically significant decrease in glass former fragility when comparing the desiccation-tolerant life stage to the desiccation-sensitive life stage.

Figure 6 provides an overview of what we envision is occurring to enable changes in the *T_g_* (Fig. 6a) and glass former fragility (Fig. 6b and 6c) to be protective during desiccation. In Figure 6a we depict a glass with a decreased *T_g_* as one with relatively few and/or weak bonds, which do not effectively slow down molecular motions leading to the destabilization and/or aggregation of embedded clients over time. In contrast, a glass with an increased *T_g_* is one with increased and/or stronger bonds leading to reduction in molecular motion and an increase in stability of a client over time. This is in line with previous suggestions that inducing a super viscous (glassy) state is stabilizing because to unfold a protein would need to displace the embedding media [5,14]. In this light, increased *T_g_* would be protective as materials will become more fluid as the temperature they are stored at approaches *T_g_*. Figure 6b illustrates what we envision is happening when a glass forming mixture forms a weak glass. As water is lost, the weak glass forming ability of the mixture does not become viscous (green) soon enough to prevent drying induced damage (tan). Specifically, a weak glass former material will only begin gaining sufficient viscosity to confer protection after significant water loss. Conversely, Figure 6c illustrates how a strong glass forming mixture starts to gain viscosity (purple) much earlier in the desiccation process compared to weaker glass forming materials. This steady increase in viscosity, allows for the slowing detrimental perturbations (tan) that manifest early during drying. This model implies to some degree that it is the drying process, in addition to being in a dry state, that must be protected against.

**Figure 6:**
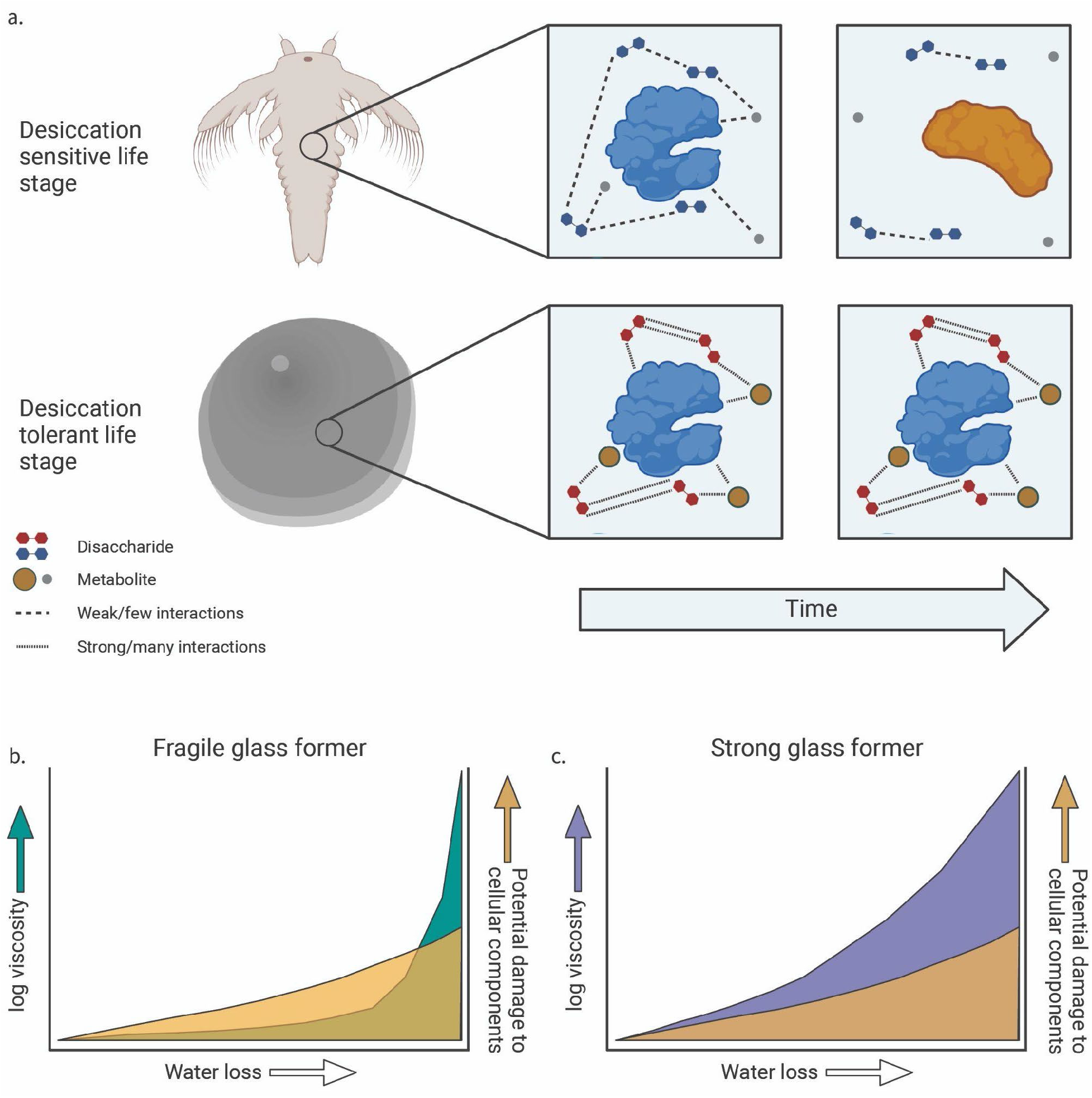
Model of protection conferred by increased glass transition temperature and reduced glass former fragility. a) Schematic representation of the potential mechanism of protection conferred by increasing glass transition temperature. b) Schematic representation of the potential mechanism underlying how damage is accrued in fragile glass former mixtures. c) Schematic of the potential mechanism underlying how damage is prevented in strong glass former systems.

While previous studies have examined the glass former fragility of seeds in relationship to their desiccation tolerance, to our knowledge this is the first study examining changes in *T_g_* and glass former fragility in animal and fungal systems [3,9,66–69]. These comparative organismal studies show a stark contrast to our *in vitro* data, in that rather than a single material property correlating with protection, it appears that in living anhydrobiotic systems both increased *T_g_* and reduced glass former fragility are generally increased. This may be due the nature of the *in vitro* systems being simple, or less complex, in their interactions while the *in vivo* studies are by their nature much more complex, both in their material makeup and interactions.

These results hint at the fact that living systems likely make use of multiple mediators of desiccation tolerance to produce protective glasses. Indeed, an emerging paradigm in the anhydrobiosis field is that beyond disaccharides, other molecules, such as intrinsically disordered proteins play vital roles in preserving biological function in the solid state.

Our study advances our understanding of what properties of a vitrified system promote desiccation tolerance and the phenomenon of anhydrobiosis both *in vitro* and *in vivo*. A deeper understanding of natural desiccation tolerance promises to provide avenues for pursuing real world applications such as biobanking of seeds and tissues, stabilization of pharmaceuticals, and the generation of stress tolerant crops.

## 9.1 Funding sources and acknowledgments

### 9.2 Funding sources

This work was primarily supported by NSF DBI grant # 2213983. This work was partially funded through a fellowship to JFR administered by Wyoming NASA EPSCoR (NASA Grant #80NSSC19M0061). JFR is supported by the USDA National Institute of Food and Agriculture, Hatch project #1012152. The authors gratefully acknowledge financial support from the NSF (CHE 0619920 to NA and IntBio 2128069 to TCB) and the Institutional Development Award (IDeA) from the National Institute of General Medical Sciences of the National Institutes of Health (Grant # 2P20GM103432). Any opinions, findings, conclusions, or recommendations expressed in this publication are those of the author(s) and do not necessarily reflect the view of the National Institute of Food and Agriculture (NIFA) or the United States Department of Agriculture (USDA).

## 9.3 Acknowledgments

Caenorhabditis elegans daf-2 strains were donated by the Fay Laboratory at the University of Wyoming, department of Molecular biology. Saccharomyces cerevisiae strain BY4742 was donated by Dr. Peter Thorsness Emeritus Professor at the University of Wyoming, department of Molecular Biology. We thank members of the Water and Life Interface Institute (WALII), supported by NSF DBI grant # 2213983, in particular Christina Walters, for helpful discussions.The authors are thankful to Dr. Silvia Sanchez Martinez and members of the Boothby Lab for discussions and reading of this manuscript.

## 10.1 Author contributions

**John F. Ramirez:** Conceptualization, Methodology, Investigation, Formal analysis, Data curation, Writing - Original Draft, Writing - Review and Editing, Visualization. **U.G.V.S.S. Kumara:** Methodology, Investigation, Data Curation, Writing - Review and Editing. **Navamoney Arulsamy:** Investigation, Writing - Original Draft, Writing - Review and Editing. **Thomas C. Boothby:** Conceptualization, Methodology, Writing - Original Draft, Writing - Review and Editing, Supervision, Visualization, Project administration, Funding acquisition.

## 11.1 Data availability statement

All raw data used in this paper is supplied in supplemental files.

## 12.1 Additional information

The authors declare no competing interests.

## 13.1 Supplemental figures

**Figure S1:**
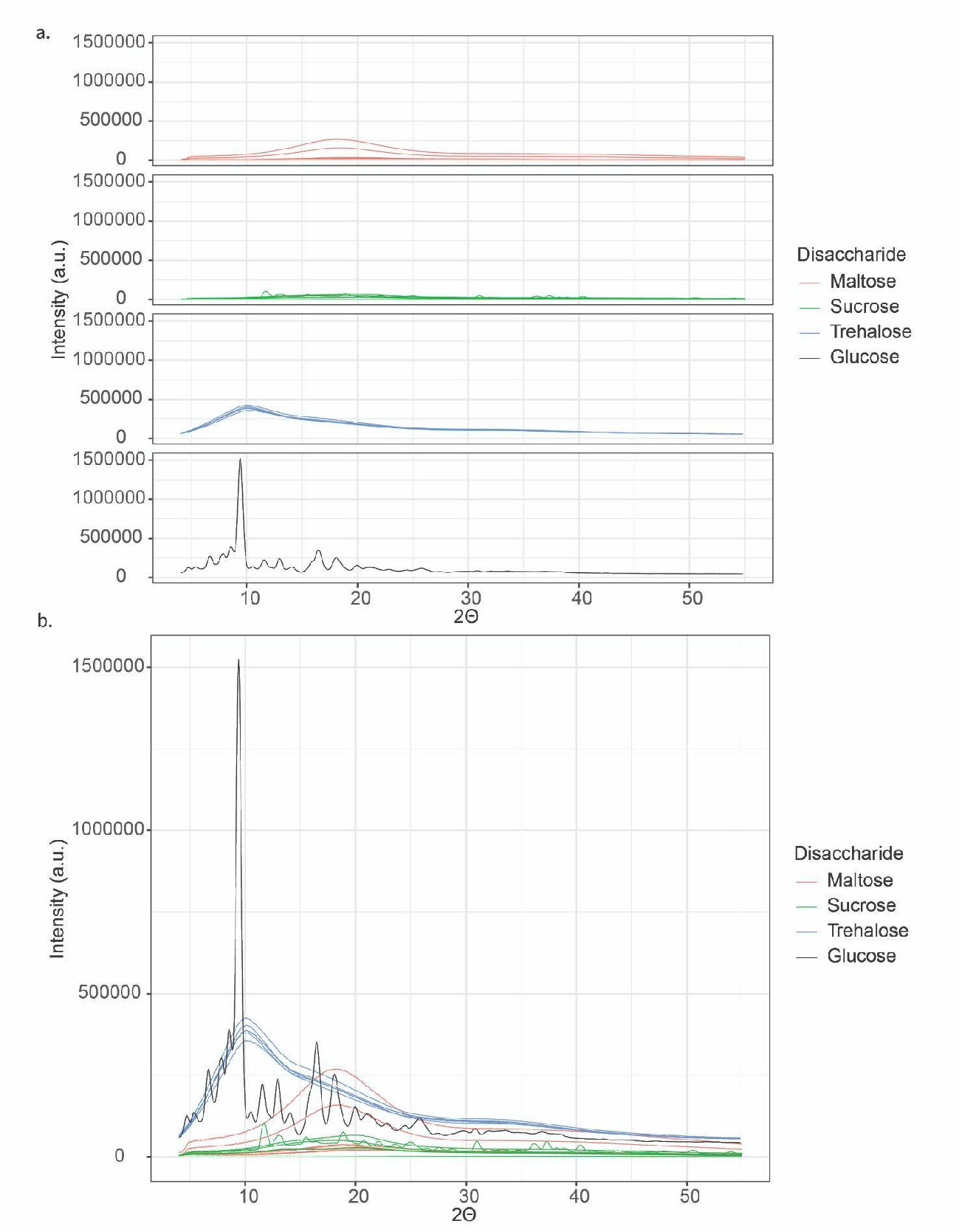
*In vitro* disaccharide-glycerol mixtures produce non-crystalline amorphous (“glassy”) solids when desiccated. a) XRD intensity (a.u.) values for disaccharide-glycerol mixtures organized by disaccharide. b) XRD intensity (a.u.) values for disaccharide-glycerol mixtures organized by disaccharide content.

**Figure S2:**
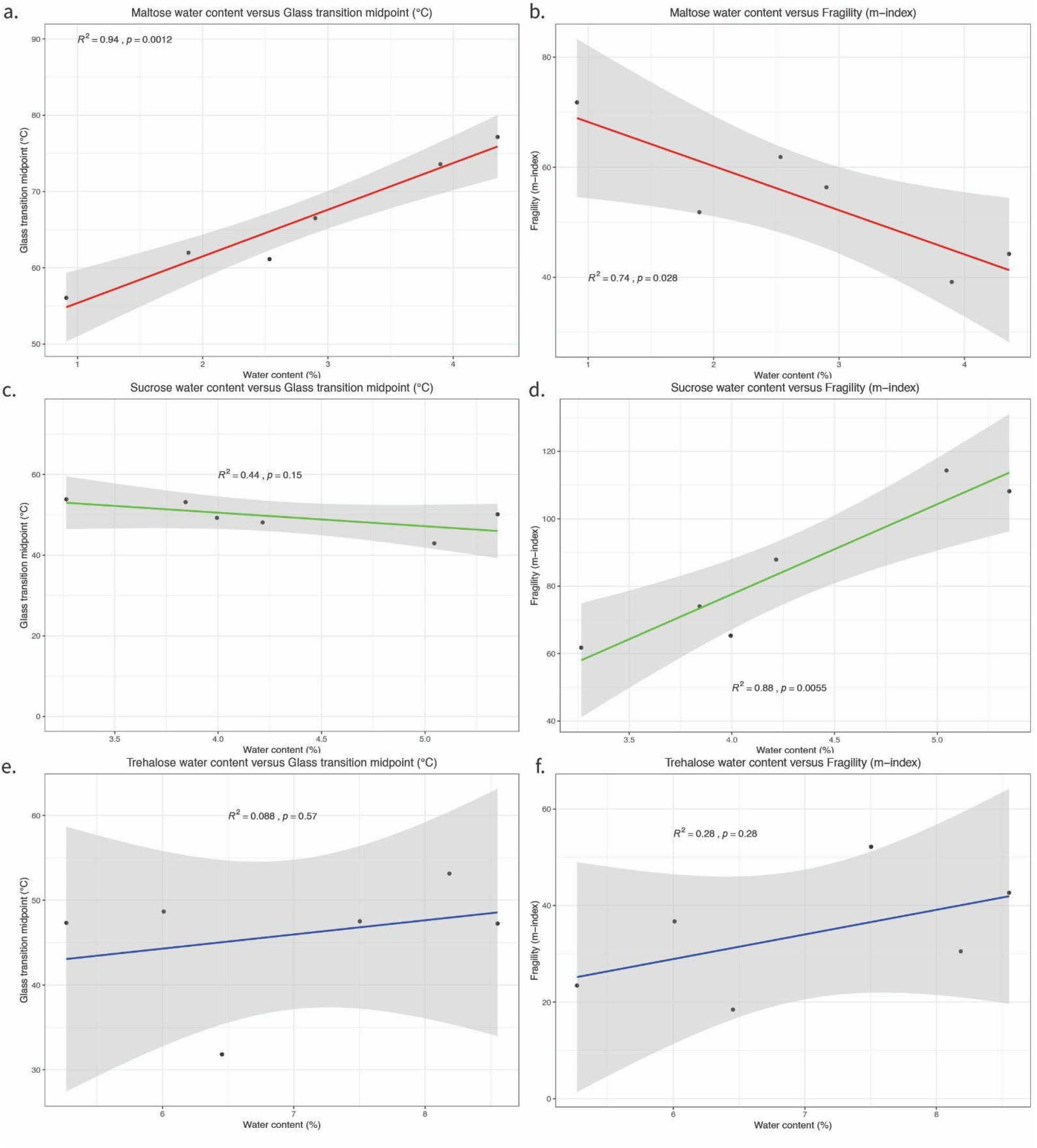
Water content appears to correlate with other material properties for some disaccharide-glycerol mixtures. a) Correlation plot of water content versus glass transition midpoint for maltose-glycerol mixtures. b) Correlation plot of water content versus glass former fragility (m-index) for maltose-glycerol mixtures. c) Correlation plot of water content versus glass transition midpoint for sucrose-glycerol mixtures. d) Correlation plot of water content versus glass former fragility (m-index) for sucrose-glycerol mixtures. e) Correlation plot of water content versus glass transition midpoint for trehalose-glycerol mixtures. f) Correlation plot of water content versus glass former fragility (m-index) for trehalose-glycerol mixtures, error bars represent 95% CI.

**Figure S3:**
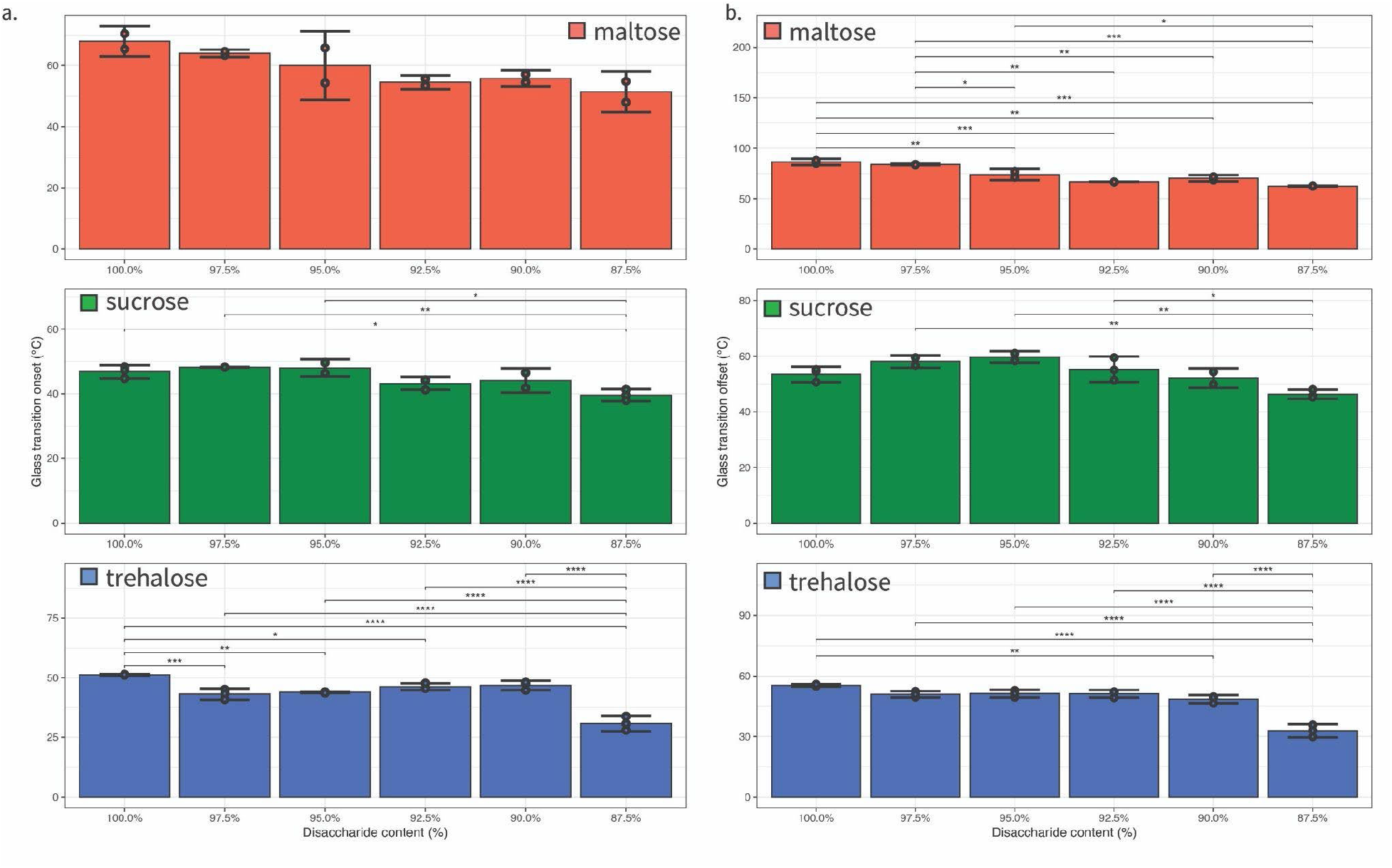
Glass transition onset and offset vary by disaccharide-glycerol mixtures differently than the glass transition midpoint. a) Glass transition onset values for disaccharide-glycerol mixtures organized by disaccharide. b) Glass transition offset values for disaccharide-glycerol mixtures organized by disaccharide. Statistics calculated using a one-way ANOVA and Tukey post-hoc test: p-value ≤ 0.05 (*) p-value ≤ 0.01 (**), p-value ≤ 0.001 (***), pairwise relationships not shown are not significant, error bars represent 95% CI.

**Figure S4:**
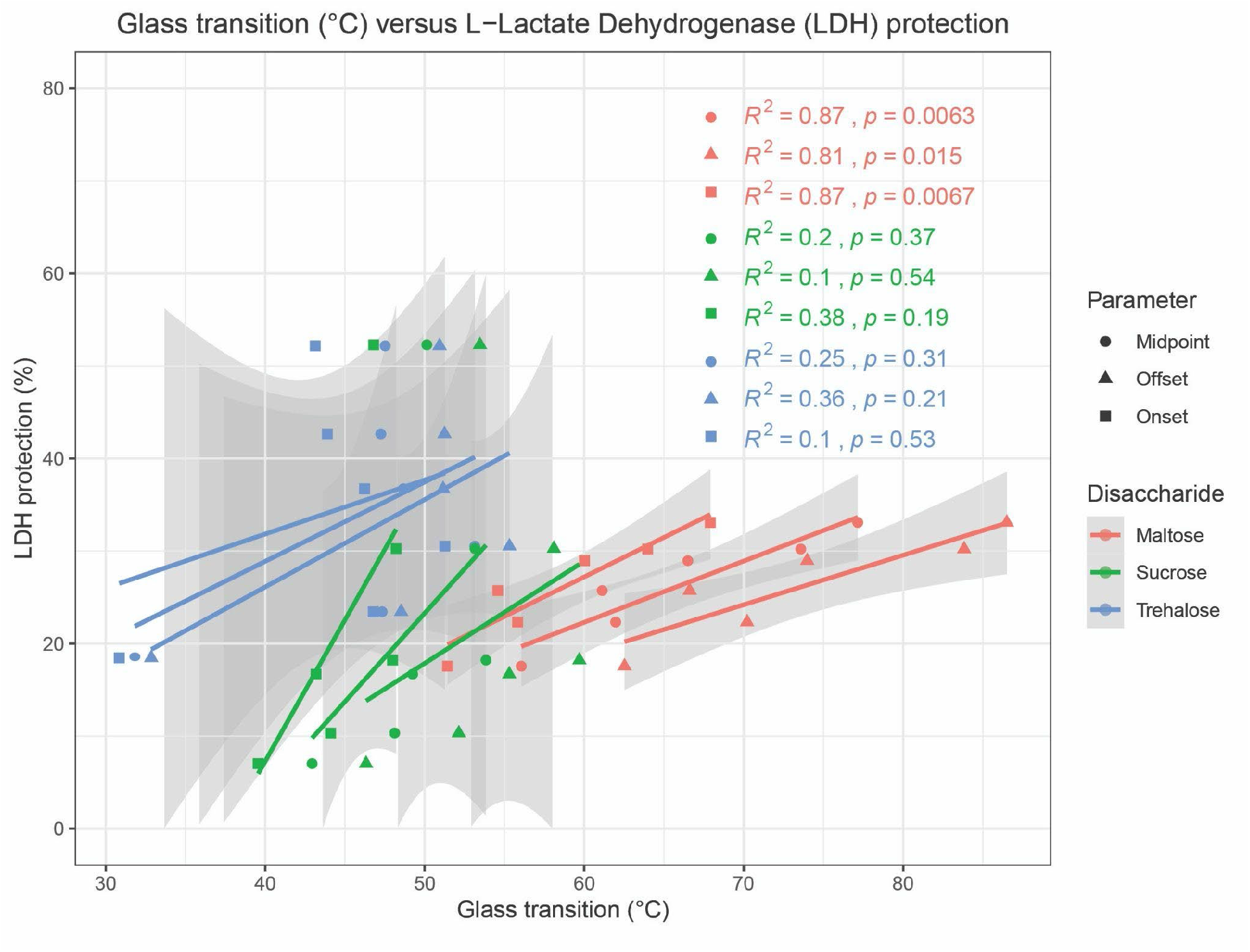
Glass transition onset, midpoint, and offset are better correlated with protection for different disaccharide-glycerol mixtures. Correlation plot of glass transition onset, midpoint, and offset versus protection for maltose-, sucrose-, and trehalose-glycerol mixtures, error bars represent 95% CI.

**Figure S5:**
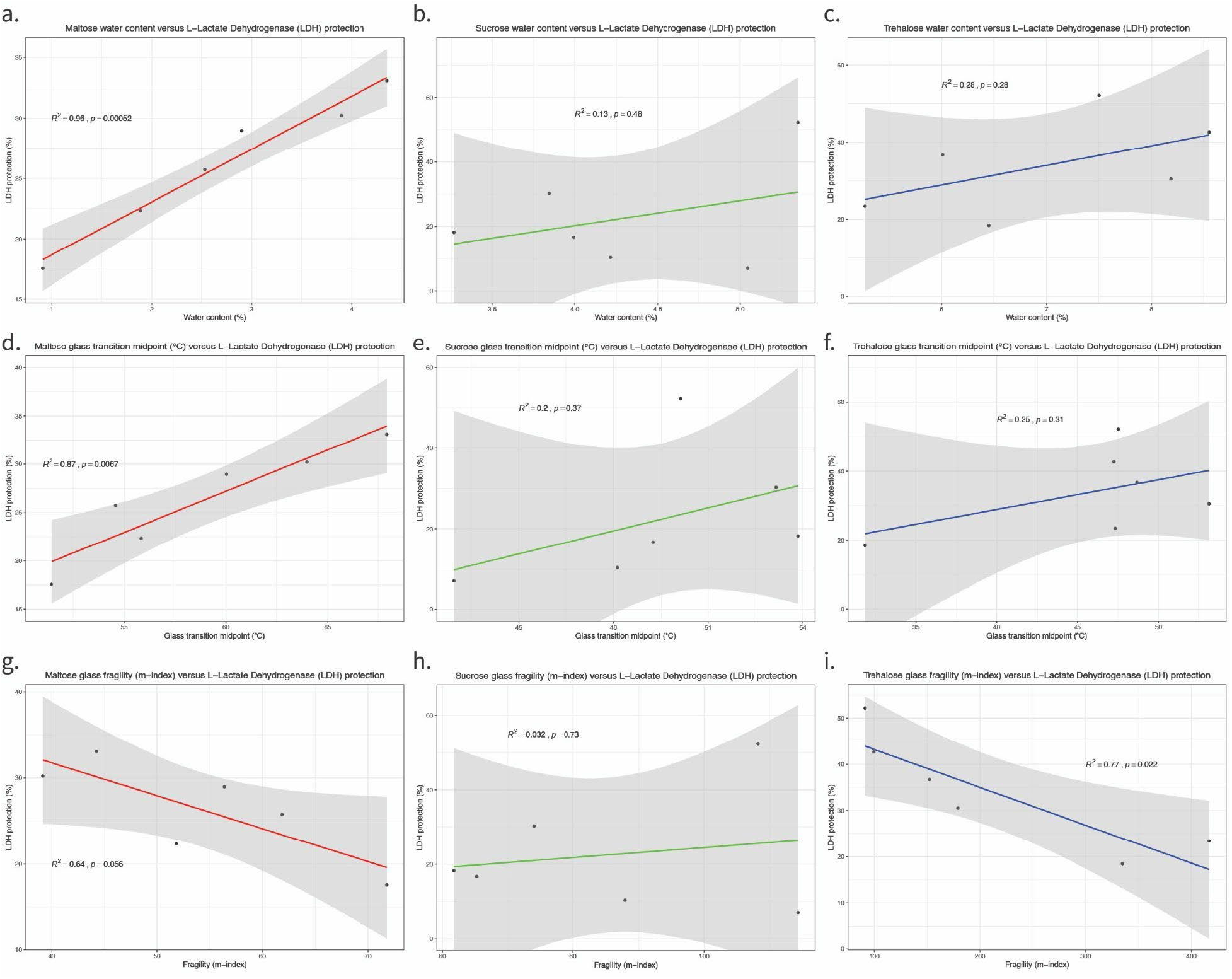
Material property measurements appear to correlate with protection differently for different disaccharides. a) Correlation plot of water content versus LDH protection for maltose-glycerol mixtures. b) Correlation plot of water content versus LDH protection for sucrose-glycerol mixtures. c) Correlation plot of water content versus LDH protection for trehalose-glycerol mixtures. d) Correlation plot of glass transition midpoint temperature versus LDH protection for maltose-glycerol mixtures. e) Correlation plot of glass transition midpoint temperature versus LDH protection for sucrose-glycerol mixtures. f) Correlation plot of glass transition midpoint temperature versus LDH protection for trehalose-glycerol mixtures. g) Correlation plot of glass former fragility (m-index) versus LDH protection for maltose-glycerol mixtures. h) Correlation plot of glass former fragility (m-index) versus LDH protection for sucrose-glycerol mixtures. i) Correlation plot of glass former fragility (m-index) versus LDH protection for trehalose-glycerol mixtures, error bars represent 95% CI.

**Figure S6:**
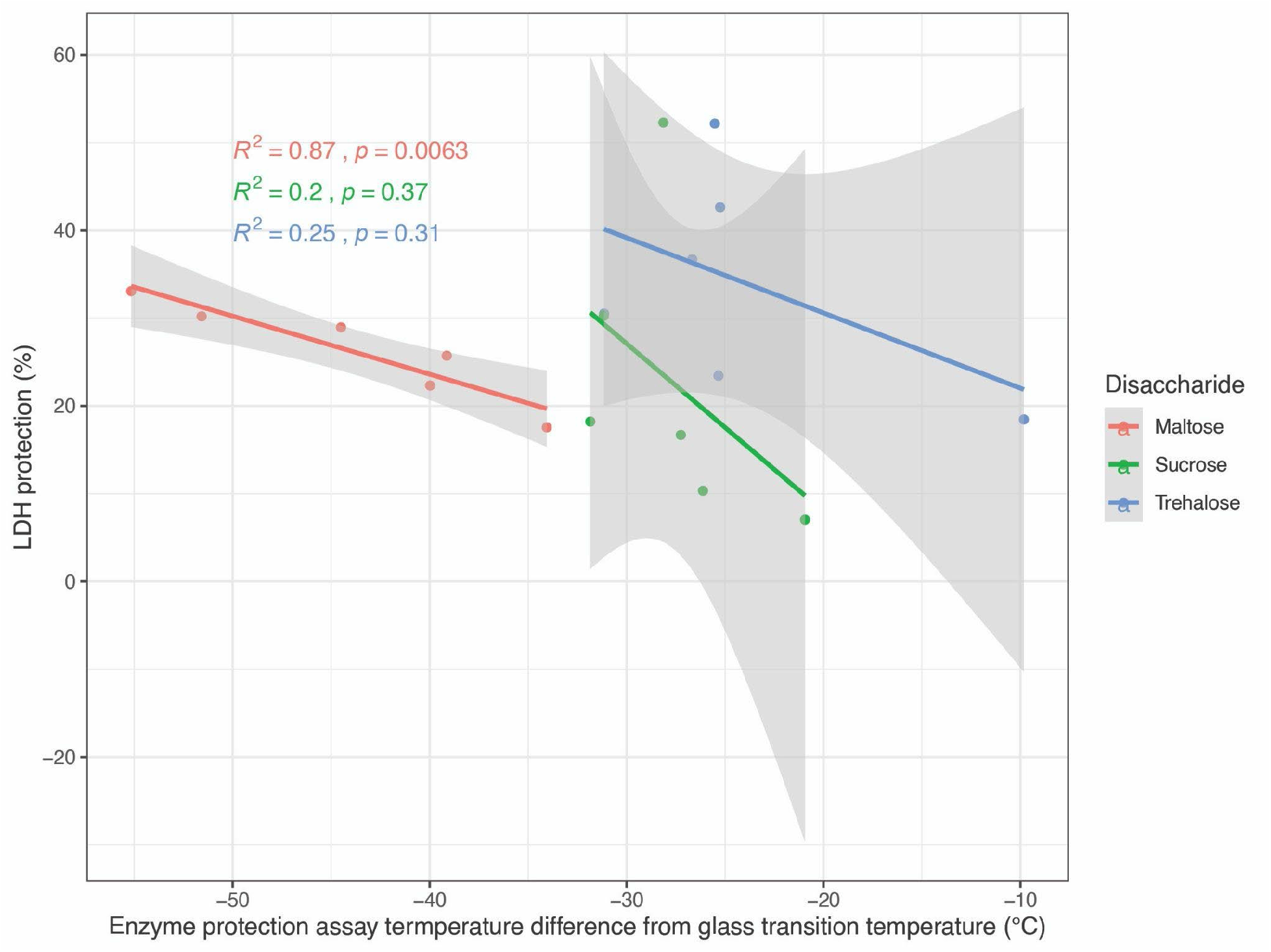
The difference between glass transition temperature and the temperature at which the enzyme protection assay was performed was only significant for maltose-glycerol mixtures. Correlation plot of T_exp_-T_g_ (22 °C) versus protection for maltose-, sucrose-, and trehalose-glycerol mixtures, error bars represent 95% CI.

**Figure S7:**
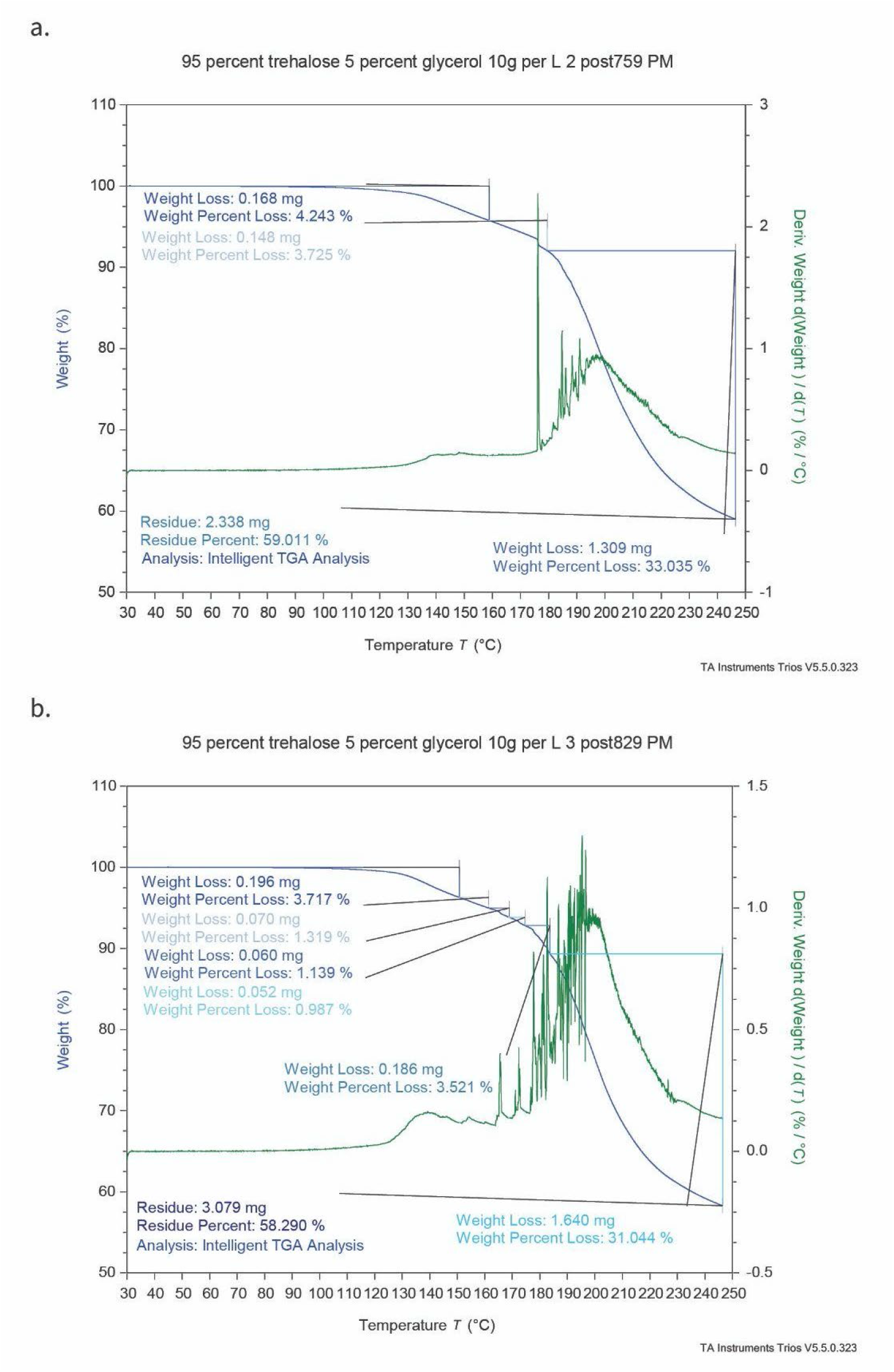
Comparison of simple versus complex interpretation of thermogravimetric analysis results. Step transitions identified by TRIOS Intelligent TGA Analysis, a) Sample TGA thermogram of 95% maltose sample demonstrating easily interpreted results, b) Sample TGA thermogram of 95% maltose sample demonstrating unclear or complex results.

**Figure S8:**
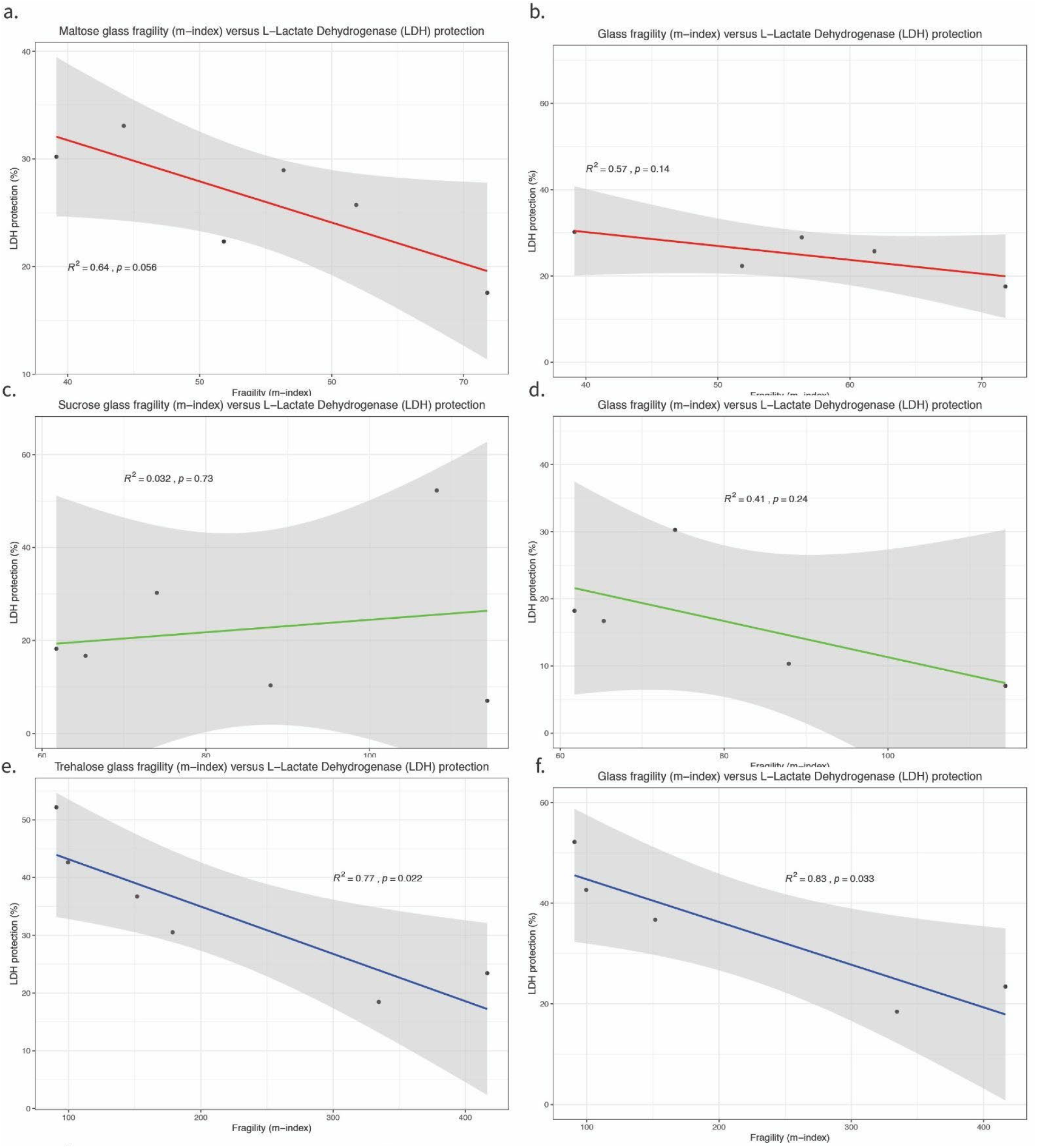
The material properties of pure (100%) disaccharide samples might be different from the material properties of samples with the addition of glycerol, a) Correlation plot of fragility (m-index) versus enzyme protective capacity for maltose-glycerol mixtures, b) Correlation plot of fragility (m-index) versus enzyme protective capacity for maltose-glycerol mixtures without the 100% disaccharide samples, c) Correlation plot of fragility (m-index) versus enzyme protective capacity for sucrose-glycerol mixtures, d) Correlation plot of fragility (m-index) versus enzyme protective capacity for sucrose-glycerol mixtures without the 100% disaccharide samples, e) Correlation plot of fragility (m-index) versus enzyme protective capacity for trehalose-glycerol mixtures, f) Correlation plot of fragility (m-index) versus enzyme protective capacity for trehlose-glycerol mixtures without the 100% disaccharide samples, error bars represent 95% Cl.

## 14.1 Supplemental files

File S1: Processed LDH, TGA, and DSC data

File S2: Raw TGA data

File S3: TGA thermograms

File S4: Raw DSC data

File S5: DSC thermograms

File S6: R scripts used in this study

File S7: XRD data

File S8: Raw LDH data

## References

[1] J.H. Crowe, F.A. Hoekstra, L.M. Crowe, Anhydrobiosis, Annu. Rev. Physiol., 54 (1992) 579–599.

[2] C. Hesgrove, T.C. Boothby, The biology of tardigrade disordered proteins in extreme stress tolerance, Cell Commun. Signal., 18 (2020) 178.

[3] D. Ballesteros, H.W. Pritchard, C. Walters, Dry architecture: towards the understanding of the variation of longevity in desiccation-tolerant germplasm, Seed Sci. Res., 30 (2020) 142–155.

[4] J.D. Hibshman, J.S. Clegg, B. Goldstein, Mechanisms of Desiccation Tolerance: Themes and Variations in Brine Shrimp, Roundworms, and Tardigrades, Front. Physiol., 11 (2020) 592016.

[5] J.H. Crowe, J.F. Carpenter, L.M. Crowe, The role of vitrification in anhydrobiosis, Annu. Rev. Physiol., 60 (1998) 73–103.

[6] M. Sakurai, T. Furuki, K.-I. Akao, D. Tanaka, Y. Nakahara, T. Kikawada, M. Watanabe, T. Okuda, Vitrification is essential for anhydrobiosis in an African chironomid, Polypedilum vanderplanki, Proc. Natl. Acad. Sci. U. S. A., 105 (2008) 5093–5098.

[7] T.C. Boothby, H. Tapia, A.H. Brozena, S. Piszkiewicz, A.E. Smith, I. Giovannini, L. Rebecchi, G.J. Pielak, D. Koshland, B. Goldstein, Tardigrades Use Intrinsically Disordered Proteins to Survive Desiccation, Mol. Cell, 65 (2017) 975–984.e5.

[8] T.C. Boothby, Water content influences the vitrified properties of CAHS proteins, Mol. Cell, 81 (2021) 411–413.

[9] D. Ballesteros, C. Walters, Solid-State Biology and Seed Longevity: A Mechanical Analysis of Glasses in Pea and Soybean Embryonic Axes, Front. Plant Sci., 10 (2019) 920.

[10] L. Weng, G.D. Elliott, Local minimum in fragility for trehalose/glycerol mixtures: implications for biopharmaceutical stabilization, J. Phys. Chem. B, 119 (2015) 6820–6827.

[11] J.C. Wright, Desiccation tolerance and water-retentive mechanisms in tardigrades, J. Exp. Biol., 142 (1989) 267–292.

[12] F.A. Hoekstra, E.A. Golovina, J. Buitink, Mechanisms of plant desiccation tolerance, Trends Plant Sci., 6 (2001) 431–438.

[13] P.S. Belton, A.M. Gil, IR and Raman spectroscopic studies of the interaction of trehalose with hen egg white lysozyme, Biopolymers, 34 (1994) 957–961.

[14] G.I. Olgenblum, L. Sapir, D. Harries, Properties of Aqueous Trehalose Mixtures: Glass Transition and Hydrogen Bonding, J. Chem. Theory Comput., 16 (2020) 1249–1262.

[15] R. Politi, D. Harries, Enthalpically driven peptide stabilization by protective osmolytes, Chem. Commun., 46 (2010) 6449–6451.

[16] S.N. Timasheff, Protein hydration, thermodynamic binding, and preferential hydration, Biochemistry, 41 (2002) 13473–13482.

[17] T.Y. Lin, S.N. Timasheff, On the role of surface tension in the stabilization of globular proteins, Protein Sci., 5 (1996) 372–381.

[18] S. Shimizu, D.J. Smith, Preferential hydration and the exclusion of cosolvents from protein surfaces, J. Chem. Phys., 121 (2004) 1148–1154.

[19] G. Bellavia, S. Giuffrida, G. Cottone, A. Cupane, L. Cordone, Protein thermal denaturation and matrix glass transition in different protein-trehalose-water systems, J. Phys. Chem. B, 115 (2011) 6340–6346.

[20] F. Francia, M. Dezi, A. Mallardi, G. Palazzo, L. Cordone, G. Venturoli, Protein−Matrix Coupling/Uncoupling in “Dry” Systems of Photosynthetic Reaction Center Embedded in Trehalose/Sucrose: The Origin of Trehalose Peculiarity, J. Am. Chem. Soc., 130 (2008) 10240–10246.

[21] A. Lerbret, F. Affouard, Molecular Packing, Hydrogen Bonding, and Fast Dynamics in Lysozyme/Trehalose/Glycerol and Trehalose/Glycerol Glasses at Low Hydration, J. Phys. Chem. B, 121 (2017) 9437–9451.

[22] J.C. Lee, S.N. Timasheff, The stabilization of proteins by sucrose, J. Biol. Chem., 256 (1981) 7193–7201.

[23] C.A. Angell, The old problems of glass and the glass transition, and the many new twists, Proc. Natl. Acad. Sci. U. S. A., 92 (1995) 6675–6682.

[24] C.A. Angell, Entropy, fragility, “landscapes”, and the glass transition, in: Complex Behaviour of Glassy Systems, Springer Berlin Heidelberg, 1997: pp. 1–21.

[25] K. Ito, C.T. Moynihan, C.A. Angell, Thermodynamic determination of fragility in liquids and a fragile-to-strong liquid transition in water, Nature, 398 (1999) 492–495.

[26] E.H. Immergut, H.F. Mark, Principles of Plasticization, in: Plasticization and Plasticizer Processes, AMERICAN CHEMICAL SOCIETY, 1965: pp. 1–26.

[27] A. Anopchenko, T. Psurek, D. VanderHart, J.F. Douglas, J. Obrzut, Dielectric study of the antiplasticization of trehalose by glycerol, Phys. Rev. E Stat. Nonlin. Soft Matter Phys., 74 (2006) 031501.

[28] T.C. Boothby, Mechanisms and evolution of resistance to environmental extremes in animals, Evodevo, 10 (2019) 30.

[29] C.A. Angell, Entropy, fragility, “landscapes”, and the glass transition, Complex Behaviour of Glassy Systems, (n.d.) 1–21.

[30] C. Dinakar, D. Bartels, Desiccation tolerance in resurrection plants: new insights from transcriptome, proteome and metabolome analysis, Front. Plant Sci., 4 (2013) 482.

[31] T.S. Gechev, M. Benina, T. Obata, T. Tohge, N. Sujeeth, I. Minkov, J. Hille, M.-R. Temanni, A.S. Marriott, E. Bergström, J. Thomas-Oates, C. Antonio, B. Mueller-Roeber, J.H.M. Schippers, A.R. Fernie, V. Toneva, Molecular mechanisms of desiccation tolerance in the resurrection glacial relic Haberlea rhodopensis, Cell. Mol. Life Sci., 70 (2013) 689– 709.

[32] L.M. Crowe, Lessons from nature: the role of sugars in anhydrobiosis, Comp. Biochem. Physiol. A Mol. Integr. Physiol., 131 (2002) 505–513.

[33] H. Tapia, D.E. Koshland, Trehalose is a versatile and long-lived chaperone for desiccation tolerance, Curr. Biol., 24 (2014) 2758–2766.

[34] H. Tapia, L. Young, D. Fox, C.R. Bertozzi, D. Koshland, Increasing intracellular trehalose is sufficient to confer desiccation tolerance to Saccharomyces cerevisiae, Proc. Natl. Acad. Sci. U. S. A., 112 (2015) 6122–6127.

[35] S.H. Loomis, K.A.C. Madin, Anhydrobiosis in nematodes: biosynthesis of trehalose, Journal of Experimental, (1980).

[36] C. Womersley, L. Smith, Anhydrobiosis in nematodes—I The role of glycerol myo-inositol and trehalose during desiccation, Comparative Biochemistry and Physiology Part B: Comparative Biochemistry, 70 (1981) 579–586.

[37] C. Womersley, Biochemical and physiological aspects of anhydrobiosis, Comparative Biochemistry and Physiology Part B: Comparative Biochemistry, 70 (1981) 669–678.

[38] M. Robert Michaud, J.B. Benoit, G. Lopez-Martinez, M.A. Elnitsky, R.E. Lee, D.L. Denlinger, Metabolomics reveals unique and shared metabolic changes in response to heat shock, freezing and desiccation in the Antarctic midge, Belgica antarctica, J. Insect Physiol., 54 (2008) 645–655.

[39] K.J. Crowley, G. Zografi, The use of thermal methods for predicting glass-former fragility, Thermochim. Acta, 380 (2001) 79–93.

[40] J.A. Lewis, J.T. Fleming, Basic culture methods, Methods Cell Biol., 48 (1995) 3–29.

[41] YPD media, Cold Spring Harb. Protoc., 2010 (2010) db.rec12315.

[42] T.E. Dirama, G.A. Carri, A.P. Sokolov, Role of hydrogen bonds in the fast dynamics of binary glasses of trehalose and glycerol: a molecular dynamics simulation study, J. Chem. Phys., 122 (2005) 114505.

[43] PubChem, Maltose, https://pubchem.ncbi.nlm.nih.gov/compound/Maltose#section=Chemical-and-Physical-Properties, (n.d.).

[44] PubChem, Sucrose, https://pubchem.ncbi.nlm.nih.gov/compound/Sucrose#section=Chemical-and-Physical-Properties, (n.d.).

[45] PubChem, Trehalose, https://pubchem.ncbi.nlm.nih.gov/compound/Trehalose#section=Chemical-and-Physical-Properties, (n.d.).

[46] A. Chauhan, Powder XRD technique and its applications in science and technology, J. Anal. Bioanal. Tech., 5 (2014).

[47] I.C. Madsen, N.V.Y. Scarlett, A. Kern, Description and survey of methodologies for the determination of amorphous content via X-ray powder diffraction, Zeitschrift Für Kristallographie, 226 (2011) 944–955.

[48] S. Piszkiewicz, K.H. Gunn, O. Warmuth, A. Propst, A. Mehta, K.H. Nguyen, E. Kuhlman, A.J. Guseman, S.S. Stadmiller, T.C. Boothby, S.B. Neher, G.J. Pielak, Protecting activity of desiccated enzymes, Protein Sci., 28 (2019) 941–951.

[49] K. Goyal, L.J. Walton, A. Tunnacliffe, LEA proteins prevent protein aggregation due to water stress, Biochem. J, 388 (2005) 151–157.

[50] M. Rizzuto, A. Mugica, M. Zubitur, D. Caretti, A.J. Müller, Plasticization and anti-plasticization effects caused by poly(lactide-ran-caprolactone) addition to double crystalline poly(l-lactide)/poly(ε-caprolactone) blends, CrystEngComm, 18 (2016) 2014–2023.

[51] E. Luk, A.J. Sandoval, A. Cova, A.J. Müller, Anti-plasticization of cassava starch by complexing fatty acids, Carbohydr. Polym., 98 (2013) 659–664.

[52] Y. Chen, T. Tang, C. Ayranci, Moisture-induced anti-plasticization of polylactic acid: Experiments and modeling, J. Appl. Polym. Sci., 139 (2022) 52369.

[53] Y. Figueroa, M. Guevara, A. Pérez, A. Cova, A.J. Sandoval, A.J. Müller, Effect of sugar addition on glass transition temperatures of cassava starch with low to intermediate moisture contents, Carbohydr. Polym., 146 (2016) 231–237.

[54] Y. Liu, A.K. Roy, A.A. Jones, P.T. Inglefield, P. Ogden, An NMR study of plasticization and antiplasticization of a polymeric glass, Macromolecules, 23 (1990) 968–977.

[55] J. Tarique, S.M. Sapuan, A. Khalina, Effect of glycerol plasticizer loading on the physical, mechanical, thermal, and barrier properties of arrowroot (Maranta arundinacea) starch biopolymers, Sci. Rep., 11 (2021) 13900.

[56] C.T. Moynihan, Correlation between the width of the glass transition region and the temperature dependence of the viscosity of high-tg glasses, J. Am. Ceram. Soc., 76 (1993) 1081–1087.

[57] C.T. Moynihan, S.-K. Lee, M. Tatsumisago, T. Minami, Estimation of activation energies for structural relaxation and viscous flow from DTA and DSC experiments, Thermochim. Acta, 280–281 (1996) 153–162.

[58] Y.I. Matveev, V.Y. Grinberg, V.B. Tolstoguzov, The plasticizing effect of water on proteins, polysaccharides and their mixtures Glassy state of biopolymers, food and seeds, Food Hydrocoll., 14 (2000) 425–437.

[59] G.N. Ruiz, M. Romanini, A. Hauptmann, T. Loerting, E. Shalaev, J.L. Tamarit, L.C. Pardo, R. Macovez, Genuine antiplasticizing effect of water on a glass-former drug, Sci. Rep., 7 (2017) 7470.

[60] I. Combarro Palacios, C. Olsson, C.S. Kamma-Lorger, J. Swenson, S. Cerveny, Motions of water and solutes-Slaving versus plasticization phenomena, J. Chem. Phys., 150 (2019) 124902.

[61] D. Calahan, M. Dunham, C. DeSevo, D.E. Koshland, Genetic analysis of desiccation tolerance in Sachharomyces cerevisiae, Genetics, 189 (2011) 507–519.

[62] B. Janis, C. Belott, M.A. Menze, Role of Intrinsic Disorder in Animal Desiccation Tolerance, Proteomics, 18 (2018) e1800067.

[63] C. Erkut, S. Penkov, H. Khesbak, D. Vorkel, J.-M. Verbavatz, K. Fahmy, T.V. Kurzchalia, Trehalose renders the dauer larva of Caenorhabditis elegans resistant to extreme desiccation, Curr. Biol., 21 (2011) 1331–1336.

[64] J.C. Dyre, Colloquium: The glass transition and elastic models of glass-forming liquids, Rev. Mod. Phys., 78 (2006) 953–972.

[65] G. Roudaut, D. Simatos, D. Champion, E. Contreras-Lopez, M. Le Meste, Molecular mobility around the glass transition temperature: a mini review, Innov. Food Sci. Emerg. Technol., 5 (2004) 127–134.

[66] C. Walters, Temperature dependency of molecular mobility in preserved seeds, Biophys. J., 86 (2004) 1253–1258.

[67] D. Ballesteros, C. Walters, Detailed characterization of mechanical properties and molecular mobility within dry seed glasses: relevance to the physiology of dry biological systems, Plant J., 68 (2011) 607–619.

[68] C. Walters, Understanding the mechanisms and kinetics of seed aging, Seed Sci. Res., 8 (1998) 223–244.

[69] O. Leprince, C. Walters-Vertucci, A Calorimetric Study of the Glass Transition Behaviors in Axes of Bean Seeds with Relevance to Storage Stability, Plant Physiol., 109 (1995) 1471–1481.

