## Supplementary figures and images for "Water content, transition temperature and fragility influence protection and anhydrobiotic capacity"

### 87.5 percent trehalose 12.5 percent glycerol 10g per L 1 post143 PM.pdf

87.5 percent trehalose 12.5 percent glycerol 10g per L 1 post143 PM

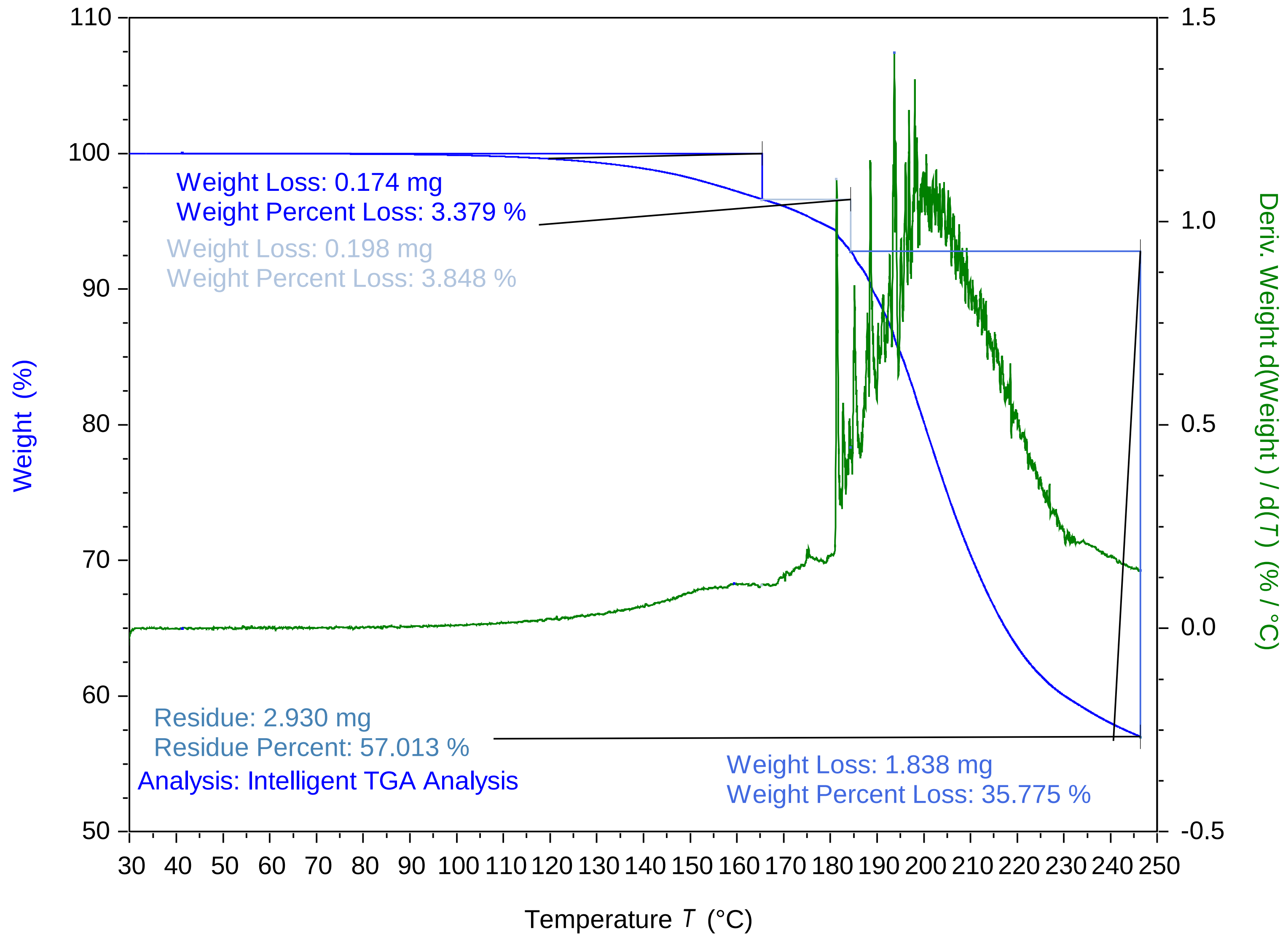


TA Instruments Trios V5.5.0.323

### 87.5 percent trehalose 12.5 percent glycerol 10g per L 2 post213 PM.pdf

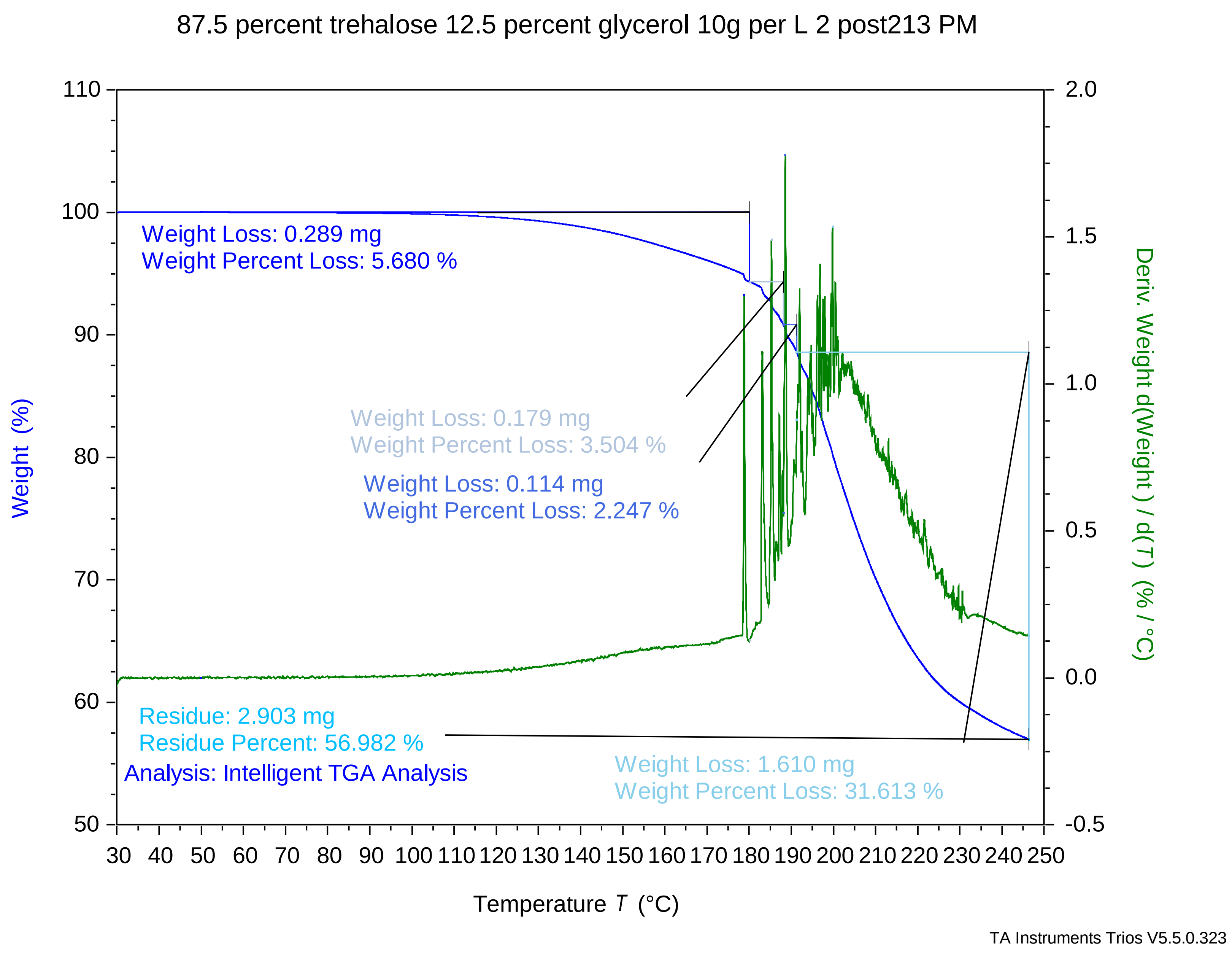

### 90 percent trehalose 10 percent glycerol 10g per L 1 Post144 PM.pdf

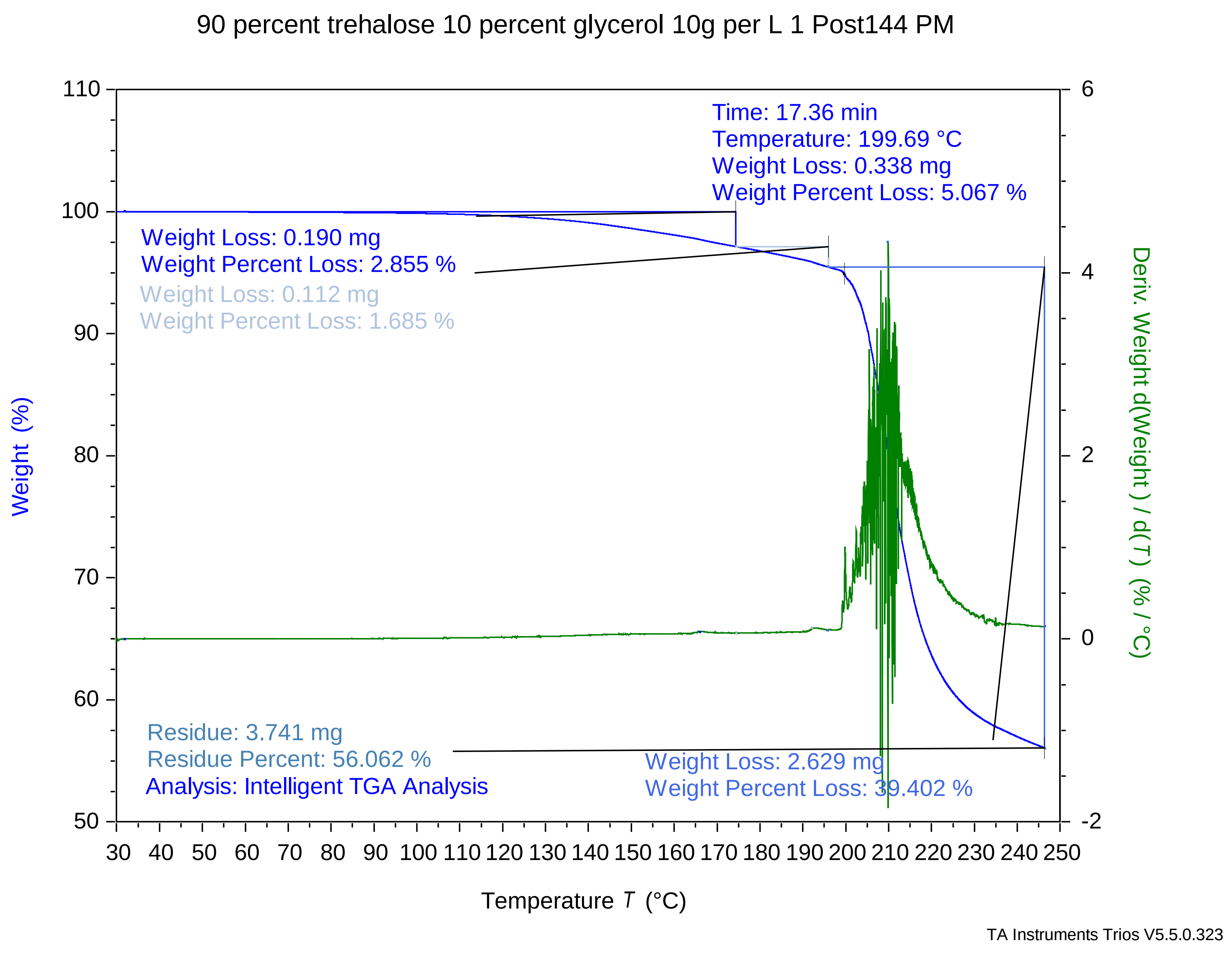

### 90 percent trehalose 10 percent glycerol 10g per L 2 Post214 PM.pdf

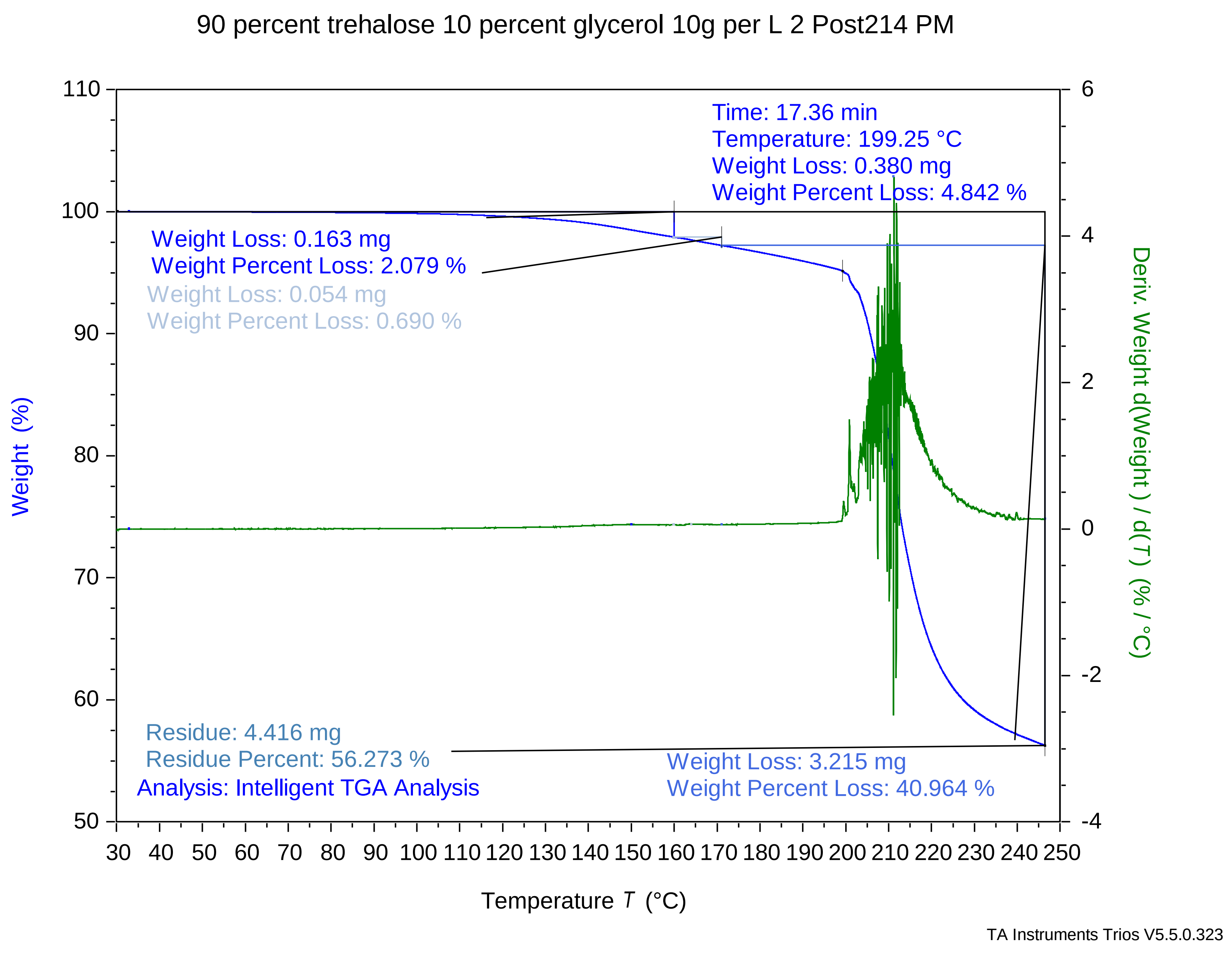

### 90 percent trehalose 10 percent glycerol 10g per L 3 Post245 PM.pdf

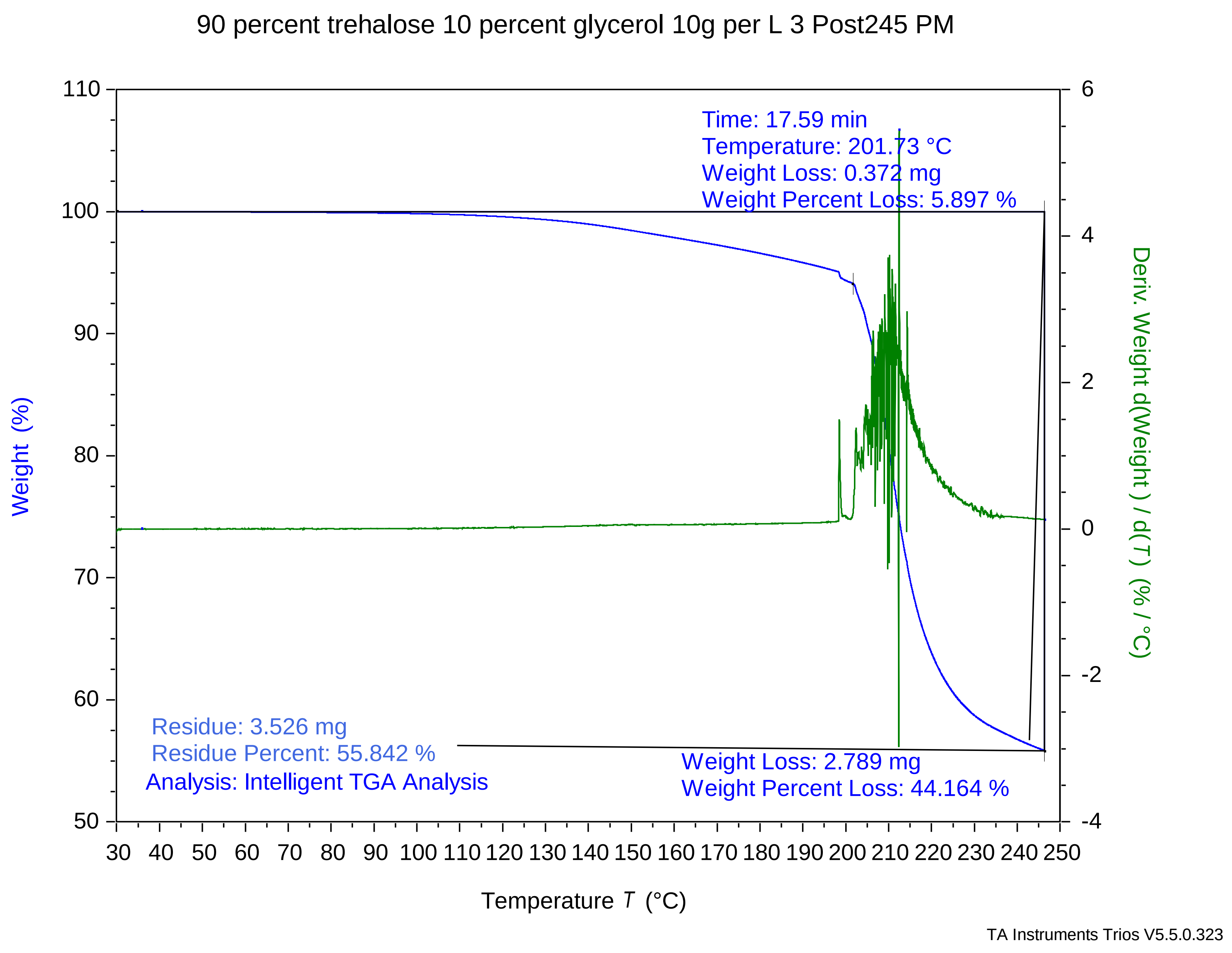

### 92.5 percent trehalose 7.5 percent glycerol 10g per L 1 post1212 PM.pdf

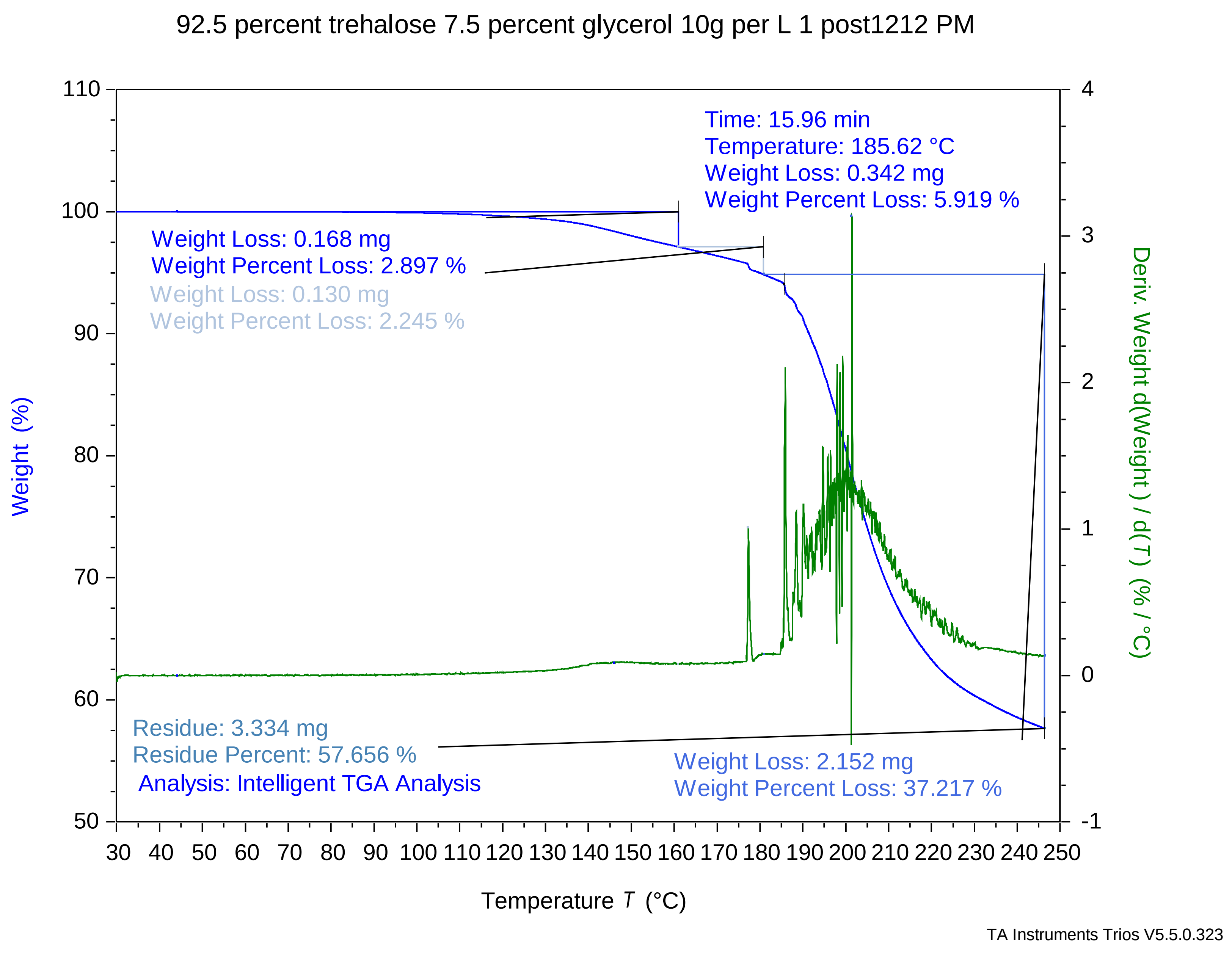

### 92.5 percent trehalose 7.5 percent glycerol 10g per L 2 post1242 PM.pdf

#### 92.5 percent trehalose 7.5 percent glycerol 10g per L 2 post1242 PM

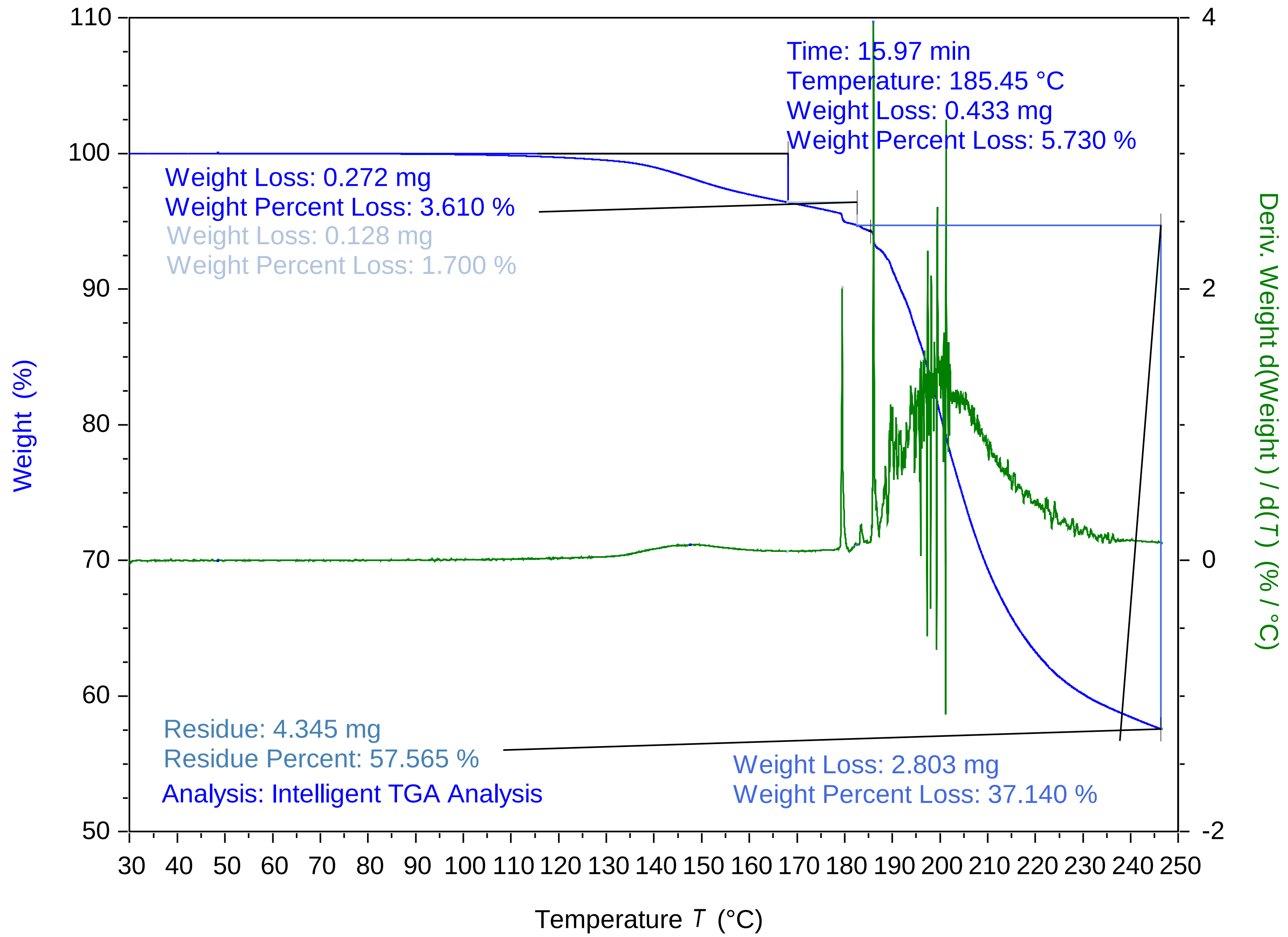


TA Instruments Trios V5.5.0.323

### 92.5 percent trehalose 7.5 percent glycerol 10g per L 3 post113 PM.pdf

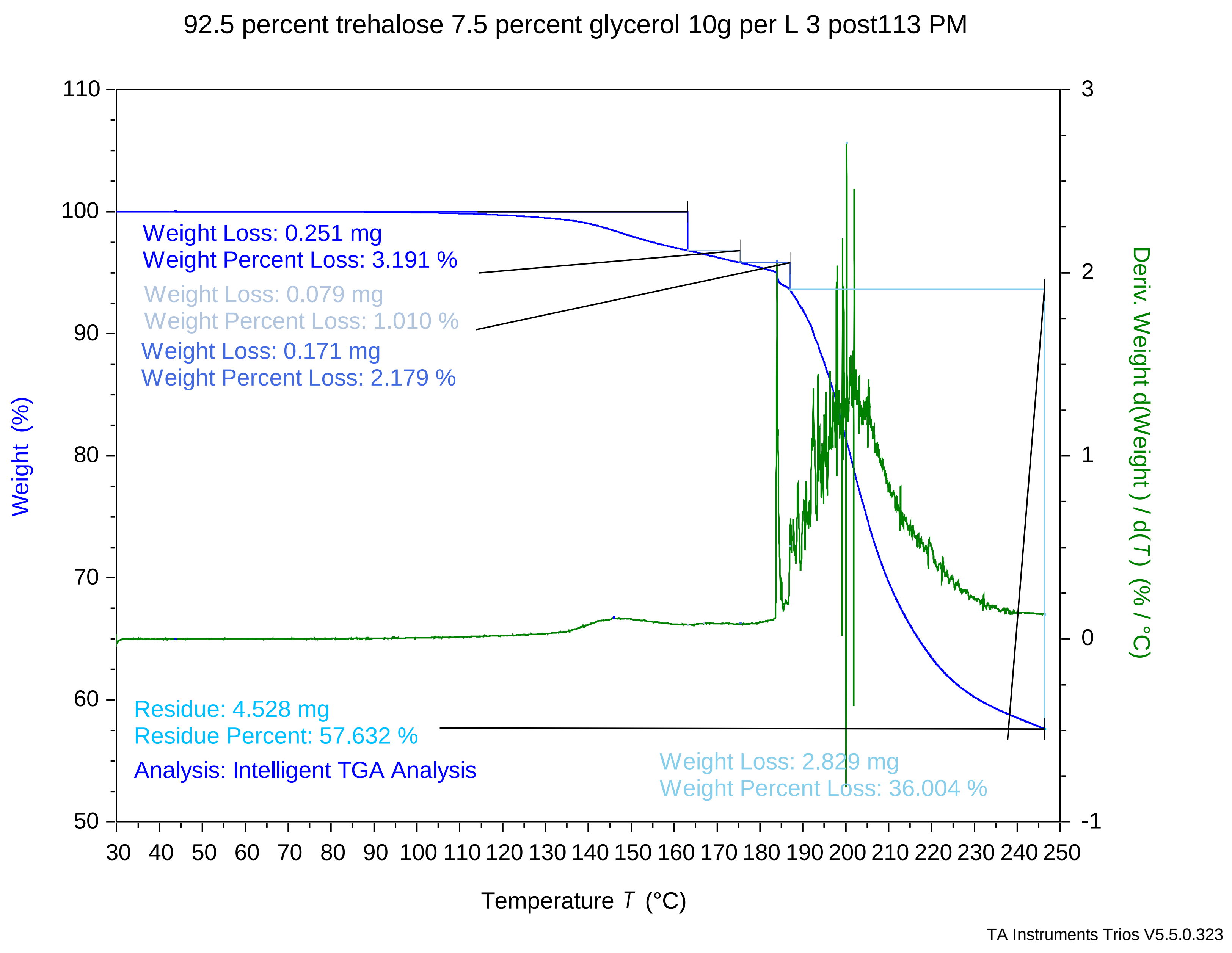

### 95 percent trehalose 5 percent glycerol 10g per L 1 post729 PM.pdf

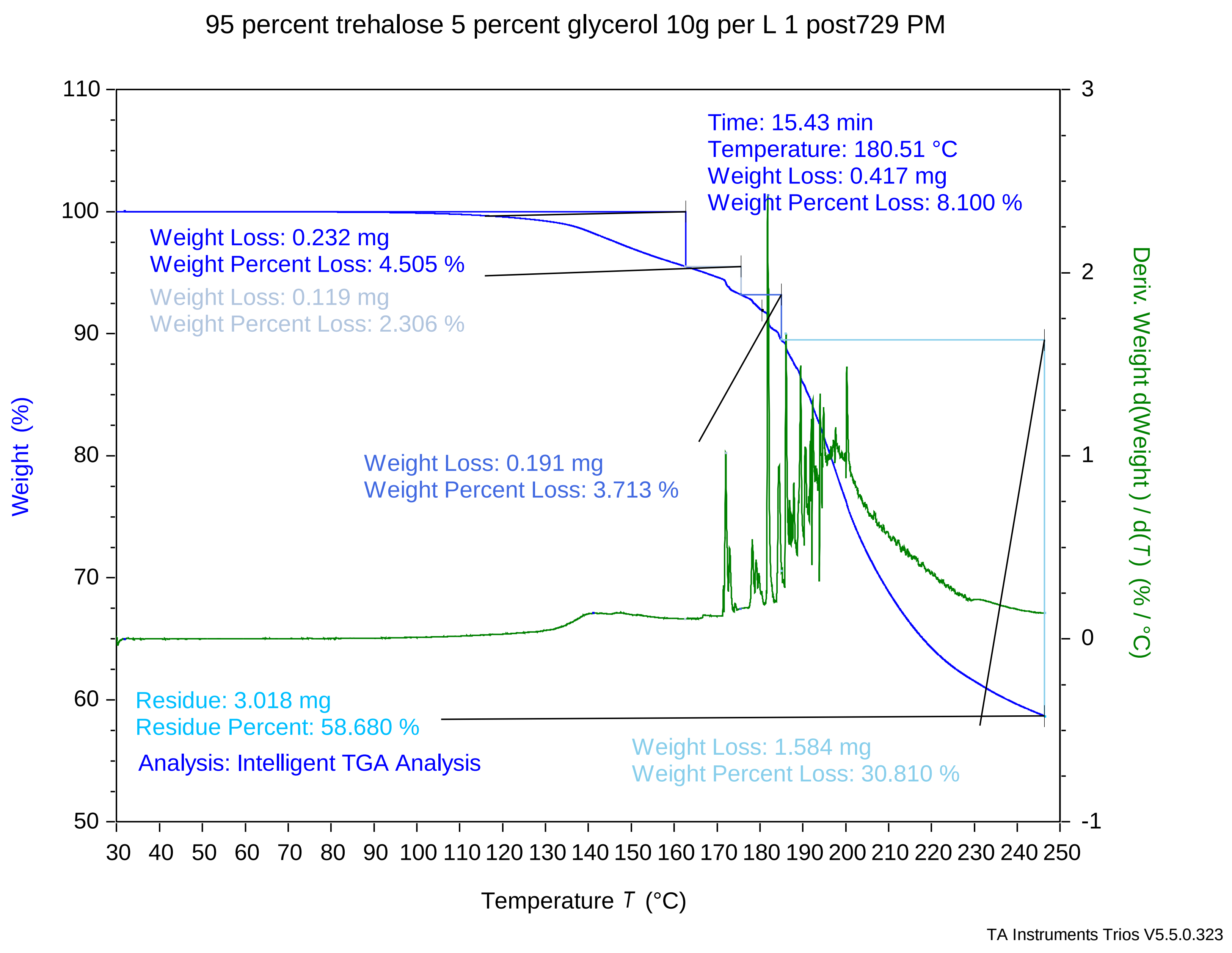

### 95 percent trehalose 5 percent glycerol 10g per L 2 post759 PM.pdf

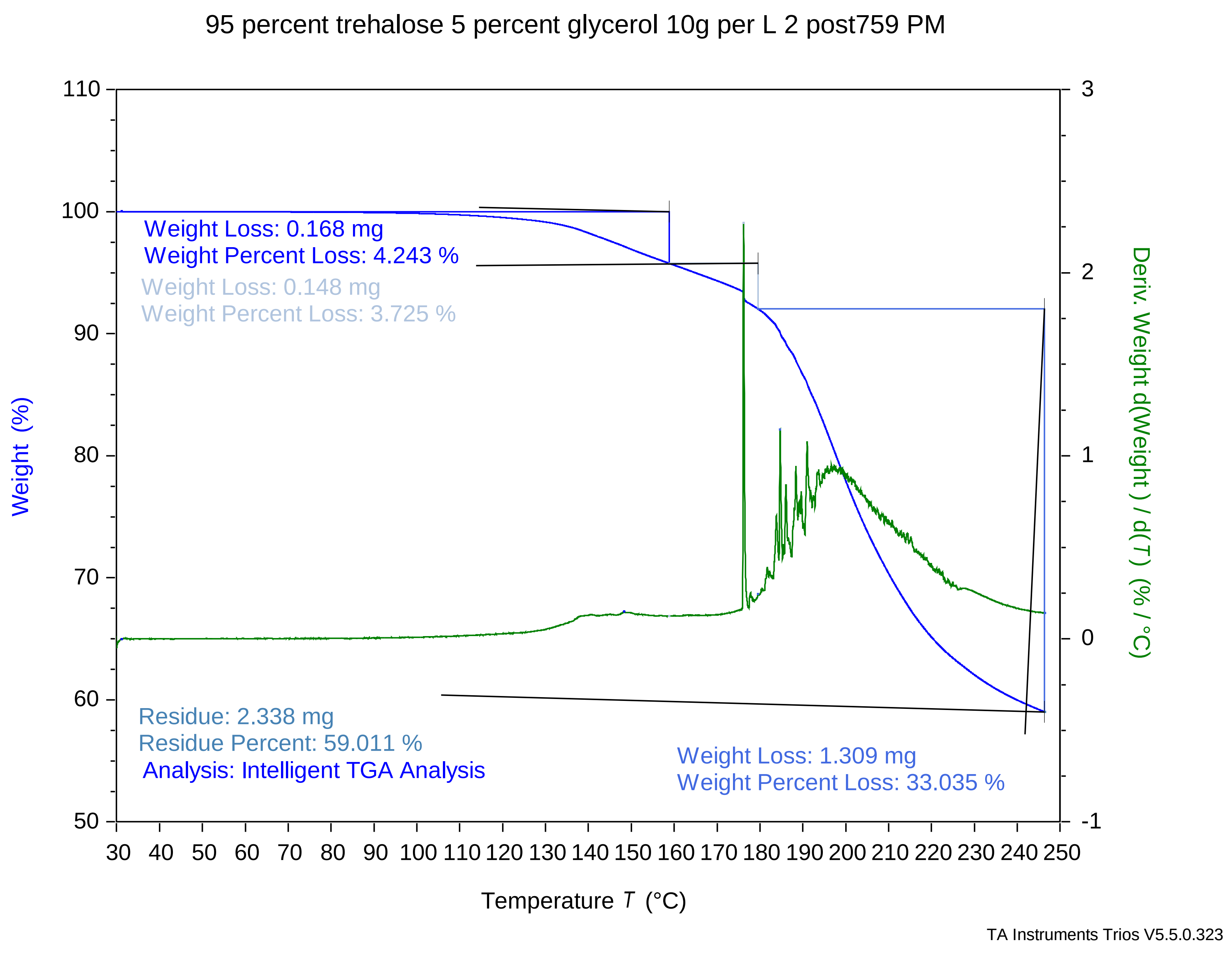

### 95 percent trehalose 5 percent glycerol 10g per L 3 post829 PM.pdf

#### 95 percent trehalose 5 percent glycerol 10g per L 3 post829 PM

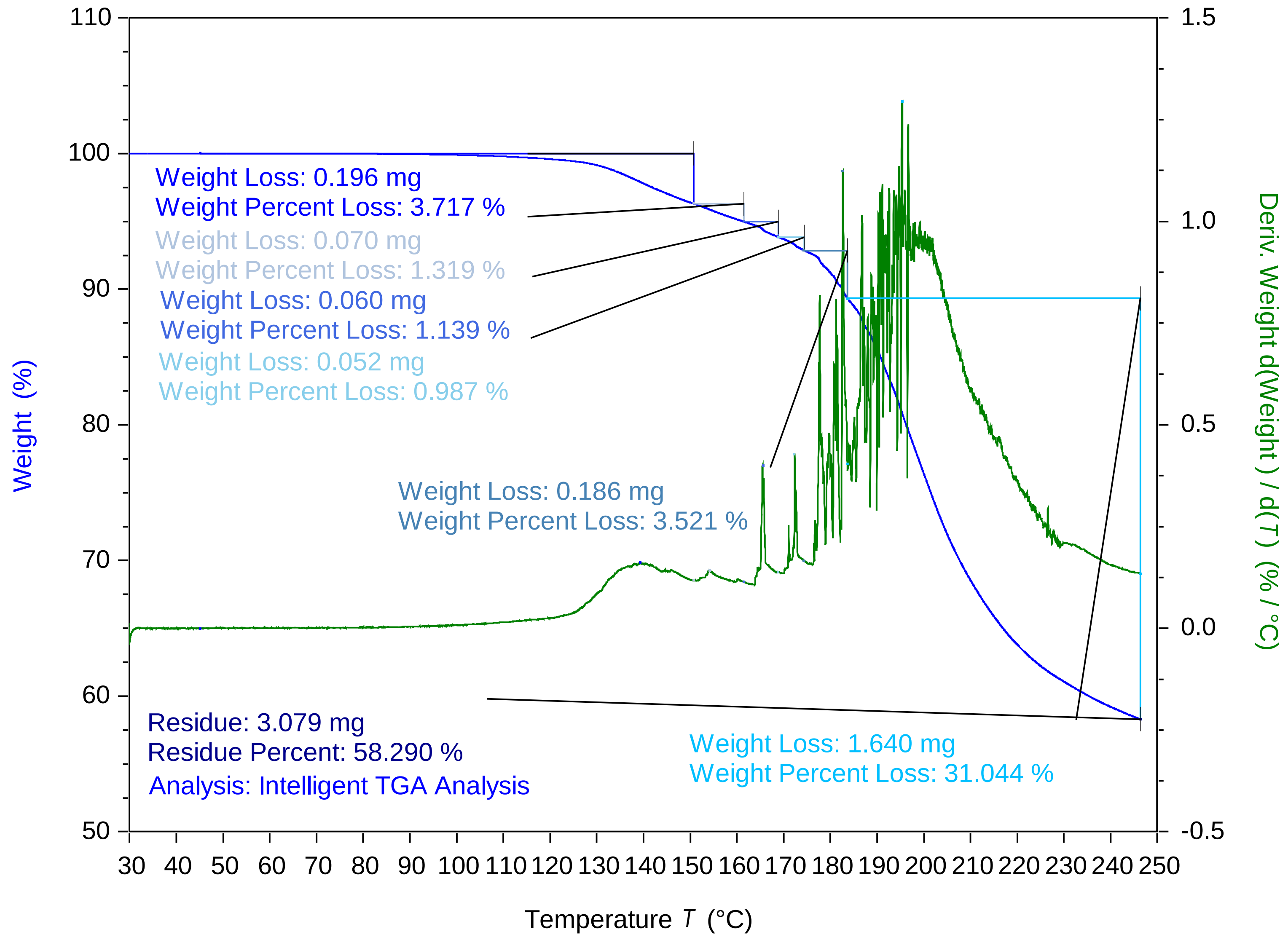


TA Instruments Trios V5.5.0.323

### 97.5 percent trehalose 2.5 percent glycerol 10g per L 1 post449 PM.pdf

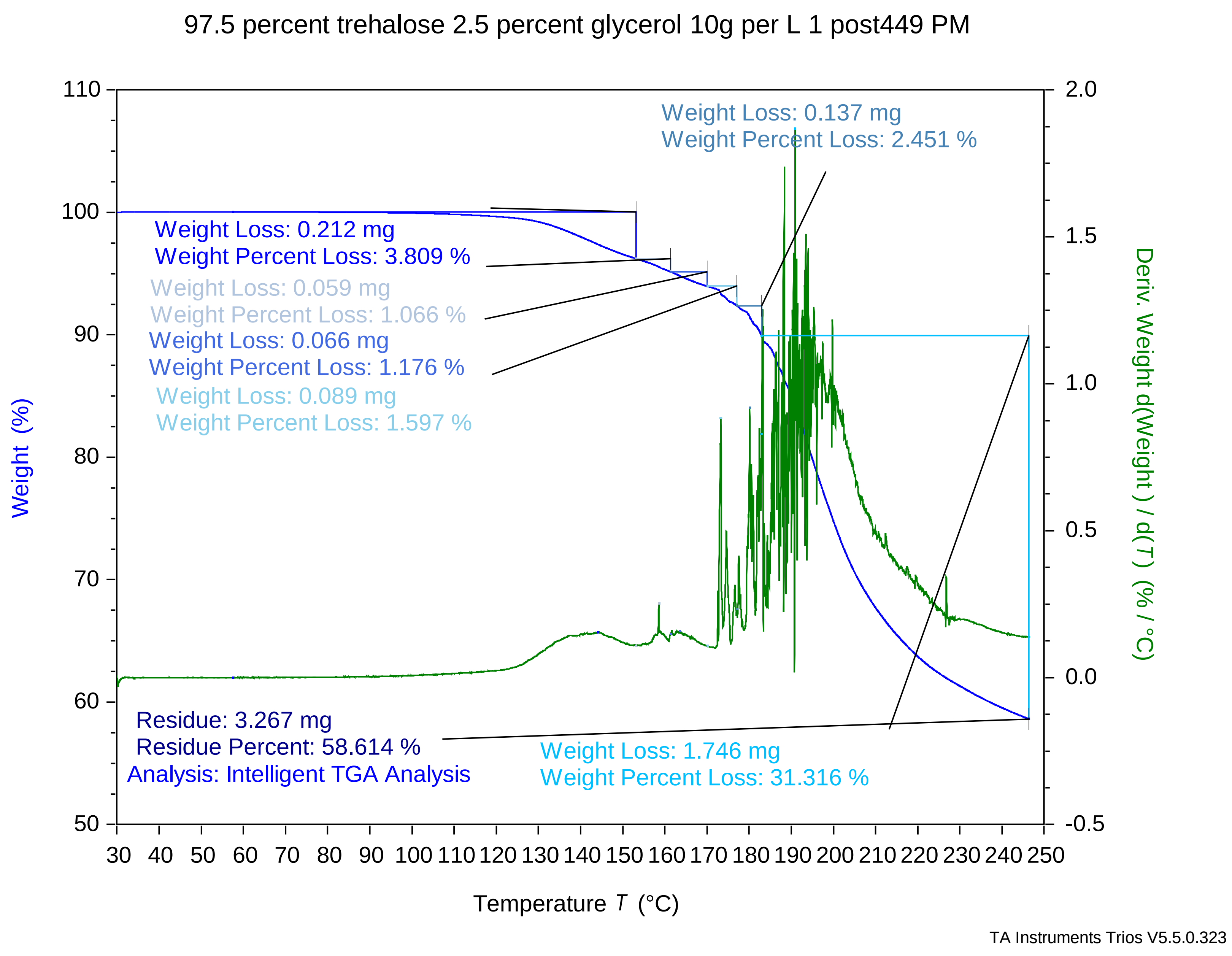

### 97.5 percent trehalose 2.5 percent glycerol 10g per L 2 post519 PM.pdf

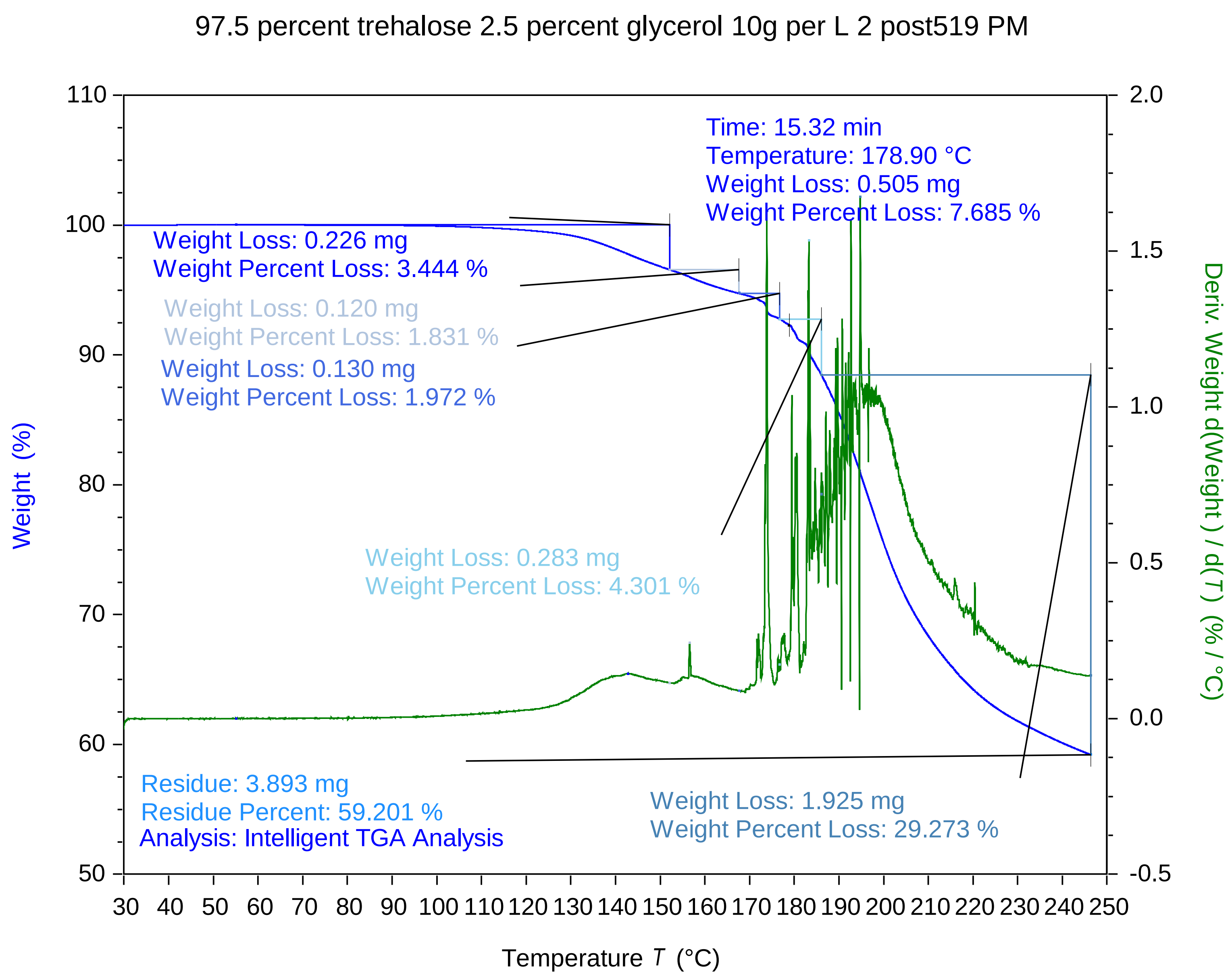

TA Instruments Trios V5.5.0.323

### 97.5 percent trehalose 2.5 percent glycerol 10g per L 3 post549 PM.pdf

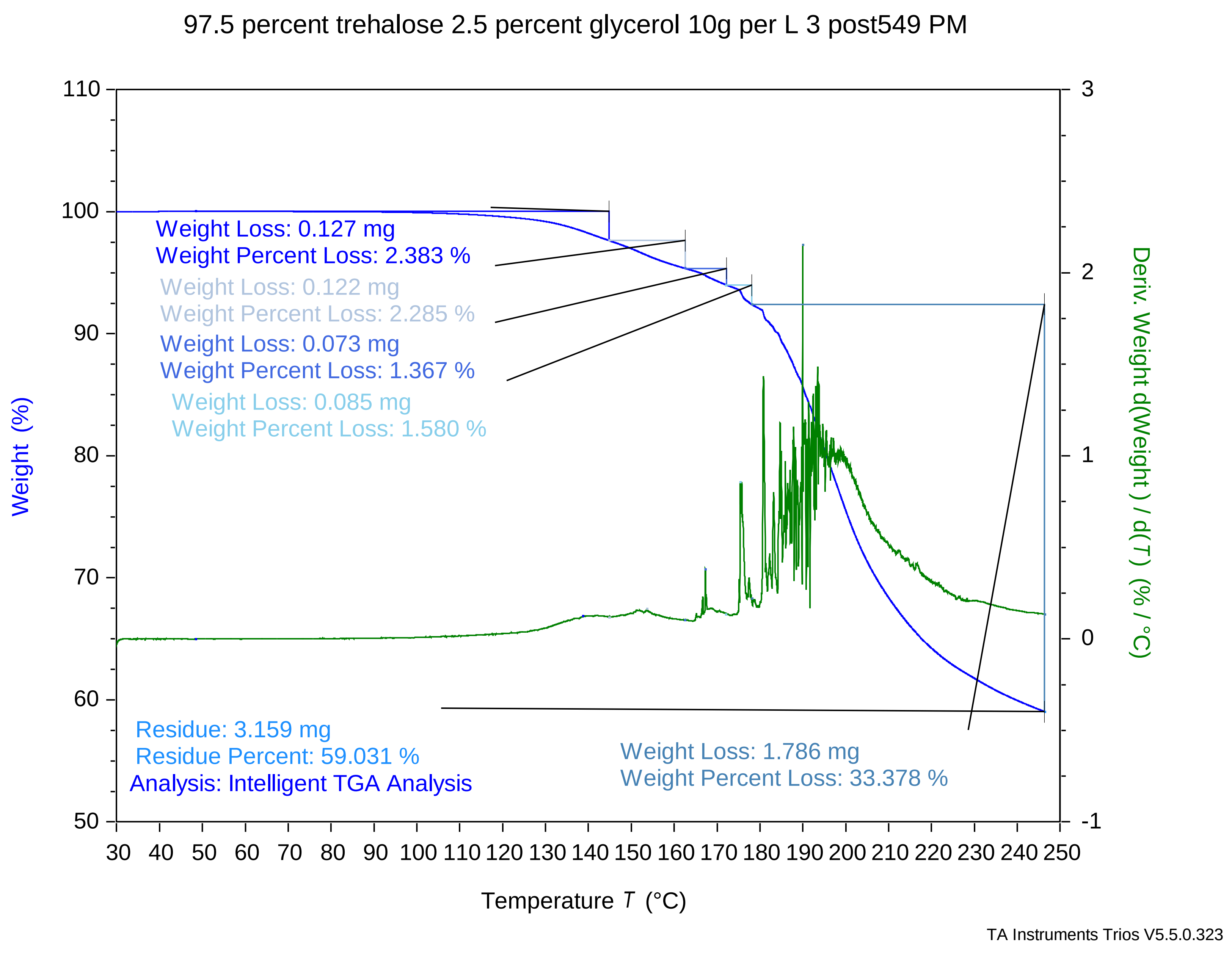

### 100 percent trehalose 0 percent glycerol 10g per L 1 post437 PM.pdf

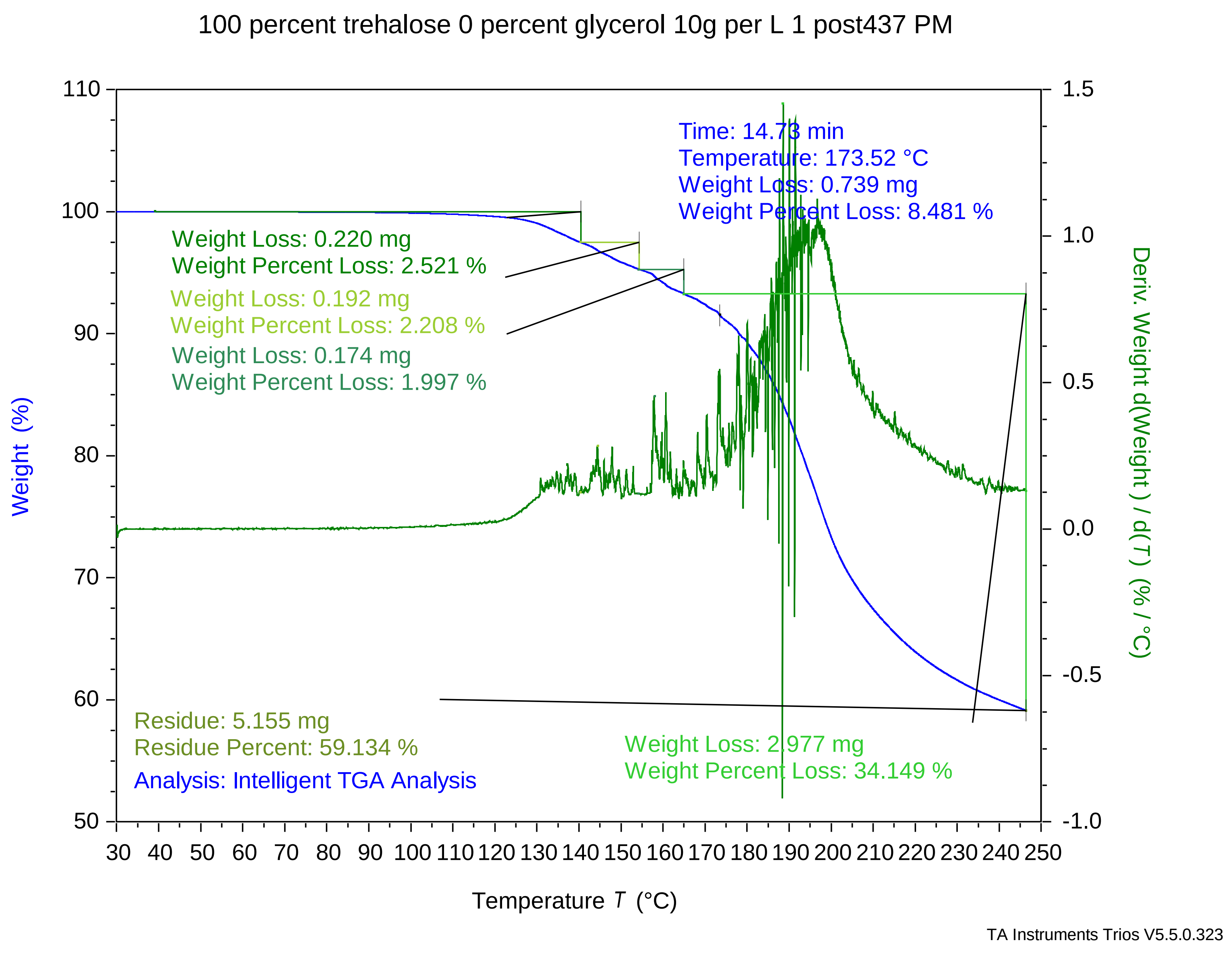

### 100 percent trehalose 0 percent glycerol 10g per L 2 post507 PM.pdf

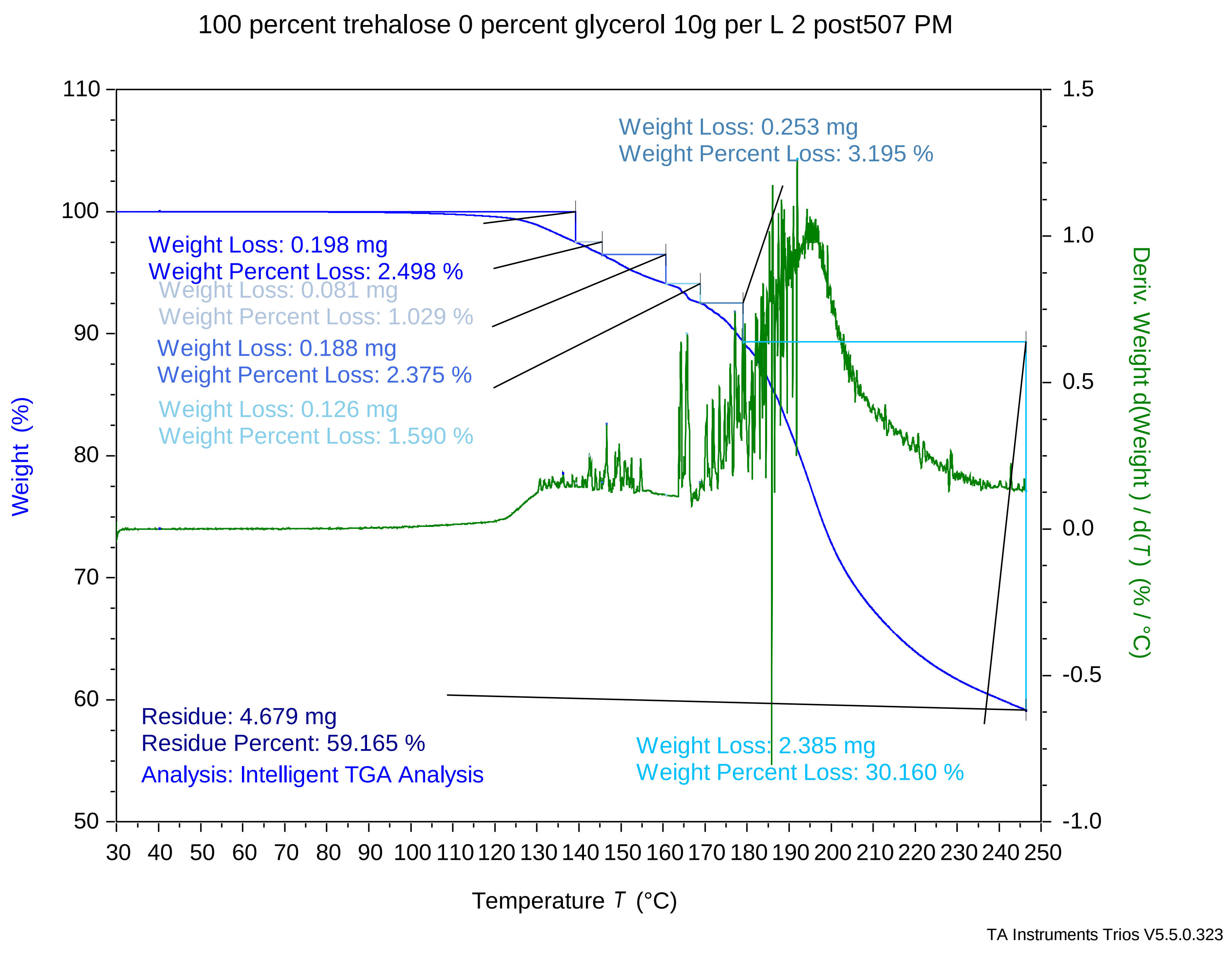

### 100 percent trehalose 0 percent glycerol 10g per L 3 post537 PM.pdf

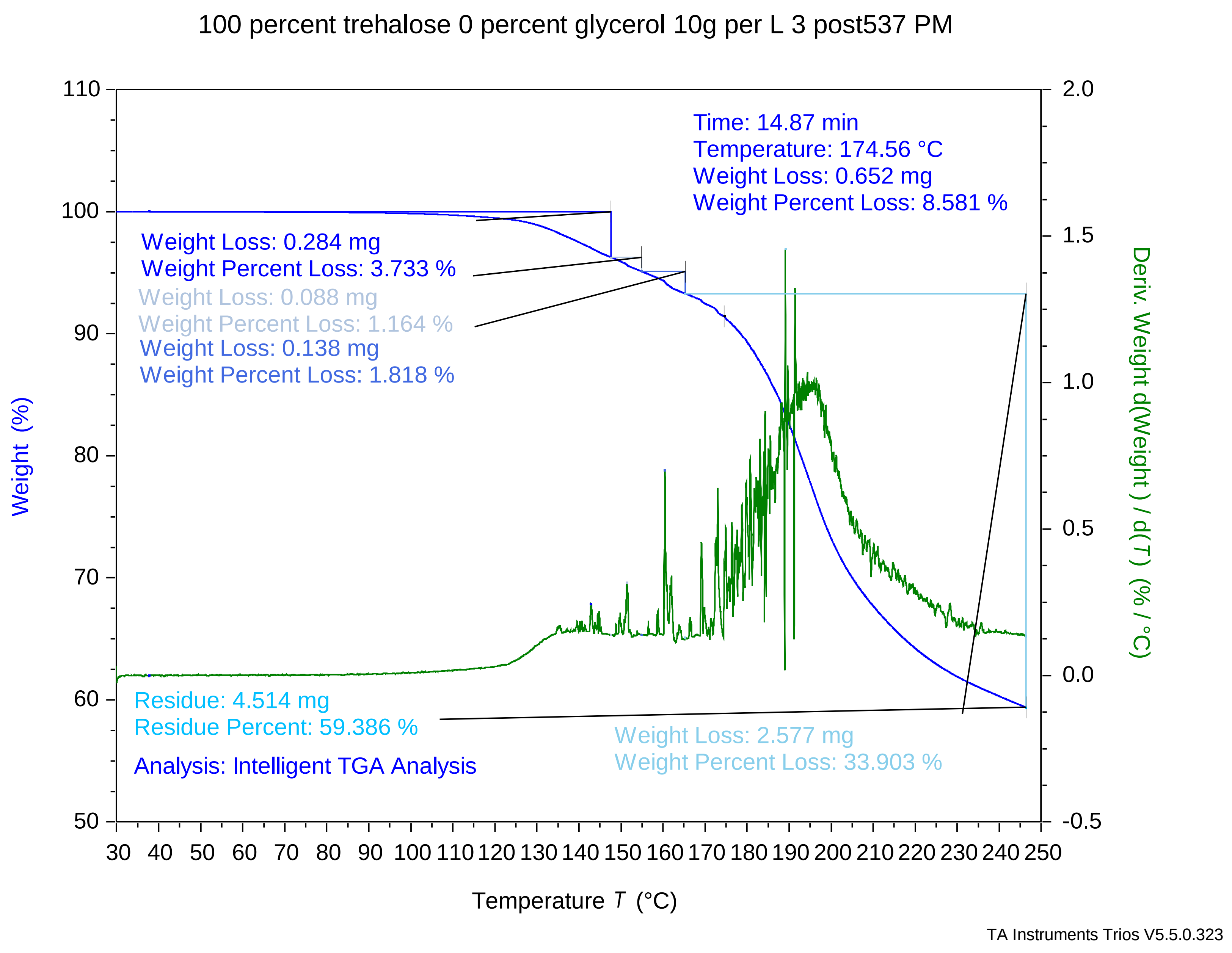

### Maltose + Glycerol 87.5 12212022 150 PM.pdf

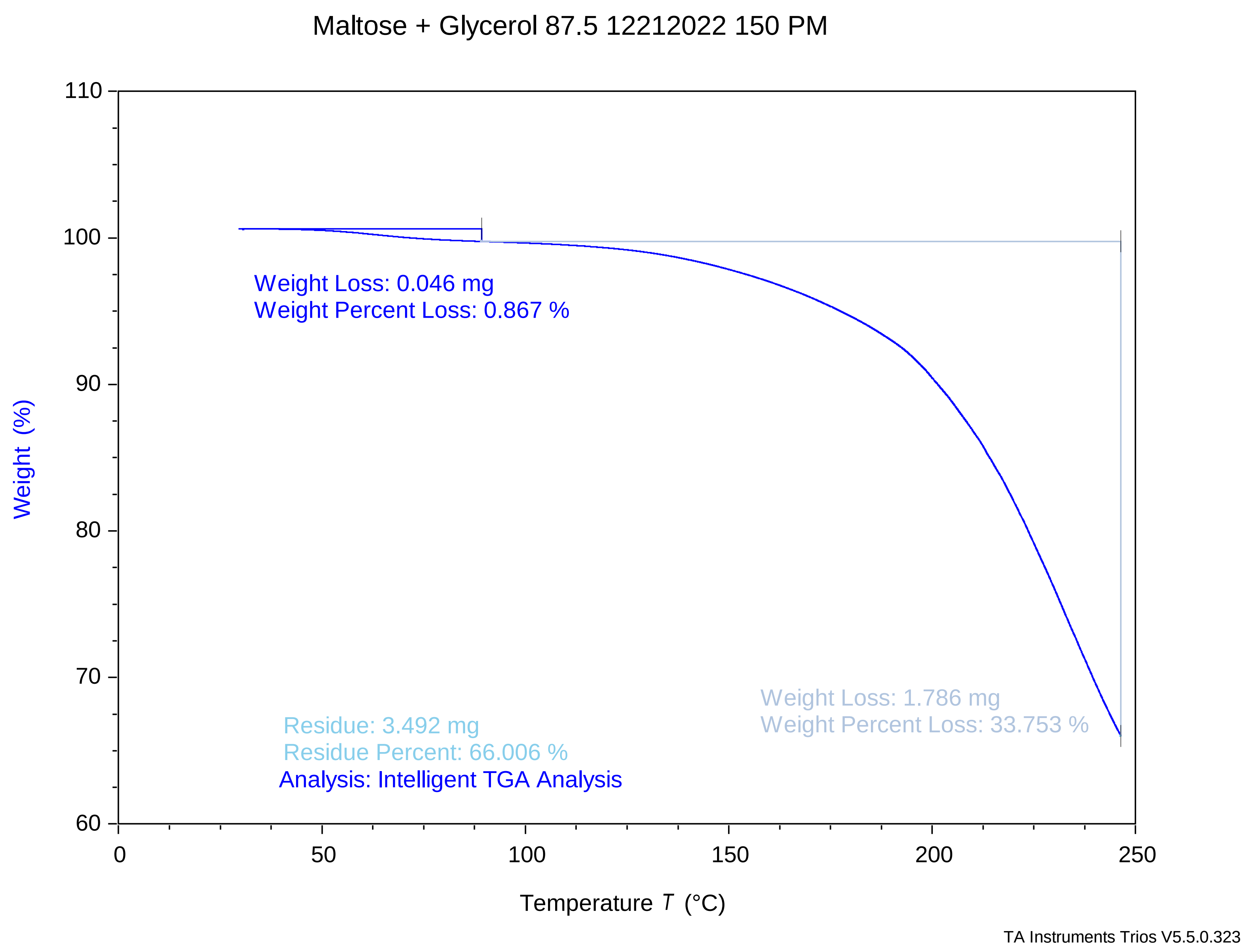

### Maltose + Glycerol 87.5 12282022 1153 AM.pdf

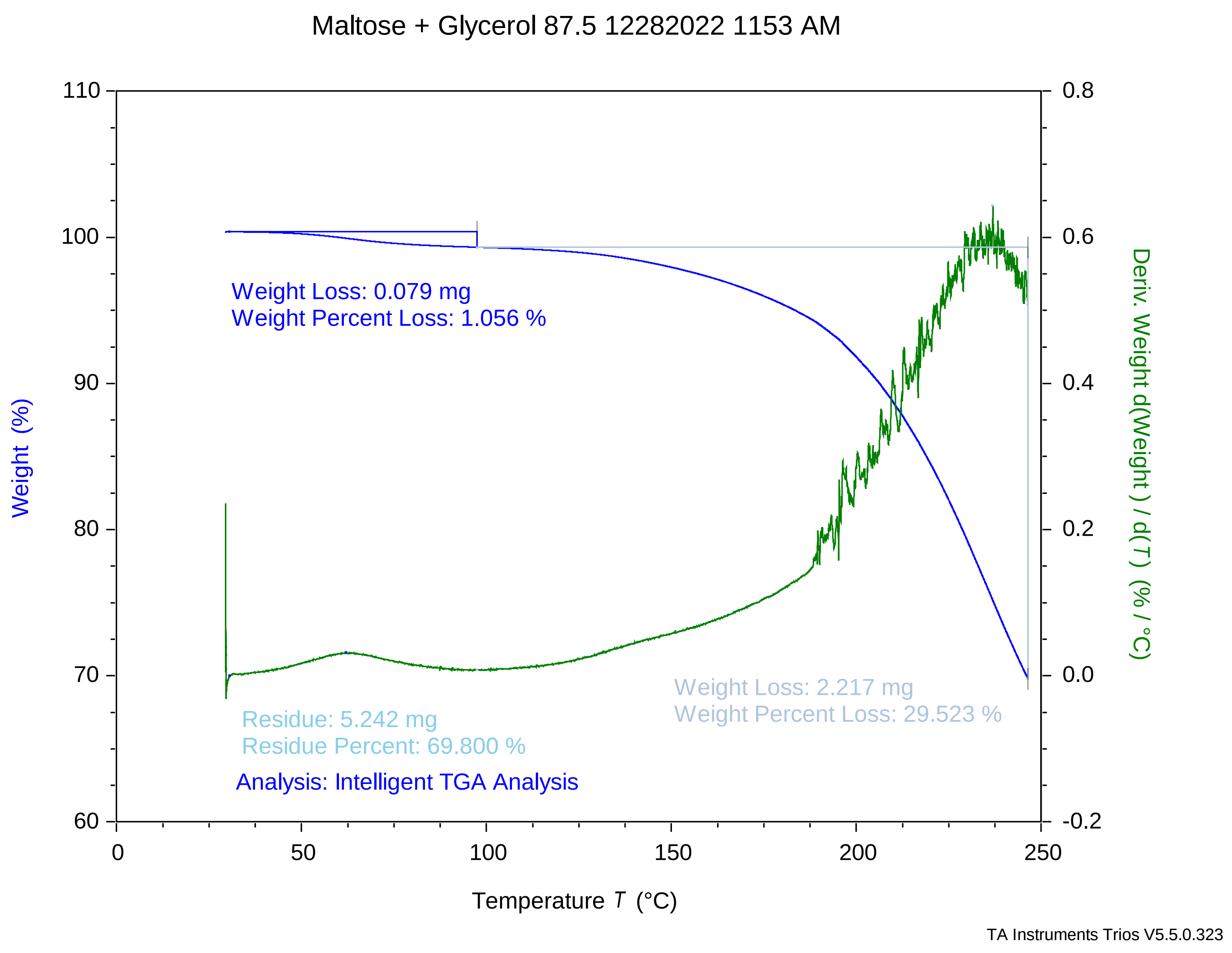

### Maltose + Glycerol 90 12202022 254 PM.pdf

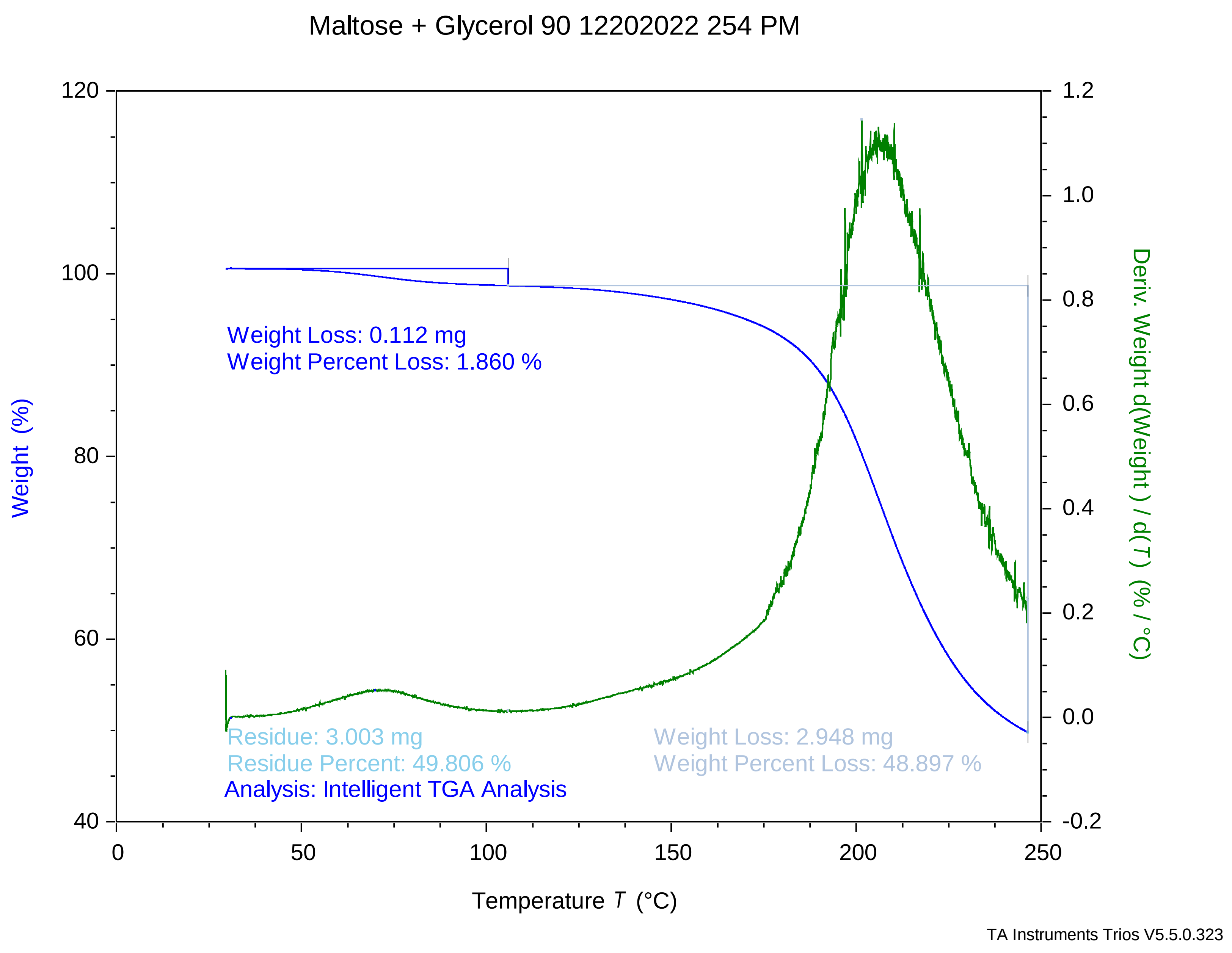

### Maltose + Glycerol 90 12212022 1145 AM.pdf

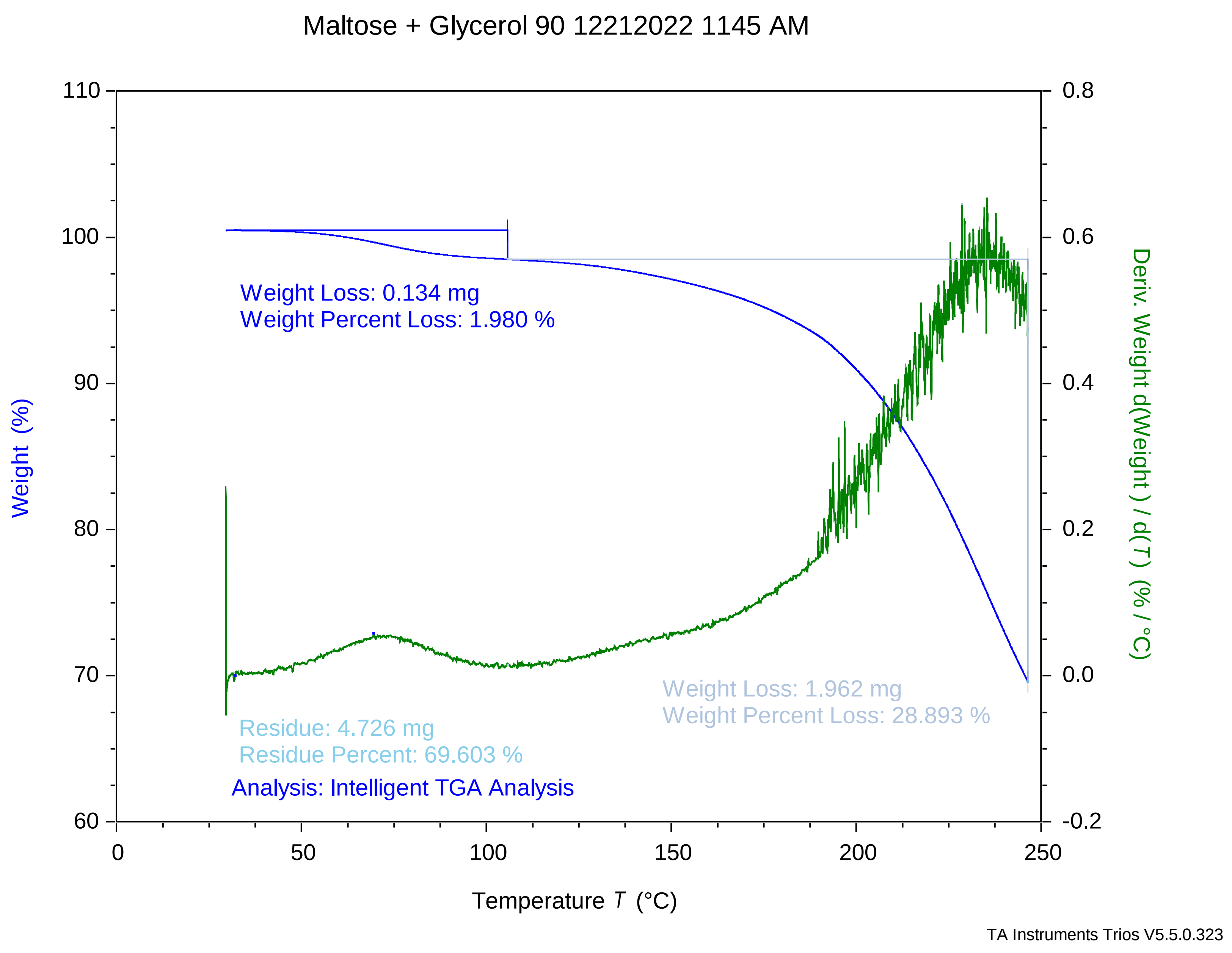

### maltose + glycerol 90 12212022 1246 pm.pdf

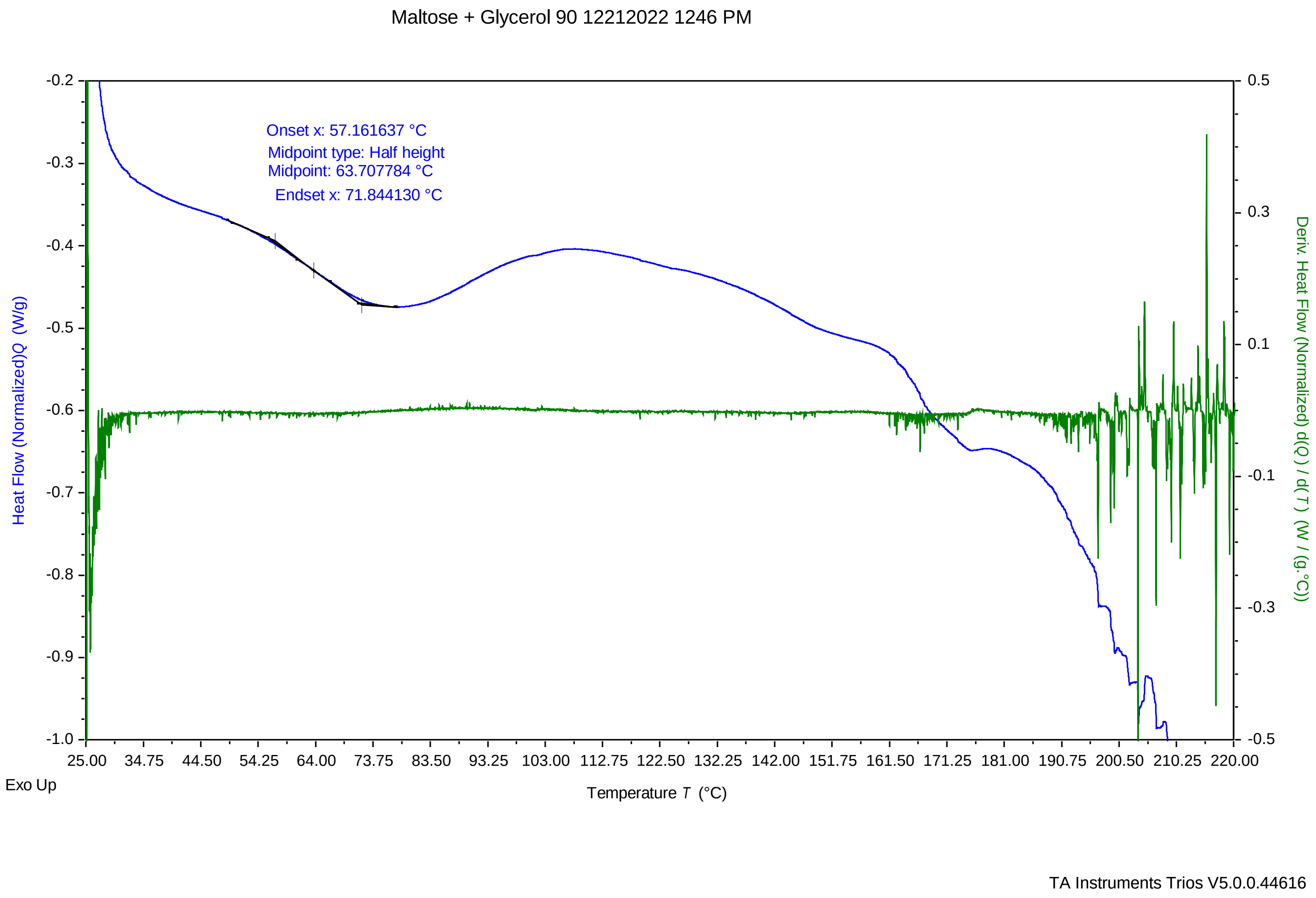

### Maltose + Glycerol 90 12282022 950 AM.pdf

Maltose + Glycerol 90 12282022 950 AM

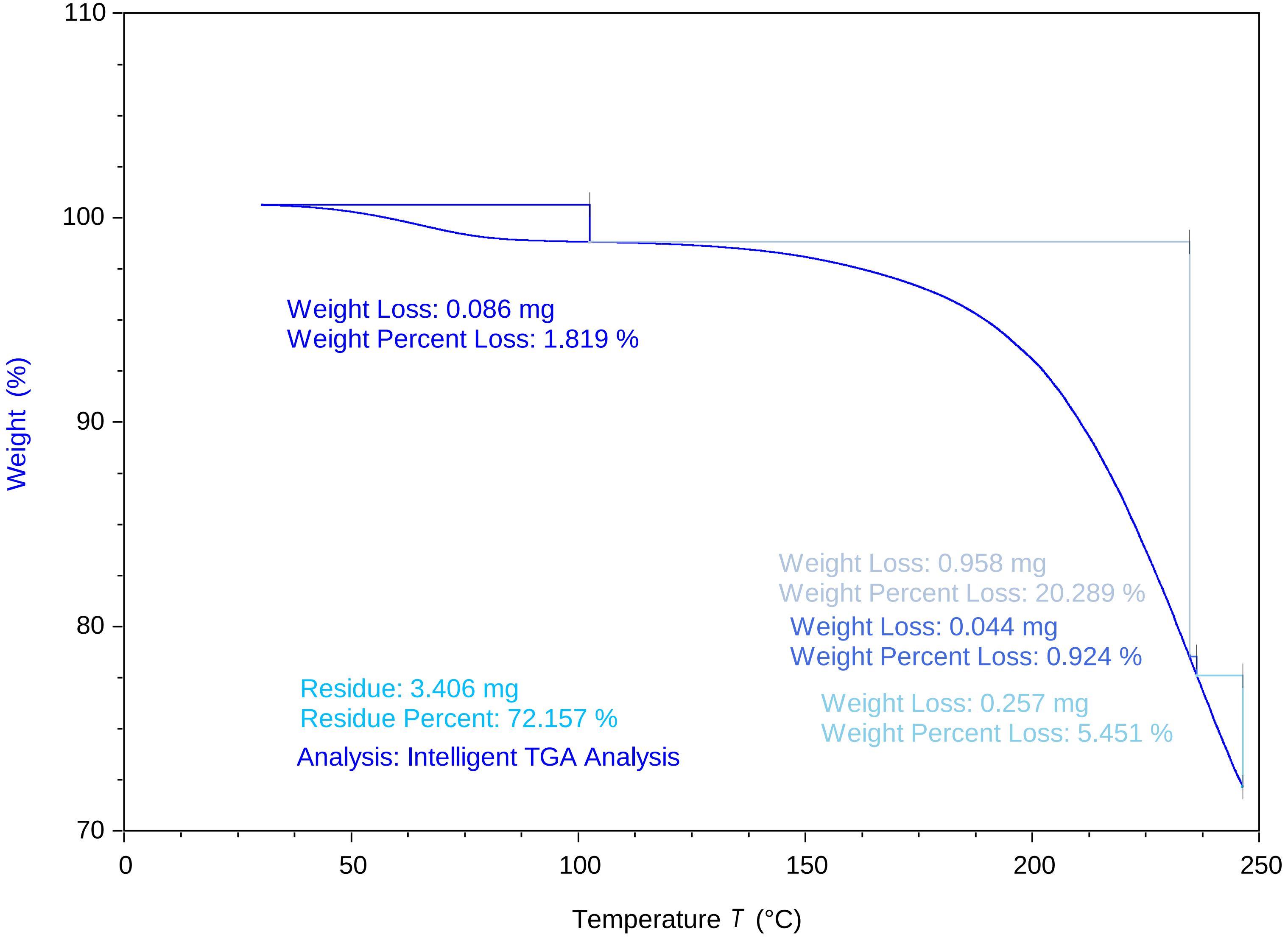


TA Instruments Trios V5.5.0.323

### maltose + glycerol 90 12282022 956 am.pdf

Maltose + Glycerol 90 12282022 956 AM

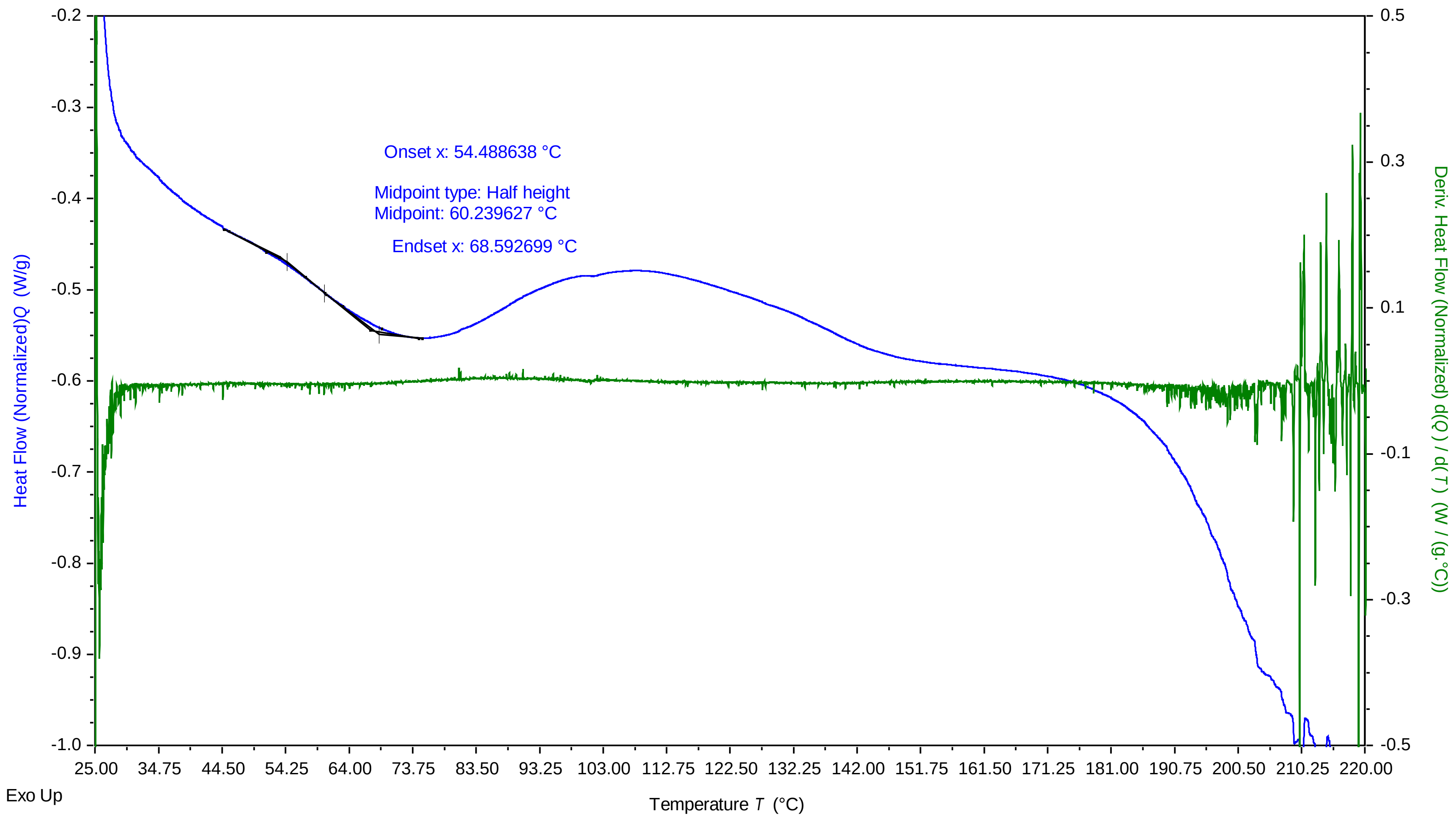


TA Instruments Trios V5.0.0.44616

### Maltose + Glycerol 92.5 12202022 326 PM.pdf

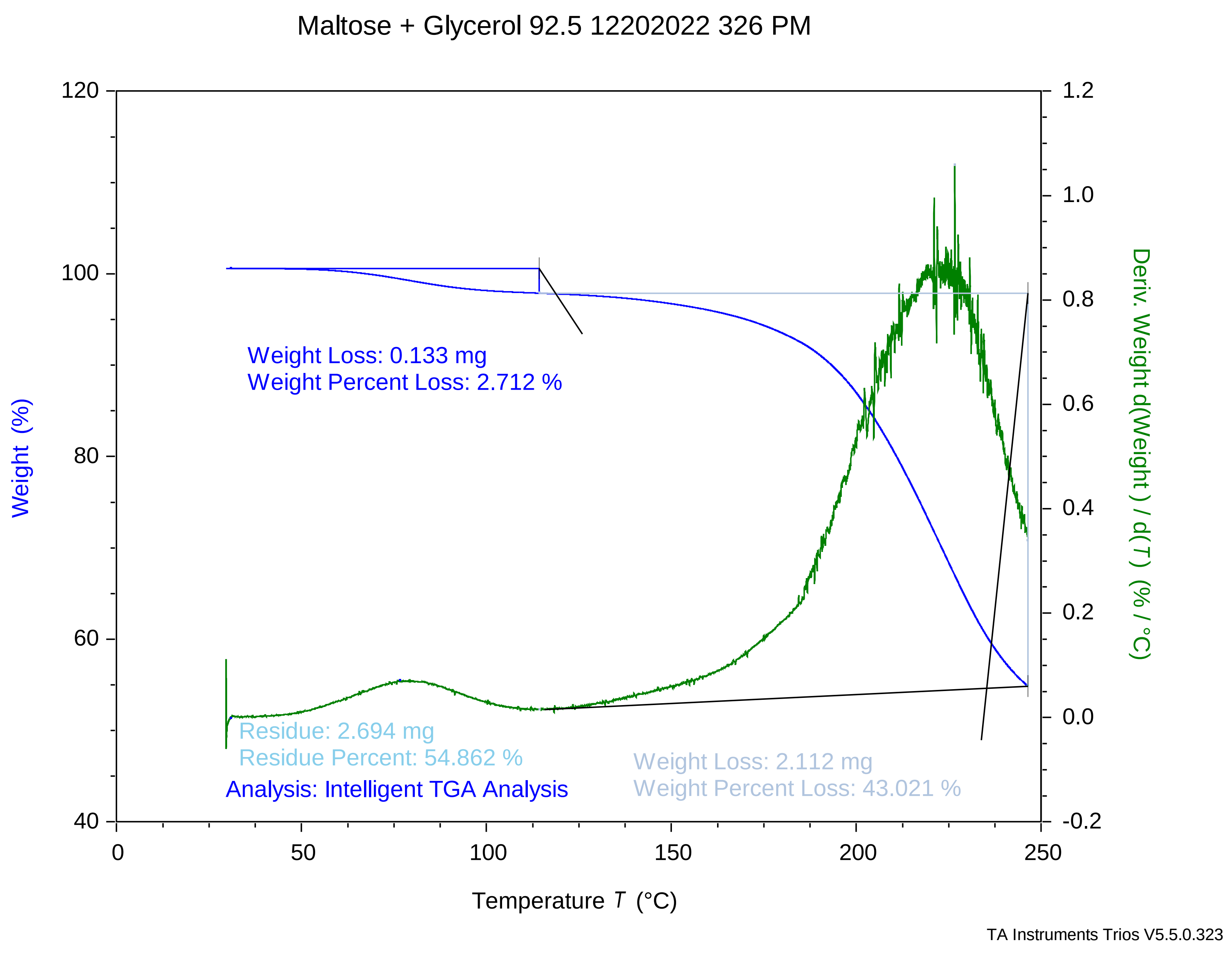

### maltose + glycerol 92.5 12212022 117 pm.pdf

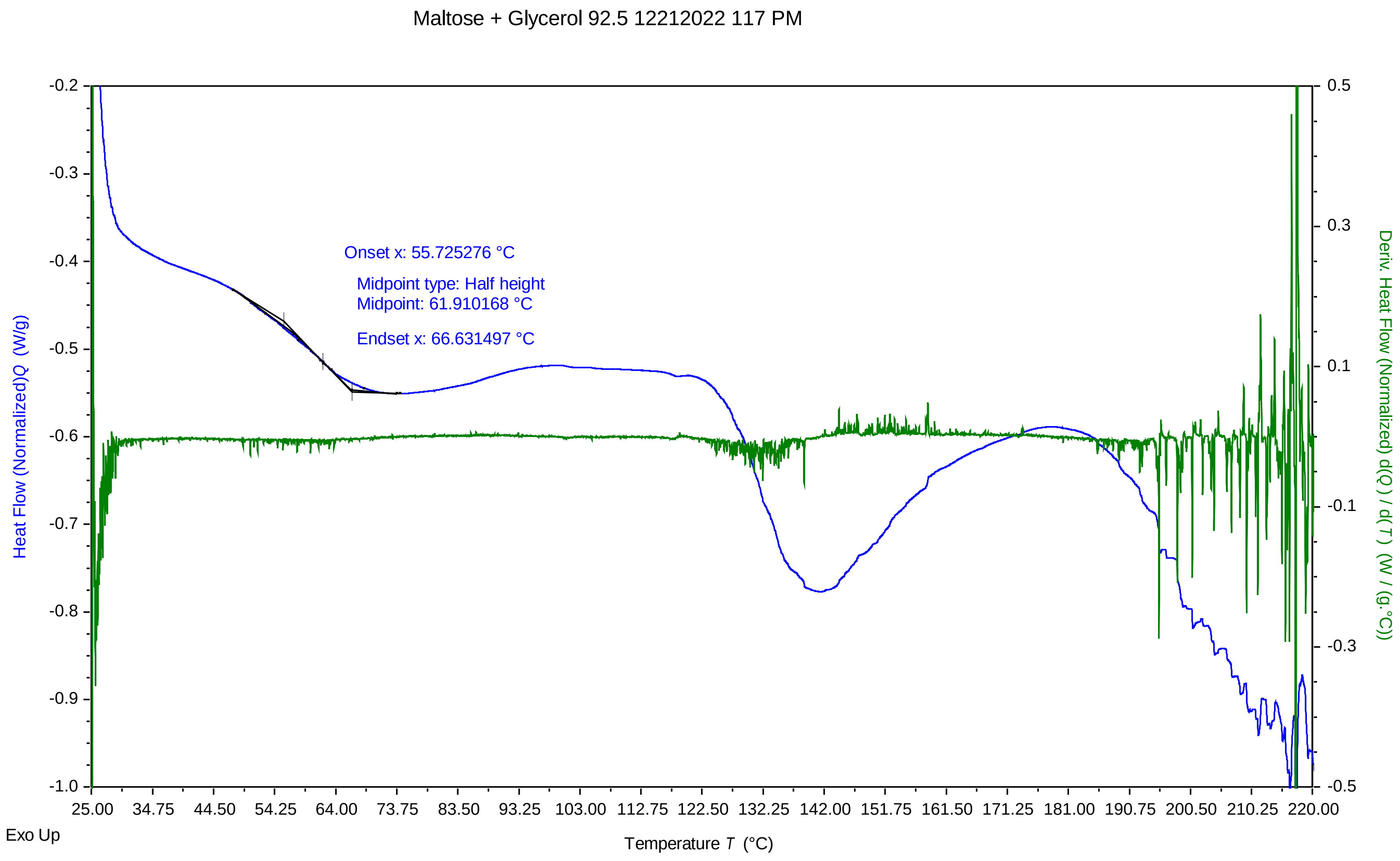


TA Instruments Trios V5.0.0.44616

### Maltose + Glycerol 92.5 12212022 1217 PM.pdf

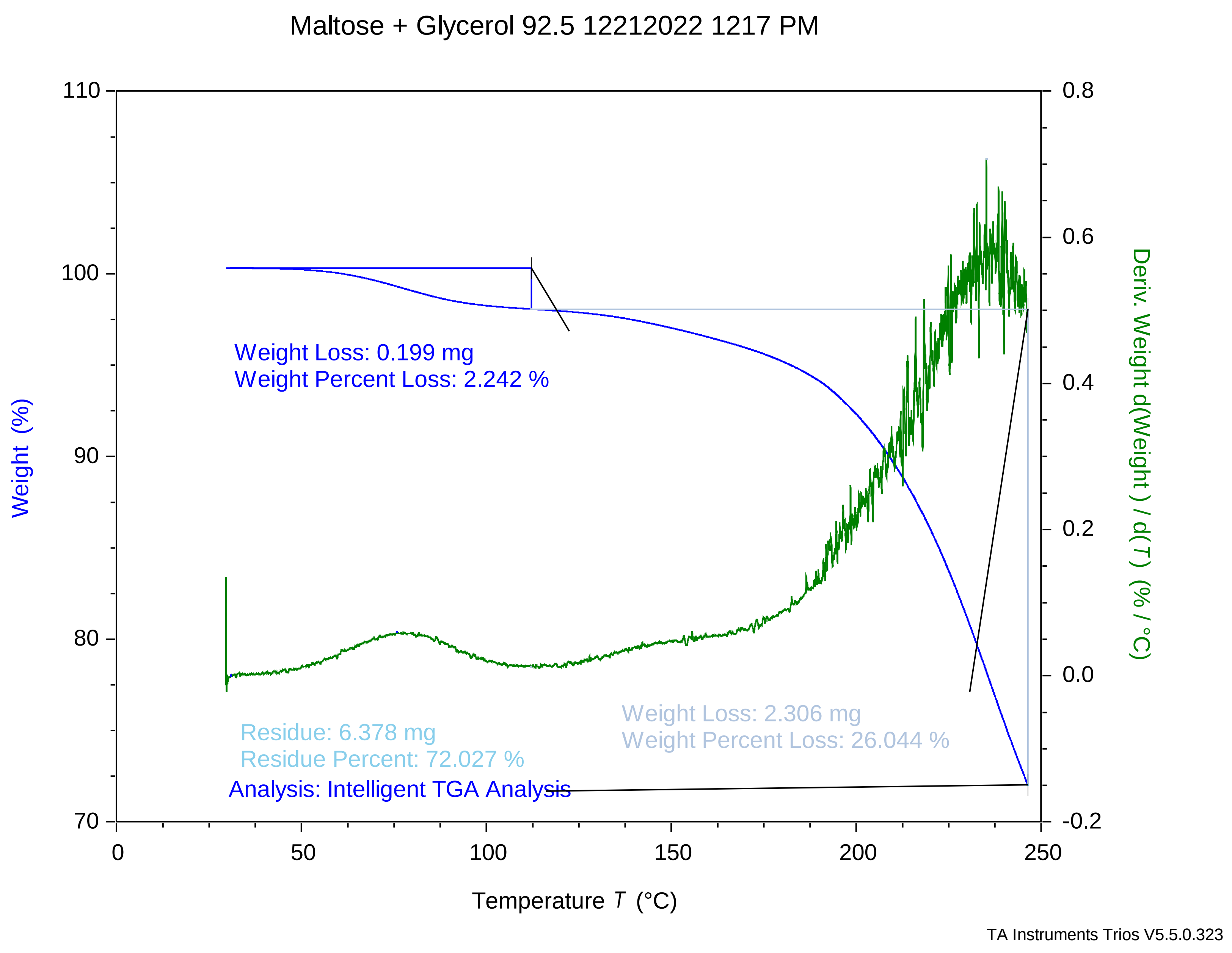

### Maltose + Glycerol 92.5 12282022 1021 AM.pdf

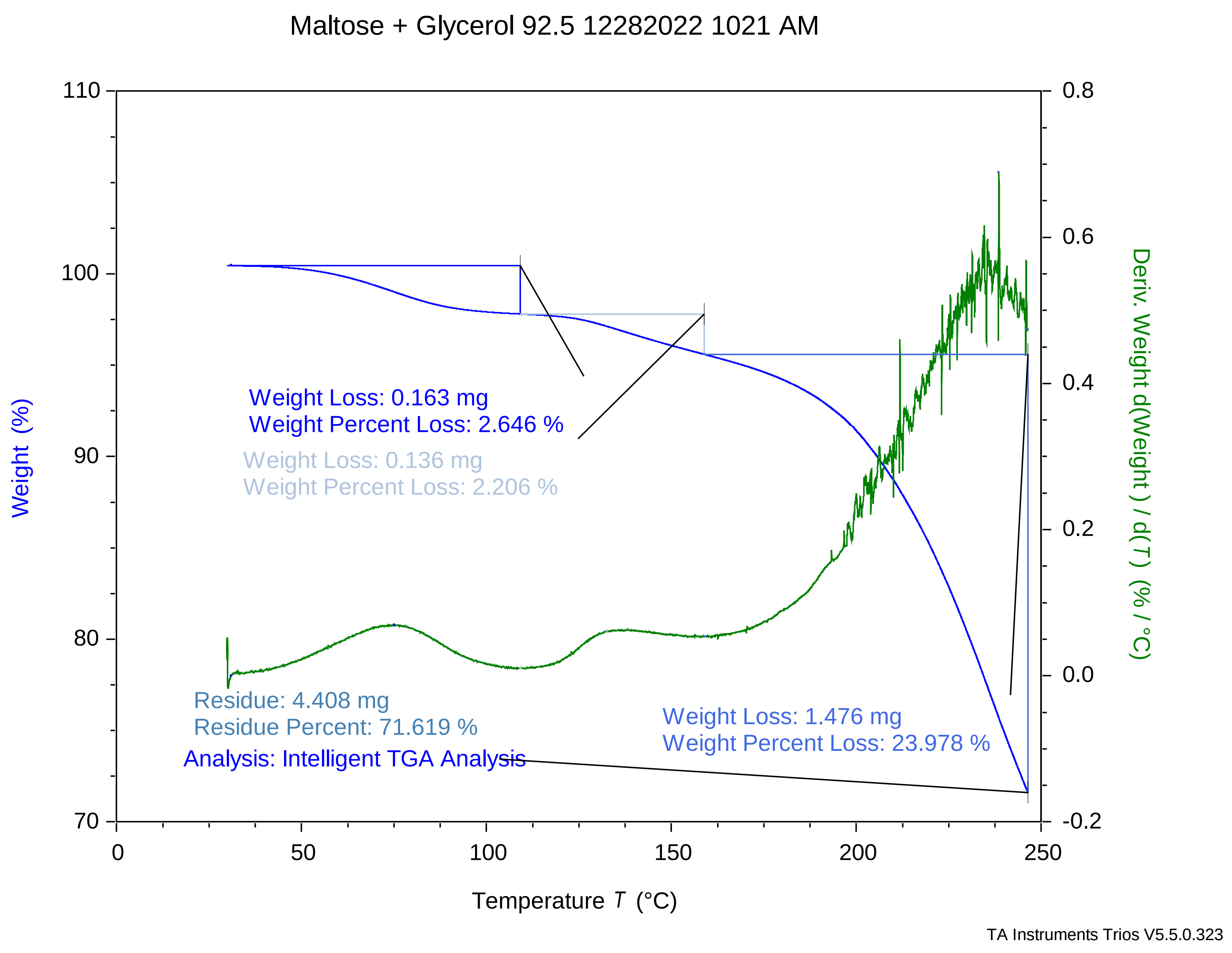

### Maltose + Glycerol 95 12202022 224 PM.pdf

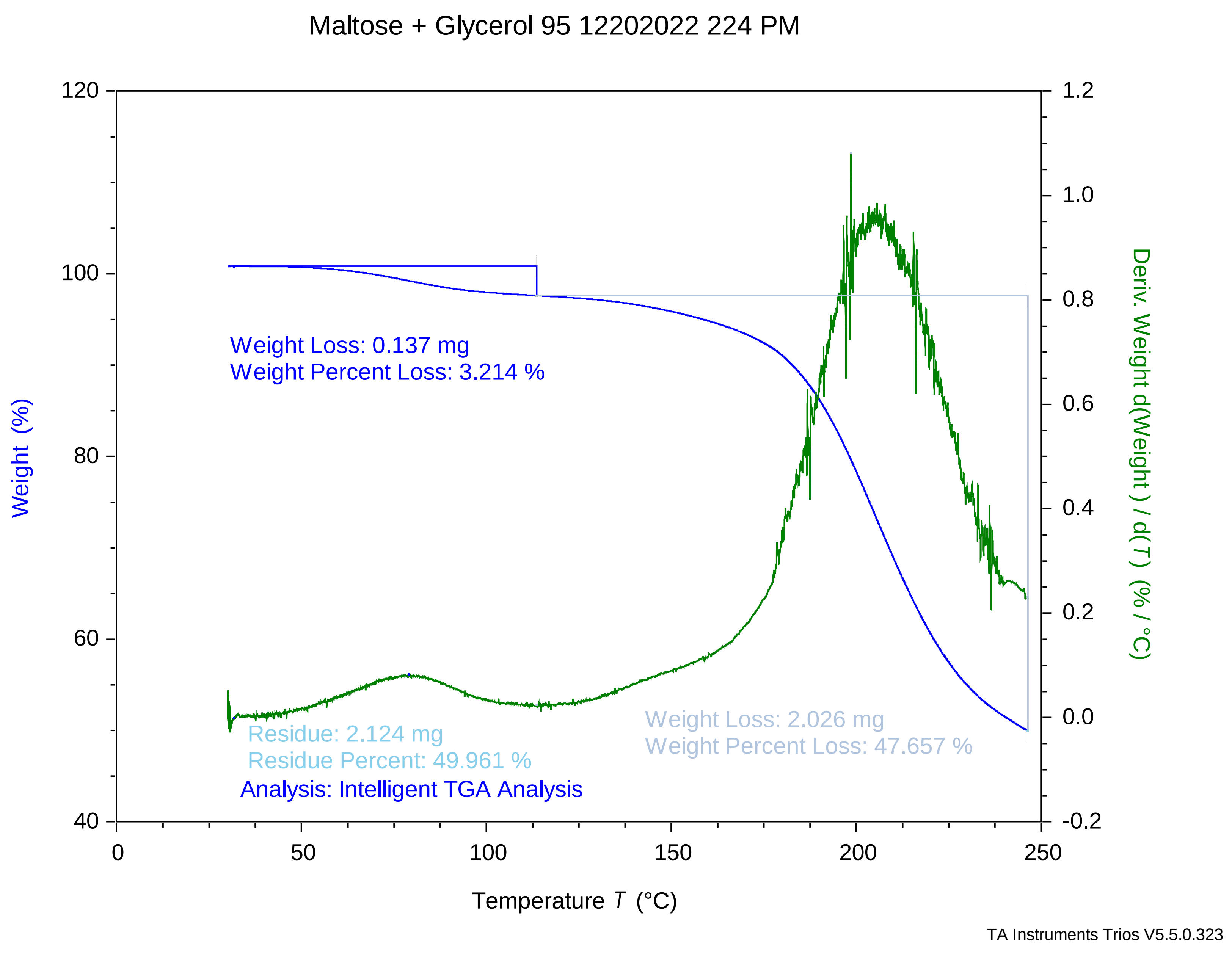

### maltose + glycerol 95 12202022 238 pm.pdf

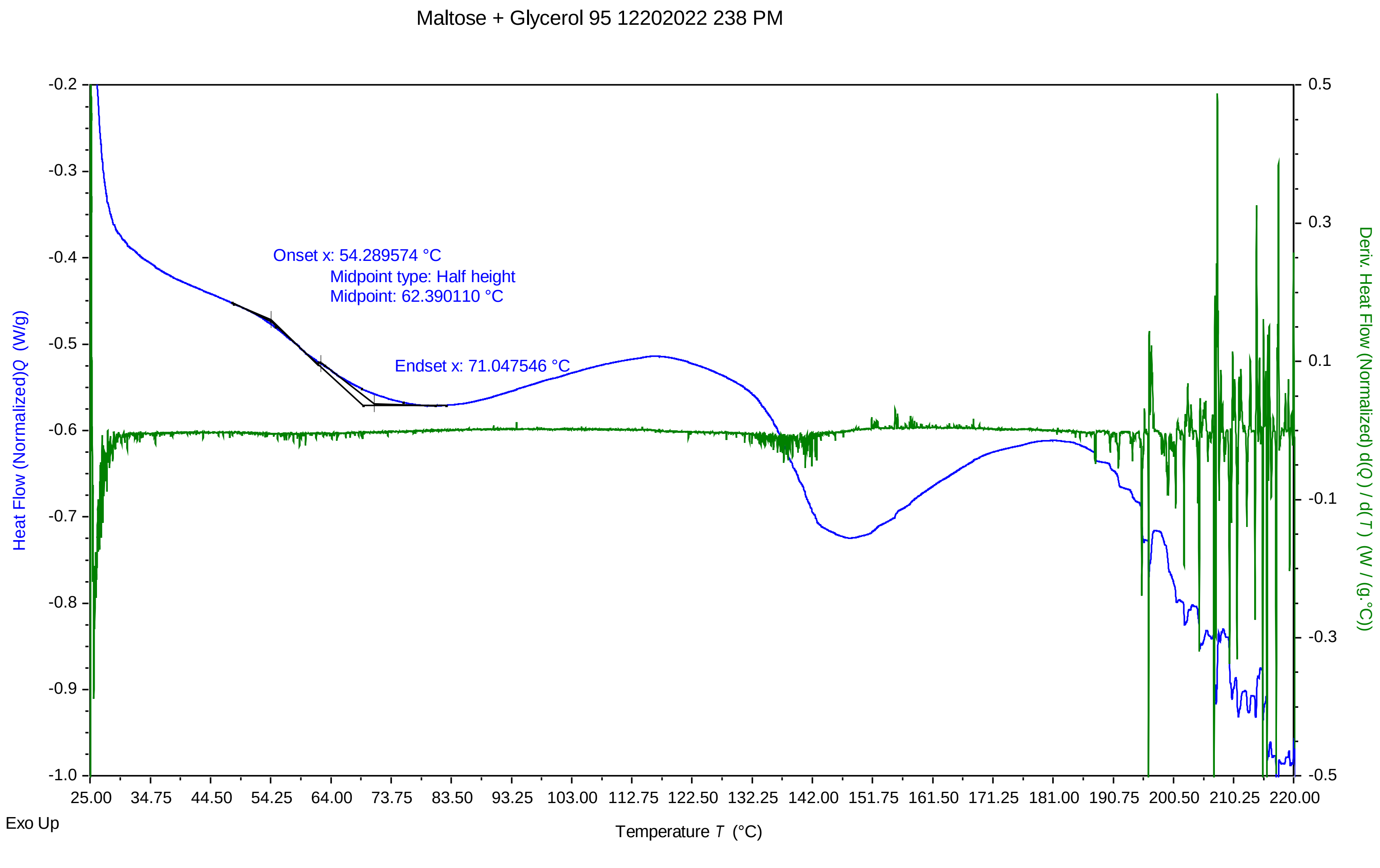


TA Instruments Trios V5.0.0.44616

### Maltose + Glycerol 95 12212022 1116 AM.pdf

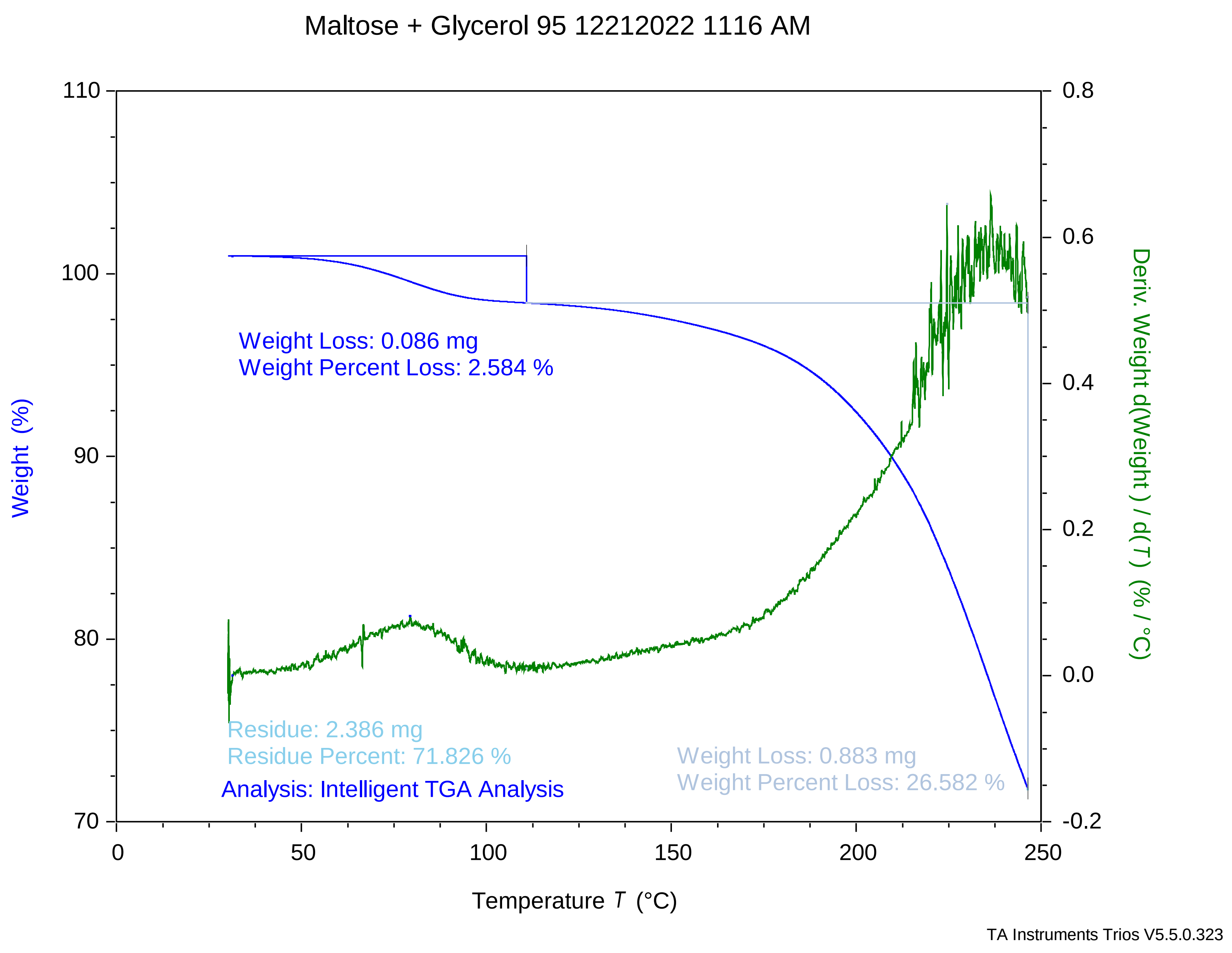

### maltose + glycerol 95 12212022 1215 pm.pdf

Maltose + Glycerol 95 12212022 1215 PM

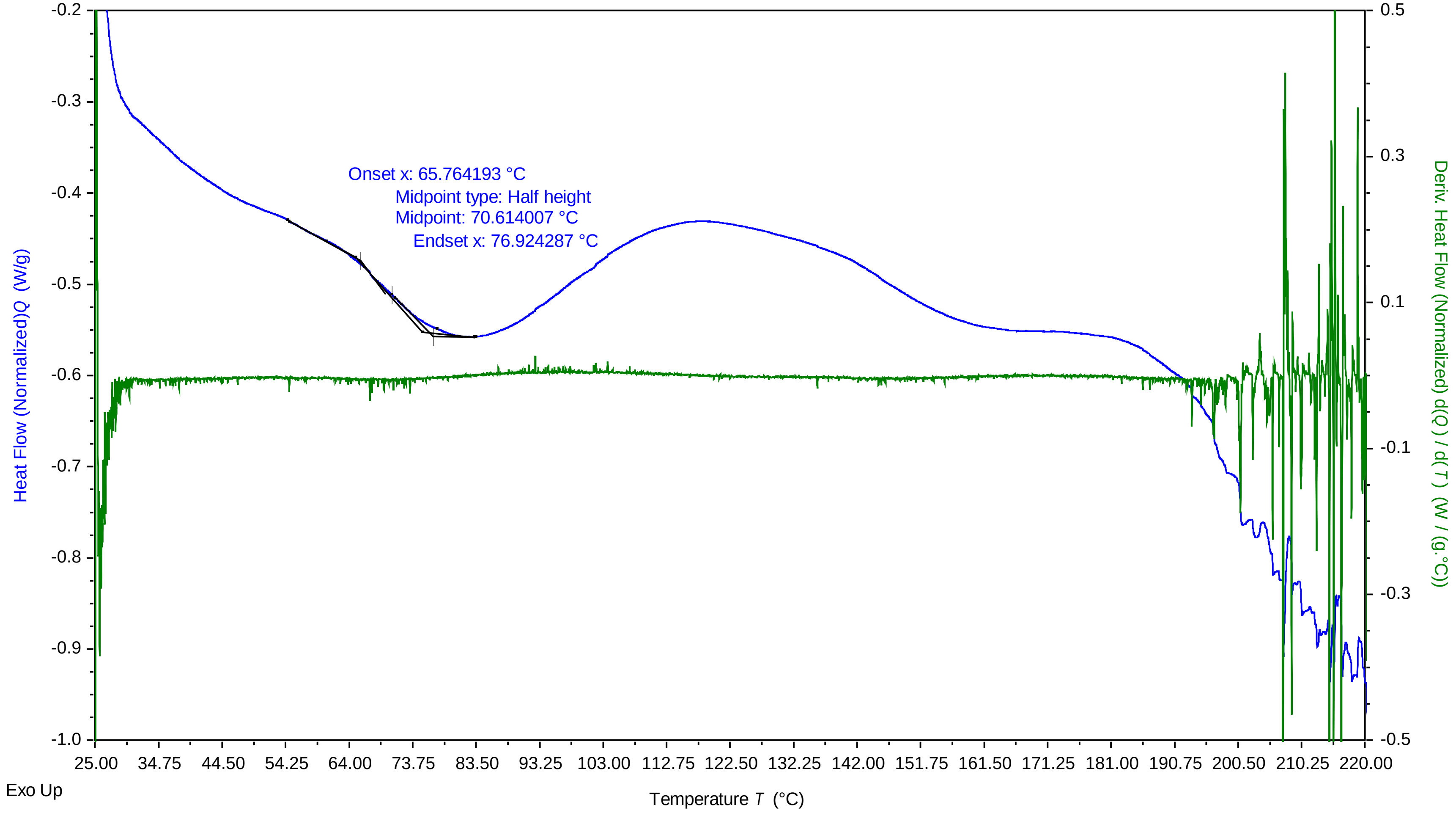


TA Instruments Trios V5.0.0.44616

### Maltose + Glycerol 97.5 12282022 1121 AM.pdf

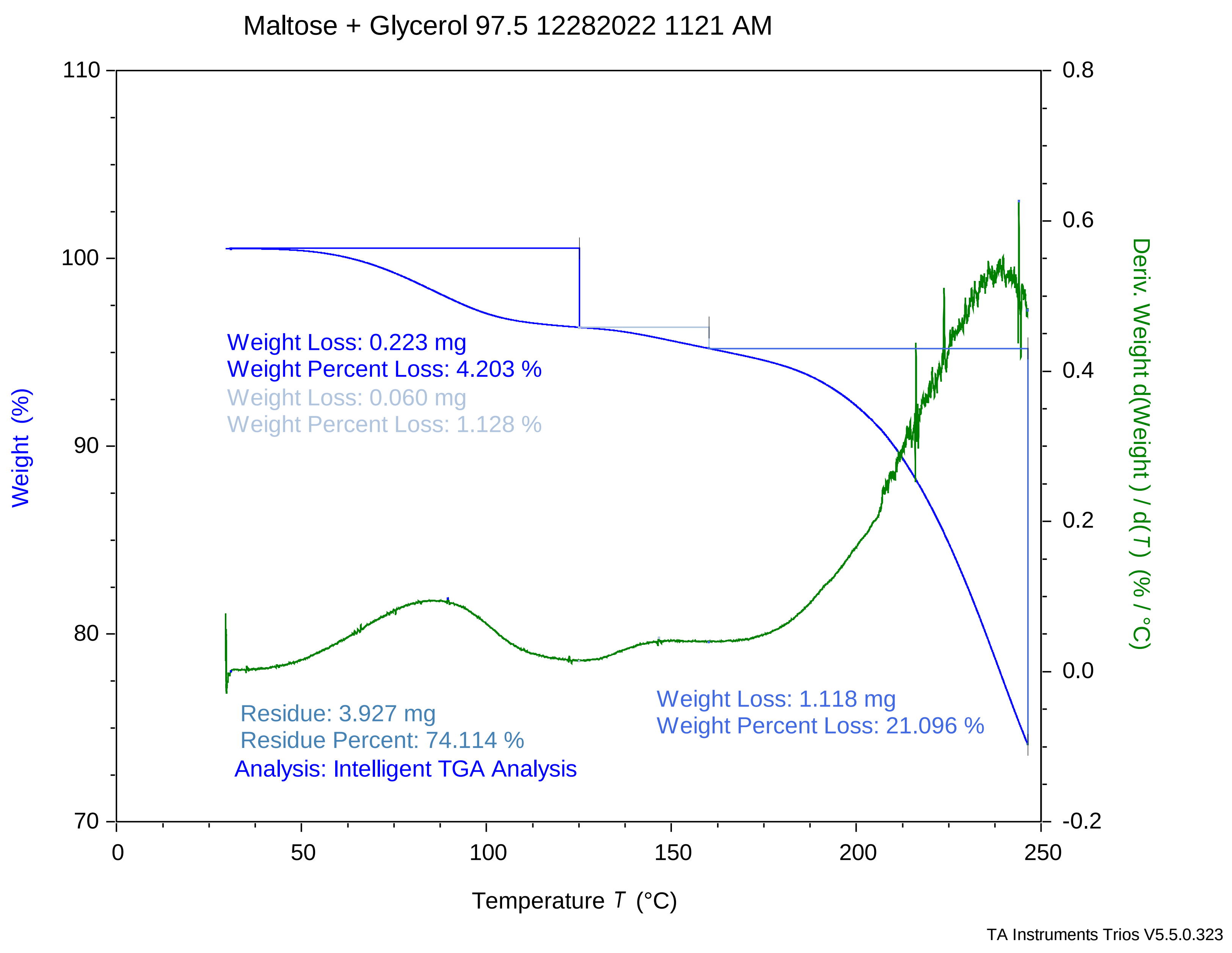

### Maltose + Glycerol 100 12202022 358 PM.pdf

Maltose + Glycerol 100 12202022 358 PM

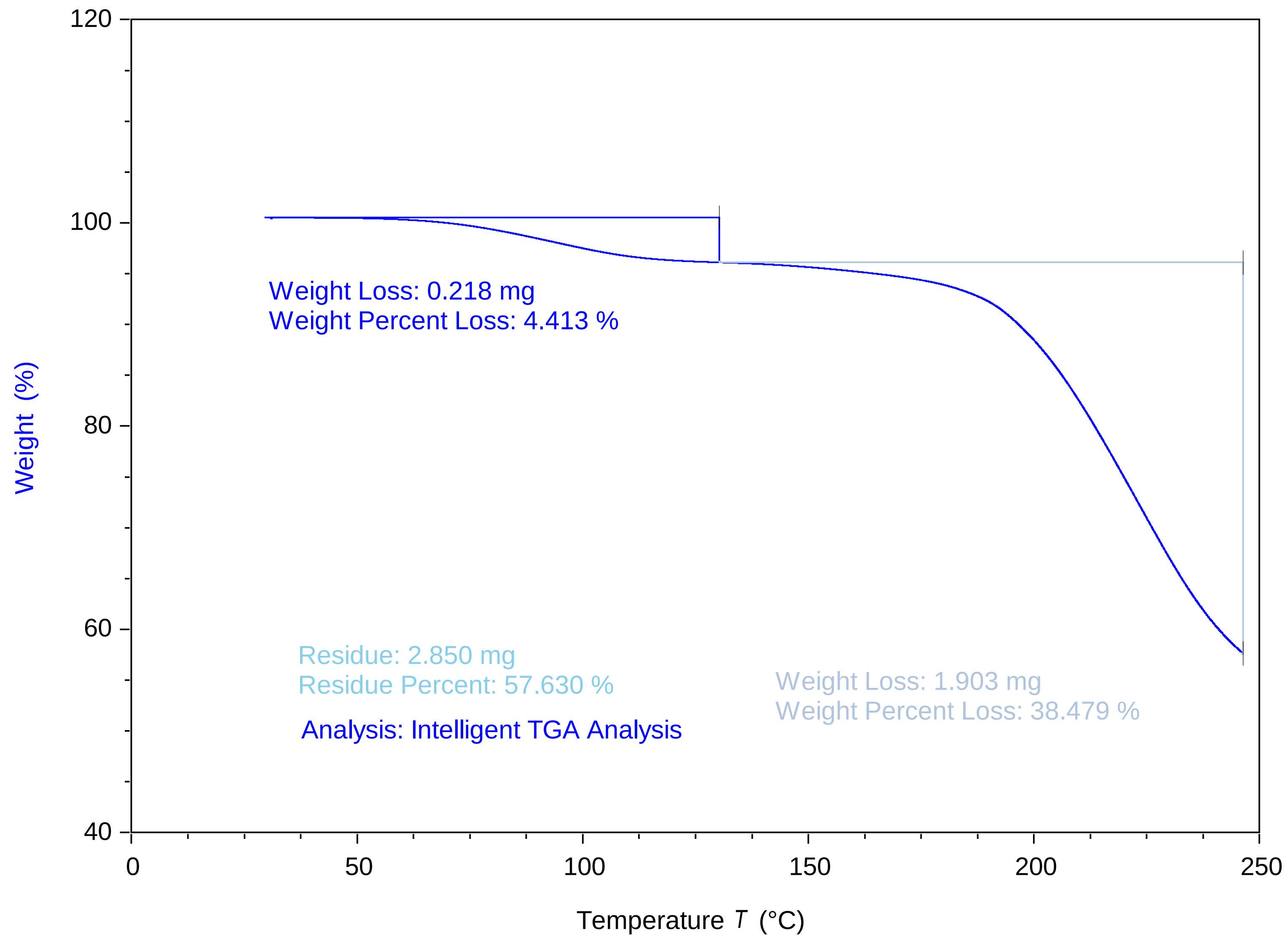


TA Instruments Trios V5.5.0.323

### maltose + glycerol 100 12202022 411 pm.pdf

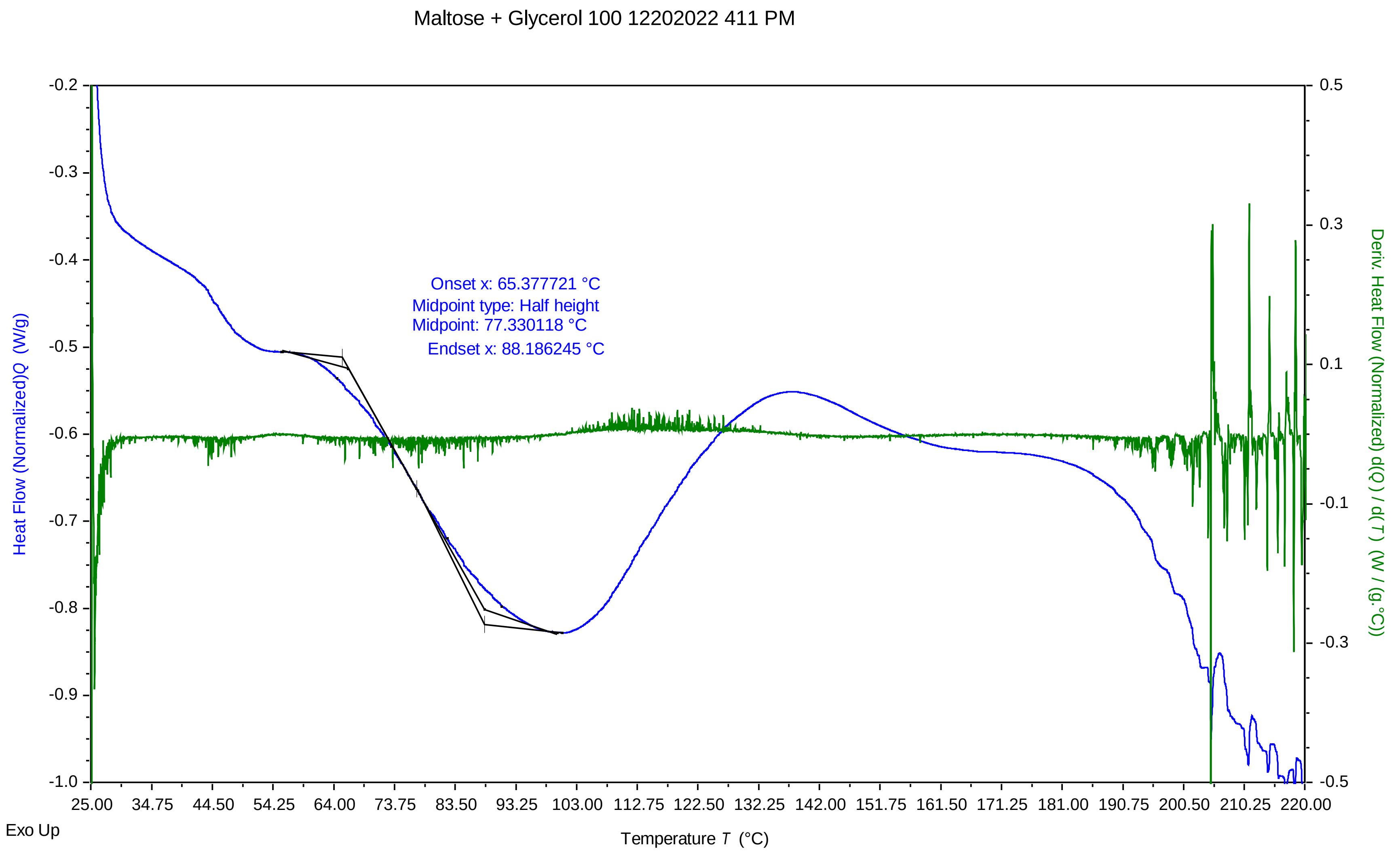


TA Instruments Trios V5.0.0.44616

### maltose + glycerol 100 12212022 148 pm.pdf

Maltose + Glycerol 100 12212022 148 PM

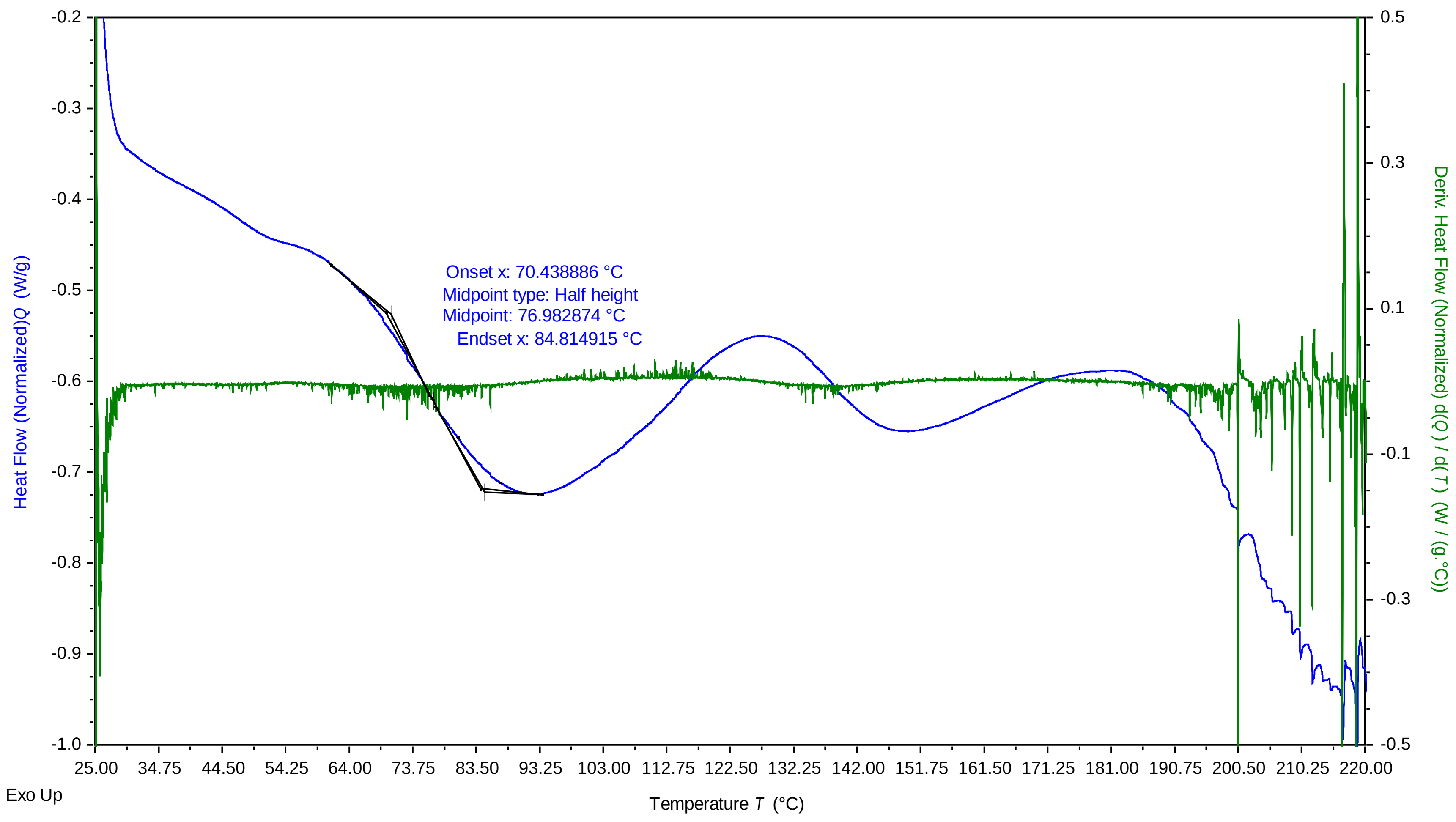


TA Instruments Trios V5.0.0.44616

### Maltose + Glycerol 100 12212022 1249 PM.pdf

Maltose + Glycerol 100 12212022 1249 PM

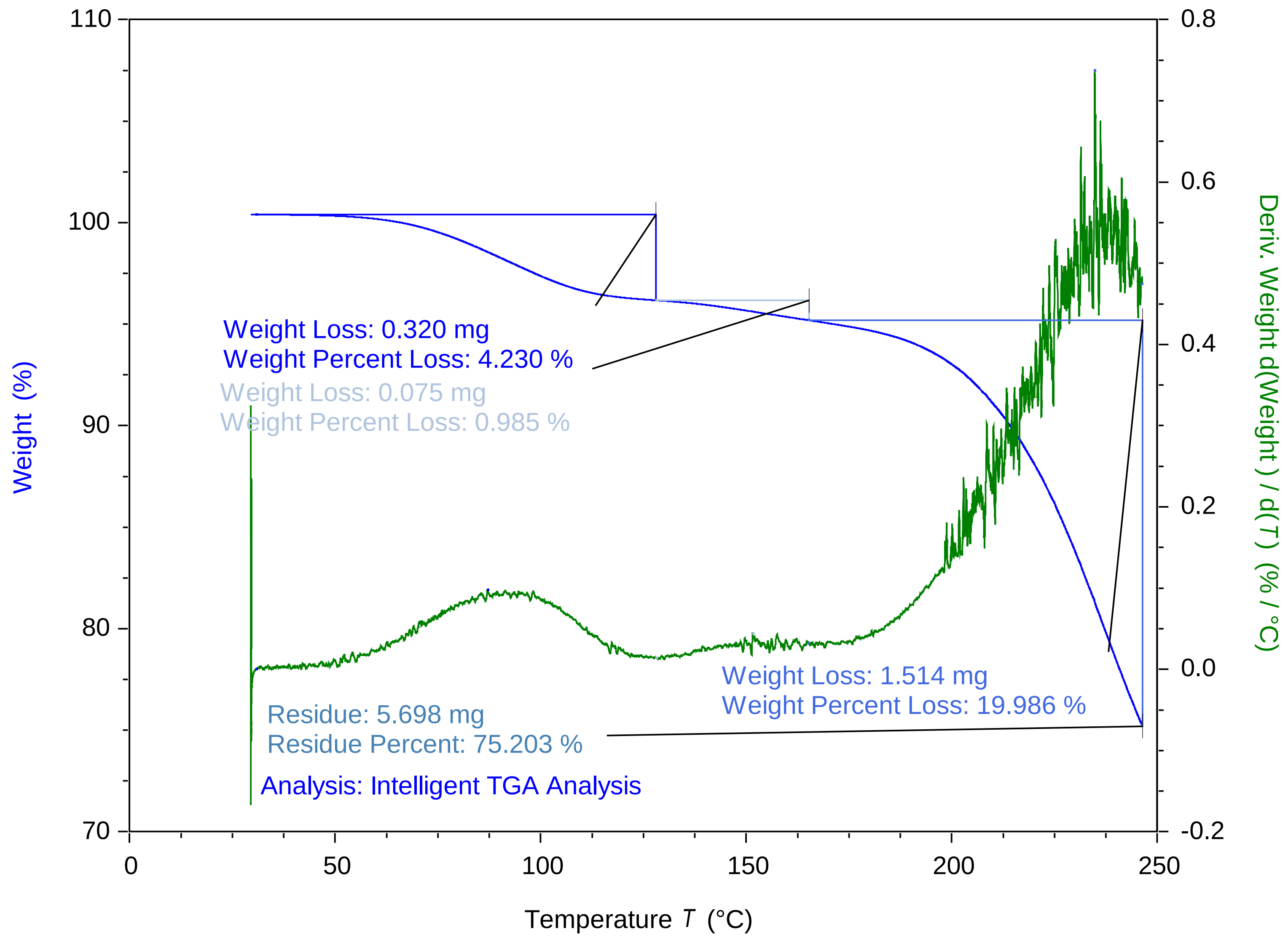


TA Instruments Trios V5.5.0.323

### Maltose + Glycerol 100 12282022 1050 AM.pdf

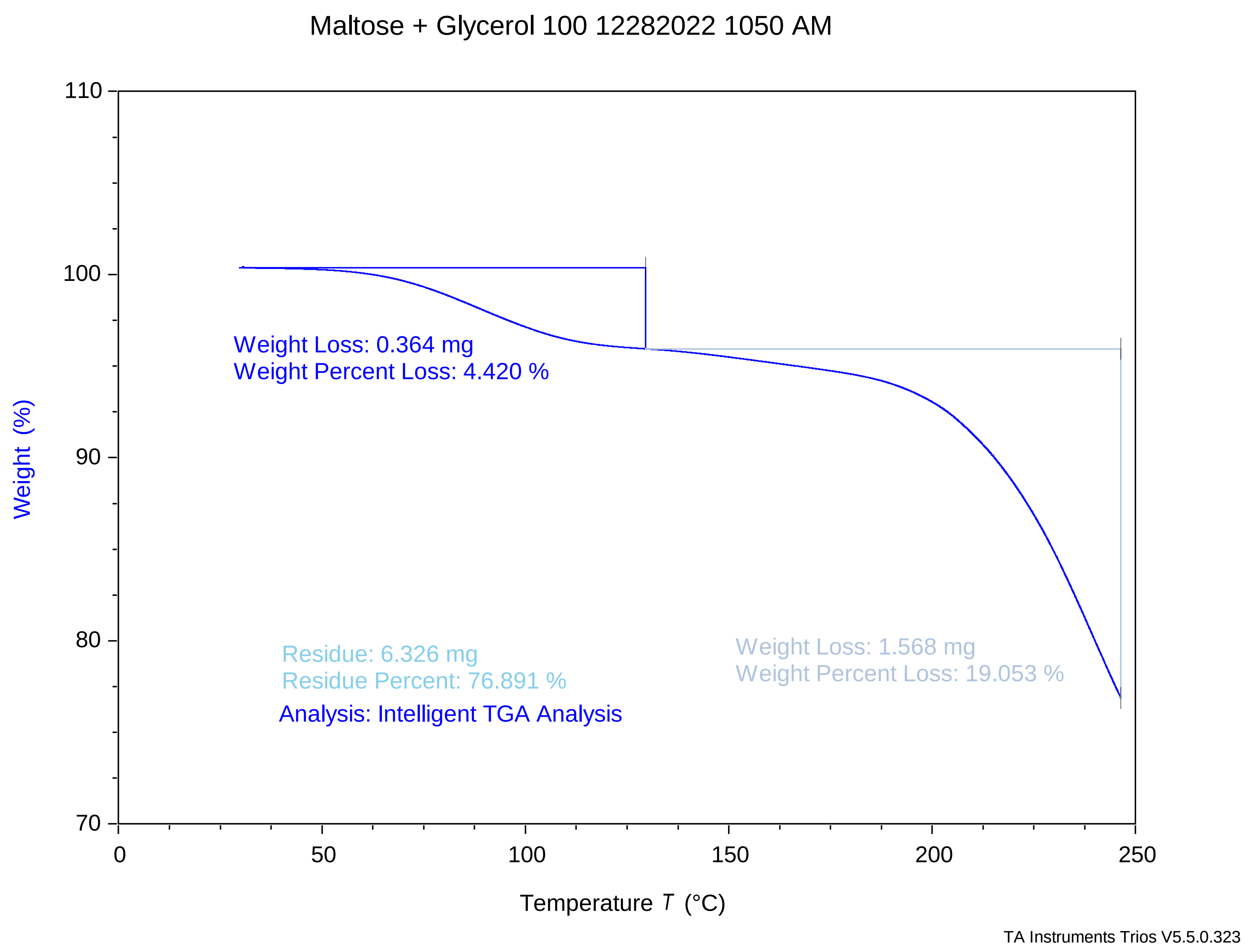

### Maltose + Glycerol 875 12202022 501 PM.pdf

Maltose + Glycerol 87.5 12202022 501 PM

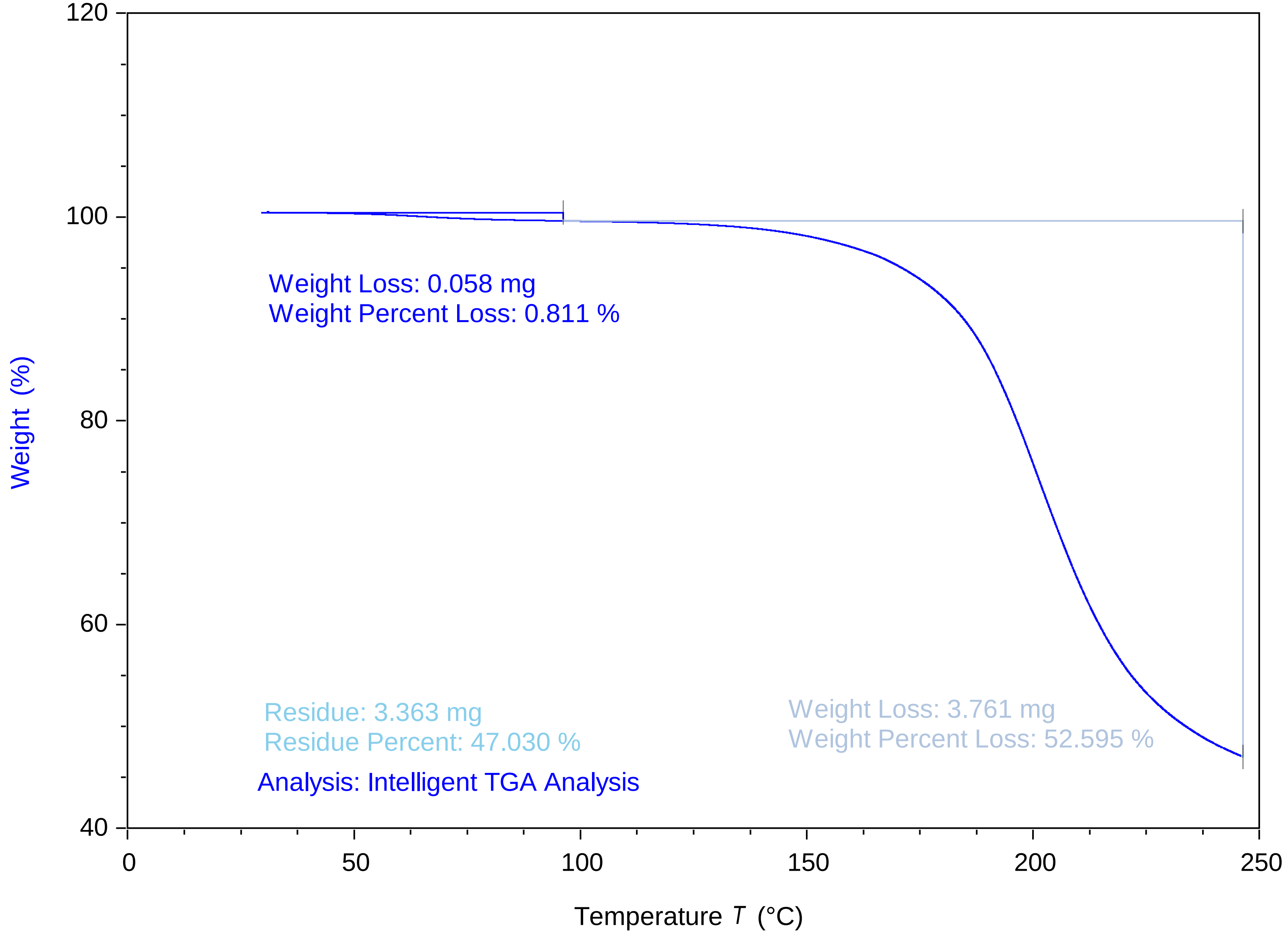


TA Instruments Trios V5.5.0.323

### maltose + glycerol 875 12202022 513 pm.pdf

Maltose + Glycerol 87.5 12202022 513 PM

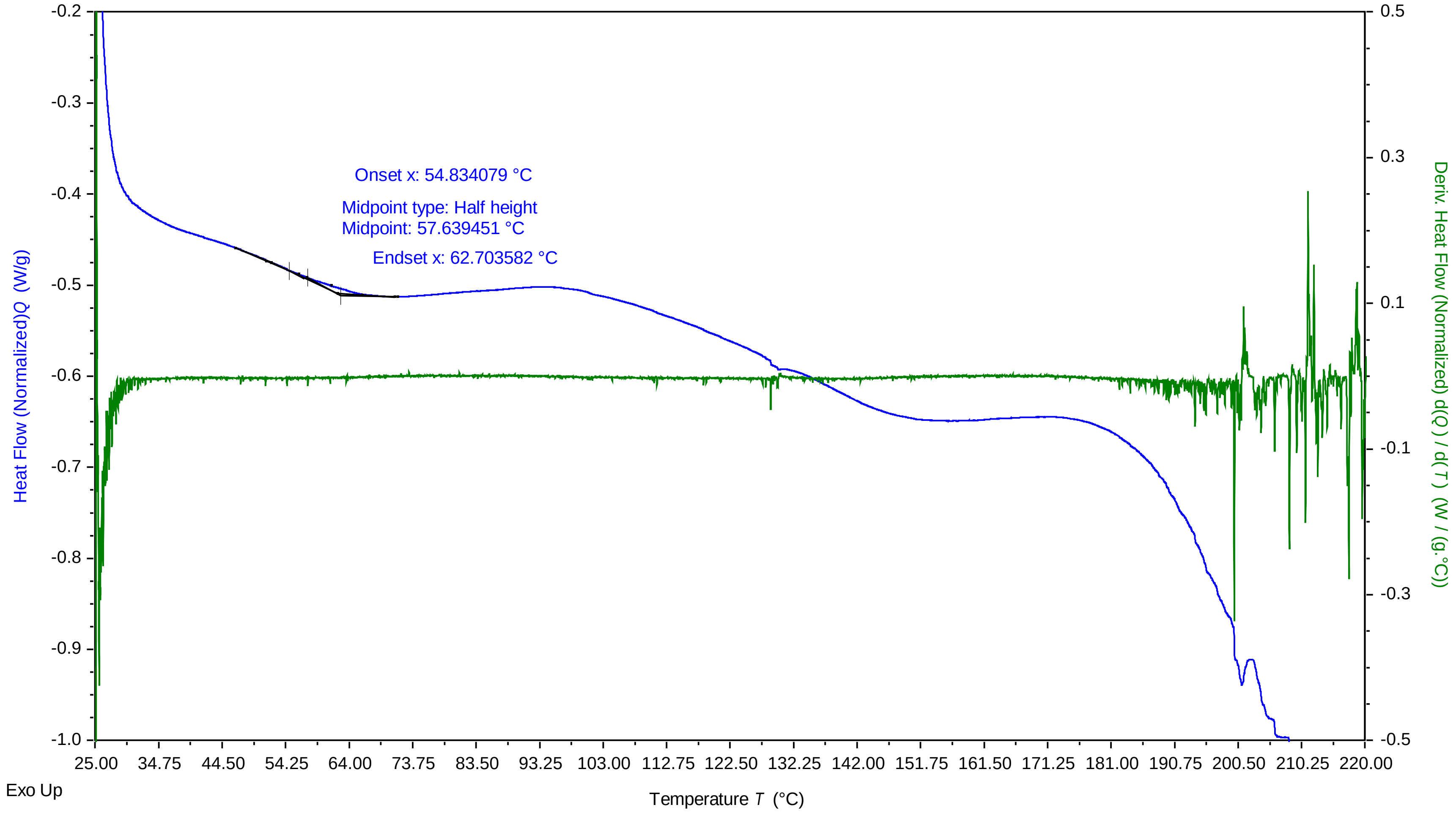


TA Instruments Trios V5.0.0.44616

### maltose + glycerol 875 12212022 250 pm.pdf

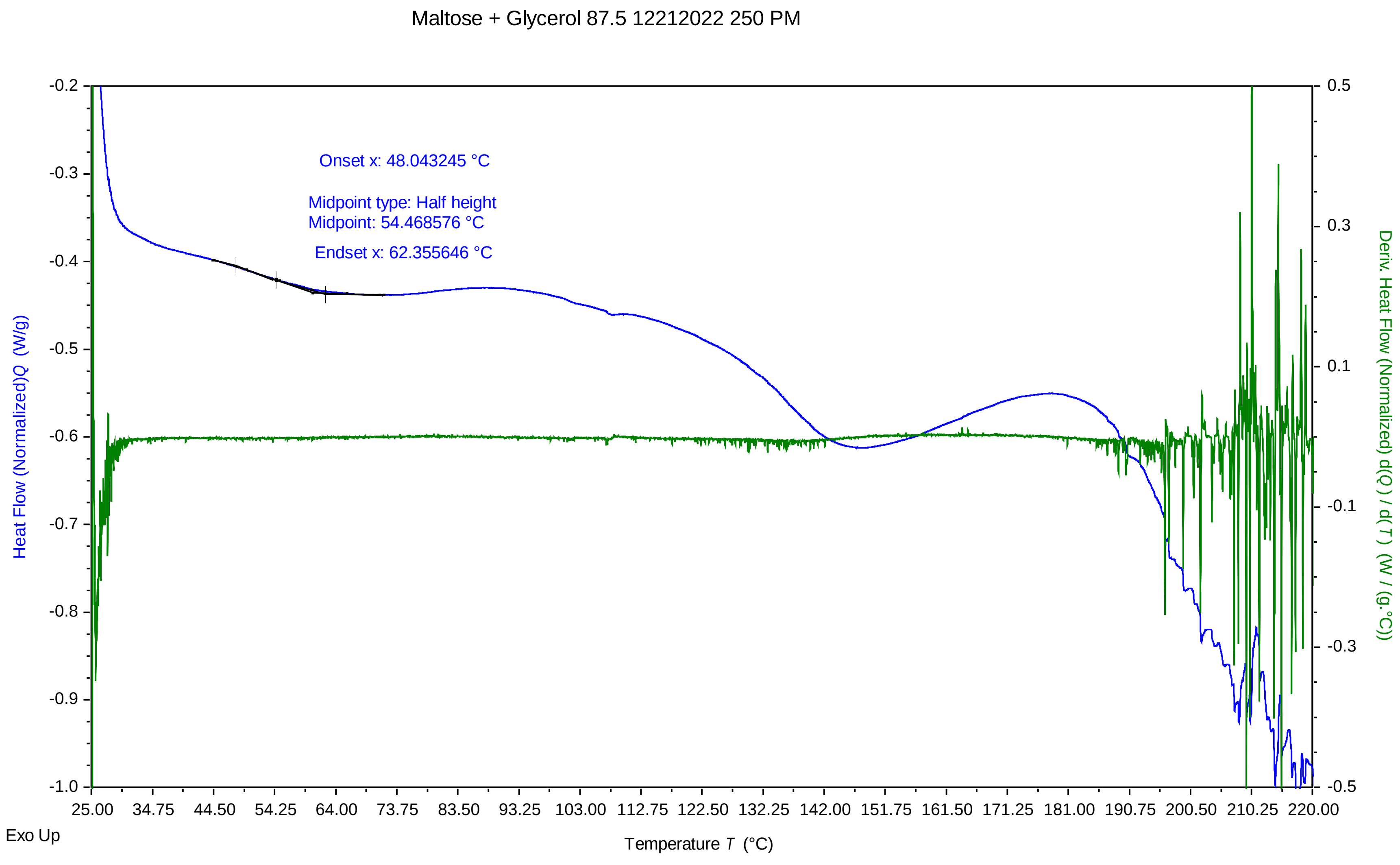


TA Instruments Trios V5.0.0.44616

### maltose + glycerol 925 12282022 1026 am.pdf

Maltose + Glycerol 92.5 12282022 1026 AM

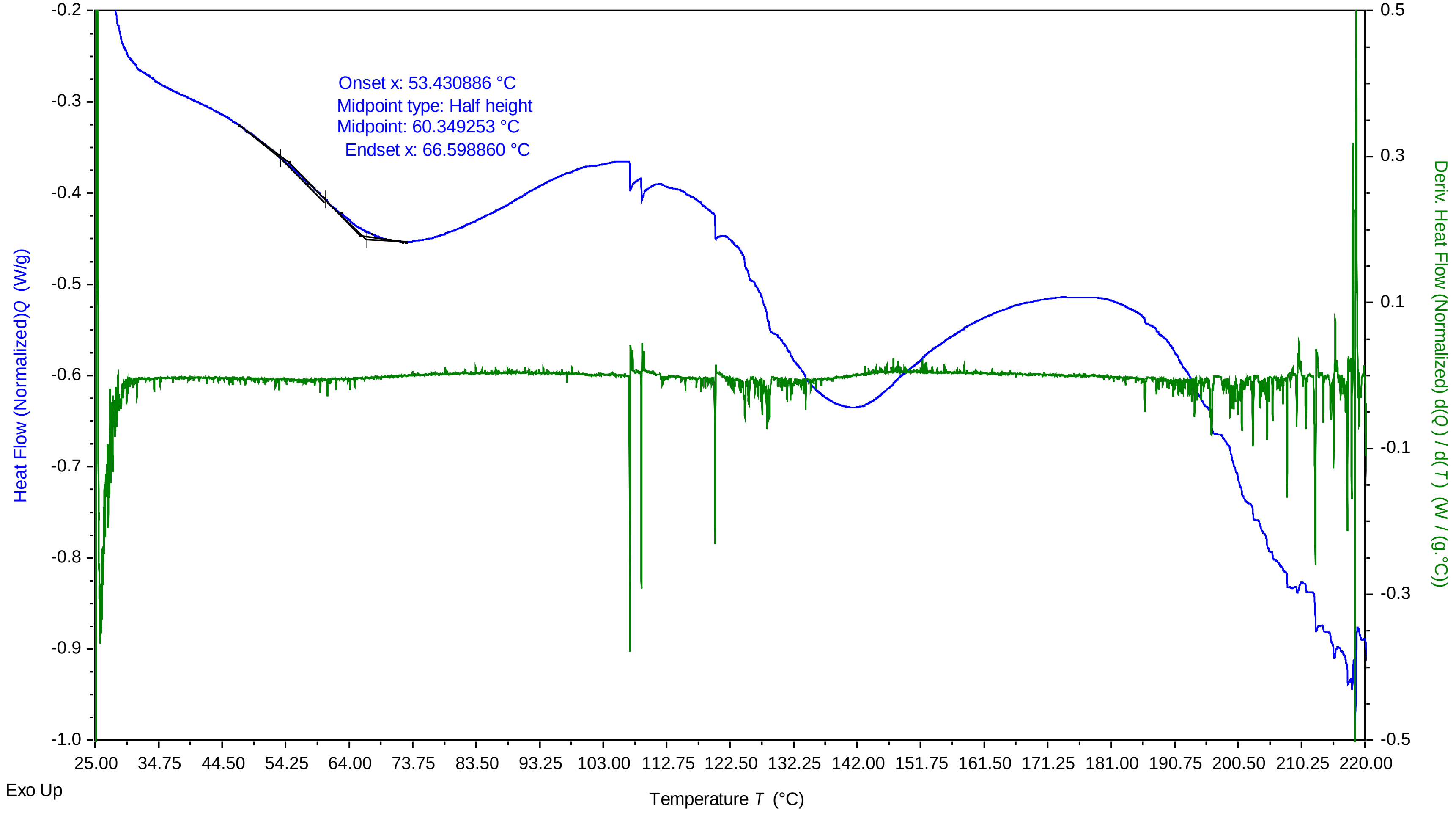


TA Instruments Trios V5.0.0.44616

### Maltose + Glycerol 975 12202022 429 PM.pdf

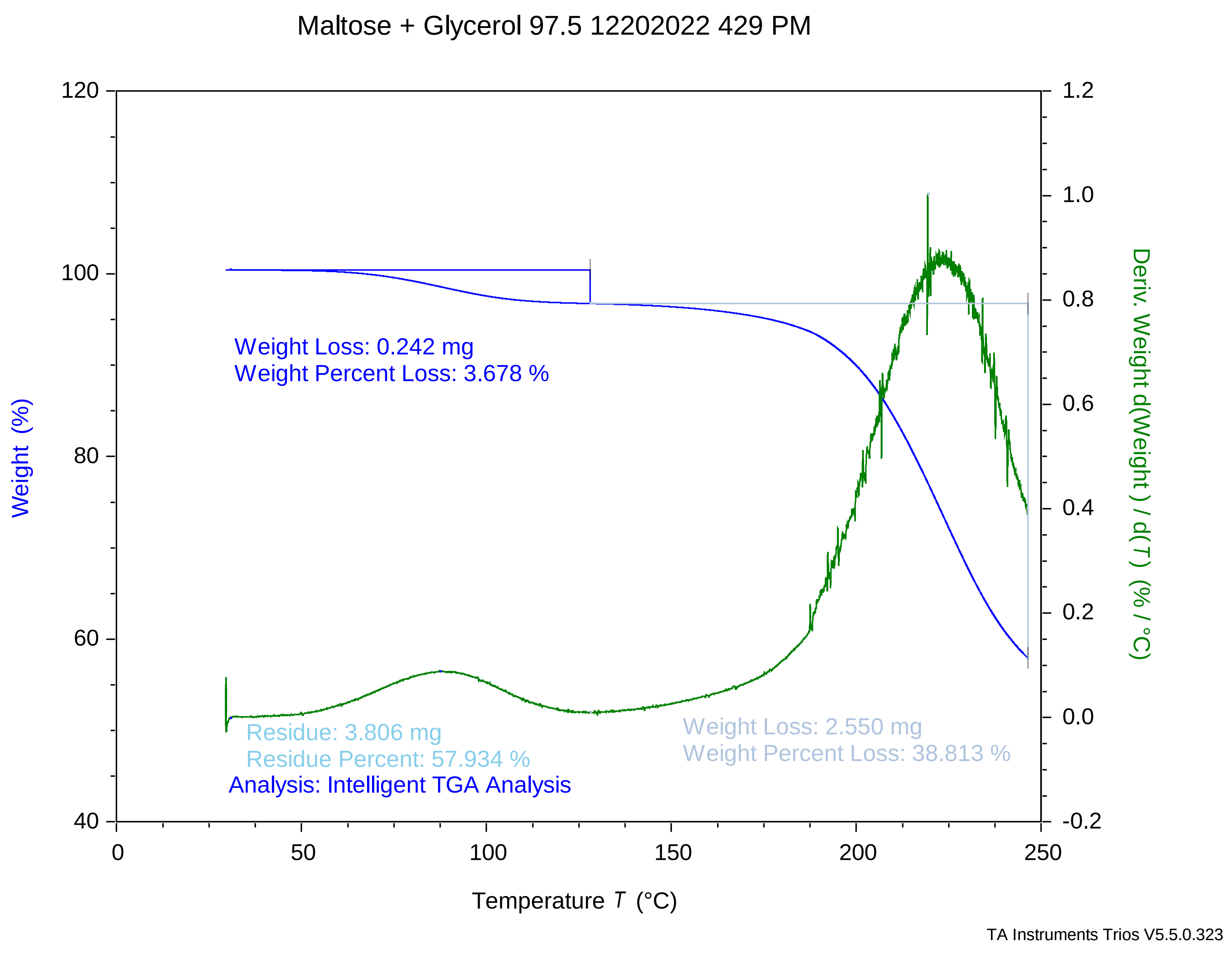

### Maltose + Glycerol 975 12212022 120 PM.pdf

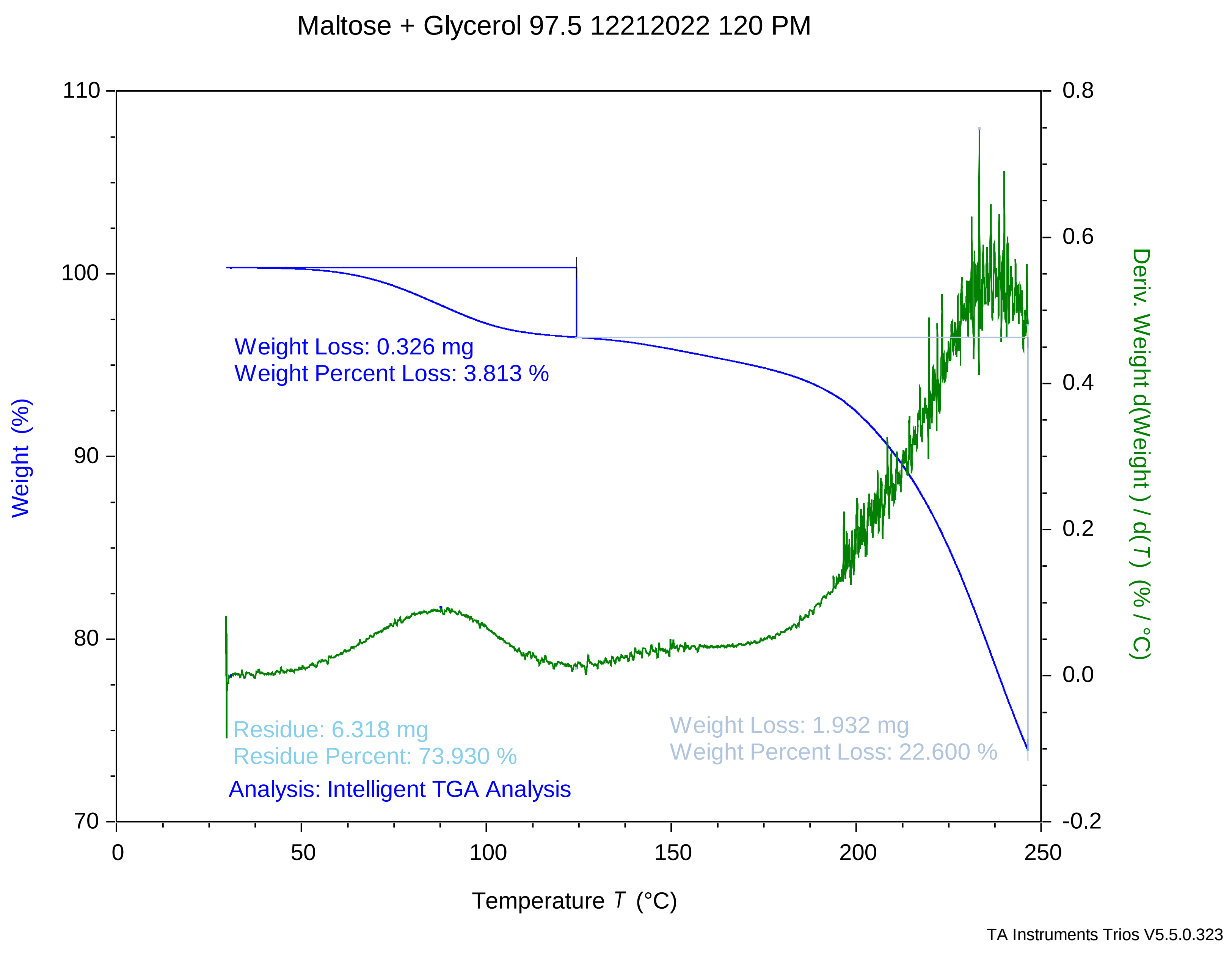

### maltose + glycerol 975 12212022 219 pm.pdf

Maltose + Glycerol 97.5 12212022 219 PM

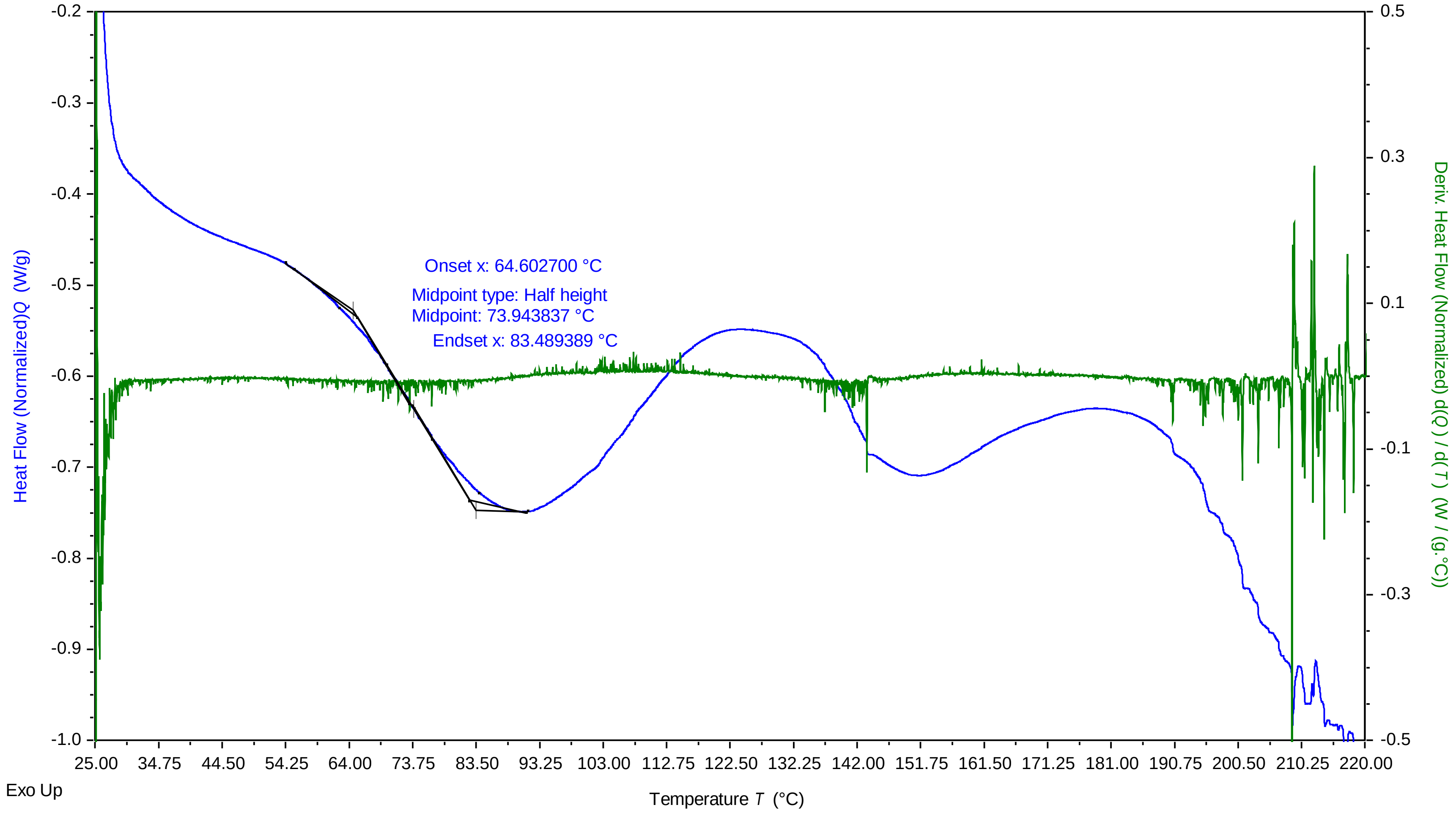


TA Instruments Trios V5.0.0.44616

### maltose + glycerol 975 12282022 1128 am.pdf

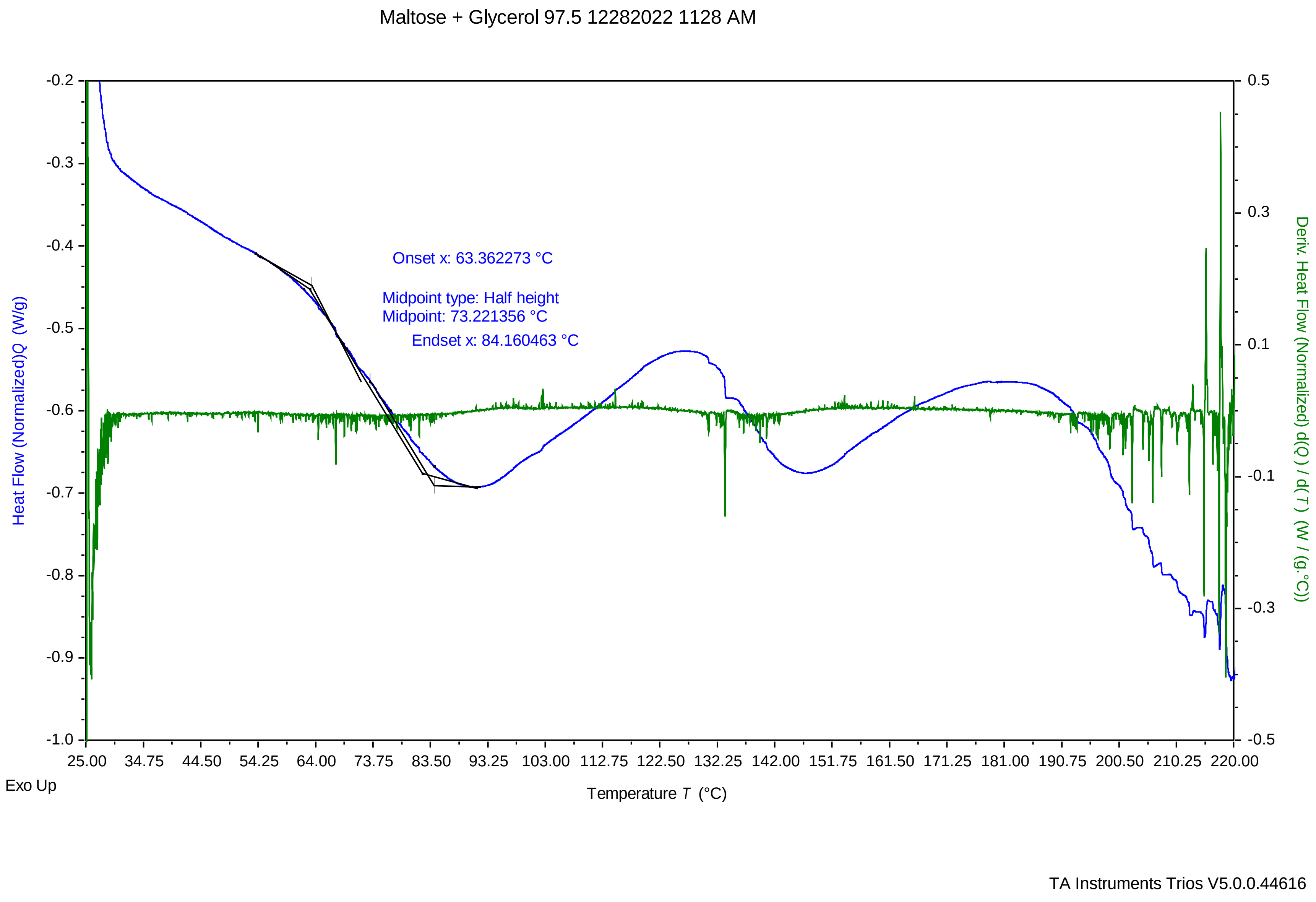

### sucrose 10mg per ml 87.5 percent 1 3222023 746 pm.pdf

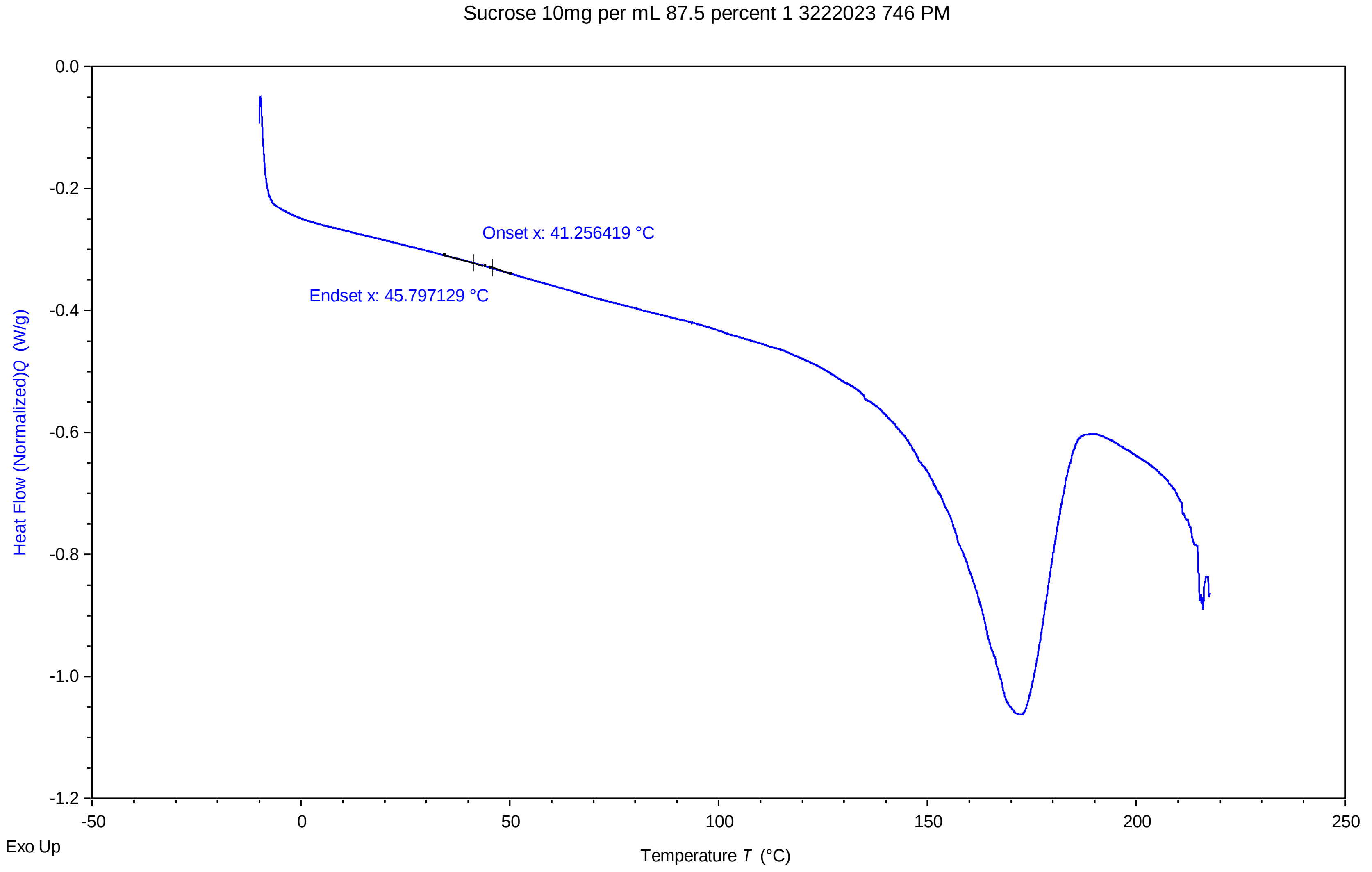


TA Instruments Trios V5.0.0.44616

### sucrose 10mg per ml 87.5 percent 2 3222023 819 pm.pdf

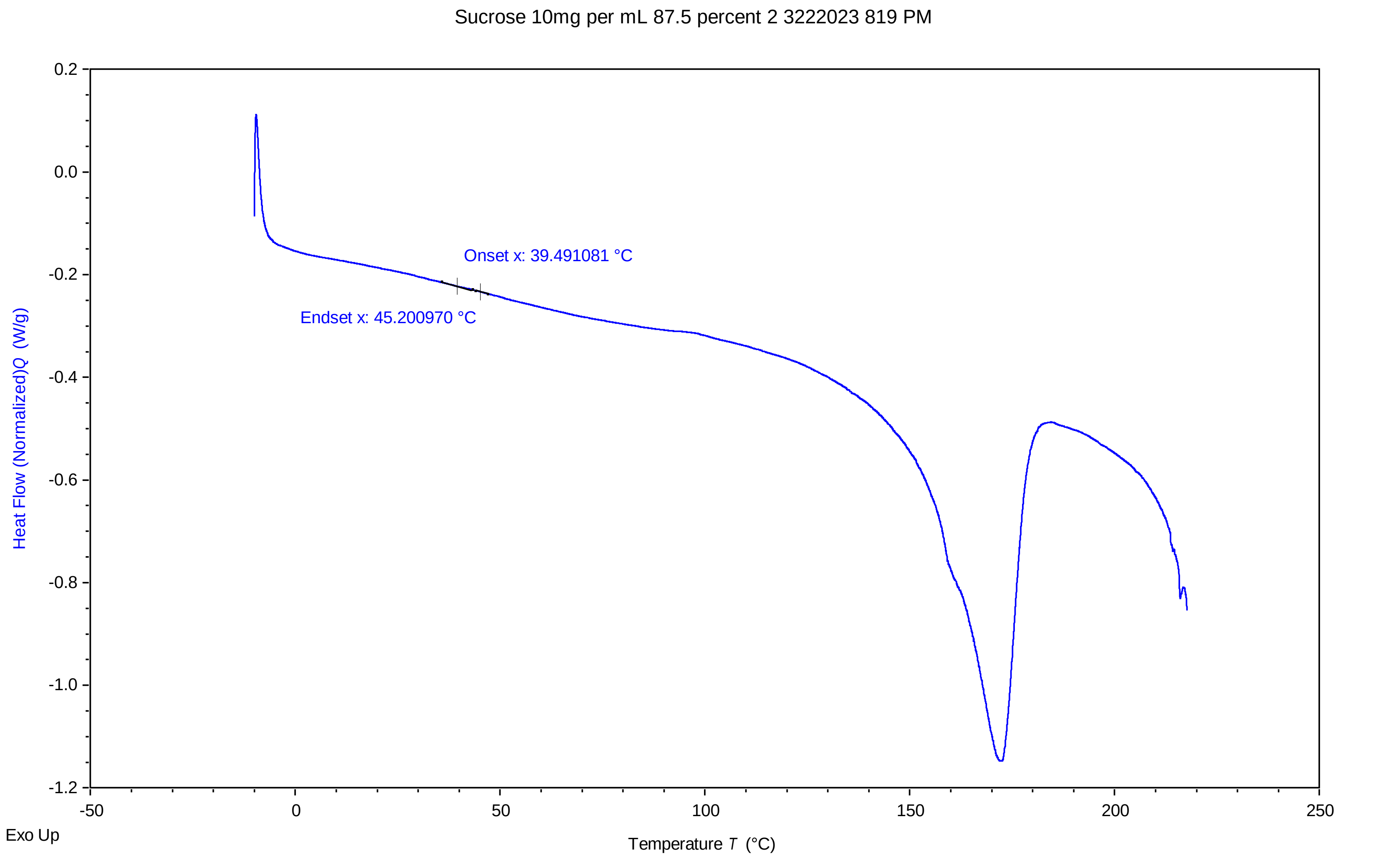


TA Instruments Trios V5.0.0.44616

### sucrose 10mg per ml 87.5 percent 3 3222023 852 pm.pdf

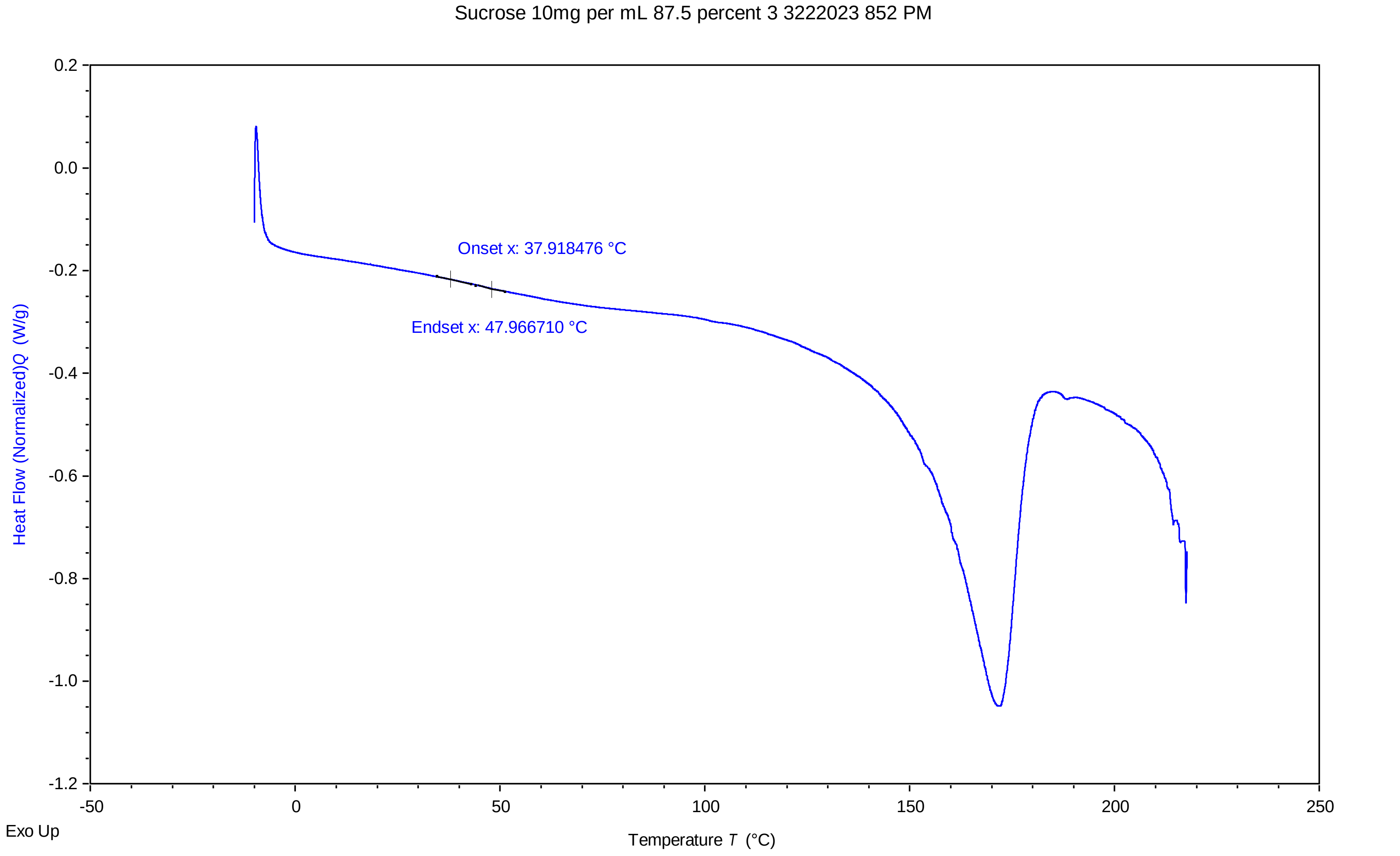


TA Instruments Trios V5.0.0.44616

### sucrose 10mg per ml 90 percent 1 3222023 606 pm.pdf

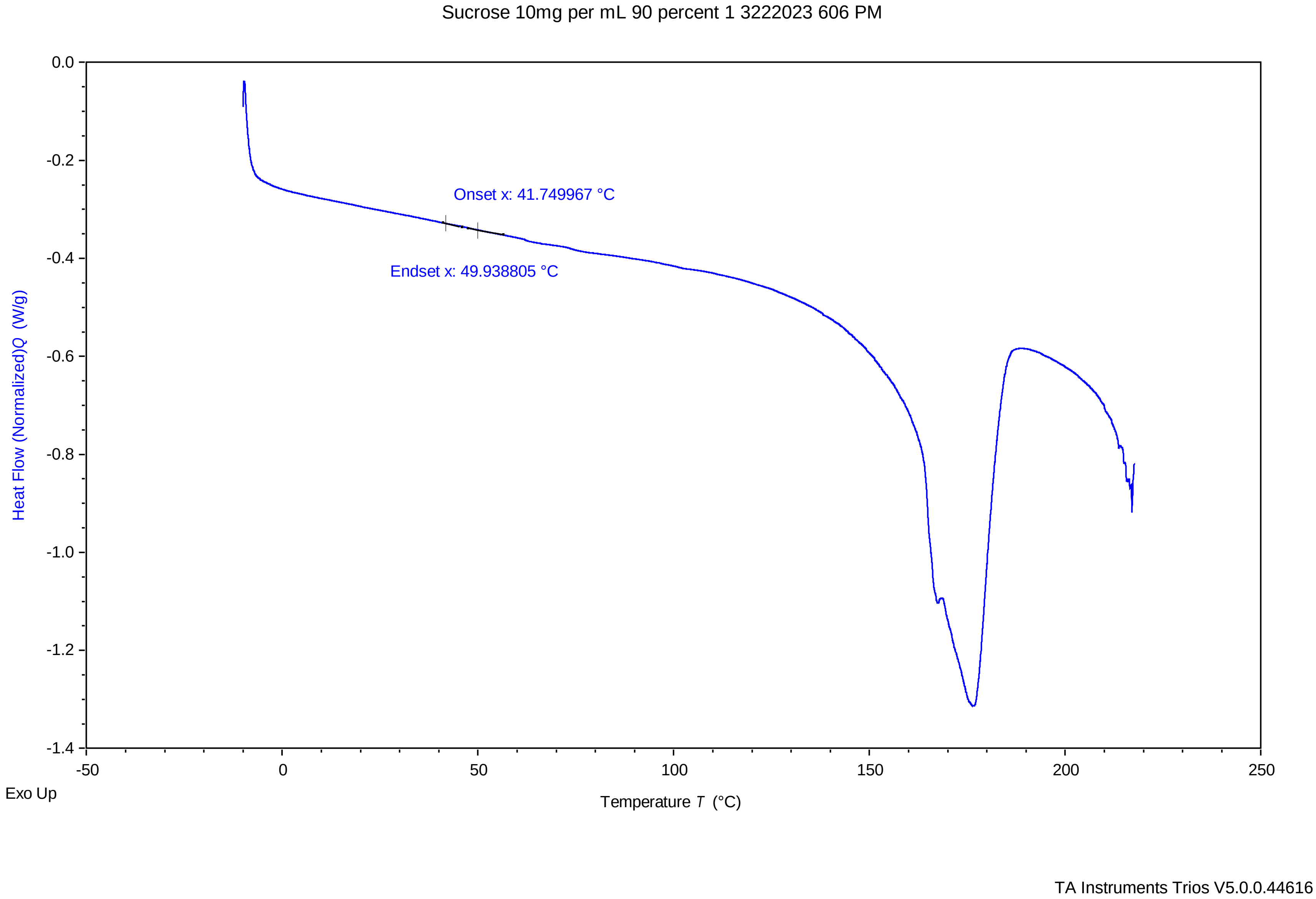

### sucrose 10mg per ml 90 percent 2 3222023 639 pm.pdf

#### Sucrose 10mg per mL 90 percent 2 3222023 639 PM

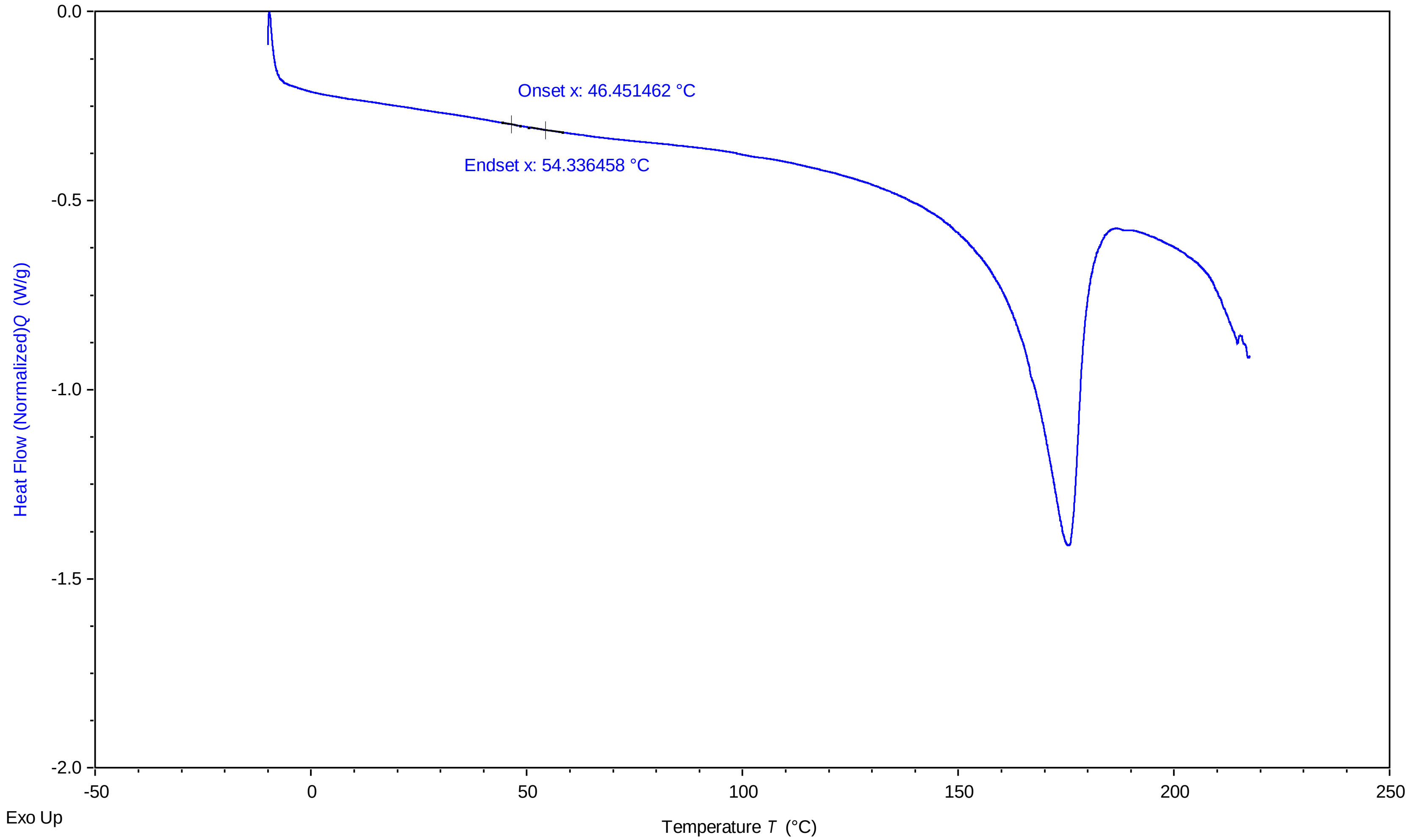


TA Instruments Trios V5.0.0.44616

### sucrose 10mg per ml 92.5 percent 1 3222023 427 pm.pdf

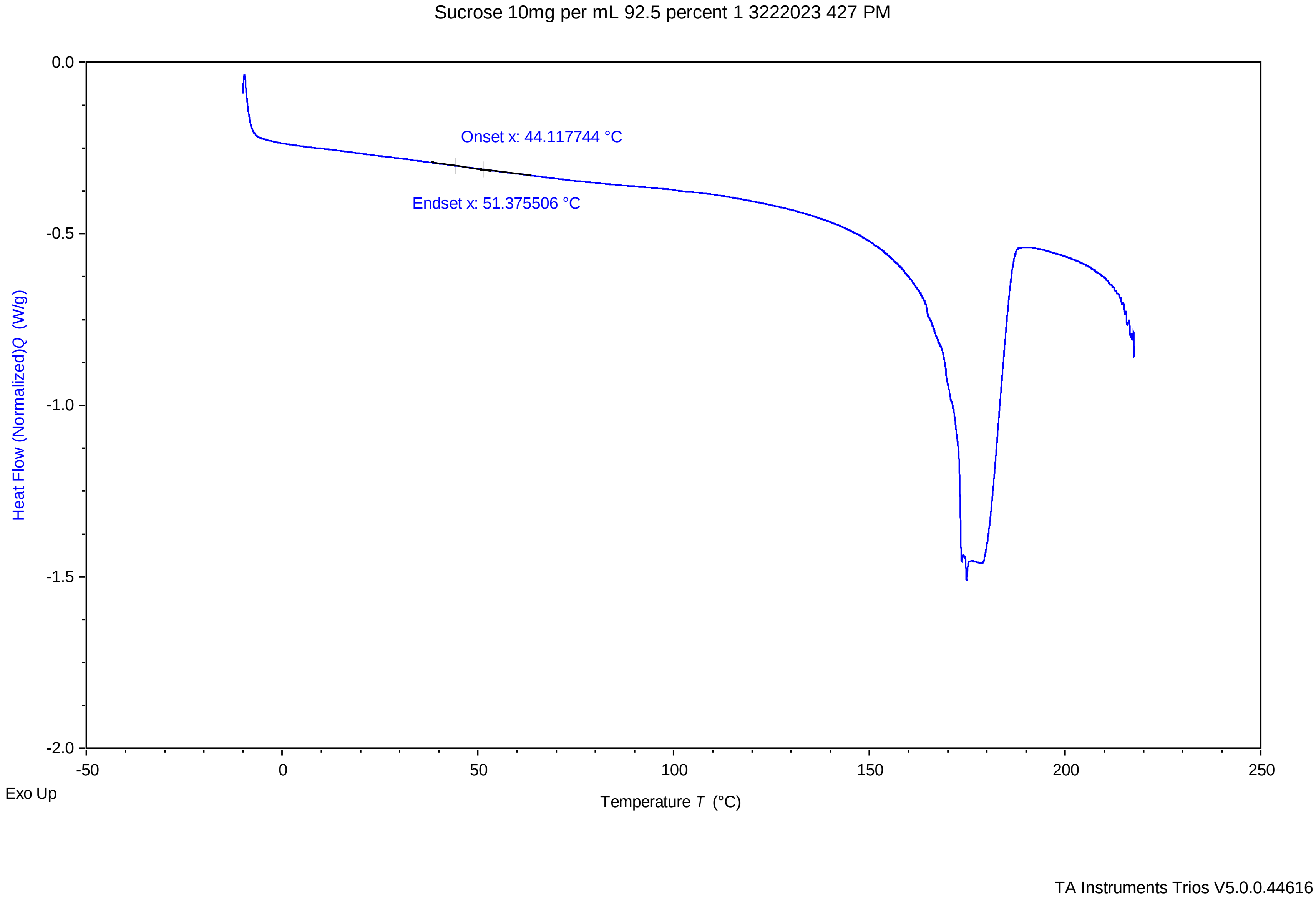

### sucrose 10mg per ml 92.5 percent 2 3222023 500 pm.pdf

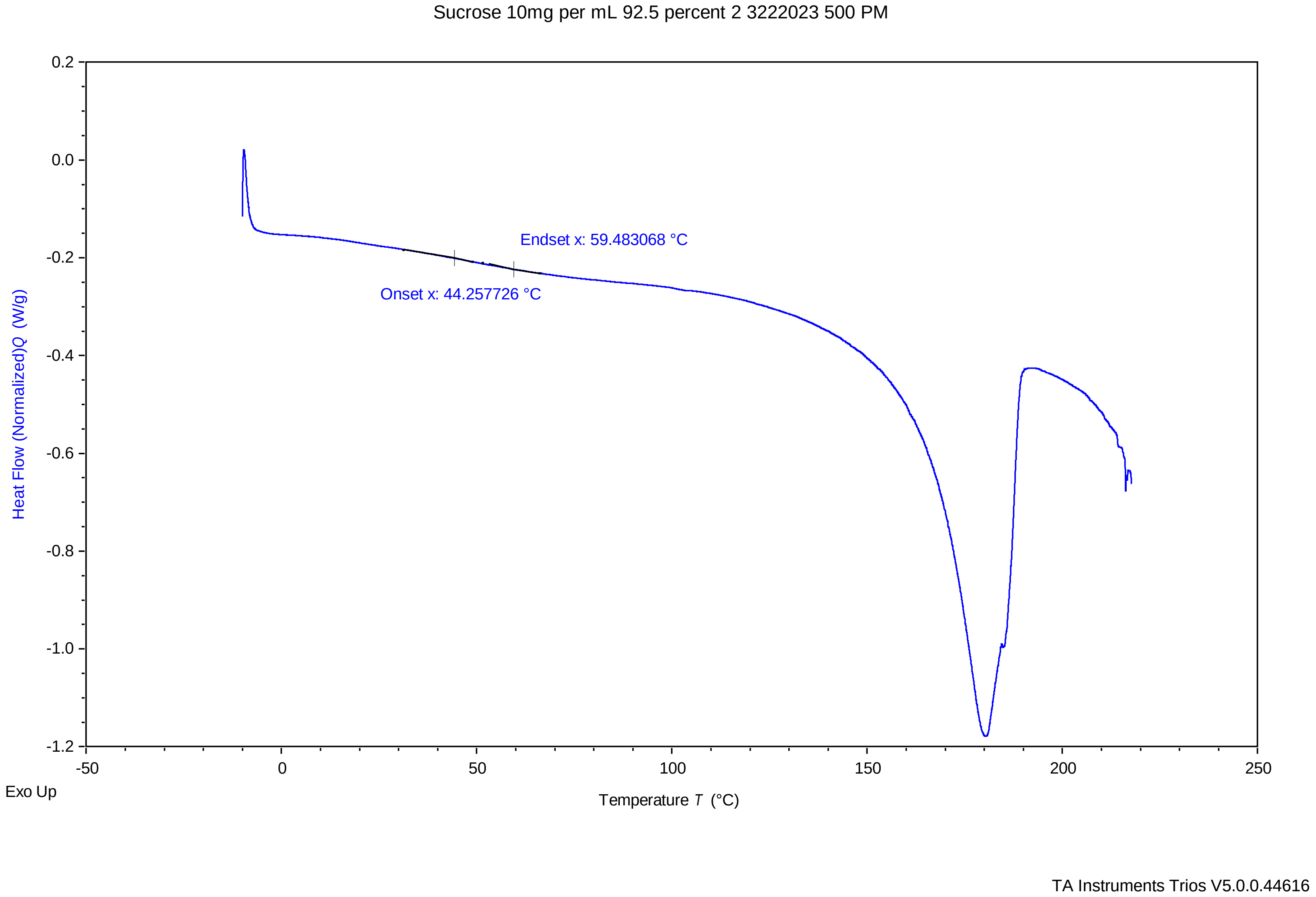

### sucrose 10mg per ml 92.5 percent 3 3222023 533 pm.pdf

Sucrose 10mg per mL 92.5 percent 3 3222023 533 PM

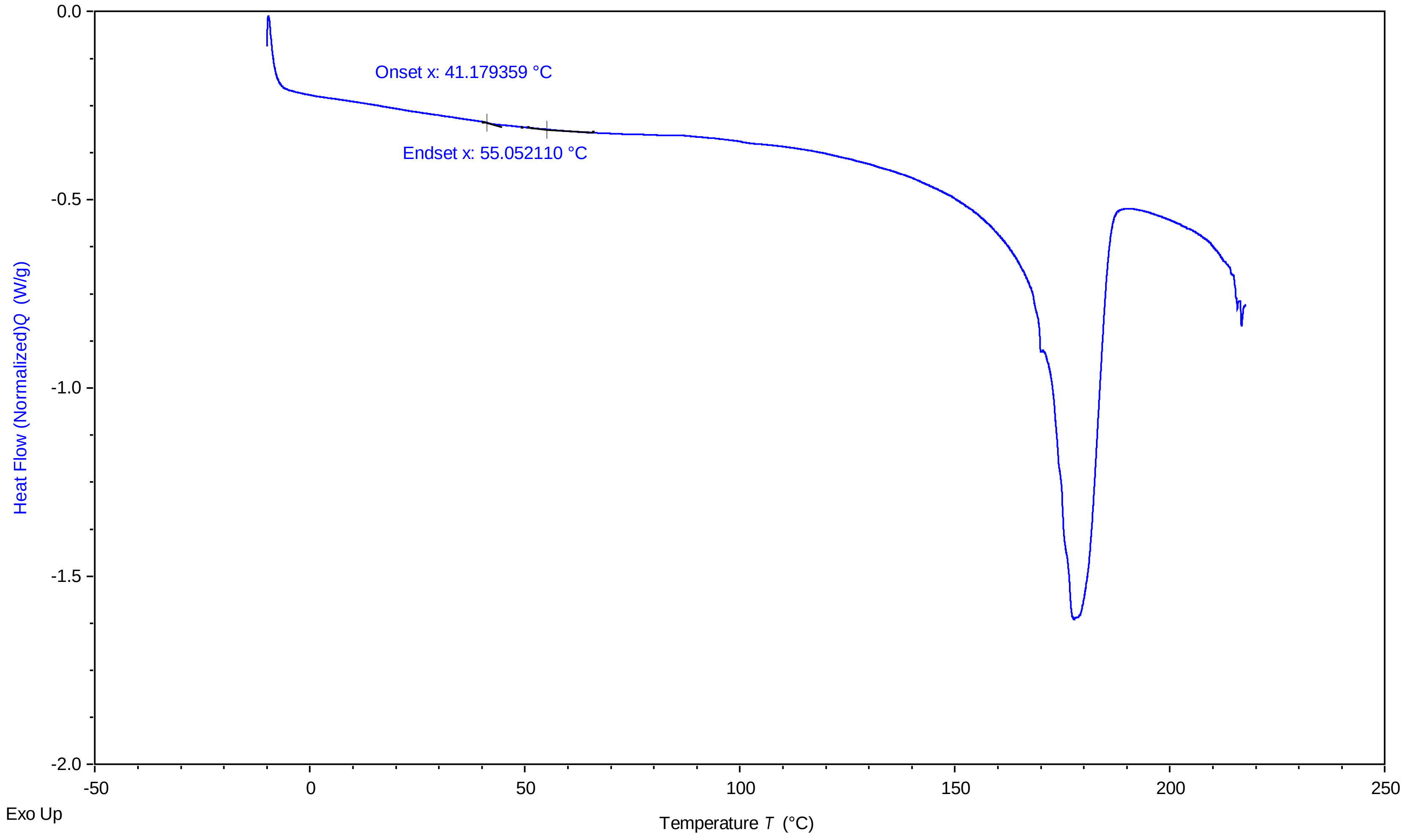


TA Instruments Trios V5.0.0.44616

### sucrose 10mg per ml 95 percent 1 3222023 248 pm.pdf

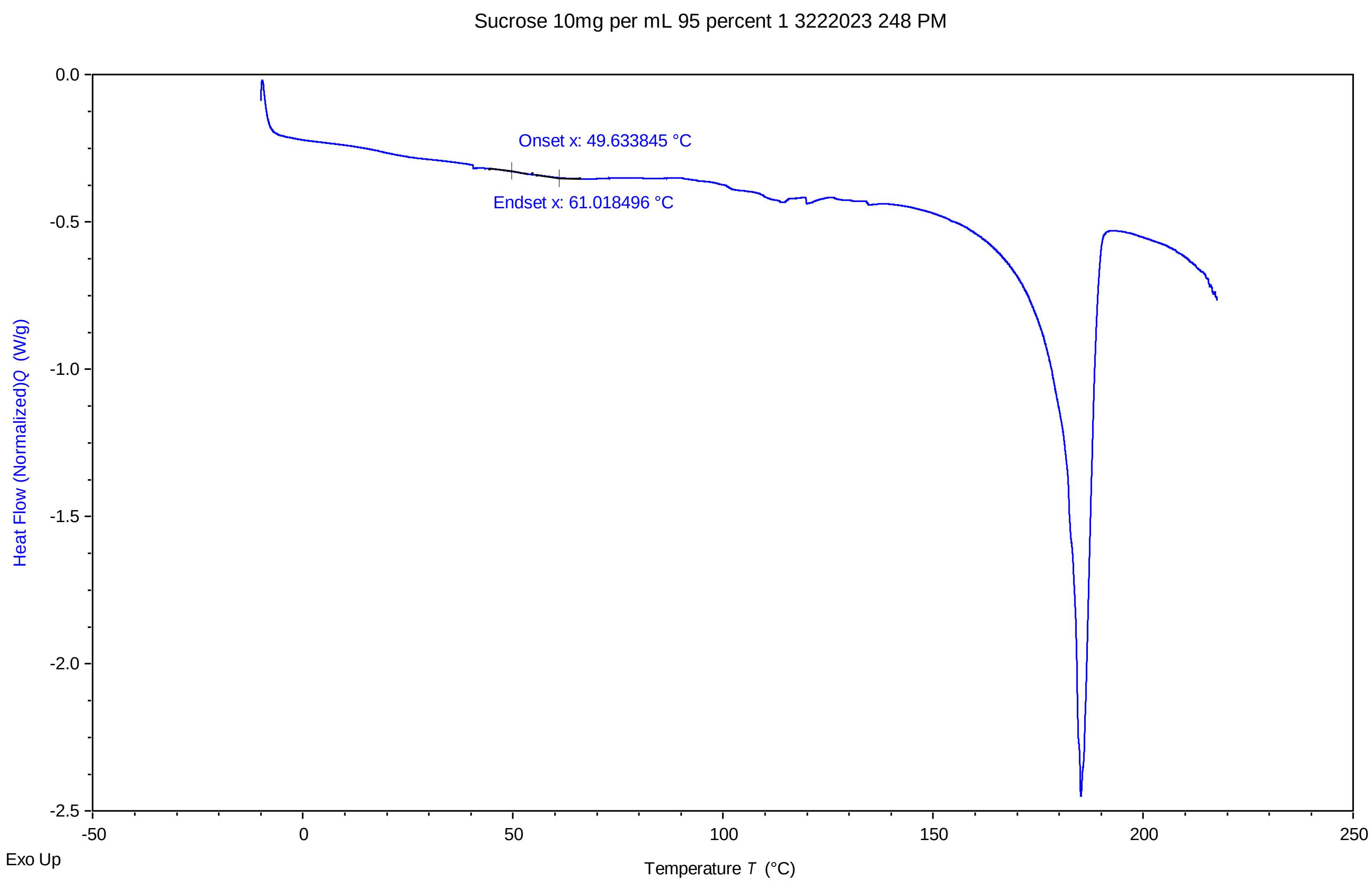


TA Instruments Trios V5.0.0.44616

### sucrose 10mg per ml 95 percent 3 3222023 354 pm.pdf

Sucrose 10mg per mL 95 percent 3 3222023 354 PM

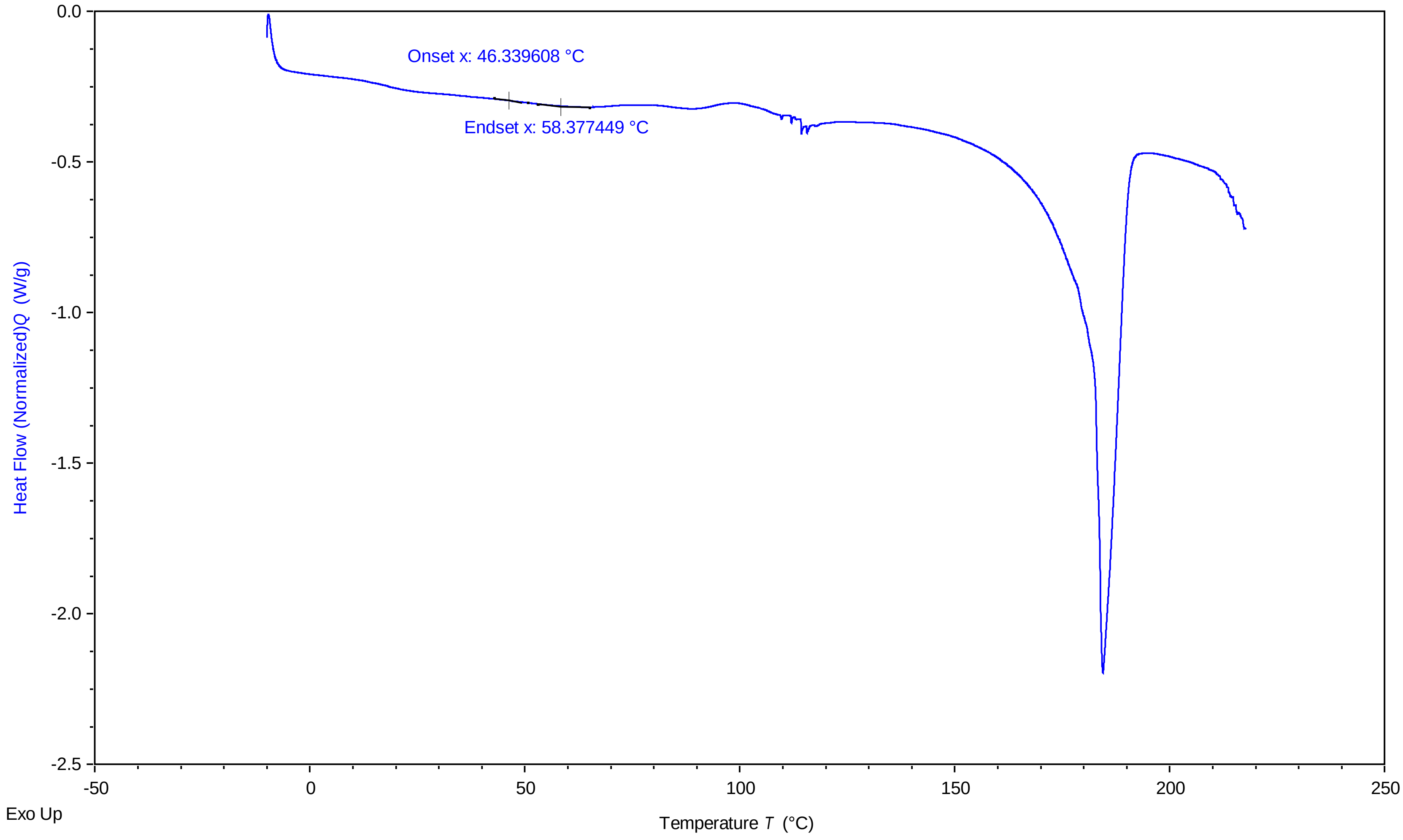


TA Instruments Trios V5.0.0.44616

### sucrose 10mg per ml 97.5 percent 1 3222023 108 pm.pdf

Sucrose 10mg per mL 97.5 percent 1 3222023 108 PM

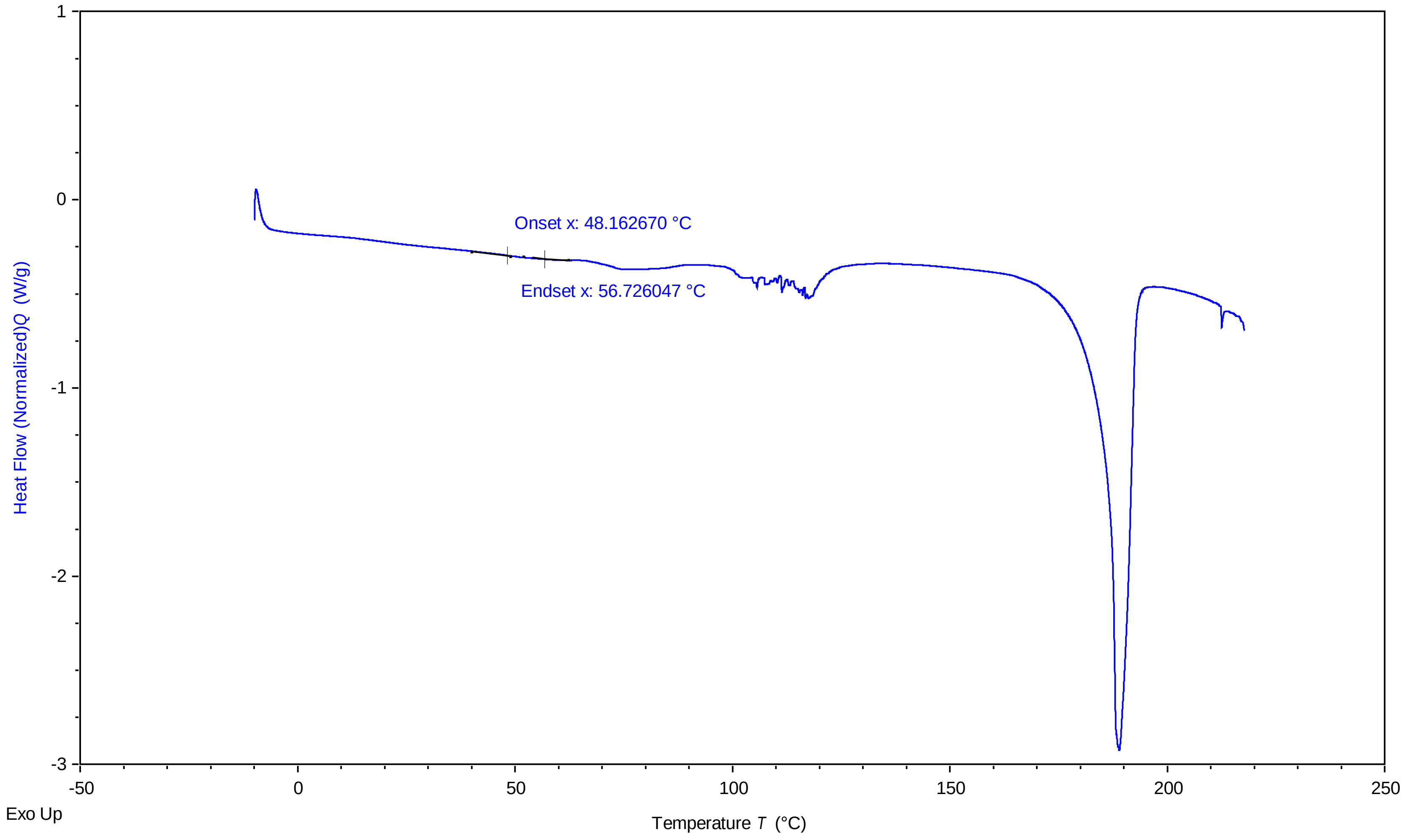


TA Instruments Trios V5.0.0.44616

### sucrose 10mg per ml 97.5 percent 3 3222023 214 pm.pdf

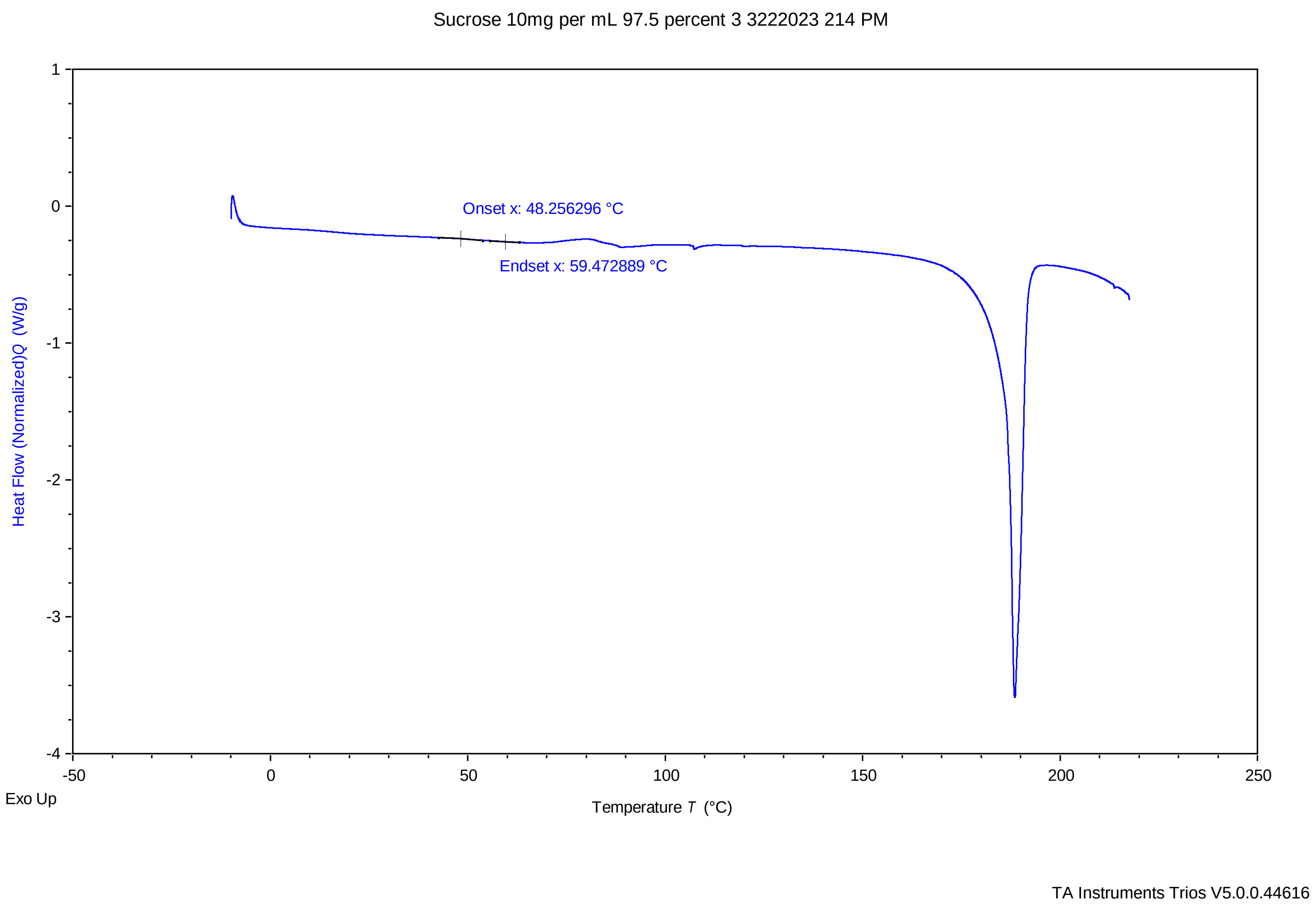

### sucrose 10mg per ml 100 percent 1 3222023 1129 am.pdf

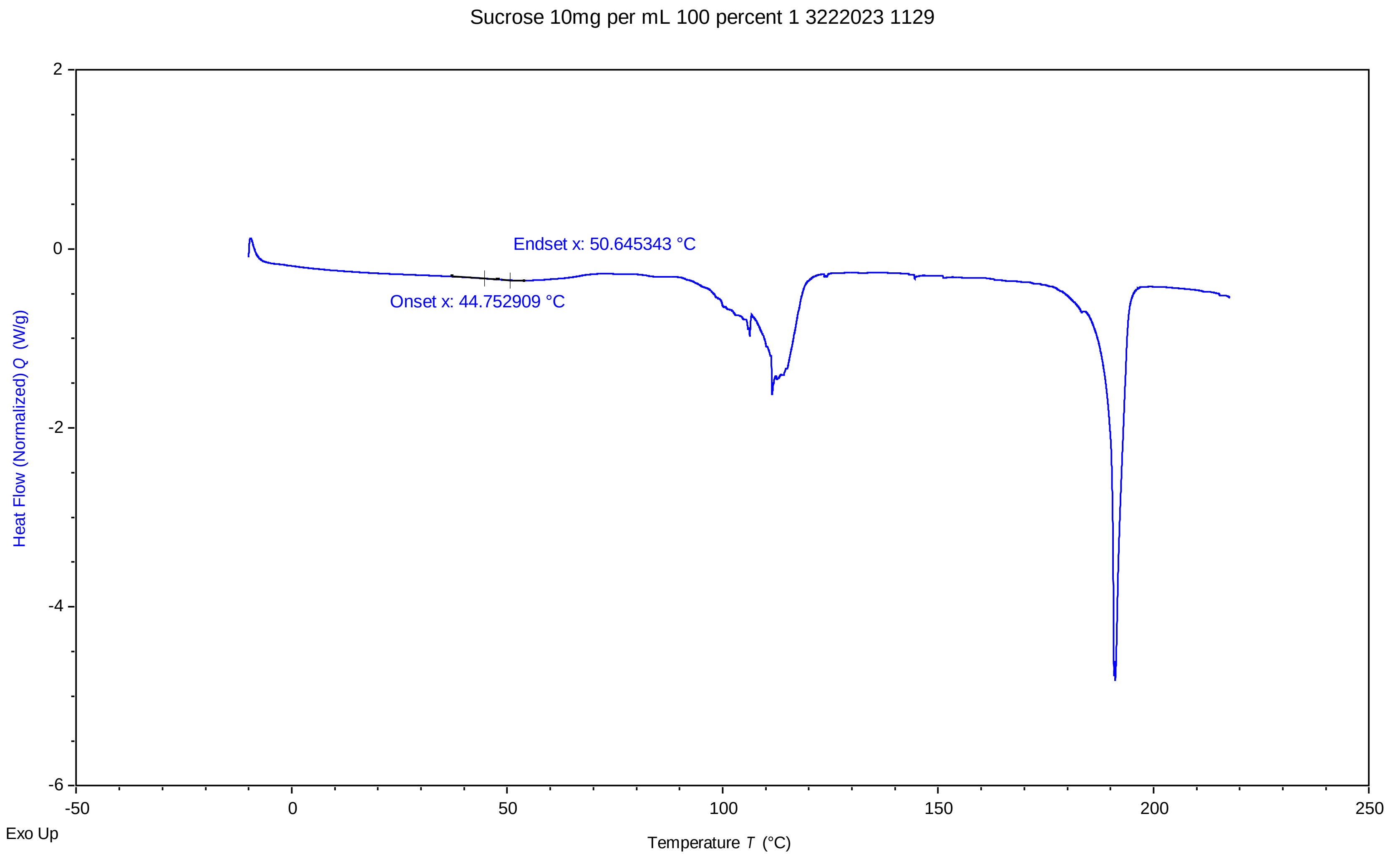


TA Instruments Trios V5.0.0.44616

### sucrose 10mg per ml 100 percent 2 3222023 1202 pm.pdf

Sucrose 10mg per mL 100 percent 2 3222023 1202 PM

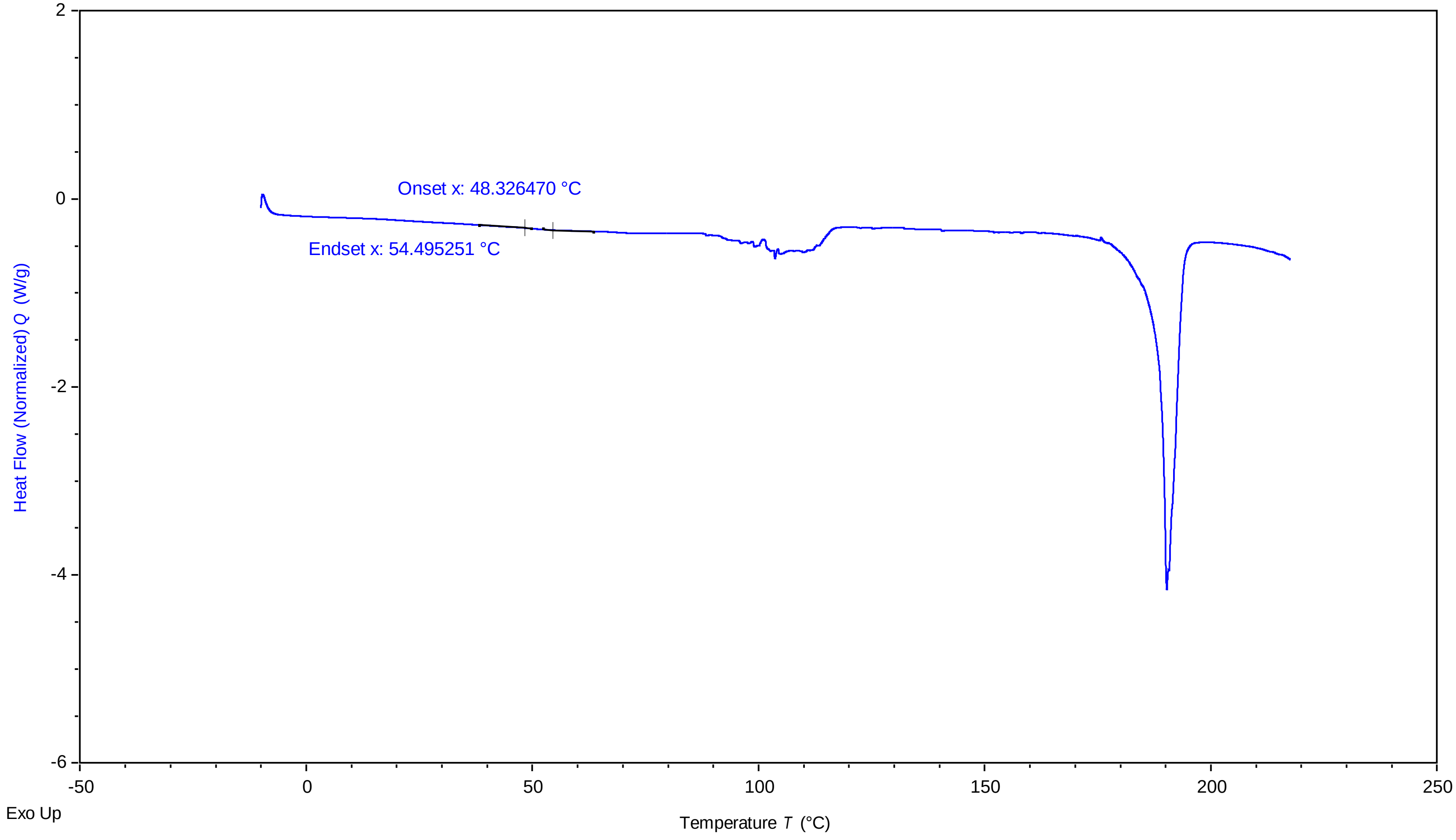


TA Instruments Trios V5.0.0.44616

### sucrose 10mg per ml 100 percent 3 3222023 1235 pm.pdf

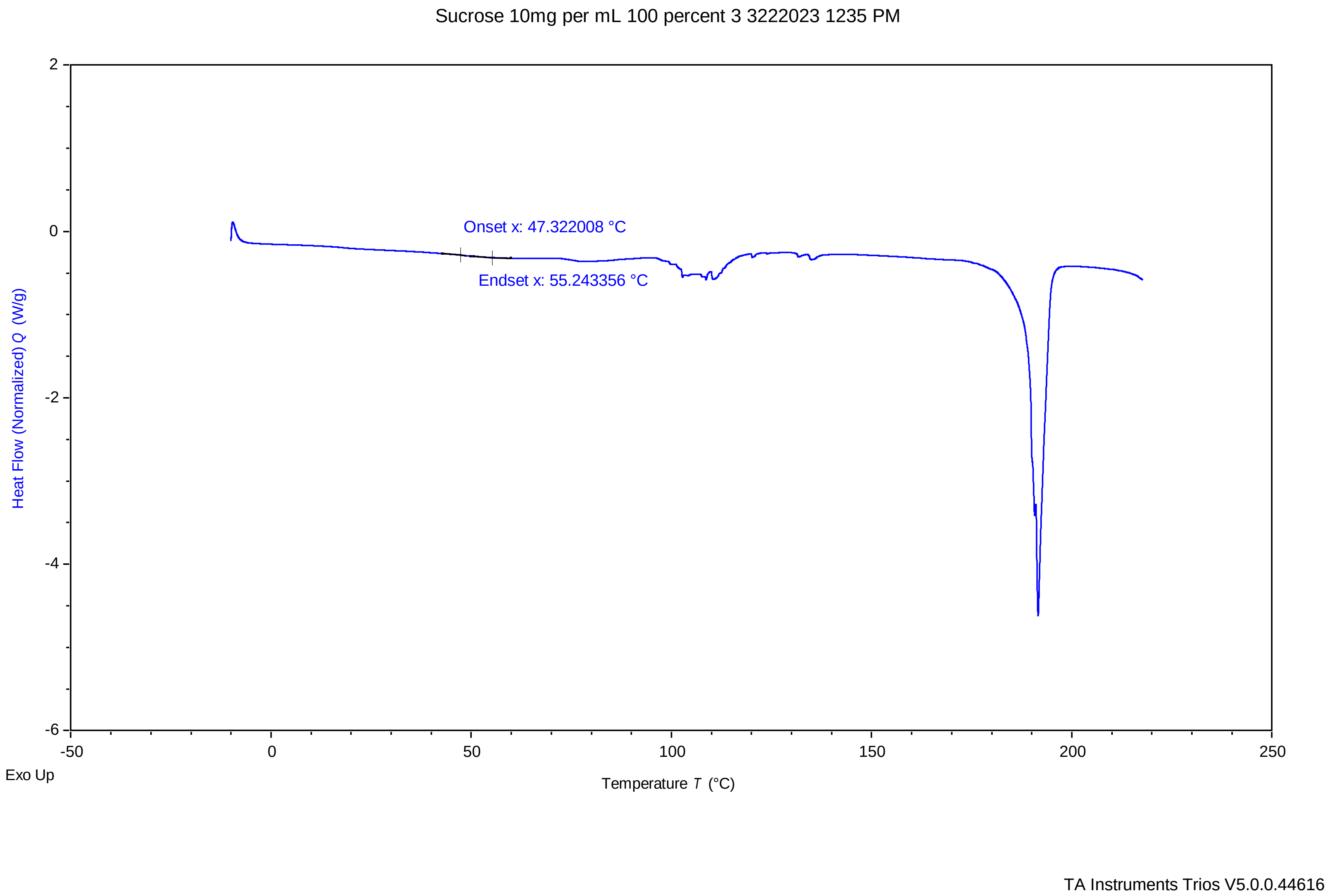

### sucrose 87.5 percent 1 3222023 820 pm.pdf

sucrose 87.5 percent 1 3222023 820 PM

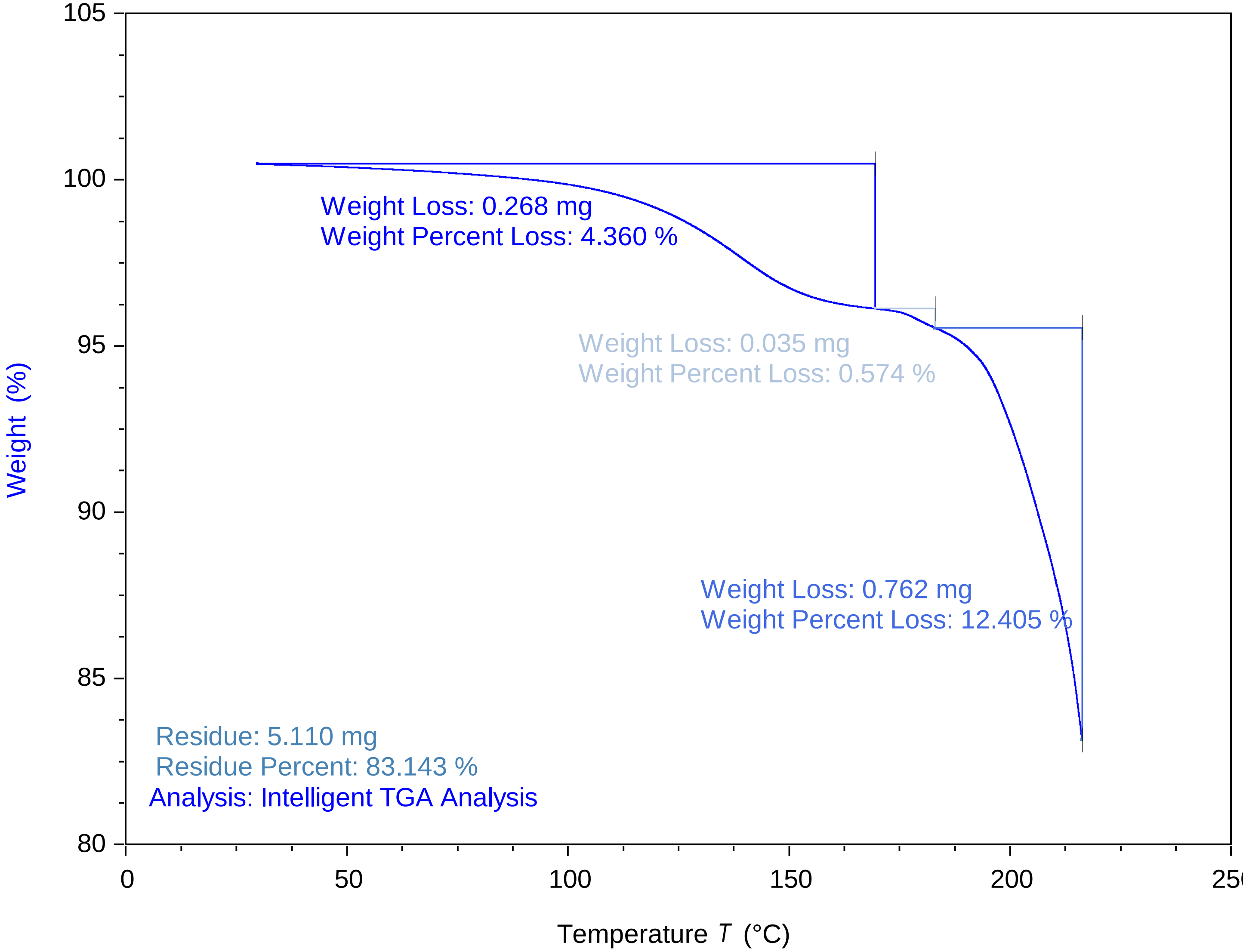


TA Instruments Trios V5.5.0.323

### sucrose 87.5 percent 2 3222023 846 pm.pdf

sucrose 87.5 percent 2 3222023 846 PM

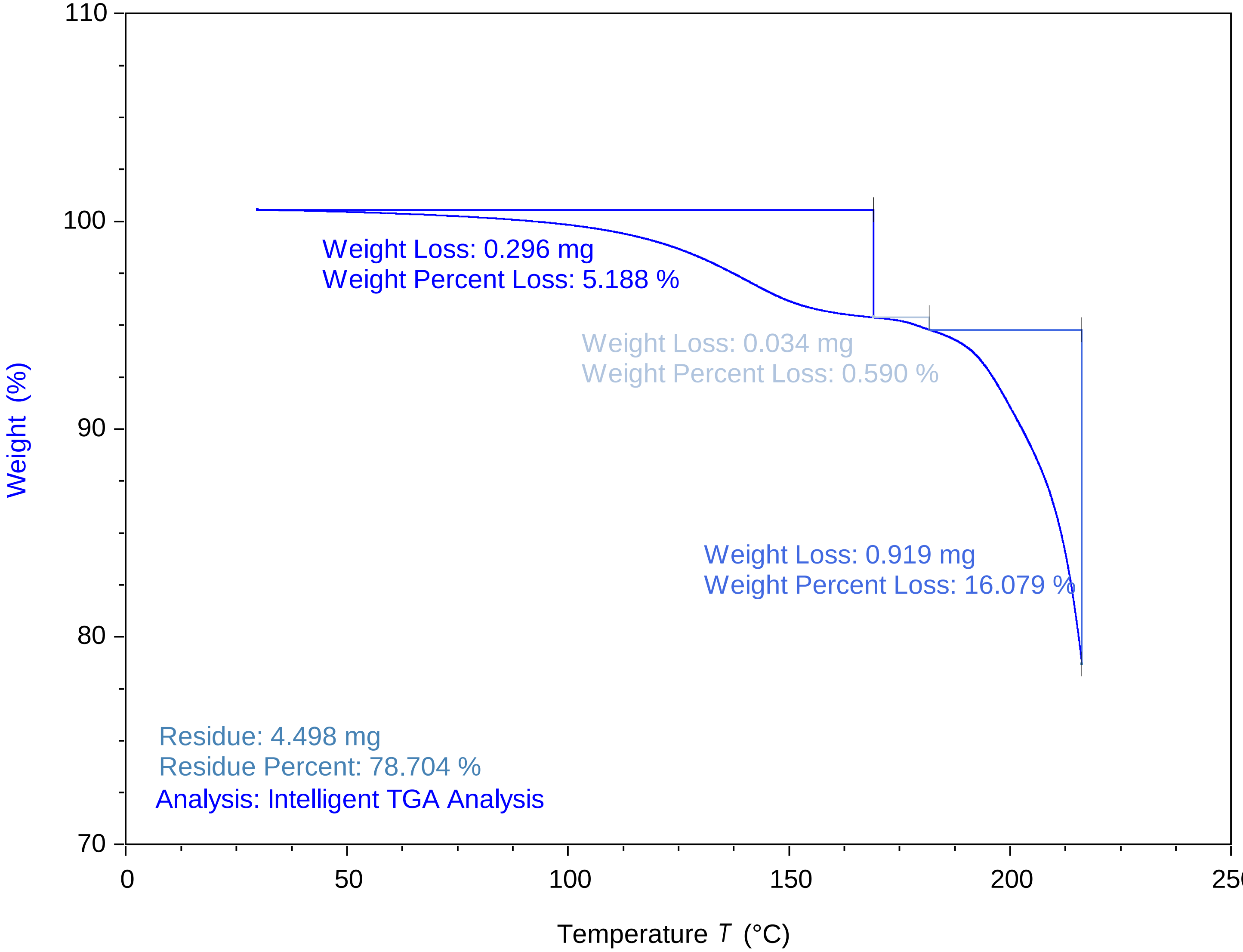


TA Instruments Trios V5.5.0.323

### sucrose 87.5 percent 3 3222023 911 pm.pdf

sucrose 87.5 percent 3 3222023 911 PM

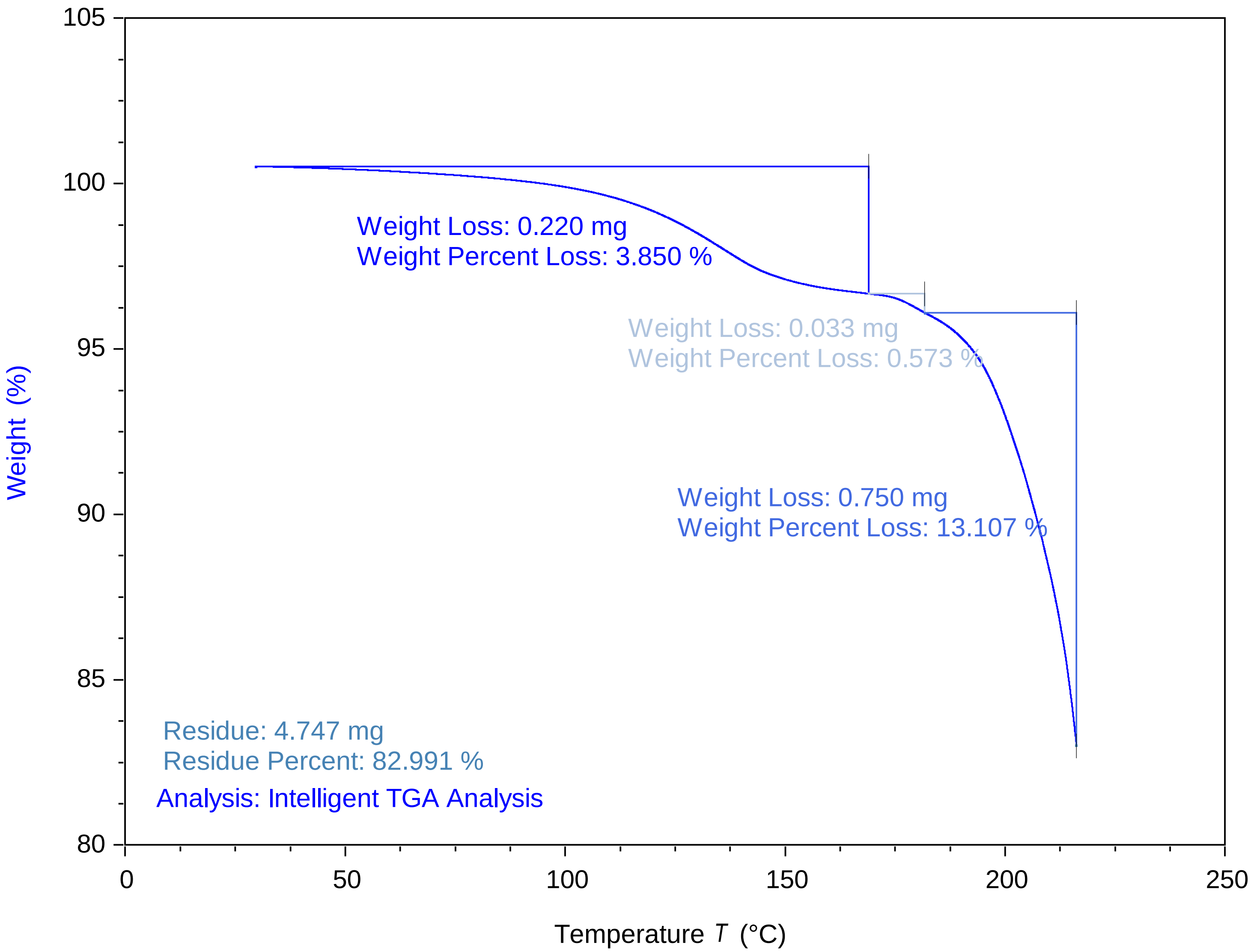


TA Instruments Trios V5.5.0.323

### sucrose 90 percent 1 3222023 702 pm.pdf

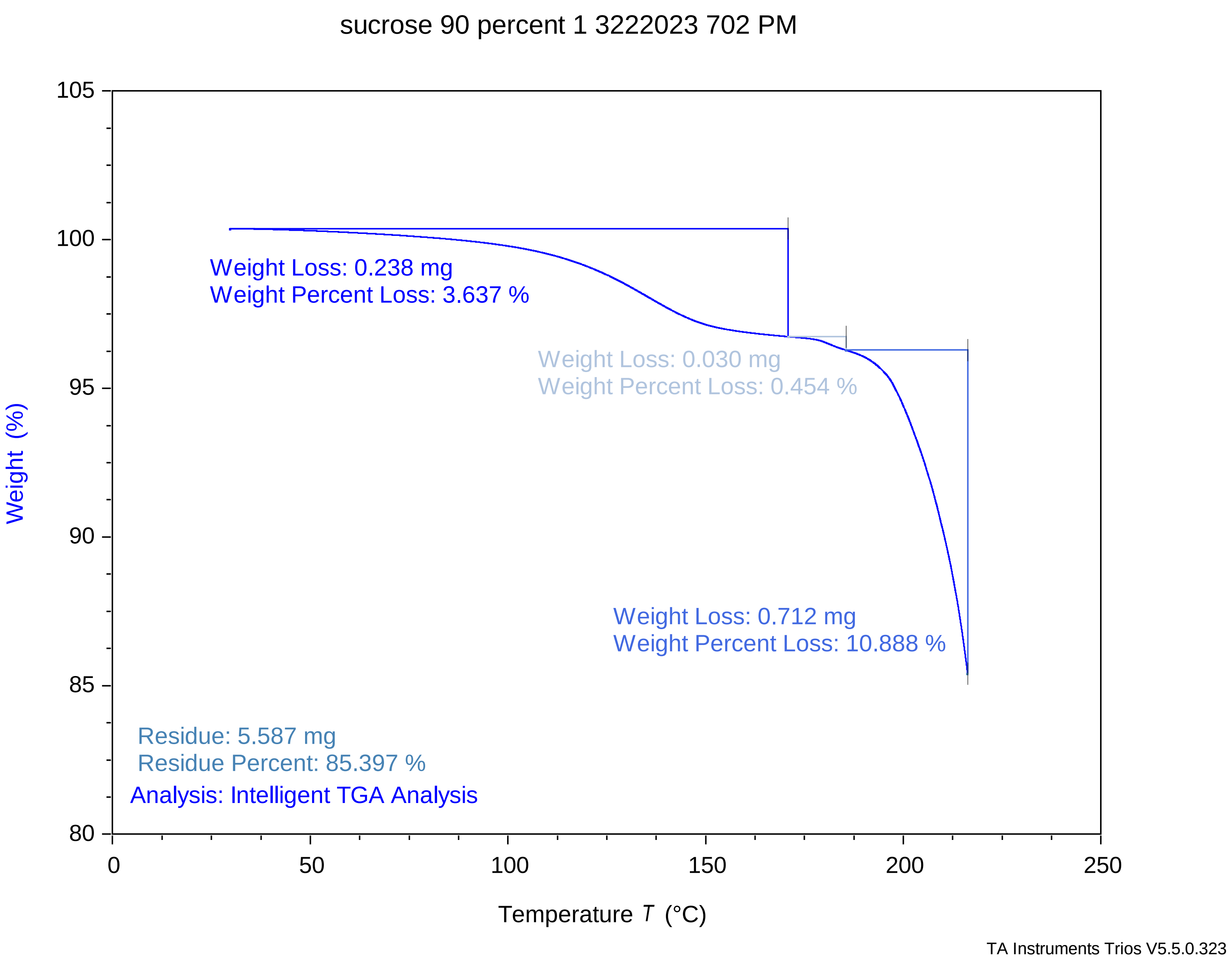

### sucrose 90 percent 2 3222023 728 pm.pdf

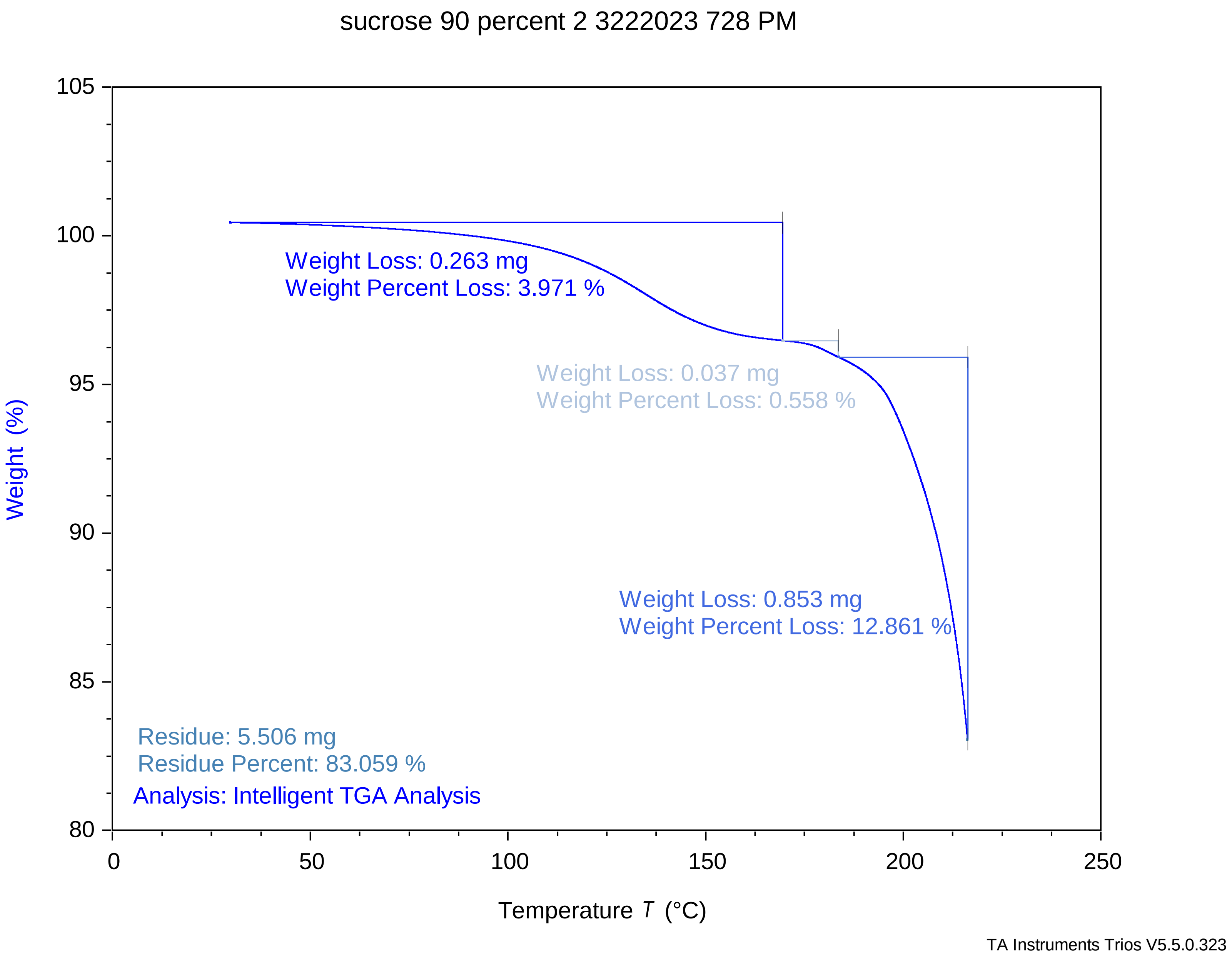

### sucrose 90 percent 3 3222023 754 pm.pdf

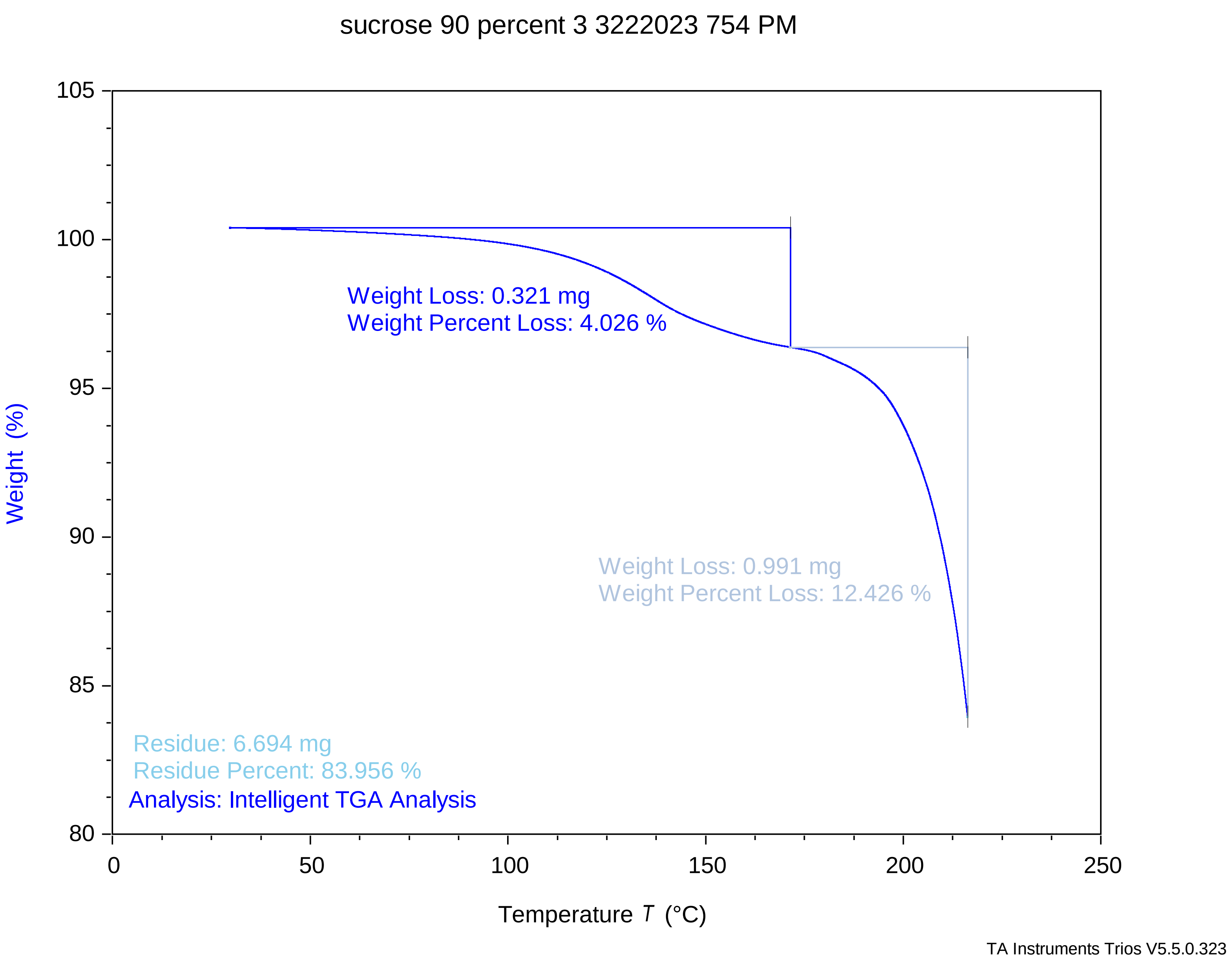

### sucrose 92.5 percent 1 3222023 540 pm.pdf

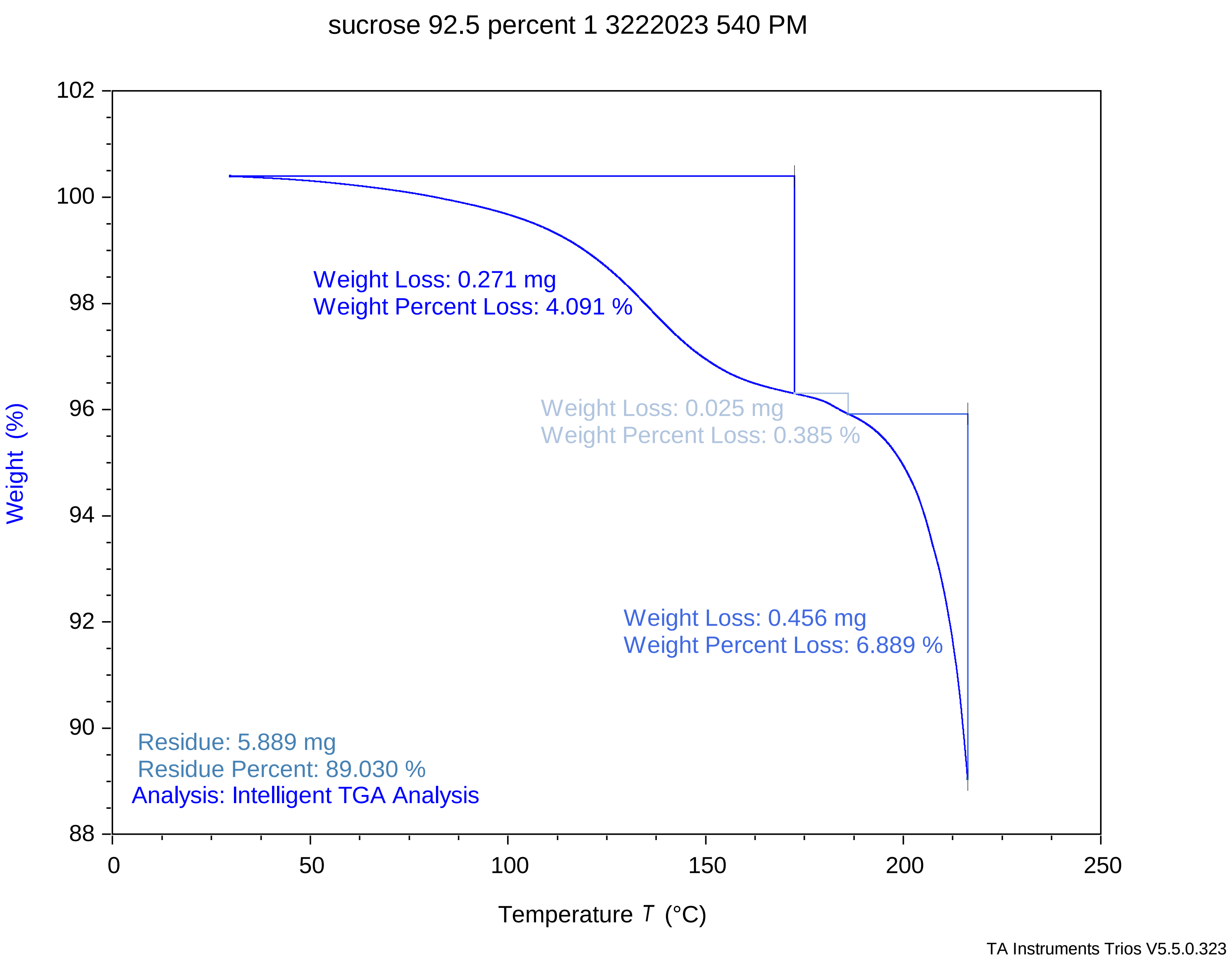

### sucrose 92.5 percent 2 3222023 607 pm.pdf

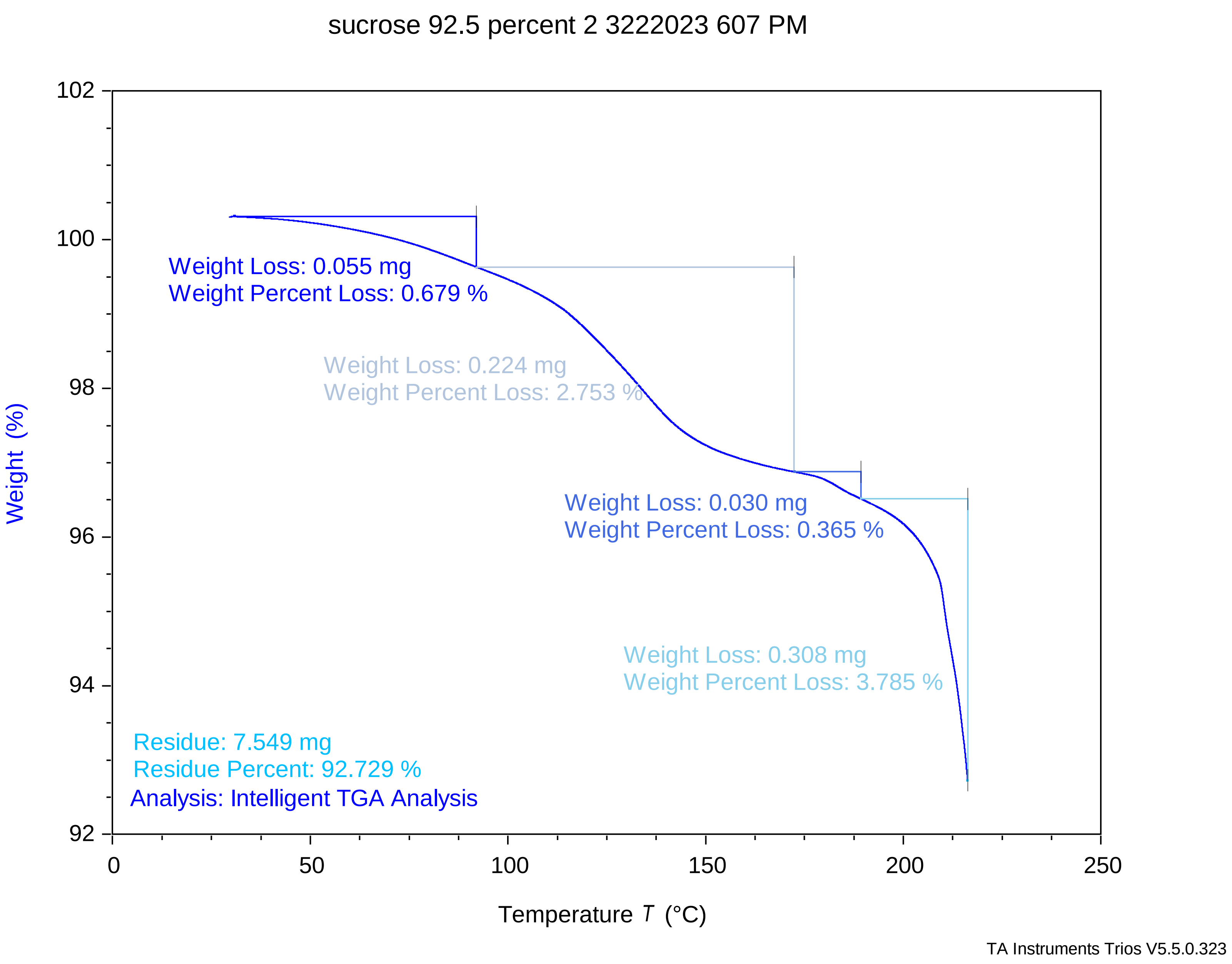

### sucrose 92.5 percent 3 3222023 633 pm.pdf

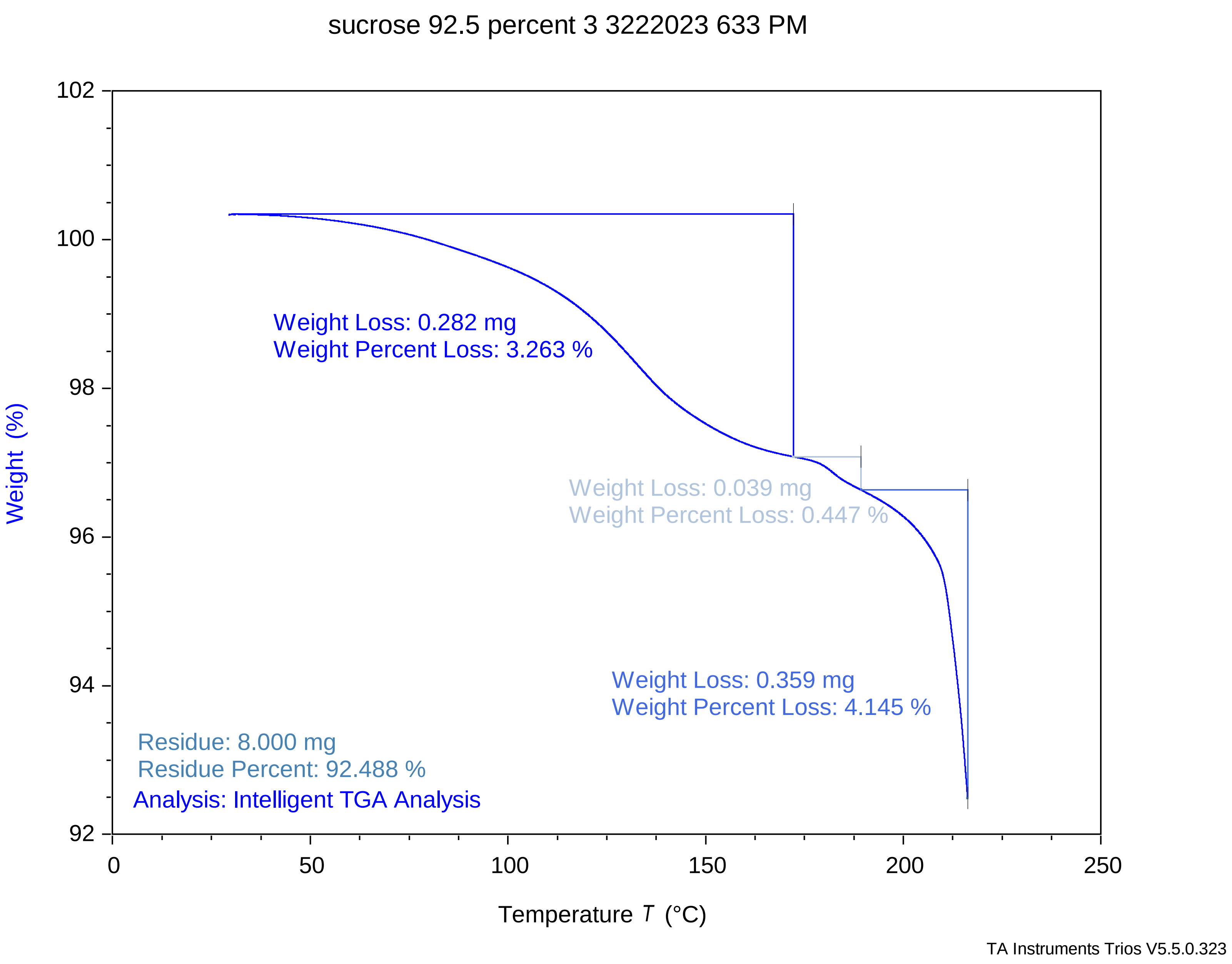

### sucrose 95 percent 1 3222023 414 pm.pdf

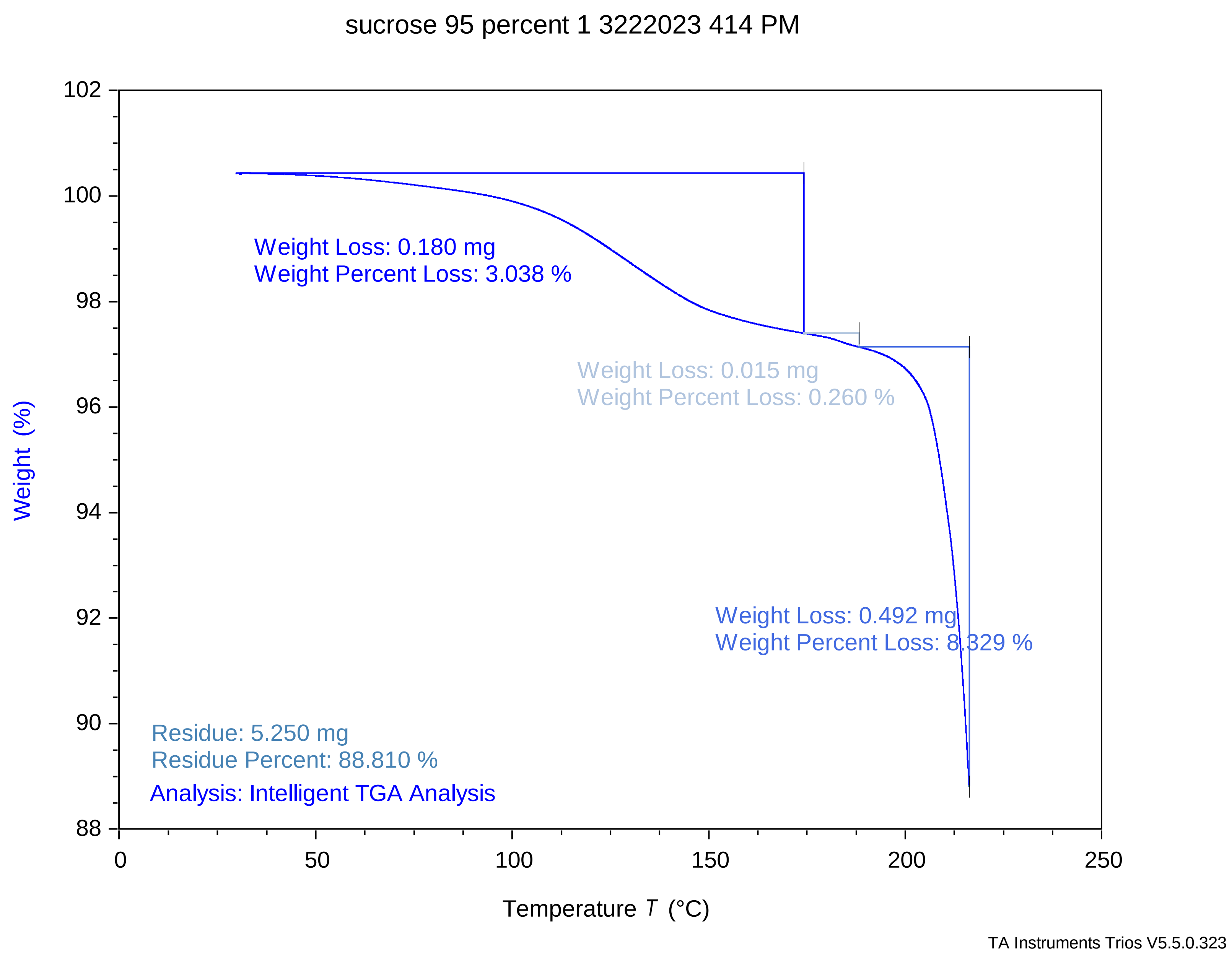

### sucrose 95 percent 2 3222023 442 pm.pdf

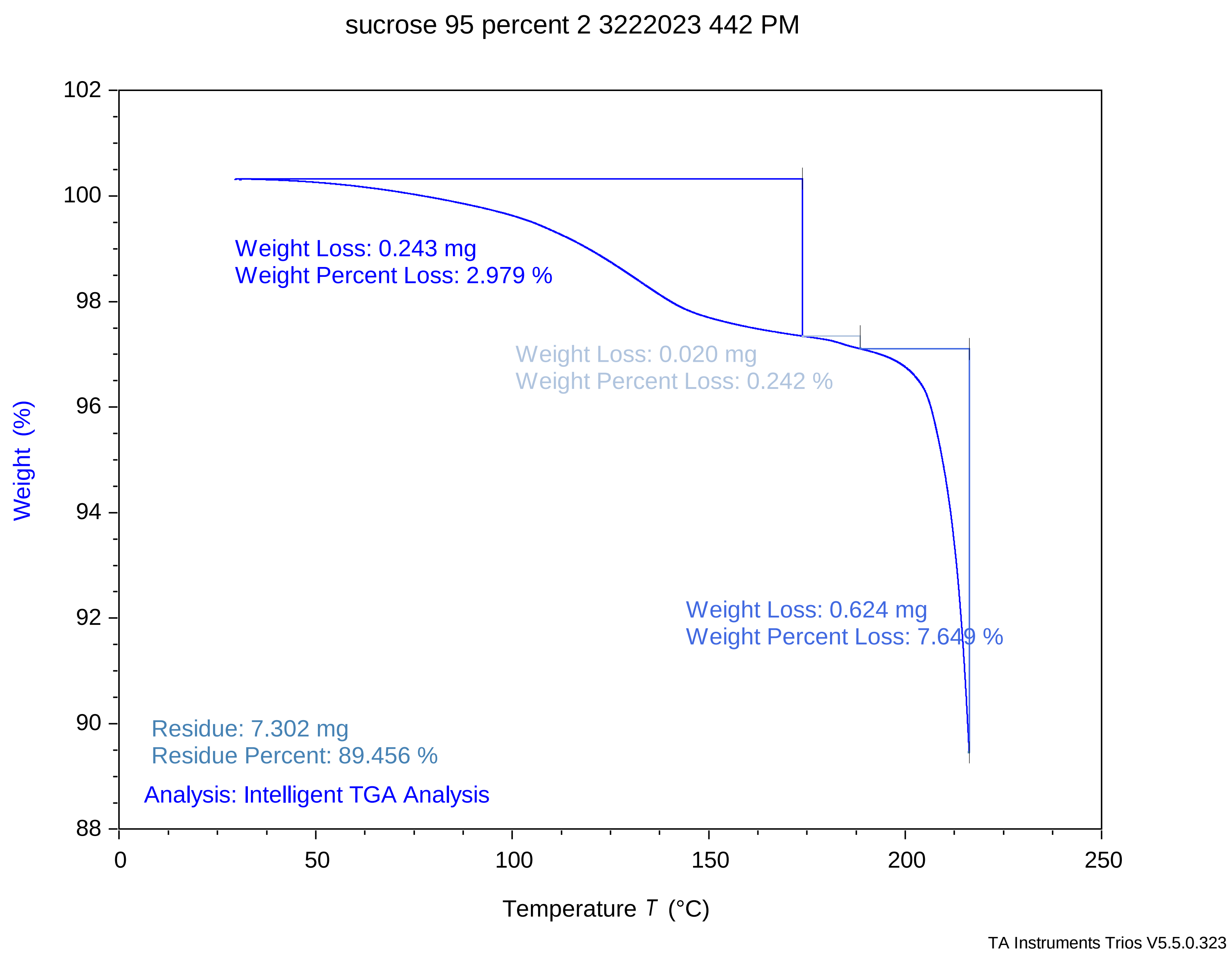

### sucrose 95 percent 3 3222023 511 pm.pdf

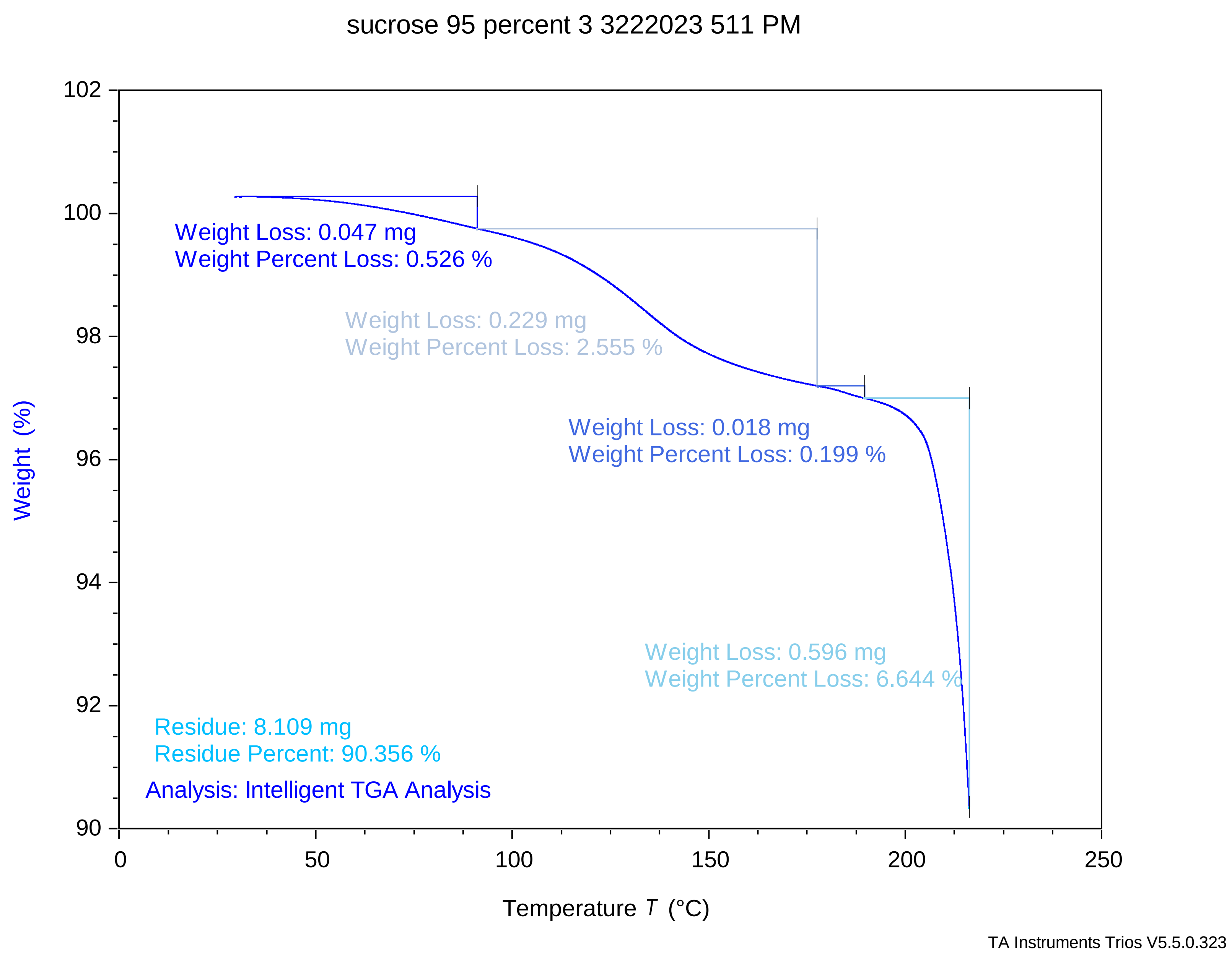

### sucrose 97.5 percent 1 3222023 250 pm.pdf

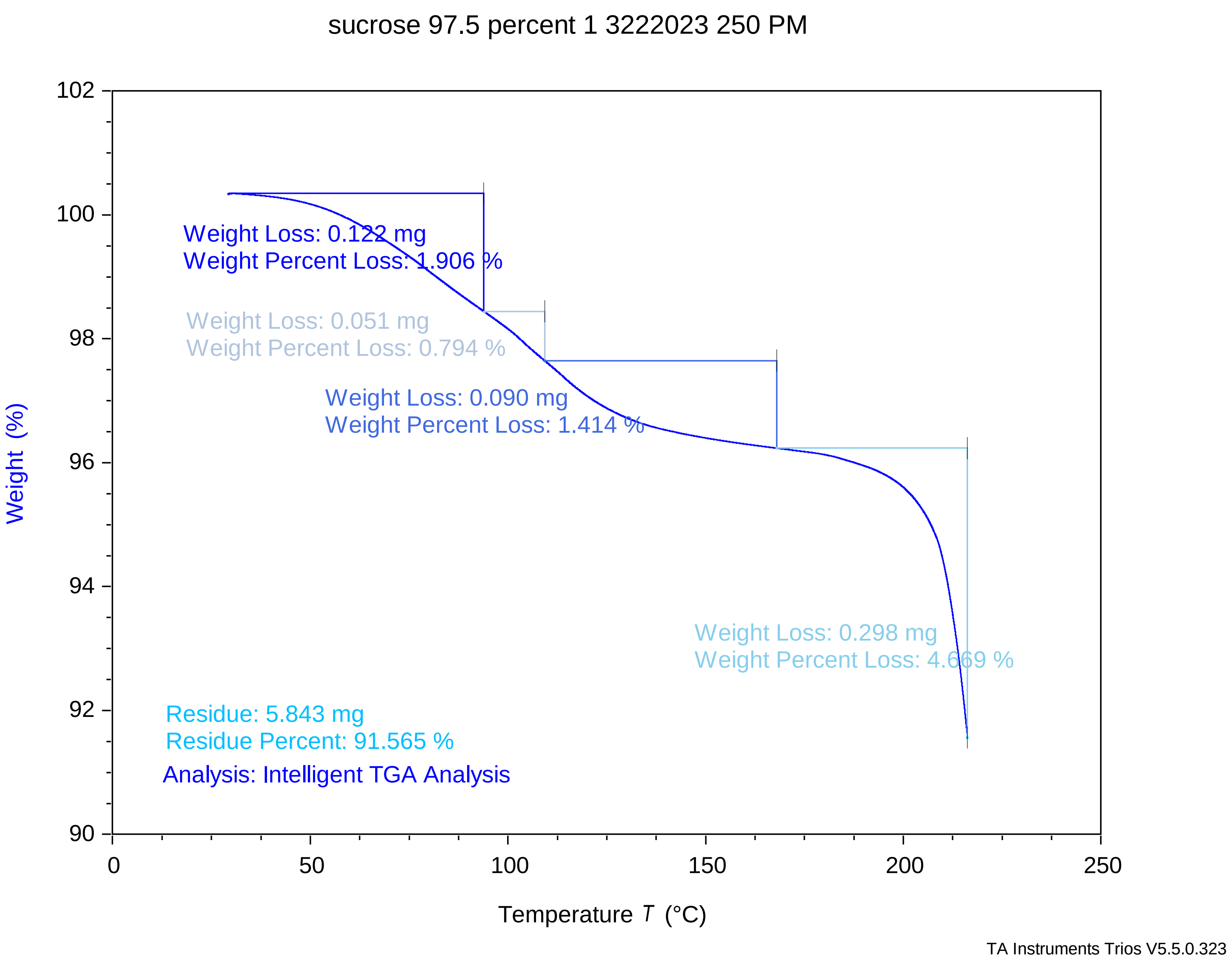

### sucrose 97.5 percent 2 3222023 319 pm.pdf

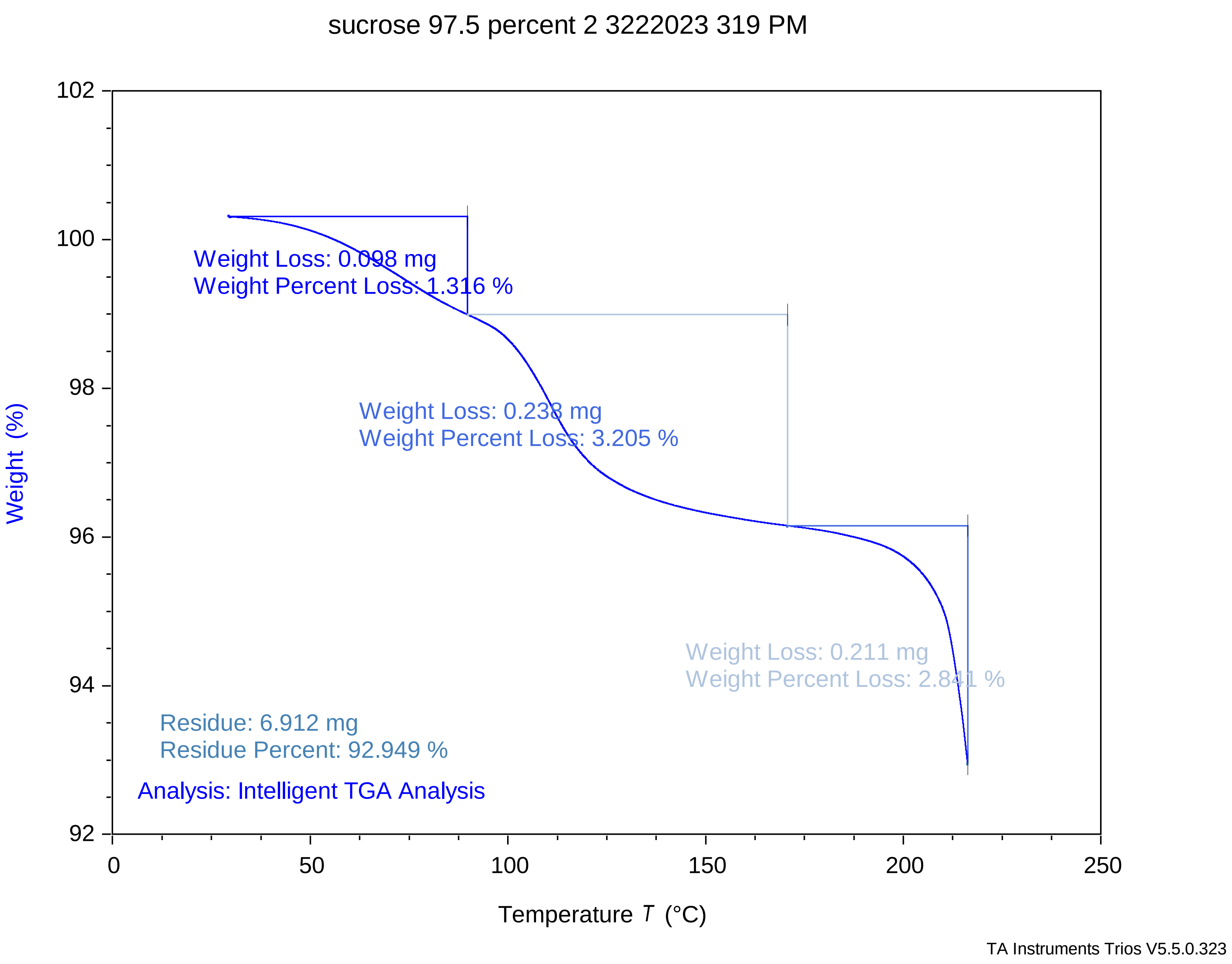

### sucrose 97.5 percent 3 3222023 348 pm.pdf

sucrose 97.5 percent 3 3222023 348 PM

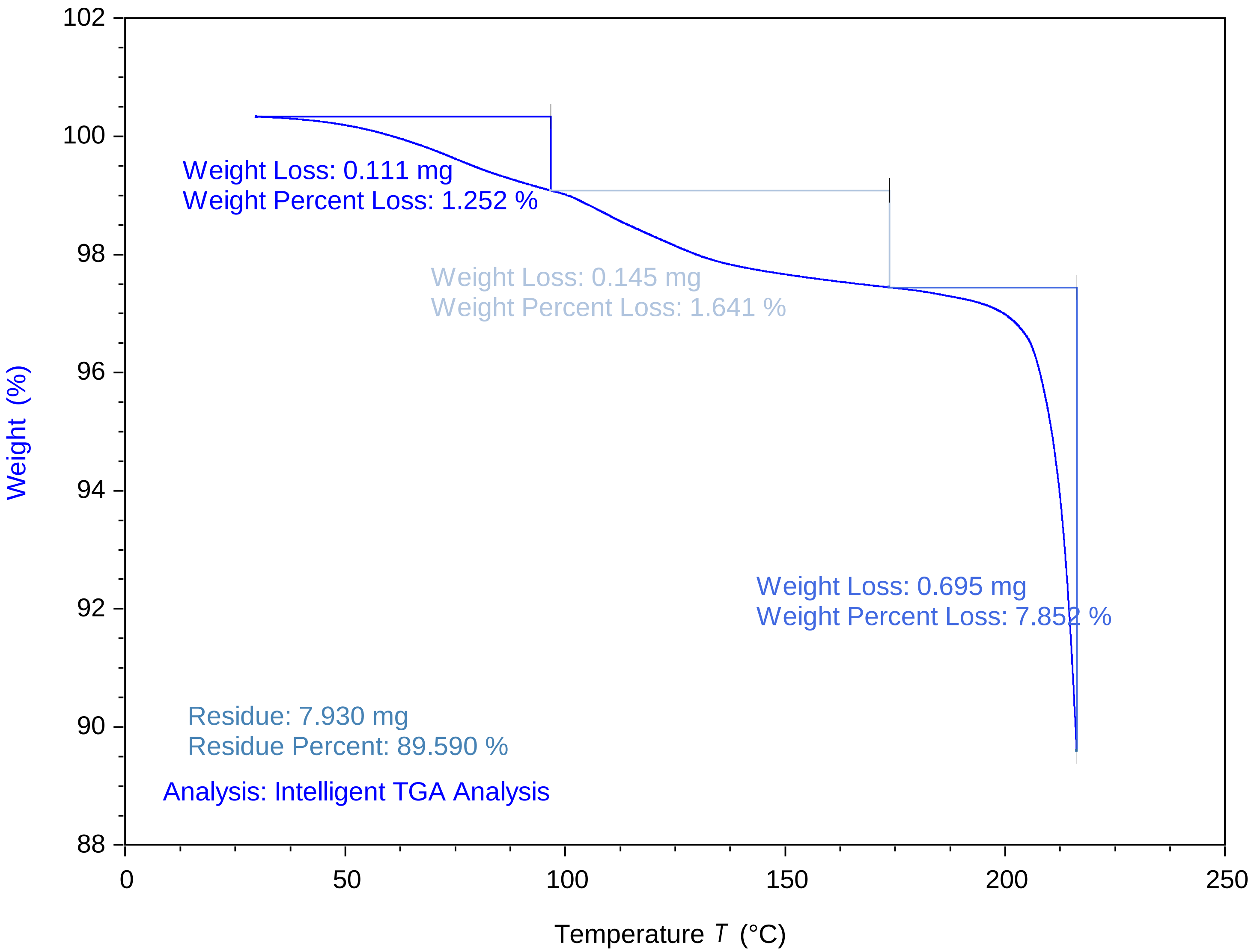


TA Instruments Trios V5.5.0.323

### sucrose 100 percent 1 3222023 126 PM.pdf

sucrose 100 percent 1 3222023 126 pm

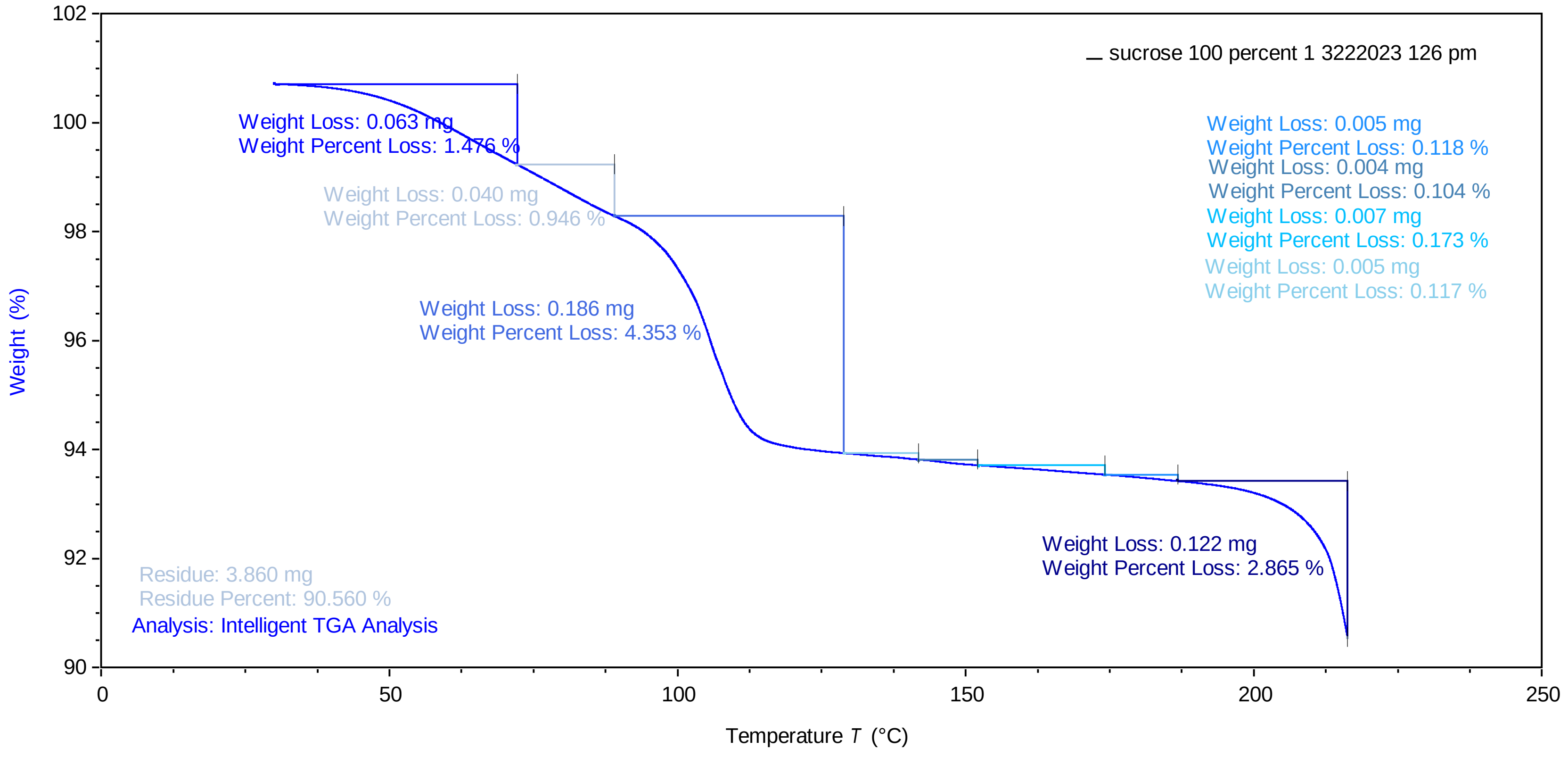


TA Instruments Trios V5.5.0.323

### sucrose 100 percent 2 3222023 153 pm.pdf

sucrose 100 percent 2 3222023 153 PM

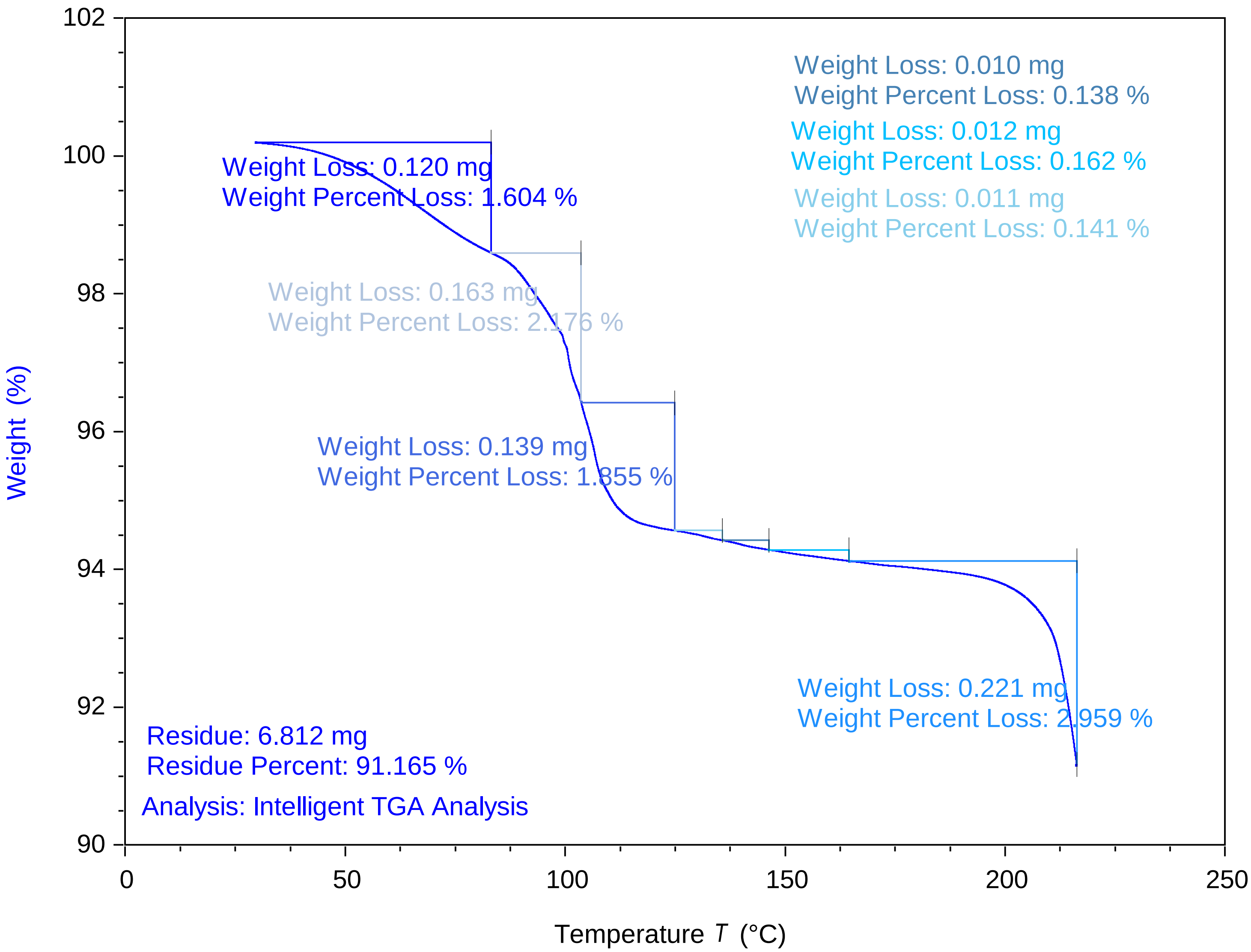


TA Instruments Trios V5.5.0.323

### sucrose 100 percent 3 3222023 222 pm.pdf

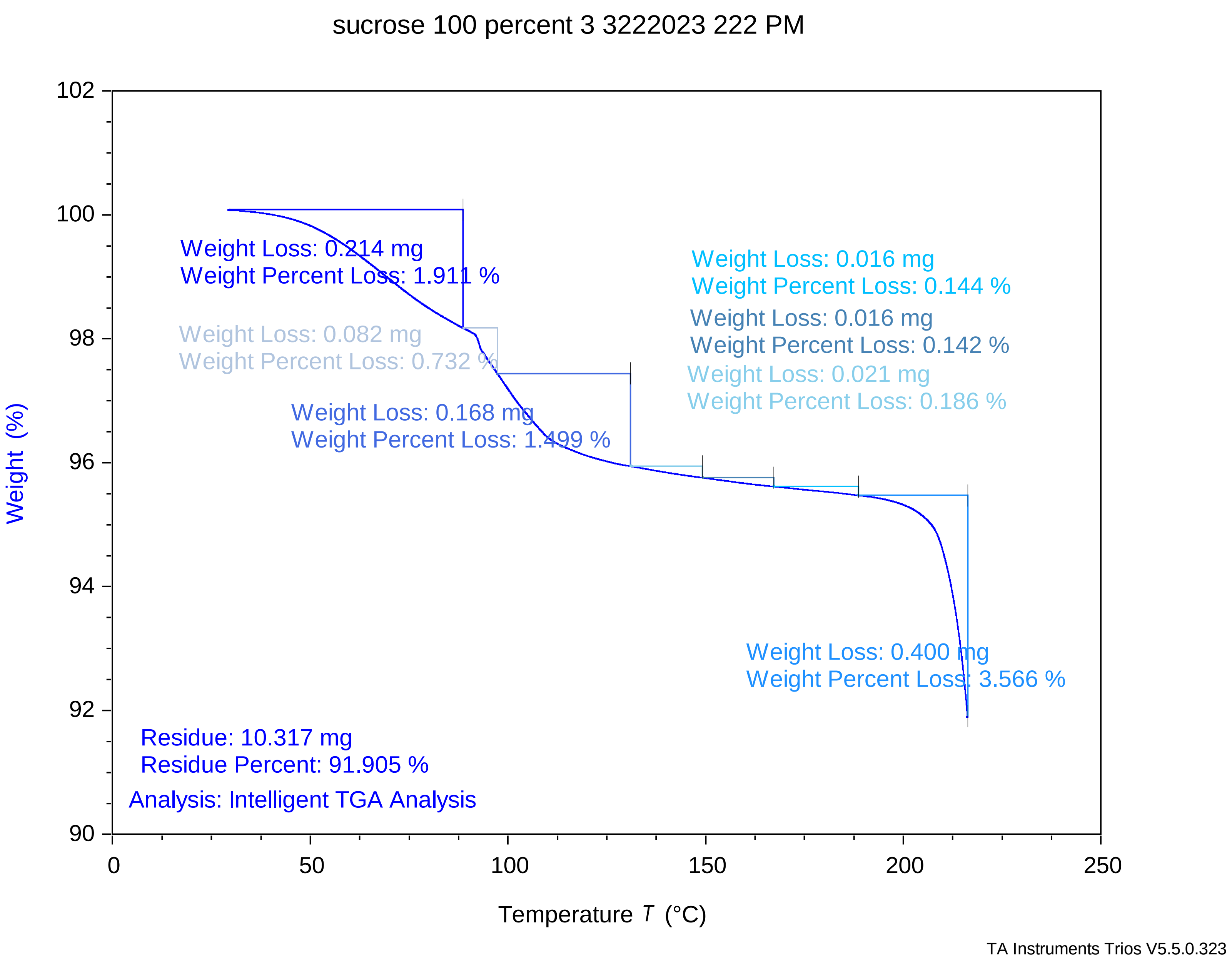

### Trehalose 10mg per mL 87.5 percent 1 412023 942 PM.pdf

Trehalose 10mg per mL 87.5 percent 1 412023 942

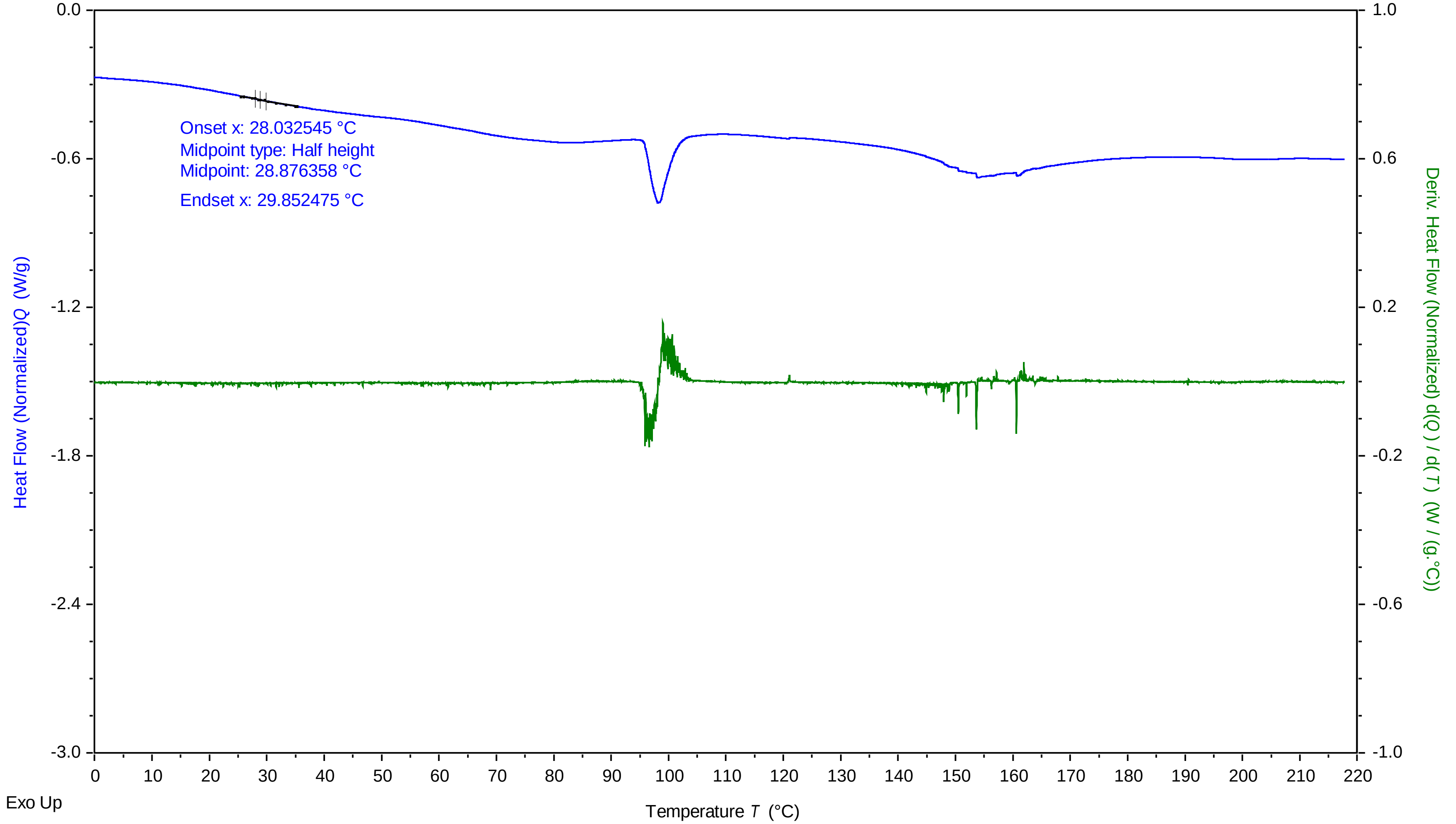


TA Instruments Trios V5.0.0.44616

### Trehalose 10mg per mL 87.5 percent 2 412023 1015 PM.pdf

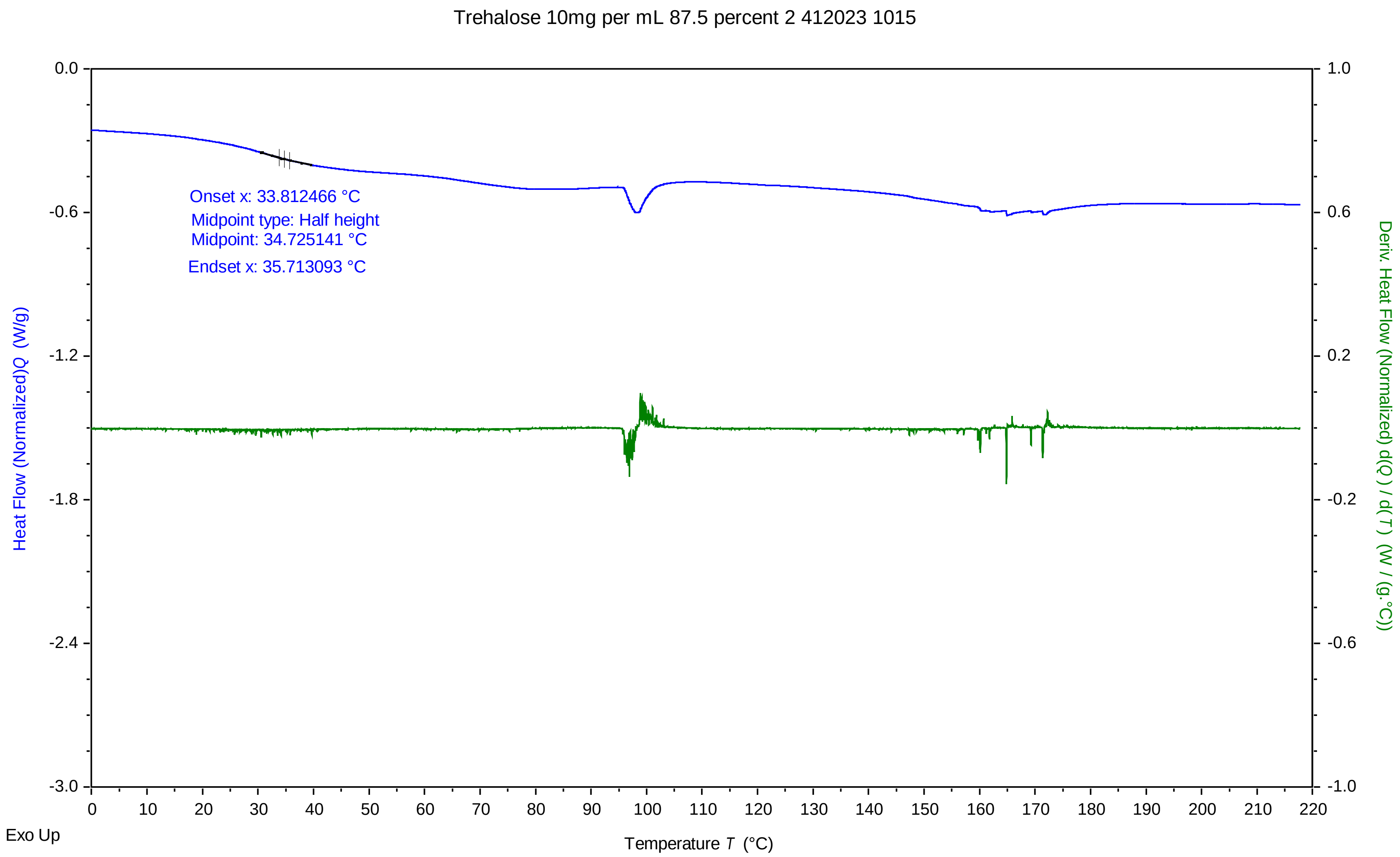


TA Instruments Trios V5.0.0.44616

### Trehalose 10mg per mL 87.5 percent 3 412023 1048 PM.pdf

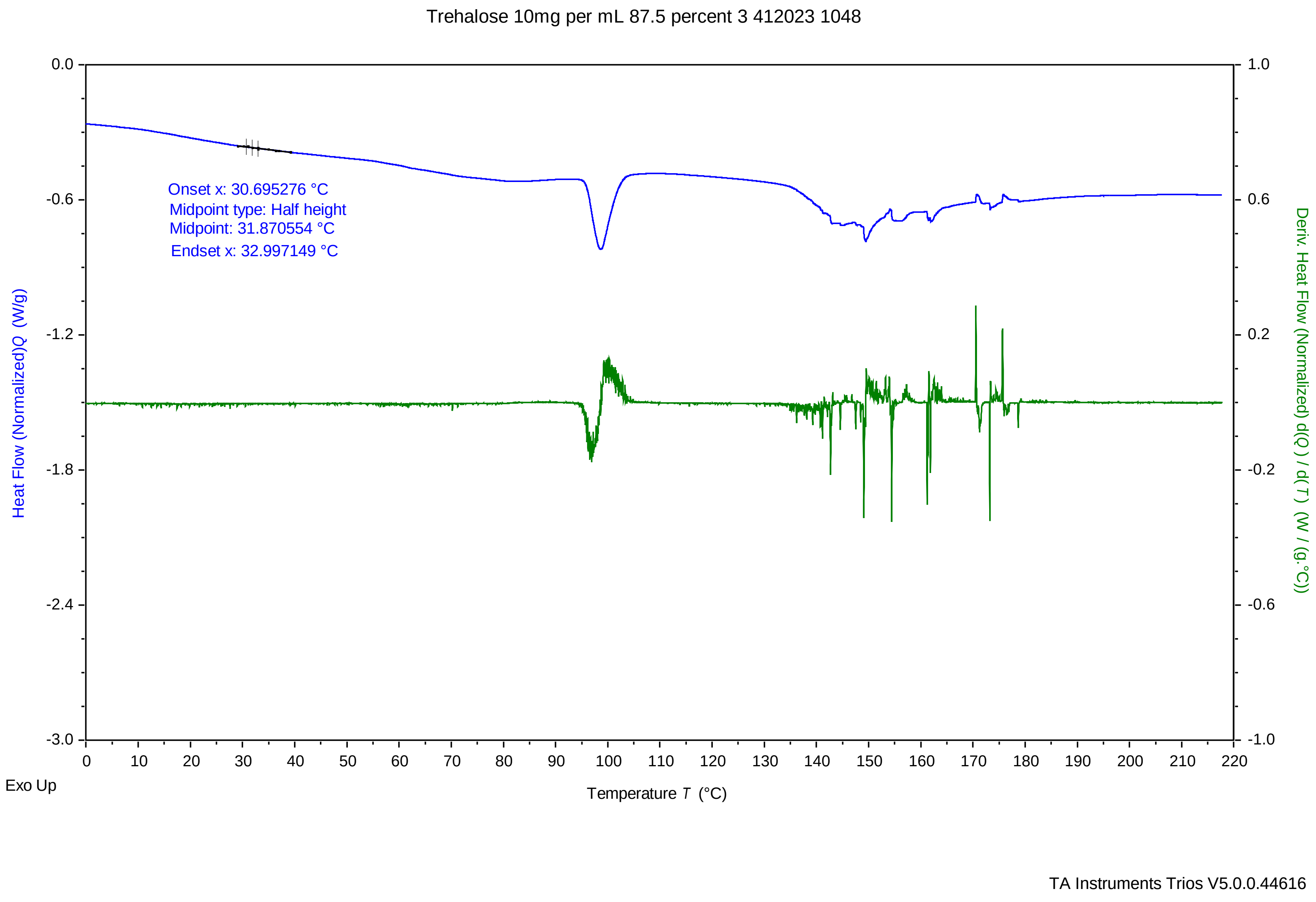

### Trehalose 10mg per mL 90 percent 1 412023 803 PM.pdf

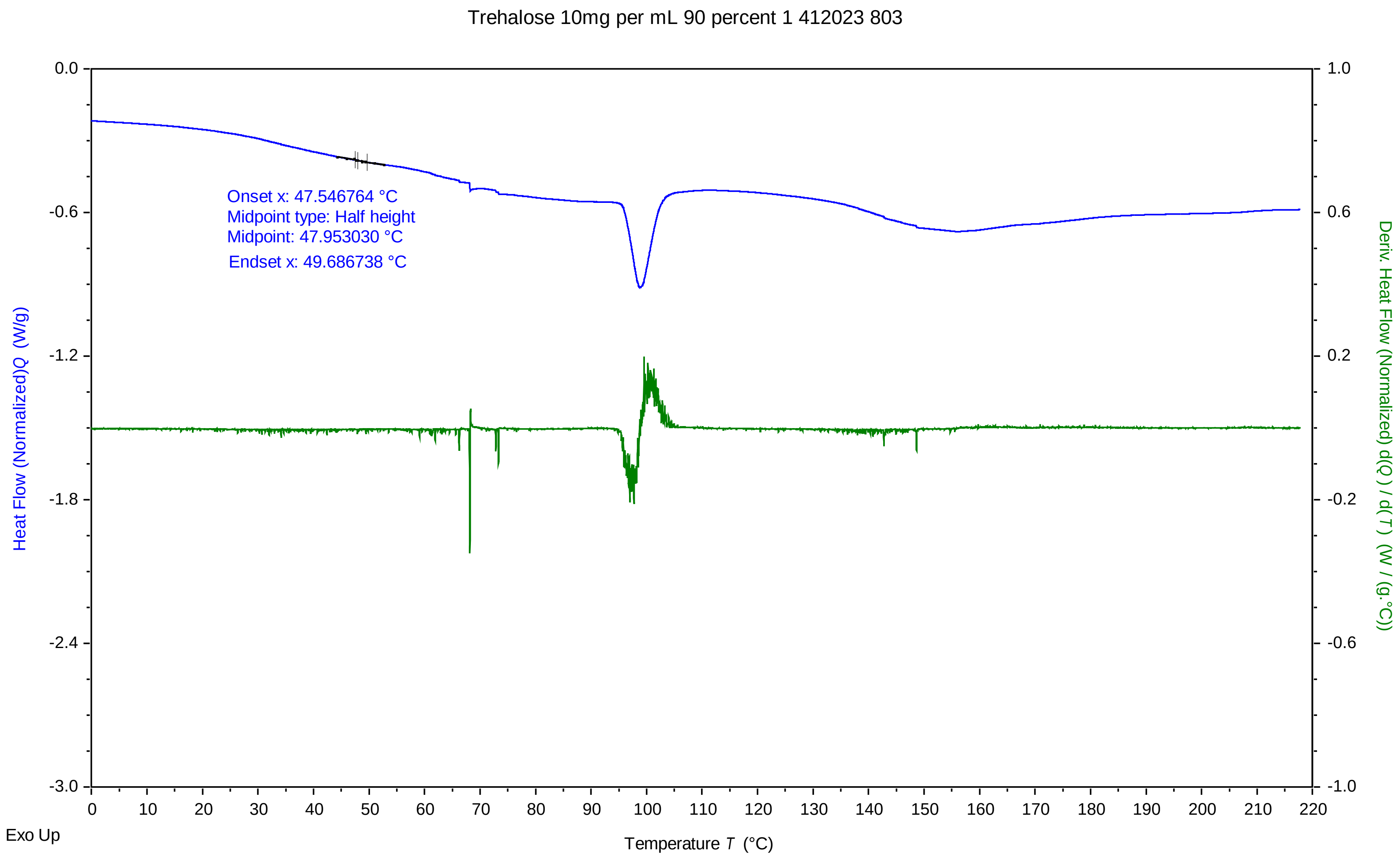


TA Instruments Trios V5.0.0.44616

### Trehalose 10mg per mL 90 percent 2 412023 836 PM.pdf

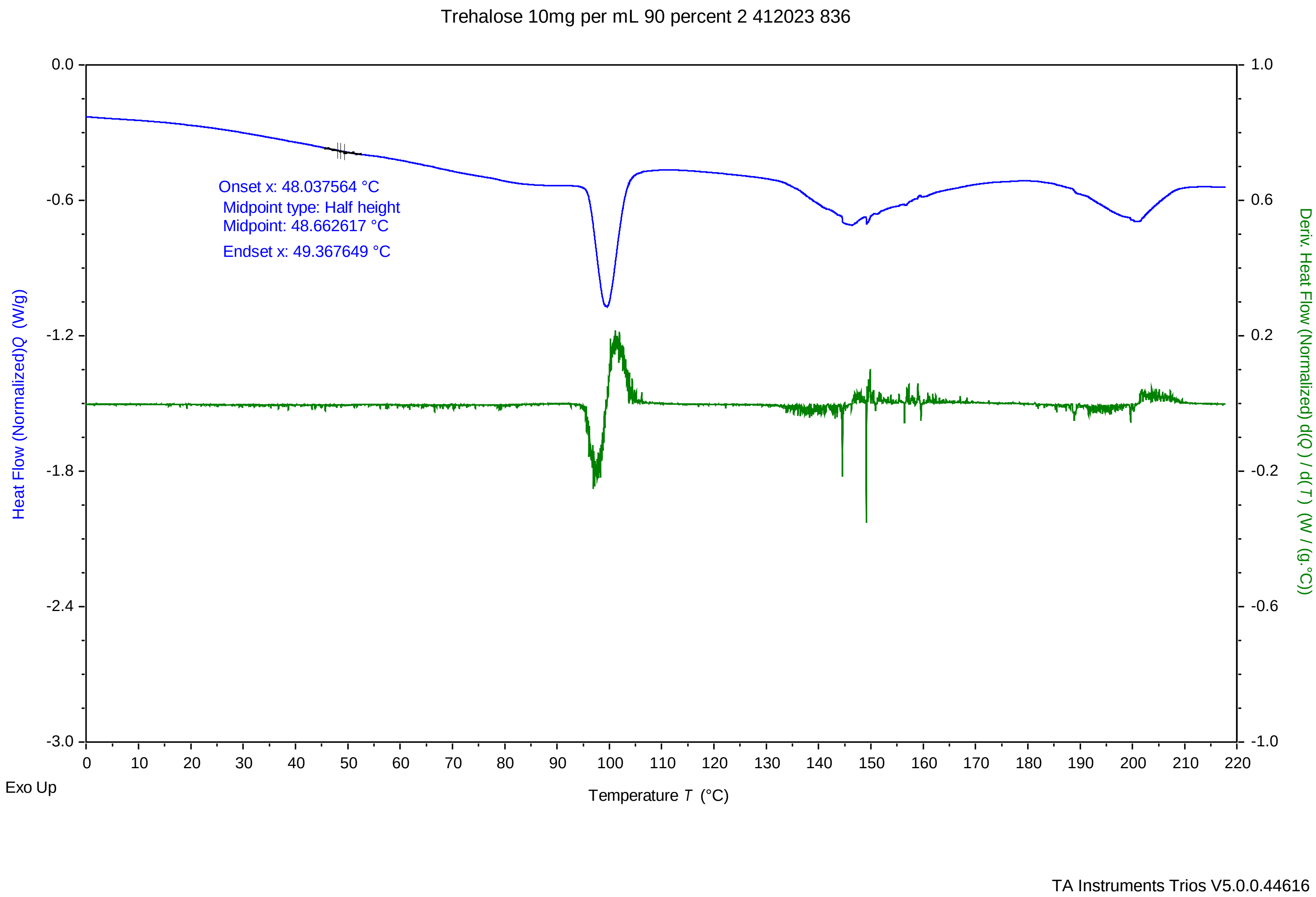

### Trehalose 10mg per mL 90 percent 3 412023 909 PM.pdf

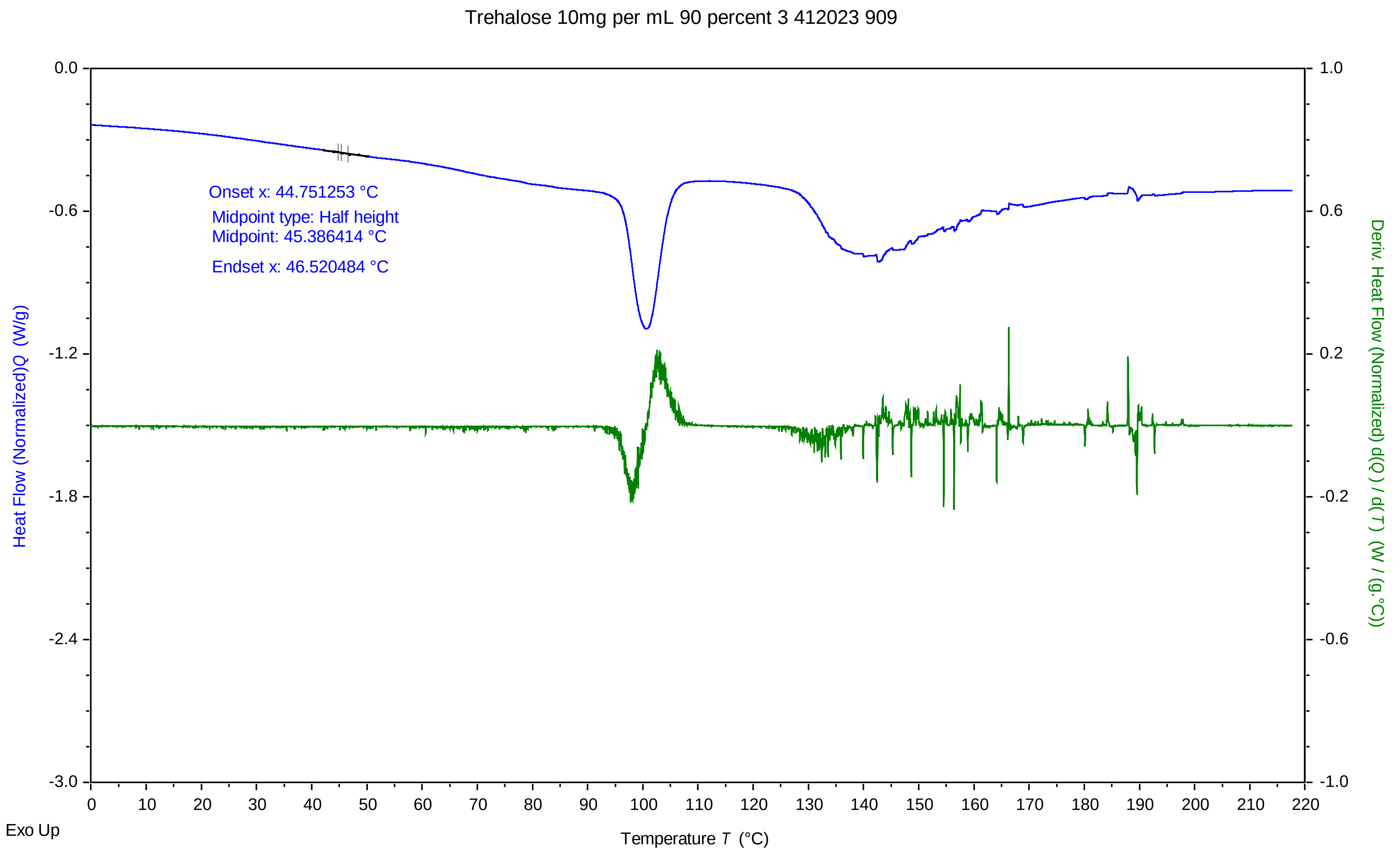


TA Instruments Trios V5.0.0.44616

### Trehalose 10mg per mL 92.5 percent 1 412023 624 PM.pdf

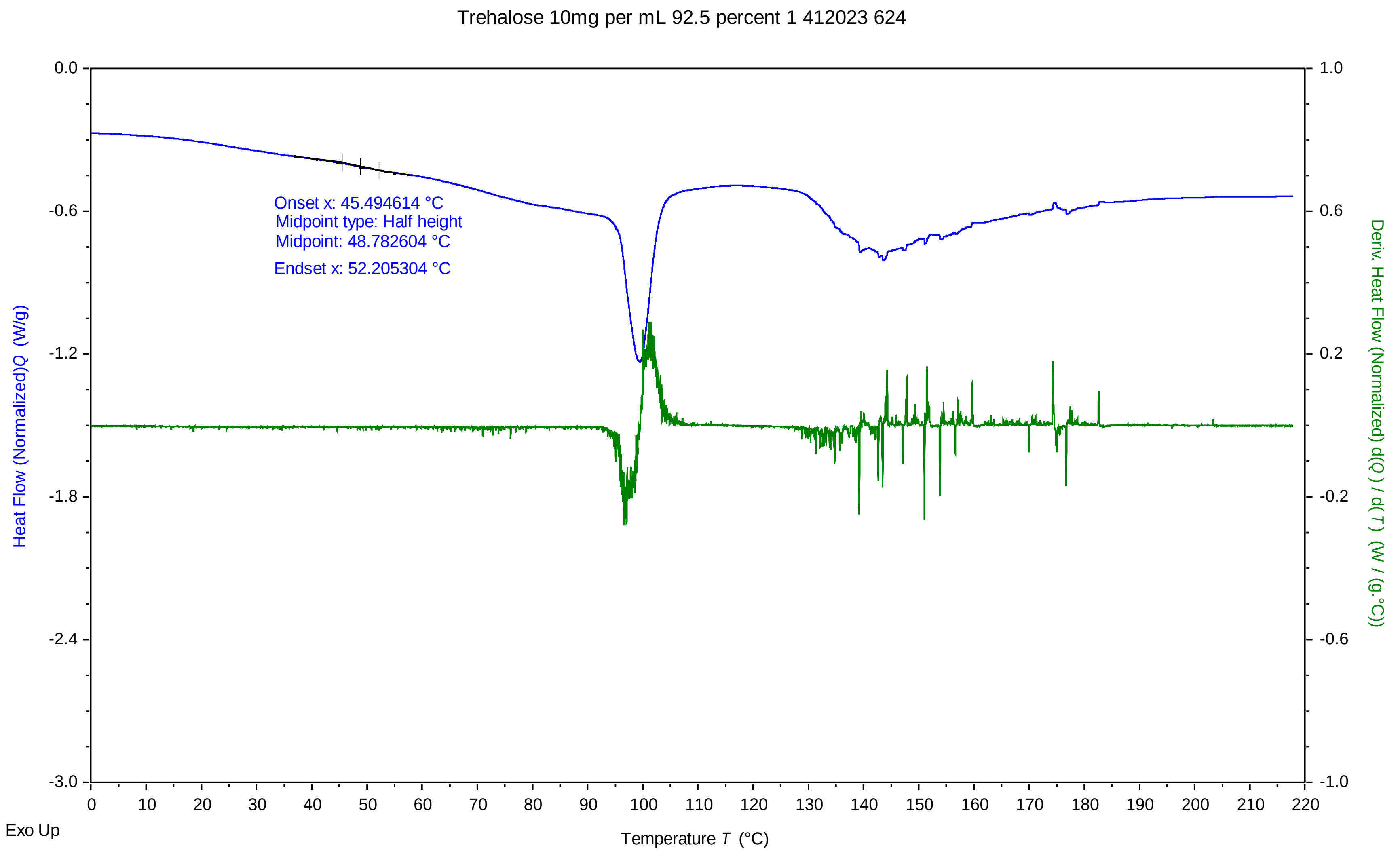


TA Instruments Trios V5.0.0.44616

### Trehalose 10mg per mL 92.5 percent 2 412023 657 PM.pdf

Trehalose 10mg per mL 92.5 percent 2 412023 657

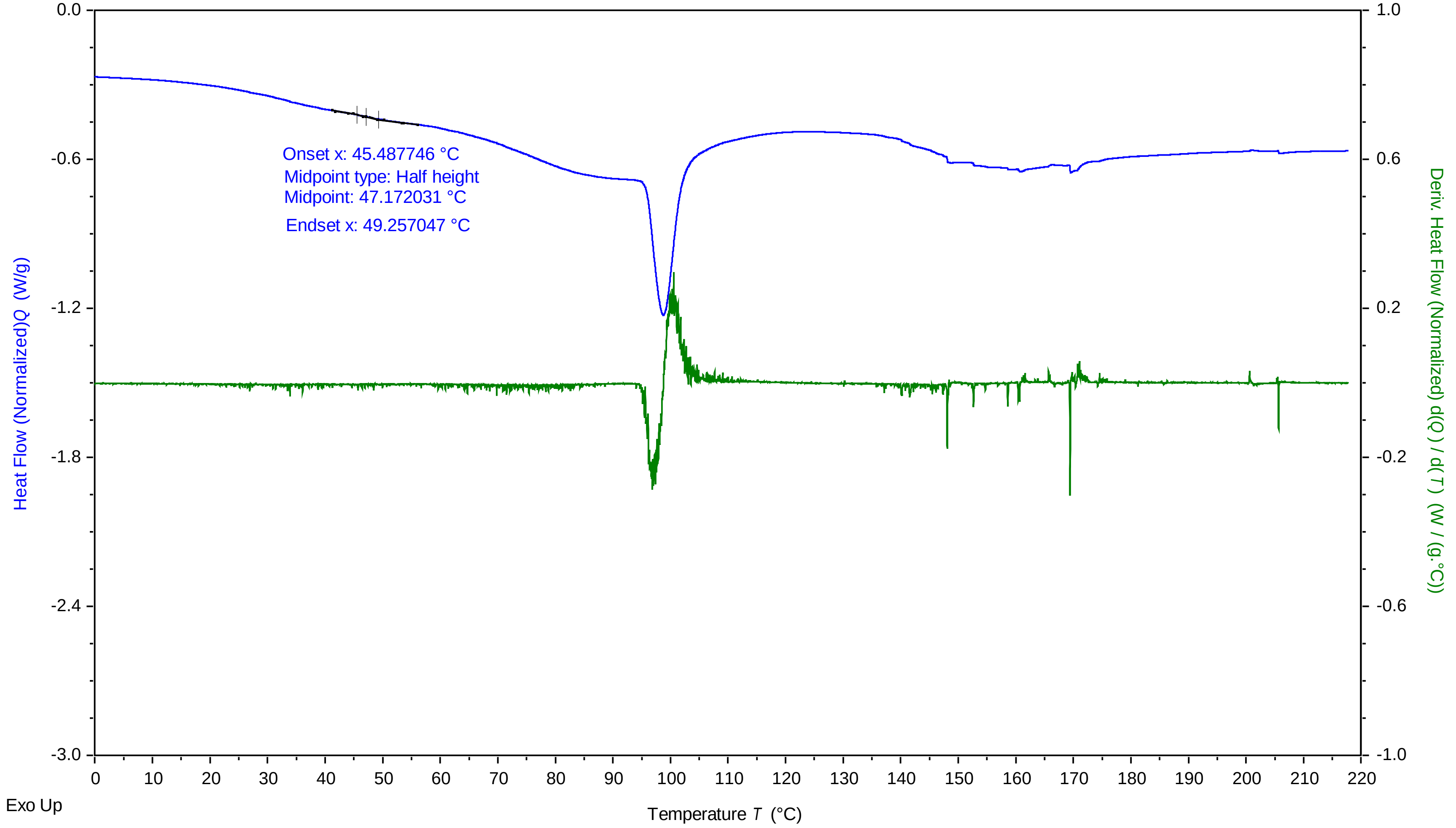


TA Instruments Trios V5.0.0.44616

### Trehalose 10mg per mL 92.5 percent 3 412023 730 PM.pdf

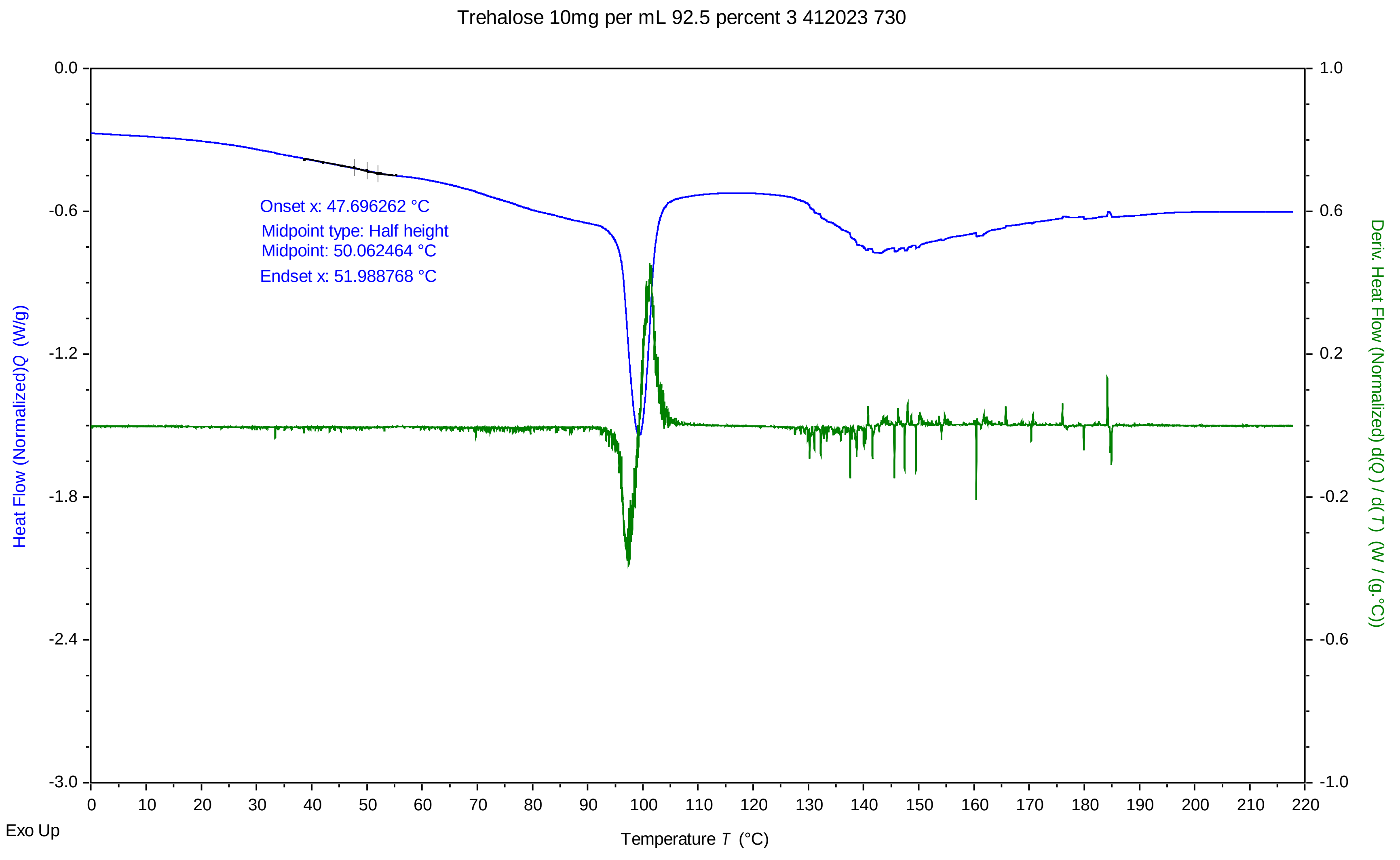


TA Instruments Trios V5.0.0.44616

### Trehalose 10mg per mL 95 percent 1 412023 444 PM.pdf

Trehalose 10mg per mL 95 percent 1 412023 444

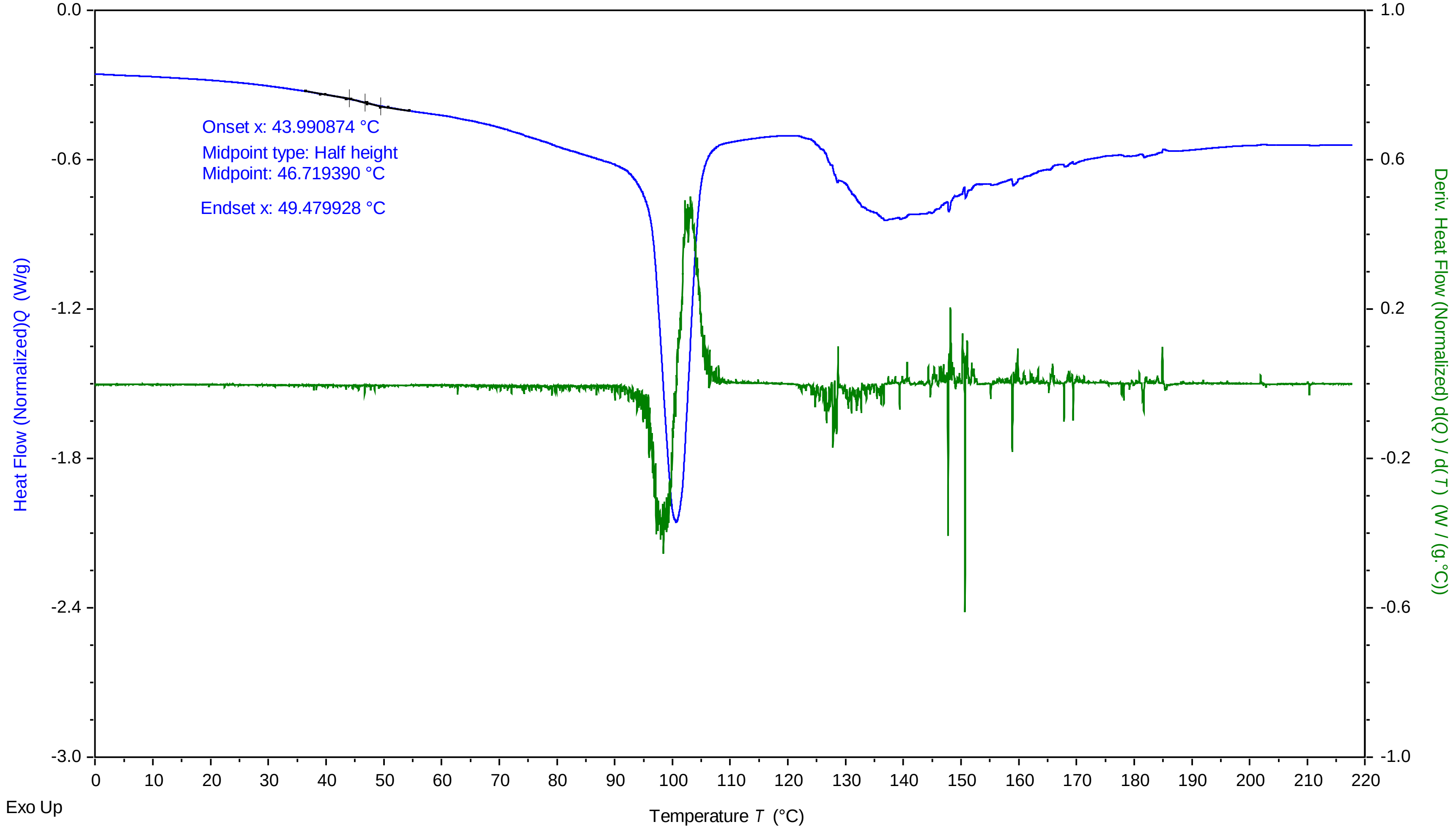


TA Instruments Trios V5.0.0.44616

### Trehalose 10mg per mL 95 percent 2 412023 517 PM.pdf

Trehalose 10mg per mL 95 percent 2 412023 517

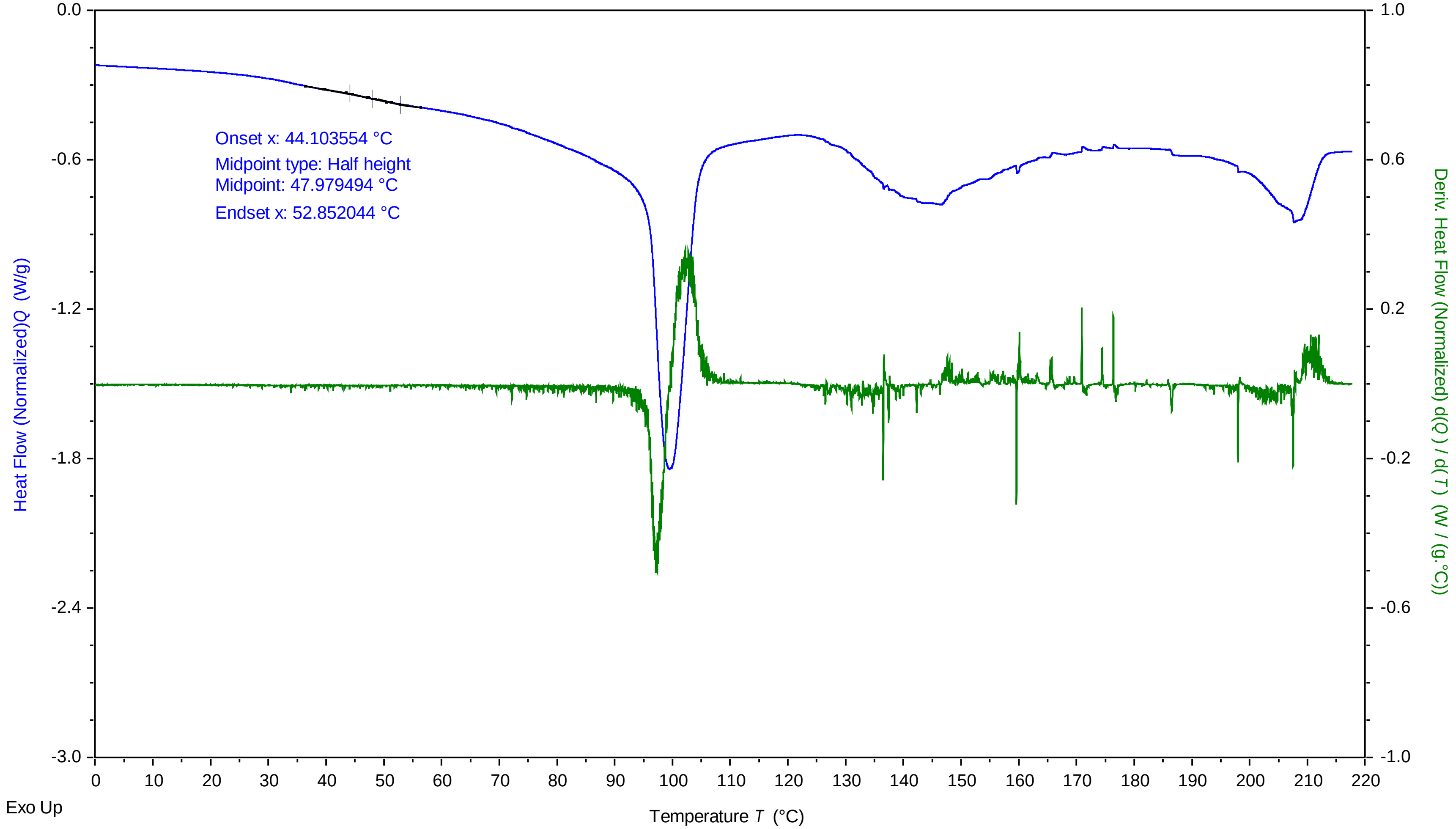


TA Instruments Trios V5.0.0.44616

### Trehalose 10mg per mL 95 percent 3 412023 551 PM.pdf

Trehalose 10mg per mL 95 percent 3 412023 551

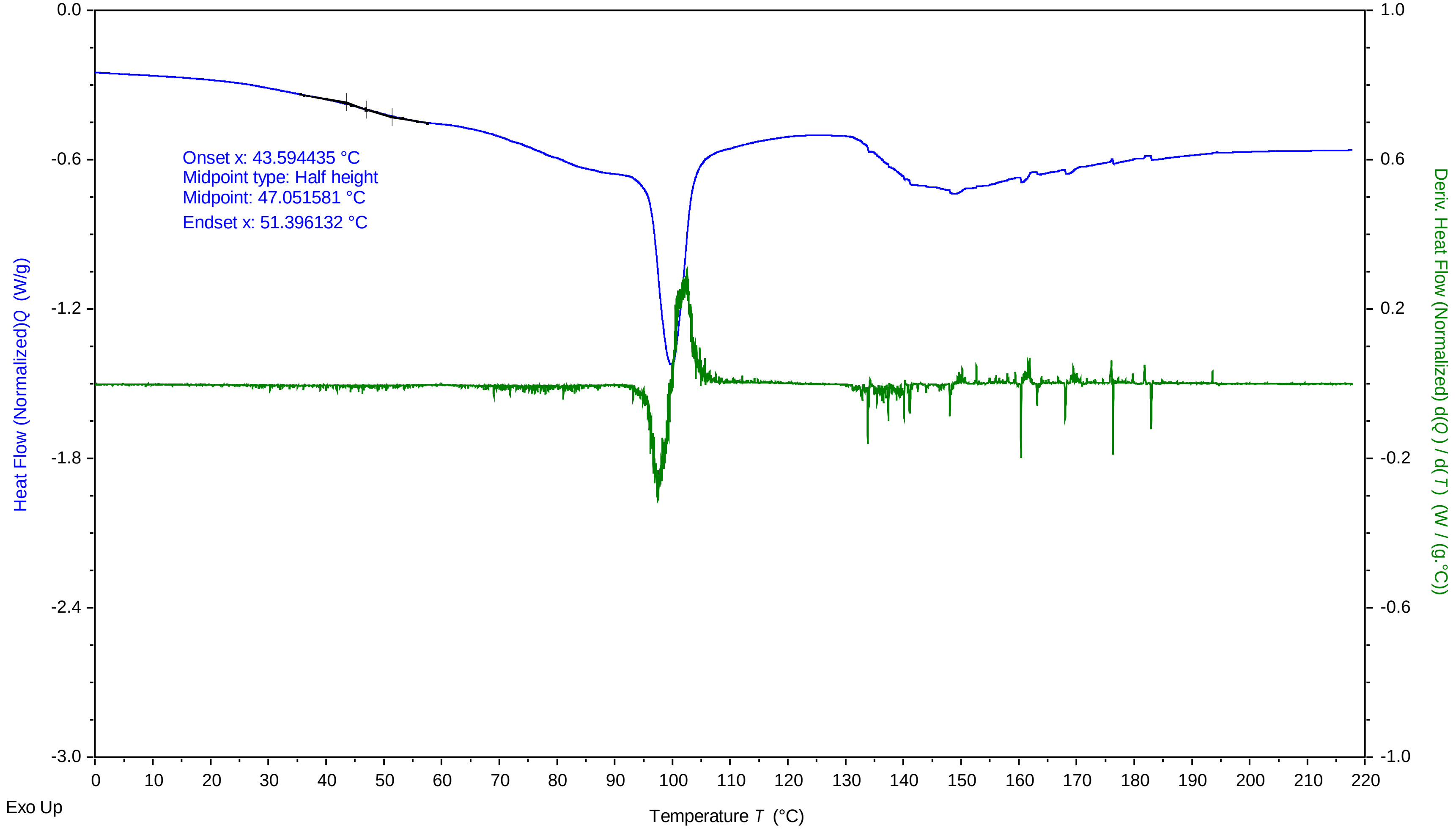


TA Instruments Trios V5.0.0.44616

### Trehalose 10mg per mL 97.5 percent 1 412023 305 PM.pdf

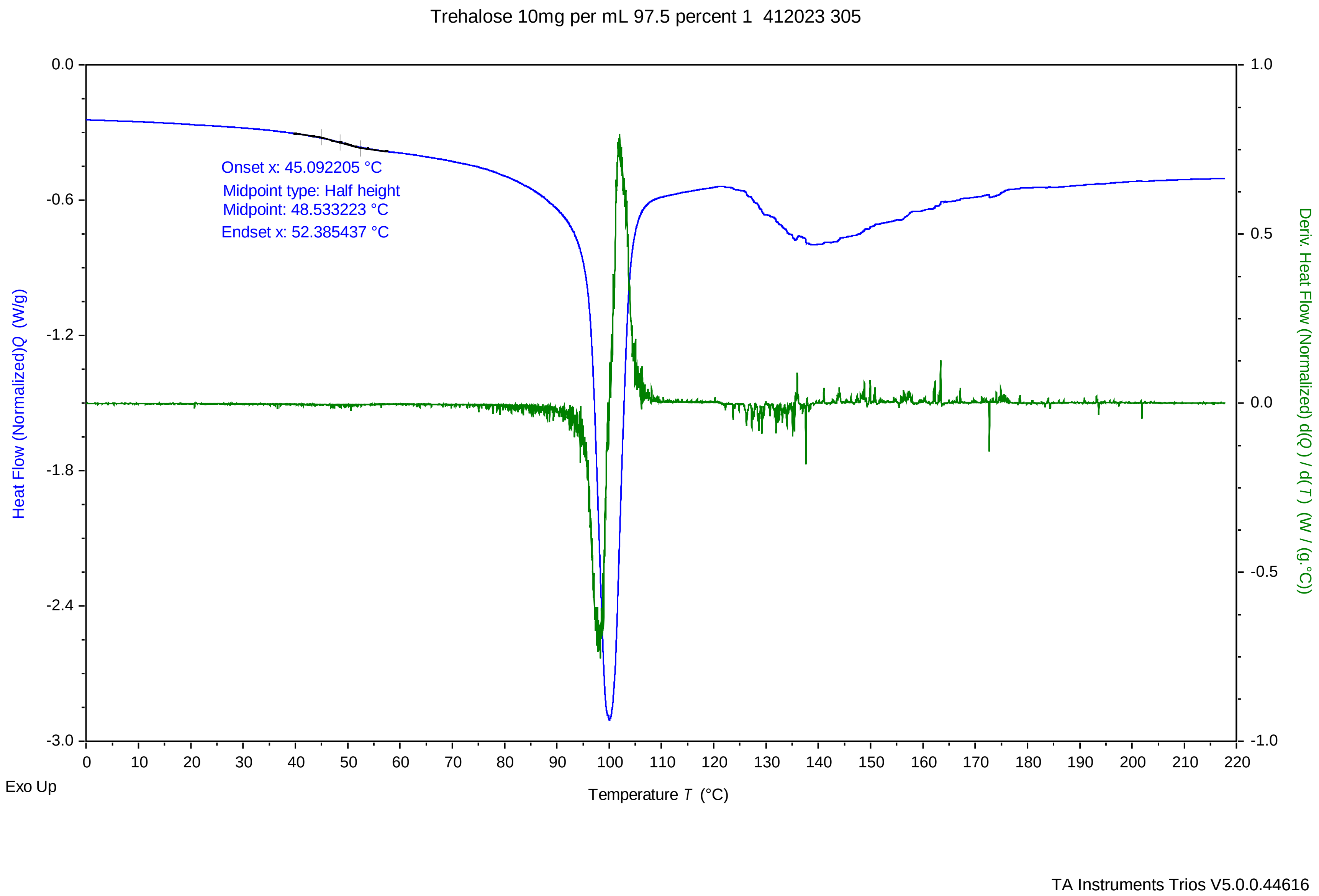

### Trehalose 10mg per mL 97.5 percent 2 412023 338 PM.pdf

Trehalose 10mg per mL 97.5 percent 2 412023 338

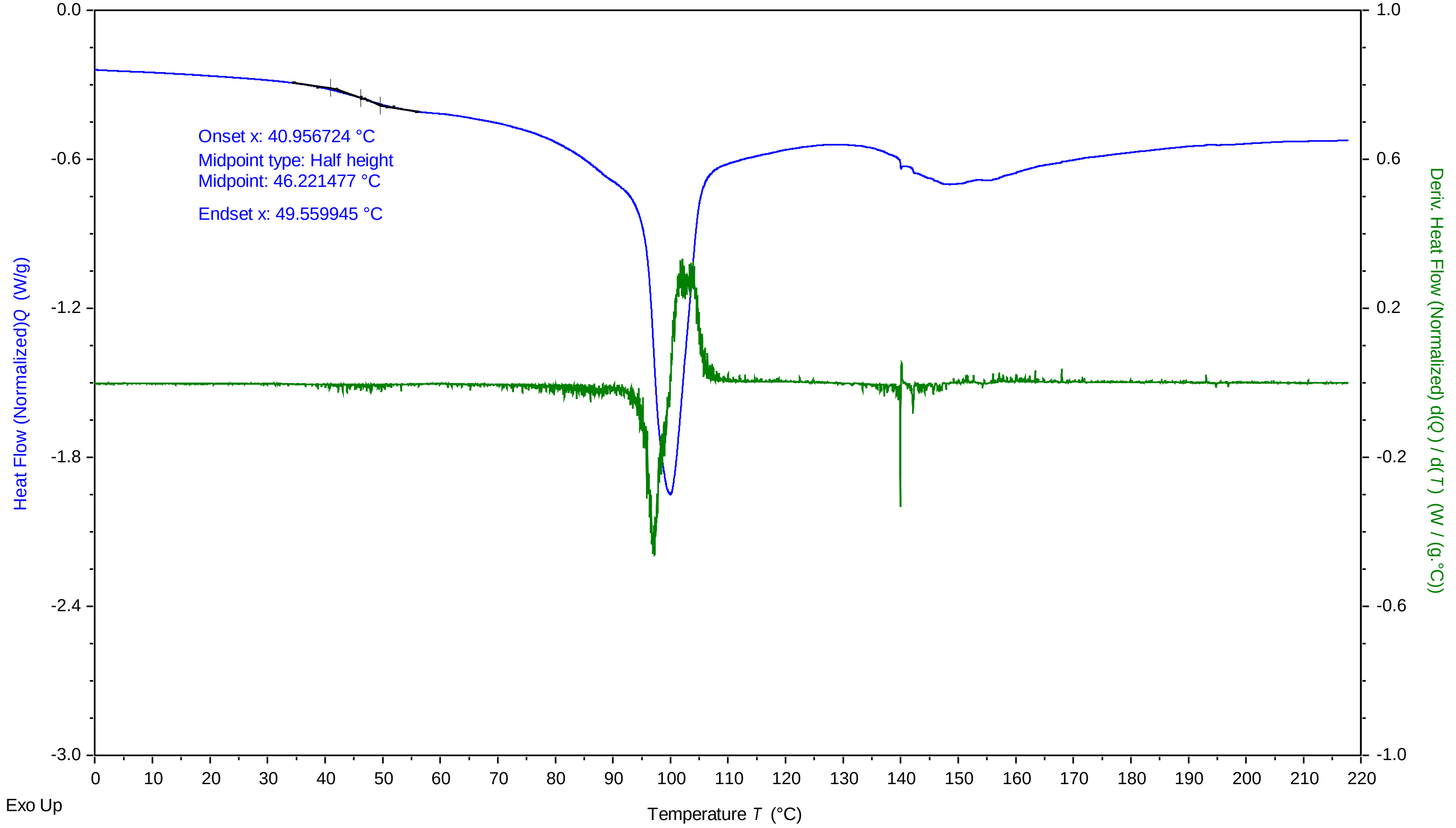


TA Instruments Trios V5.0.0.44616

### Trehalose 10mg per mL 97.5 percent 3 412023 411 PM.pdf

Trehalose 10mg per mL 97.5 percent 3 412023 411

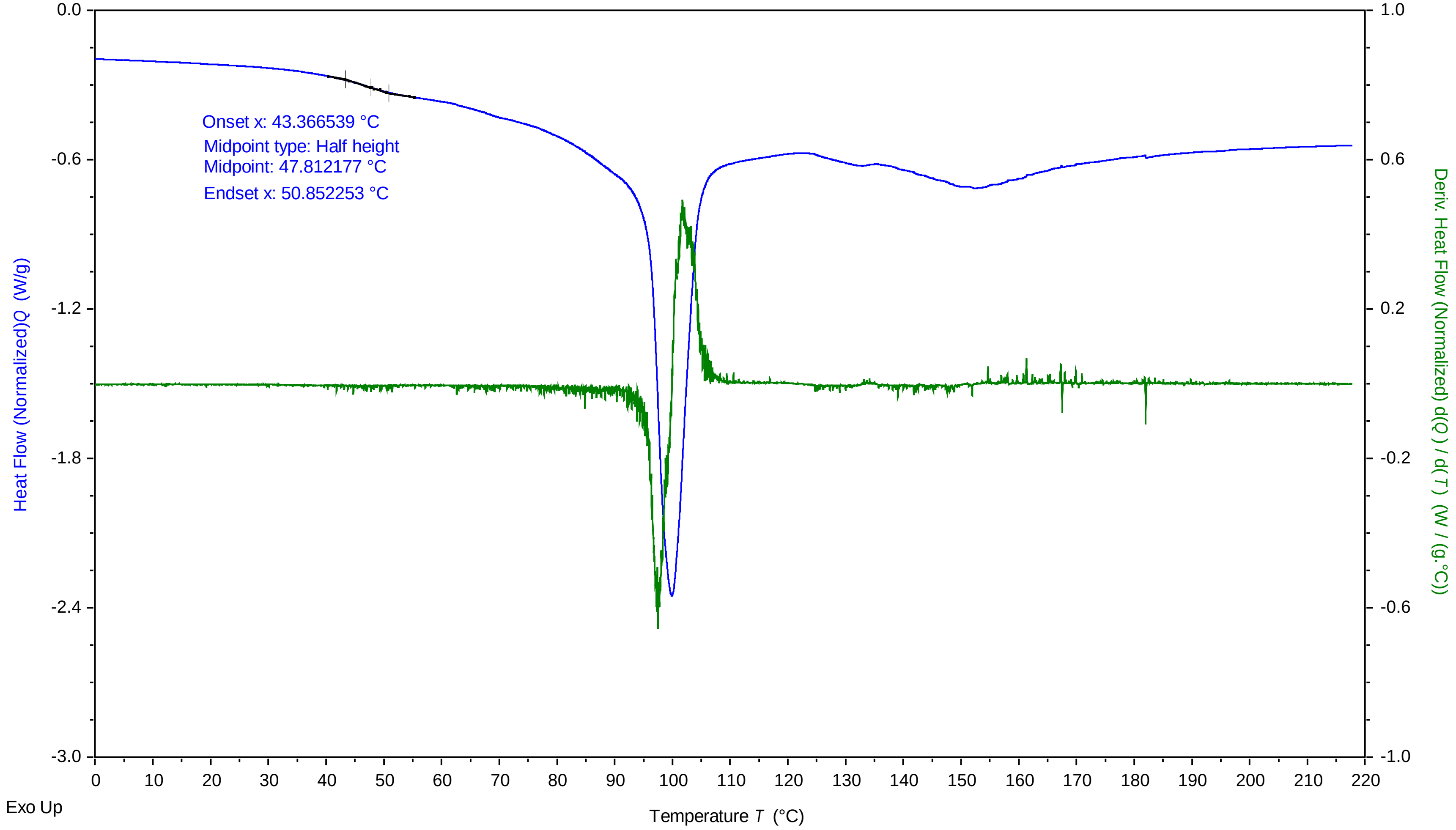


TA Instruments Trios V5.0.0.44616

### Trehalose 10mg per mL 100 percent 1 412023 126 PM.pdf

Trehalose 10mg per mL 100 percent 1 412023 126

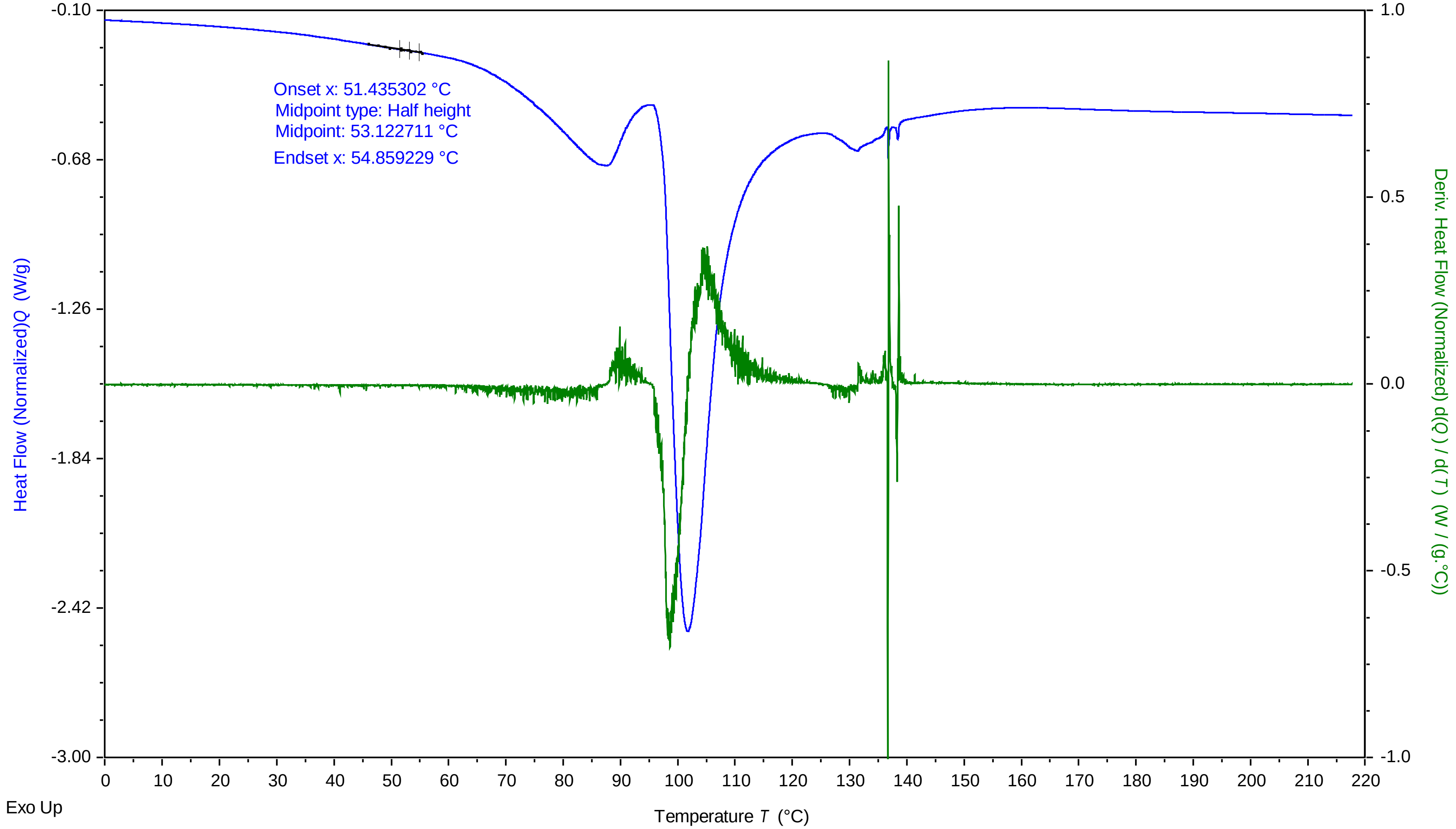


TA Instruments Trios V5.0.0.44616

### Trehalose 10mg per mL 100 percent 2 412023 159 PM.pdf

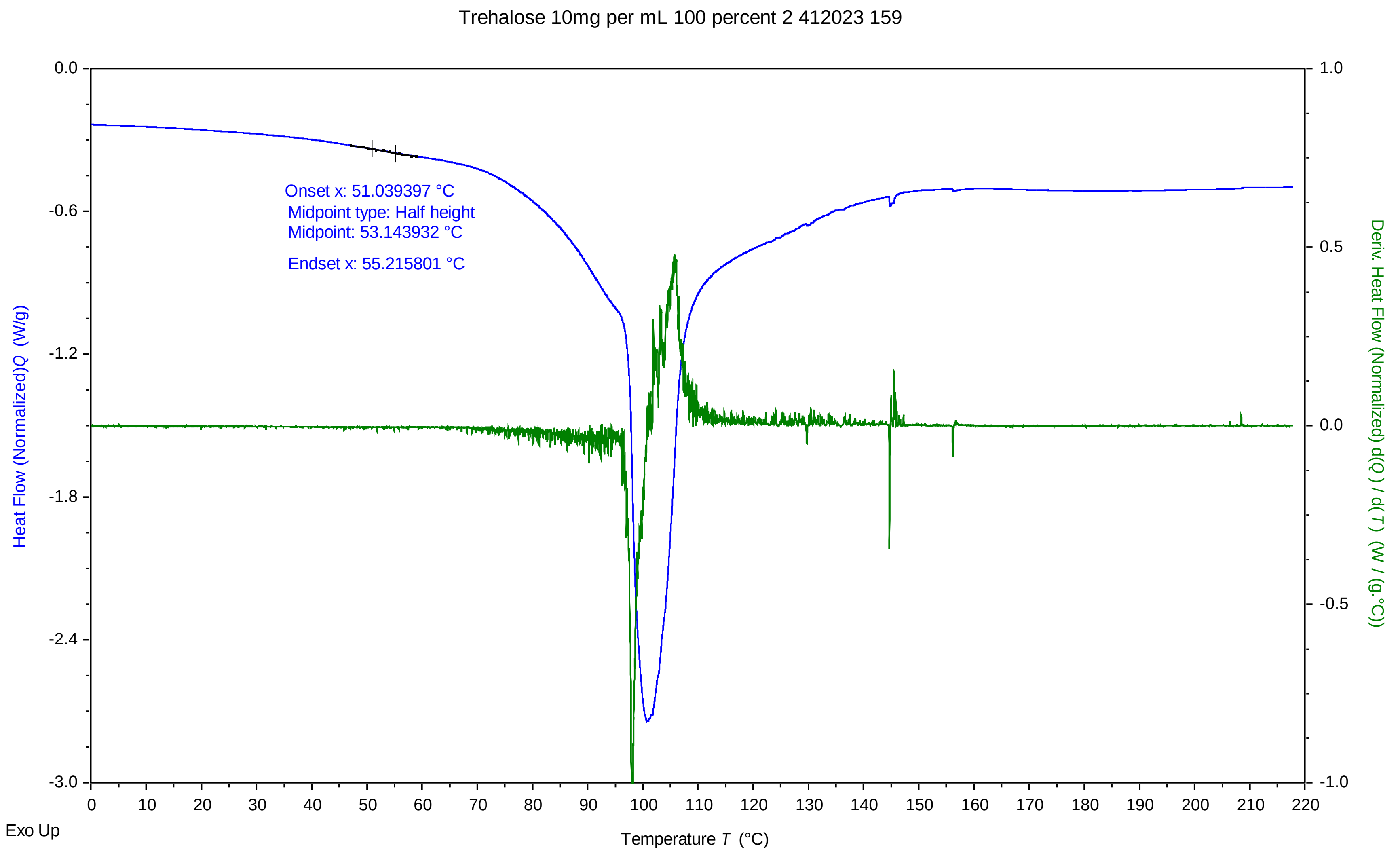


TA Instruments Trios V5.0.0.44616

### Trehalose 10mg per mL 100 percent 3 412023 232 PM.pdf

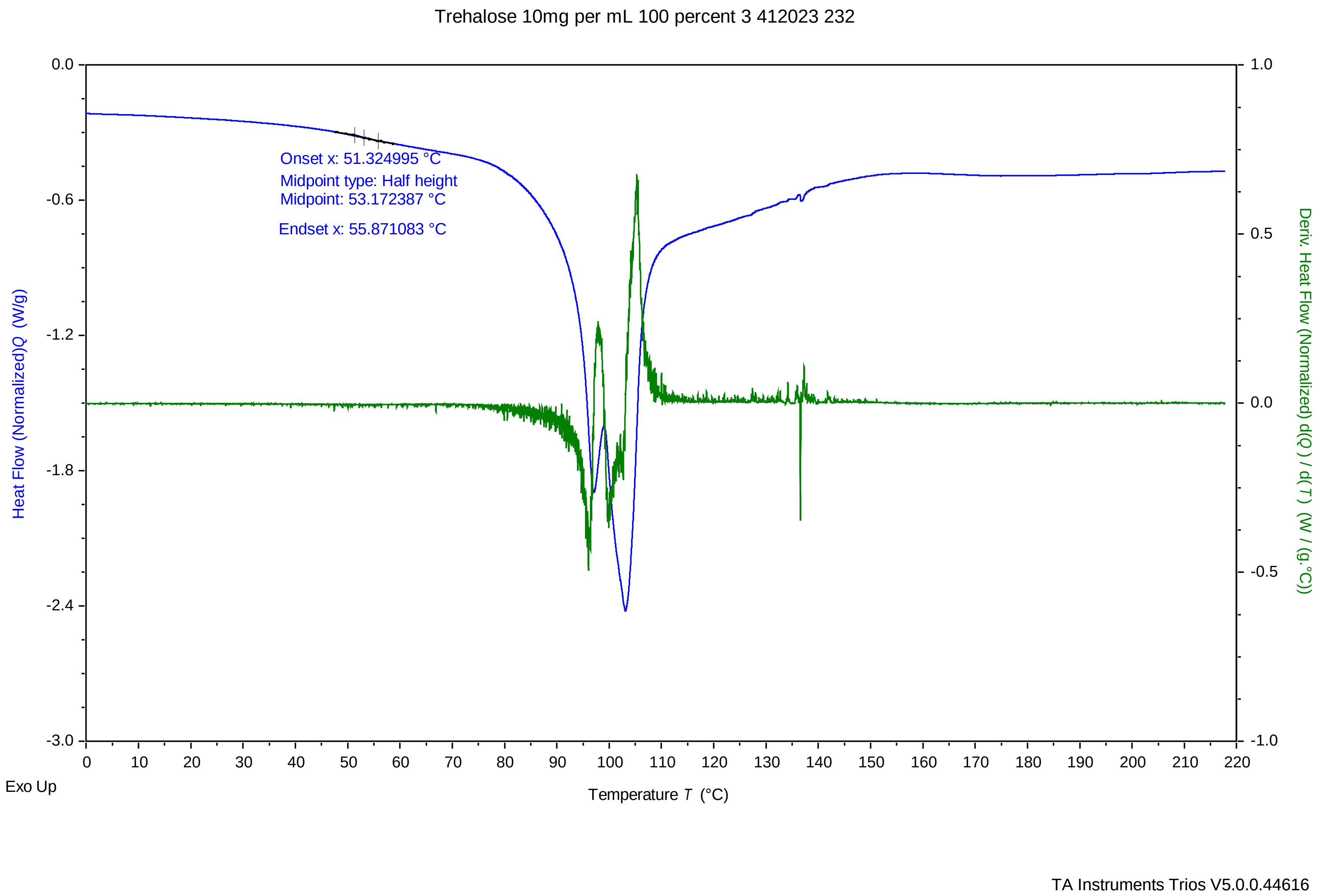

### view1.jpg

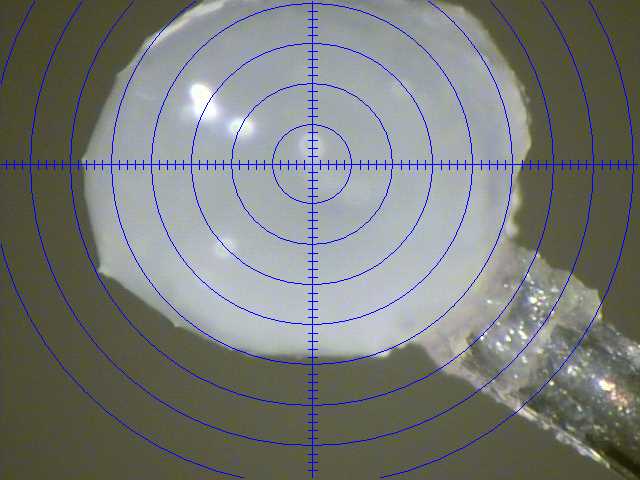

### view2.jpg

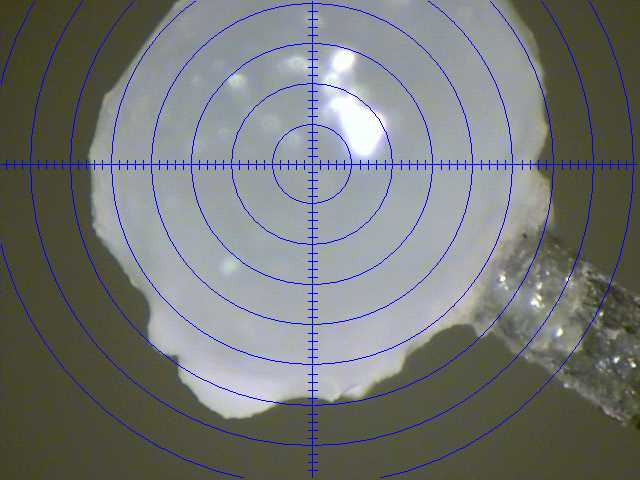
