## Supplementary material for "Water content, transition temperature and fragility influence protection and anhydrobiotic capacity": File S6: 2c_PerwaterVsGlycerol_bar.R

```
setwd("c:/R/Glass")

library(ggplot2) #load in libraries
library(gtools)
library(ggrepel)
library(readxl)
library(ggsignif)
library(ggpubr)
library(ggprism)
library(polynom)
library(hrbrthemes)
library(tidyverse)
library(rstatix)
library(nplyr)
library(patchwork)
library(dplyr)
source("stat_cor.R")
source("utilities.R")
source("utilities_label.R")
source("ANOVA_Tukey.R")

Treh <- read_excel("Treh_glyc2.xlsx", sheet = "Trehalose")
Treh2 <- read_excel("Treh_glyc2.xlsx", sheet = "Sheet1")
#reading excel sheet into var Treh

Sucr <- read_excel("Sucr_glyc.xlsx", sheet = "Sucrose")
Sucr2 <- read_excel("Sucr_glyc.xlsx", sheet = "Sheet1")
#reading excel sheet into var Sucr

Malt <- read_excel("Malt_glyc2.xlsx", sheet = "Maltose")
Malt2 <- read_excel("Malt_glyc2.xlsx", sheet = "Sheet1")
#reading excel sheet into var Sucr

df1 <- data.frame(y = Treh$Perwater, x = Treh$Glycerol, CI = Treh$Perwater_CI, 
                  Disaccharide = Treh$Sugar)
#read x and y variables into dataframe df1

df2 <- data.frame(y = Sucr$Perwater, x = Sucr$Glycerol, CI = Sucr$Perwater_CI, 
                  Disaccharide = Sucr$Sugar)
#read x and y variables into dataframe df1

df3 <- data.frame(y = Malt$Perwater, x = Malt$Glycerol, CI = Malt$Perwater_CI, 
                  Disaccharide = Malt$Sugar)
#read x and y variables into dataframe df1


df4 <- data.frame(y = Treh2$Perwater, x = Treh2$Glycerol, CI = Treh2$Perwater_CI, 
                  Disaccharide = Treh2$Sugar)
#read x and y variables into dataframe df1

df5 <- data.frame(y = Sucr2$Perwater, x = Sucr2$Glycerol, CI = Sucr2$Perwater_CI, 
                  Disaccharide = Sucr2$Sugar)
#read x and y variables into dataframe df1

df6 <- data.frame(y = Malt2$Perwater, x = Malt2$Glycerol, CI = Malt2$Perwater_CI, 
                  Disaccharide = Malt2$Sugar)

#read x and y variables into dataframe df1

df <- rbind(df1, df2, df3) %>% 
    mutate(x = as_factor(x), 
           x = fct_recode(x, "100.0%" = "0", "97.5%"= "2.5", "95.0%"= "5", 
                          "92.5%"= "7.5", "90.0%"= "10", "87.5%"= "12.5"), 
           x = fct_relevel(x, c("100.0%", "97.5%", "95.0%", "92.5%", "90.0%", 
                                "87.5%"))) 

dfpoint <- rbind(df4, df5, df6) %>% 
    mutate(x = as_factor(x), 
           x = fct_recode(x, "100.0%" = "0", "97.5%"= "2.5", "95.0%"= "5", 
                          "92.5%"= "7.5", "90.0%"= "10", "87.5%"= "12.5"), 
           x = fct_relevel(x, c("100.0%", "97.5%", "95.0%", "92.5%", "90.0%", 
                                "87.5%"))) 

#ANOVA analysis
rawdata_nest <- dfpoint %>% 
    drop_na() %>%
    group_nest(Disaccharide)

hf_fit <- rawdata_nest %>%
    mutate(fitaov = map(data, do_aov),
           fittukey = map(fitaov, do_tukey))  %>%
    nest_summarise(data, max_y = max(y)) %>%
    select(-fitaov) %>%
    unnest(cols = c(data, fittukey))

write.csv(dfpoint, "PerwaterVsGlycerol_bar.csv", row.names=FALSE)

dfpoint1 <- dfpoint %>% filter(Disaccharide == "Maltose")
hf_fit1 <- hf_fit %>% filter(Disaccharide =="Maltose") %>% filter(p.adj <= 0.05)
p1 <- df %>% filter(Disaccharide == "Maltose") %>%
    ggplot(aes(x = x, y = y)) +
    geom_bar(stat = "identity", colour = "black", position = "dodge", 
             scales = "free_y", fill = "tomato") +
    geom_errorbar(aes(ymin = y-CI, ymax = y+CI), width=0.5, size=1, 
                  position = position_dodge(width = 0.90)) +
    geom_point(data = dfpoint1, aes(x = x, y = y, fill = Disaccharide), fill = "tomato", shape = 21, 
               size = 1, stroke = 2, position = position_dodge(width = 0.90)) +
    #ggtitle("Disaccharide content (%) versus water content (%)") +
    theme_bw() +
    theme(plot.title = element_text(hjust = 0.5), strip.text.x = element_blank(), 
          legend.position = "none", axis.title.x=element_blank(), axis.title.y=element_blank()) +
    stat_pvalue_manual(data = hf_fit1, y.position = 6, step.increase = 0.2, 
                       label = "p.adj.signif", size = 4) 

dfpoint2 <- dfpoint %>% filter(Disaccharide == "Sucrose")
hf_fit2 <- hf_fit %>% filter(Disaccharide =="Sucrose") %>% filter(p.adj <= 0.05)
p2 <- df %>% filter(Disaccharide == "Sucrose") %>%
    ggplot(aes(x = x, y = y)) +
    geom_bar(stat = "identity", colour = "black", position = "dodge", 
             scales = "free_y", fill = "green4") +
    geom_errorbar(aes(ymin = y-CI, ymax = y+CI), width=0.5, size=1, 
                  position = position_dodge(width = 0.90)) +
    geom_point(data = dfpoint2, aes(x = x, y = y, fill = Disaccharide), fill = "green4", shape = 21, 
               size = 1, stroke = 2, position = position_dodge(width = 0.90)) +
    ylab("Water content (%)") + 
    theme_bw() +
    theme(plot.title = element_text(hjust = 0.5), strip.text.x = element_blank(), 
          legend.position = "none", axis.title.x=element_blank()) +
    stat_pvalue_manual(data = hf_fit2, y.position = 8, step.increase = 0.4, 
                       label = "p.adj.signif", size = 4) 

dfpoint3 <- dfpoint %>% filter(Disaccharide == "Trehalose")
hf_fit3 <- hf_fit %>% filter(Disaccharide =="Trehalose") %>% filter(p.adj <= 0.05)
p3 <- df %>% filter(Disaccharide == "Trehalose") %>%
    ggplot(aes(x = x, y = y)) +
    geom_bar(stat = "identity", colour = "black", position = "dodge", 
             scales = "free_y", fill = "royalblue1") +
    geom_errorbar(aes(ymin = y-CI, ymax = y+CI), width=0.5, size=1, 
                  position = position_dodge(width = 0.90)) +
    geom_point(data = dfpoint3, aes(x = x, y = y, fill = Disaccharide), fill = "royalblue1", shape = 21, 
               size = 1, stroke = 2, position = position_dodge(width = 0.90)) +
    xlab("Disaccharide content (%)") +     
    theme_bw() +
    theme(plot.title = element_text(hjust = 0.5), strip.text.x = element_blank(), 
          legend.position = "none", axis.title.y=element_blank()) +
    stat_pvalue_manual(data = hf_fit3, y.position = 12, step.increase = 0.2, 
                       label = "p.adj.signif", size = 4) 

p4 <- p1/p2/p3
p4
ggsave(p4, file = "Revised_Treh_glyc_PerwaterVsGlycerol_bar.jpeg")
ggsave(p4, file = "Revised_Treh_glyc_PerwaterVsGlycerol_bar.pdf", width = 8, height = 10, units = "in")
```
