## Supplementary material for "Water content, transition temperature and fragility influence protection and anhydrobiotic capacity": File S6: 5b_OrgVsTgoff_bar.R

```
setwd("c:/R/Glass")

library(ggplot2) #load in libraries
library(gtools)
library(ggrepel)
library(readxl)
library(ggsignif)
library(ggpubr)
library(ggprism)
library(polynom)
library(hrbrthemes)
library(tidyverse)
library(rstatix)
library(nplyr)
library(patchwork)
source("rsquared.R")
source("ANOVA_Tukey.R")

AS <- read_excel("org_as.xlsx", sheet = "Sheet2") 
#reading excel sheet into var Treh

#AT <- read_excel("org_at.xlsx", sheet = "Sheet1")
#reading excel sheet into var Sucr

#CE <- read_excel("org_ce.xlsx", sheet = "Sheet1")
CE <- read_excel("org_ce.xlsx", sheet = "Sheet2")
#reading excel sheet into var Sucr

SC <- read_excel("org_sc.xlsx", sheet = "Sheet2")
#reading excel sheet into var Sucr

df1 <- data.frame(y = AS$Tg_offset, Organism = AS$Sample, CI = AS$offset_CI, 
                  Disaccharide = AS$life_stage, Transition = AS$Transition)
#read x and y variables into dataframe df1

#df2 <- data.frame(y = AT$Tg_offset, Organism = AT$Sample, CI = AT$offset_CI, 
#                  Disaccharide = AT$life_stage)
#read x and y variables into dataframe df1

df3 <- data.frame(y = CE$Tg_offset, Organism = CE$Sample, CI = CE$offset_CI, 
                  Disaccharide = CE$life_stage, Transition = CE$Transition)
#read x and y variables into dataframe df1

df4 <- data.frame(y = SC$Tg_offset, Organism = SC$Sample, CI = SC$offset_CI, 
                  Disaccharide = SC$life_stage, Transition = SC$Transition)
#read x and y variables into dataframe df1

df <- rbind(df1, df4, df3)

blank_data_top <- data.frame(Organism = c("Artemia sp.", "Sacchromyces cerevisiae"), 
                         Disaccharide = c("adult", "log"), 
                         y = c(120, 85))
blank_data_bot <- data.frame(Organism = c("Caenorhabditis elegans"),
                             Transition = c("1", "2", "3"),
                             Disaccharide = c("dauer preconditioned"), 
                             y = c(75, 125, 190))

df$Disaccharide <- factor(df$Disaccharide, levels = c("adult", "cyst", "non-dauer", 
        "dauer", "dauer preconditioned", "log", "stationary"))

df$Transition <- factor(df$Transition, levels = c("1", "2", "3"))

dftop <- df %>% filter_all(all_vars(.!="Caenorhabditis elegans"))
dfbot <- filter(df, Organism == c("Caenorhabditis elegans"))

p1 <- 
  ggplot(dftop, aes(x = Disaccharide, y = y, fill = Organism)) +
  geom_bar(stat = "identity", colour = "black", position = "dodge", scales = "free_y") +
  geom_errorbar(aes(ymin = y-CI, ymax = y+CI), position = position_dodge(1), 
                width=0.5, size=1) +
  geom_blank(data = blank_data_top, aes(x = Disaccharide, y = y)) +
  #create continuous xlabels for all mixtures
  #scale_x_continuous(breaks = c(0, 2.5, 5, 7.5, 10, 12.5), labels = c("100.0%", 
  #  "97.5%", "95.0%", "92.5%", "90.0%", "87.5%")) +
  #xlab("Desiccation tolerant organism") +   
  ylab("Glass transition offset (°C)") + 
  #coord_cartesian(xlim = c(-1, 13.5), ylim = c(0, 100)) +
  ggtitle("Glass transition offset (°C) for Desiccation tolerant \norganinms' non-tolerant and tolerant life stages") +
  facet_wrap(vars(Organism), scales='free') +
  theme_bw() +
  theme(legend.position = "none", axis.title.x = element_blank(), 
        plot.title = element_text(hjust = 0.5), strip.text.x = element_blank(), 
        axis.text.x = element_text(angle = 90, hjust = 1, vjust = 0.5))

p1 + scale_x_discrete(labels = function(x) str_wrap(x, width = 10))

p2 <- 
    ggplot(dfbot, aes(x = Disaccharide, y = y)) +
    geom_bar(stat = "identity", colour = "black", position = "dodge", 
             scales = "free_y", fill = "green") +
    geom_errorbar(aes(ymin = y-CI, ymax = y+CI), position = position_dodge(1), 
                  width=0.5, size=1) +
    geom_blank(data = blank_data_bot, aes(x = Disaccharide, y = y)) +
    #create continuous xlabels for all mixtures
    #scale_x_continuous(breaks = c(0, 2.5, 5, 7.5, 10, 12.5), labels = c("100.0%", 
    #  "97.5%", "95.0%", "92.5%", "90.0%", "87.5%")) +
    xlab("Desiccation tolerant organism") +   
    ylab("Glass transition offset (°C)") + 
    #coord_cartesian(xlim = c(-1, 13.5), ylim = c(0, 100)) +
    #ggtitle("Glass transition offset (°C) for Desiccation tolerant \norganinms' 
    #      non-tolerant and tolerant life stages") +
    facet_wrap(vars(Transition), scales='free') +
    theme_bw() +
    theme(legend.position = "none", axis.title.x = element_blank(), 
          plot.title = element_text(hjust = 0.5), strip.text.x = element_blank(), 
          axis.text.x = element_text(angle = 90, hjust = 1, vjust = 0.5)) 

p2 + scale_x_discrete(labels = function(x) str_wrap(x, width = 10))

p1 / p2

ggsave(file = "Treh_glyc_OrgVsTgoff_bar_3.jpeg")
ggsave("Treh_glyc_OrgVsTgoff_bar_3.pdf", width = 8, height = 6, units = "in")
```
