## Supplementary material for "Water content, transition temperature and fragility influence protection and anhydrobiotic capacity": File S6: stat_cor.R

```
#' @include utilities.R utilities_label.R
NULL
#'Add Correlation Coefficients with P-values to a Scatter Plot
#'@description Add correlation coefficients with p-values to a scatter plot. Can
#'  be also used to add `R2`.
#'@inheritParams ggplot2::layer
#'@param method a character string indicating which correlation coefficient (or
#'  covariance) is to be computed. One of "pearson" (default), "kendall", or
#'  "spearman".
#'@param alternative a character string specifying the alternative hypothesis,
#'  must be one of "two.sided" (default), "greater" or "less". You can specify
#'  just the initial letter.
#'@param cor.coef.name character. Can be one of \code{"R"} (pearson coef),
#'  \code{"rho"} (spearman coef) and \code{"tau"} (kendall coef). Uppercase and
#'  lowercase are allowed.
#'@param label.sep a character string to separate the terms. Default is ", ", to
#'  separate the correlation coefficient and the p.value.
#'@param label.x.npc,label.y.npc can be \code{numeric} or \code{character}
#'  vector of the same length as the number of groups and/or panels. If too
#'  short they will be recycled. \itemize{ \item If \code{numeric}, value should
#'  be between 0 and 1. Coordinates to be used for positioning the label,
#'  expressed in "normalized parent coordinates". \item If \code{character},
#'  allowed values include: i) one of c('right', 'left', 'center', 'centre',
#'  'middle') for x-axis; ii) and one of c( 'bottom', 'top', 'center', 'centre',
#'  'middle') for y-axis.}
#'
#'  If too short they will be recycled.
#'@param label.x,label.y \code{numeric} Coordinates (in data units) to be used
#'  for absolute positioning of the label. If too short they will be recycled.
#'@param output.type character One of "expression", "latex", "tex" or "text".
#'@param digits,r.digits,p.digits integer indicating the number of decimal
#'  places (round) or significant digits (signif) to be used for the correlation
#'  coefficient and the p-value, respectively..
#'@param r.accuracy a real value specifying the number of decimal places of
#'  precision for the correlation coefficient. Default is NULL. Use (e.g.) 0.01
#'  to show 2 decimal places of precision. If specified, then \code{r.digits} is
#'  ignored.
#'@param p.accuracy a real value specifying the number of decimal places of
#'  precision for the p-value. Default is NULL. Use (e.g.) 0.0001 to show 4
#'  decimal places of precision. If specified, then \code{p.digits} is ignored.
#'@param ... other arguments to pass to \code{\link[ggplot2]{geom_text}} or
#'  \code{\link[ggplot2:geom_text]{geom_label}}.
#'@param na.rm If FALSE (the default), removes missing values with a warning. If
#'  TRUE silently removes missing values.
#'@seealso \code{\link{ggscatter}}
#'@section Computed variables: \describe{ \item{r}{correlation coefficient}
#'  \item{rr}{correlation coefficient squared} \item{r.label}{formatted label
#'  for the correlation coefficient} \item{rr.label}{formatted label for the
#'  squared correlation coefficient} \item{p.label}{label for the p-value}
#'  \item{label}{default labeldisplayed by \code{stat_cor()}} }
#'
#' @examples
#' # Load data
#' data("mtcars")
#' df <- mtcars
#' df$cyl <- as.factor(df$cyl)
#'
#' # Scatter plot with correlation coefficient
#' #:::::::::::::::::::::::::::::::::::::::::::::::::
#' sp <- ggscatter(df, x = "wt", y = "mpg",
#'    add = "reg.line",  # Add regressin line
#'    add.params = list(color = "blue", fill = "lightgray"), # Customize reg. line
#'    conf.int = TRUE # Add confidence interval
#'    )
#' # Add correlation coefficient
#' sp + stat_cor(method = "pearson", label.x = 3, label.y = 30)
#'
#' # Specify the number of decimal places of precision for p and r
#' # Using 3 decimal places for the p-value and
#' # 2 decimal places for the correlation coefficient (r)
#' sp + stat_cor(p.accuracy = 0.001, r.accuracy = 0.01)
#'
#' # Show only the r.label but not the p.label
#' sp + stat_cor(aes(label = ..r.label..), label.x = 3)
#'
#'# Use R2 instead of R
#'ggscatter(df, x = "wt", y = "mpg", add = "reg.line") +
#'  stat_cor(
#'    aes(label = paste(..rr.label.., ..p.label.., sep = "~`,`~")),
#'   label.x = 3
#' )
#'
#' # Color by groups and facet
#' #::::::::::::::::::::::::::::::::::::::::::::::::::::
#' sp <- ggscatter(df, x = "wt", y = "mpg",
#'    color = "cyl", palette = "jco",
#'    add = "reg.line", conf.int = TRUE)
#' sp + stat_cor(aes(color = cyl), label.x = 3)
#'
#'@export
stat_cor <- function(mapping = NULL, data = NULL,
                     method = "pearson", alternative = "two.sided",
                     cor.coef.name = c("R", "rho", "tau"), label.sep = ", ",
                     label.x.npc = "left", label.y.npc = "top",
                     label.x = NULL, label.y = NULL, output.type = "expression",
                     digits = 2, r.digits = digits, p.digits = digits,
                     r.accuracy = NULL, p.accuracy = NULL,
                     geom = "text", position = "identity",  na.rm = FALSE, show.legend = NA,
                    inherit.aes = TRUE, ...) {
  parse <- ifelse(output.type == "expression", TRUE, FALSE)
  cor.coef.name = cor.coef.name[1]
  layer(
    stat = StatCor, data = data, mapping = mapping, geom = geom,
    position = position, show.legend = show.legend, inherit.aes = inherit.aes,
    params = list(label.x.npc  = label.x.npc , label.y.npc  = label.y.npc,
                  label.x = label.x, label.y = label.y, label.sep = label.sep,
                  method = method, alternative = alternative, output.type = output.type, digits = digits,
                  r.digits = r.digits, p.digits = p.digits, r.accuracy = r.accuracy,
                  p.accuracy = p.accuracy, cor.coef.name = cor.coef.name,
                  parse = parse, na.rm = na.rm, ...)
  )
}


StatCor<- ggproto("StatCor", Stat,
                  required_aes = c("x", "y"),
                  default_aes = aes(hjust = ..hjust.., vjust = ..vjust..),

                  compute_group = function(data, scales, method, alternative, label.x.npc, label.y.npc,
                                           label.x, label.y, label.sep, output.type, digits,
                                           r.digits, p.digits, r.accuracy, p.accuracy, cor.coef.name)
                    {
                    if (length(unique(data$x)) < 2) {
                      # Not enough data to perform test
                      return(data.frame())
                    }
                    # Returns a data frame with estimate, p.value, label, method
                    .test <- .cor_test(
                      data$x, data$y, method = method, alternative = alternative,
                      label.sep = label.sep, output.type = output.type, digits = digits,
                      r.digits = r.digits, p.digits = p.digits,
                      r.accuracy = r.accuracy, p.accuracy = p.accuracy,
                      cor.coef.name = cor.coef.name
                      )
                    # Returns a data frame with label: x, y, hjust, vjust
                    .label.pms <- .label_params(data = data, scales = scales,
                                                label.x.npc = label.x.npc, label.y.npc = label.y.npc,
                                                label.x = label.x, label.y = label.y ) %>%
                      mutate(hjust = 0)
                    cbind(.test, .label.pms)
                  }
)


# Correlation test
#::::::::::::::::::::::::::::::::::::::::
# Returns a data frame: estimatel|p.value|method|label
.cor_test <- function(x, y, method = "pearson", alternative = "two.sided",
                      label.sep = ", ", output.type = "expression",
                      digits = 2, r.digits = digits, p.digits = digits,
                      r.accuracy = NULL, p.accuracy = NULL,
                      cor.coef.name = "R"){
  # Overwritting digits by accuracy, if specified
  if(!is.null(p.accuracy)){
    nb_decimal_places <- round(abs(log10(p.accuracy)))
    p.digits <- nb_decimal_places
  }
  if(!is.null(r.accuracy)){
    nb_decimal_places <- round(abs(log10(r.accuracy)))
    r.digits <- nb_decimal_places
  }

  # Correlation analyses
  .cor <- suppressWarnings(stats::cor.test(
    x, y, method = method,  alternative = alternative,
    use = "complete.obs"
    ))
  estimate <- p.value <- p <- r <- rr <-  NULL
  z <- data.frame(estimate = .cor$estimate, p.value = .cor$p.value, method = method) %>%
    mutate(
      r = signif(estimate, r.digits),
      rr = signif(estimate^2, r.digits),
      p = signif(p.value, p.digits)
    )

  # Defining p and r labels
  pval <- .cor$p.value
  z <- z %>%
    dplyr::mutate(
      r.label = get_corcoef_label(
        r, accuracy = r.accuracy, prefix = "R",
        cor.coef.name = cor.coef.name, type = output.type
        ),
      rr.label = get_corcoef_label(
        rr, accuracy = r.accuracy, prefix = "R2",
        cor.coef.name = cor.coef.name, type = output.type
        ),
      p.label = get_p_label(
        p, accuracy = p.accuracy, type = output.type
        )
  )

  # Defining correlation labels
  if(output.type == "expression"){
    if(label.sep == "\n"){
      # Line break at each comma
      cortxt <- paste0("atop(", z$r.label, ",",  z$p.label, ")")
    }
    else{
      label.sep <- trimws(label.sep)
      if(label.sep == "") label.sep <- "~"
      #  Using "*" to avoid the space between the R2 value and comma
      else label.sep <- paste0("*`", label.sep, "`~")
      cortxt <- paste0(z$r.label, label.sep,  z$p.label)
    }
  }
  else if (output.type %in% c("latex", "tex", "text")){
    cortxt <- paste0(z$r.label, label.sep,  z$p.label)
  }
  z$label <- cortxt
  z
}


# Formatting R and P ----------------------
get_p_label <- function(x, accuracy = 0.0001, type = "expression"){
  if(is.null(accuracy)){
    label <- ifelse(x < 2.2e-16, "p < 2.2e-16", paste0("p = ", x))
  }
  else if (!(accuracy < 1)){
    stop(
      "Accuracy should be < 1; For example use 0.01, 0.001, 0.0001, etc.",
      call. = FALSE
    )
  }
  else{
    label <- scales::pvalue(x, accuracy = accuracy, add_p = TRUE)
    # Add space before and after: = or <
    label <- gsub(pattern = "(=|<)", replacement = " \\1 ", x = label)
  }
  if(type == "expression"){
    label <- gsub(pattern = "p = ", replacement = "italic(p)~`=`~", x = label, fixed = TRUE)
    label <- gsub(pattern = "p < ", replacement = "italic(p)~`<`~", x = label, fixed = TRUE)
  }
  label
}

# Prefix can be R or R^2.
# cor.coef.name: R, rho, tau
get_corcoef_label <- function(x, accuracy = 0.01, prefix = "R", cor.coef.name = "R", type = "expression"){
  if(is.null(accuracy)){
    label <- paste0(prefix, " = ", x)
  }
  else if(!(accuracy < 1)){
    stop(
      "Accuracy should be < 1; For example use 0.01, 0.001, 0.0001, etc.",
      call. = FALSE
    )
  }
  else{
    nb_decimal_places <- round(abs(log10(accuracy)))
    label <- formatC(x, digits = nb_decimal_places, format = "f")
    label <- paste0(prefix, " = ", label)
  }
  if(type == "expression"){
    label <- gsub(pattern = "R2 = ", replacement = "italic(R)^2~`=`~", x = label, fixed = TRUE)
    label <- gsub(pattern = "R = ", replacement = "italic(R)~`=`~", x = label, fixed = TRUE)
  }
  label <- gsub(pattern = "R", cor.coef.name, x = label, fixed = TRUE)
  label
}
```
