## Supplementary material for "Water content, transition temperature and fragility influence protection and anhydrobiotic capacity": File S6: utilities.R

```
#' @include desc_statby.R utilities_base.R utilities_color.R
NULL
#' @import ggplot2
#' @importFrom magrittr %>%
#' @importFrom rstatix df_group_by df_nest_by df_select df_arrange
#' @importFrom tibble as_tibble
#' @importFrom dplyr group_by mutate mutate_if group_nest arrange desc
#' @importFrom purrr map2 map
#' @importFrom tidyr unite
#' @importFrom dplyr do select distinct
#' @importFrom dplyr summarise
#' @importFrom dplyr everything
#' @importFrom grid drawDetails
#' @importFrom rlang !!
#' @importFrom rlang !!!
#' @importFrom rlang syms .data


required_package <- function(pkg){
  if (!requireNamespace(pkg, quietly = TRUE)) {
    stop(
      pkg, " package needed to be installed before using this function. ",
      "Type this in R: install.packages('", pkg, "')"
    )
  }
}

# Unnesting, adapt to tidyr 1.0.0
unnest <- function(data, cols = "data", ...){
  if(is_pkg_version_sup("tidyr", "0.8.3")){
    results <- tidyr::unnest(data, cols = cols, ...)
  }
  else {results <- tidyr::unnest(data, ...)}
  results
}

# Check if an installed package version is superior to a specified version
# Version, pkg: character vector
is_pkg_version_sup<- function(pkg, version){
  vv <- as.character(utils::packageVersion(pkg))
  cc <- utils::compareVersion(vv, version) > 0
  cc
}

keep_only_tbl_df_classes <- function(x){
  toremove <- setdiff(class(x), c("tbl_df", "tbl", "data.frame"))
  if(length(toremove) > 0){
    class(x) <- setdiff(class(x), toremove)
  }
  x
}

#%%%%%%%%%%%%%%%%%%%%%%%%%%%%%%%%%%%%%
# Execute a geom_* function from ggplot2
#%%%%%%%%%%%%%%%%%%%%%%%%%%%%%%%%%%%%%
# geomfunc : gem_*() functions
# data data for mapping
# ... argument accepeted by the function
# return a plot if geomfunc!=Null or a list(option, mapping) if  geomfunc = NULL
.geom_exec <- function (geomfunc = NULL, data = NULL,
                        position = NULL, ...) {
  geom_exec(geomfunc = geomfunc, data = data, position = position, ...)
}

# %%%%%%%%%%%%%%%%%%%%%%%%%%%%%%%%%%%%%%%%%%%%%%%%%%%
# Official argument from ggplot2
# %%%%%%%%%%%%%%%%%%%%%%%%%%%%%%%%%%%%%%%%%%%%%%%%%
# bar plot arguments
.barplot_params <- function(...){
  x <- list(...)
  res <- list()
  res$width  <- x$width
  res$binwidth  <- x$binwidth
  res$na.rm  <- ifelse(!is.null(x$na.rm), x$na.rm, FALSE)
  res$show.legend <- ifelse(!is.null(x$show.legend), x$show.legend, NA)
  res$inherit.aes <- ifelse(!is.null(x$inherit.aes), x$inherit.aes, TRUE)
  return(res)
}

# box plot arguments
.boxplot_params <- function(...){
  x <- list(...)
  res <- list()
  res$outlier.colour  <- x$outlier.colour
  res$outlier.shape  <- ifelse(!is.null(x$outlier.shape), x$outlier.shape, 19)
  res$outlier.size  <- ifelse(!is.null(x$outlier.size), x$outlier.size, 1.5)
  res$outlier.stroke  <- ifelse(!is.null(x$outlier.stroke), x$outlier.stroke, 0.5)
  res$notch  <- ifelse(!is.null(x$notch), x$notch, FALSE)
  res$notchwidth  <- ifelse(!is.null(x$notchwidth), x$notchwidth, 0.5)
  res$varwidth  <- ifelse(!is.null(x$varwidth), x$varwidth, FALSE)
  res$na.rm  <- ifelse(!is.null(x$na.rm), x$na.rm, FALSE)
  res$show.legend <- ifelse(!is.null(x$show.legend), x$show.legend, NA)
  res$inherit.aes <- ifelse(!is.null(x$inherit.aes), x$inherit.aes, TRUE)
  return(res)
}

.dotplot_params <- function(...){
  x <- list(...)
  res <- list()
  res$stackratio  <- ifelse(!is.null(x$stackratio ), x$stackratio, 1)
  res$width <- ifelse(!is.null(x$width), x$width, 0.9)
  return(res)
}

.violin_params <- function(...){
  x <- list(...)
  res <- list()
  res$stat  <- ifelse(!is.null(x$stat ), x$stat, "ydensity")
  res$draw_quantiles  <- x$draw_quantiles
  res$scale <- ifelse(!is.null(x$scale), x$scale, "area")
  res$trim <- ifelse(!is.null(x$trim), x$trim, TRUE)
  return(res)
}

.hist_params <- function(...){
  x <- list(...)
  res <- list()
  res$binwidth <- x$binwidth
  res$bins <- x$bins
  return(res)
}

.standard_params <- function(...){
  x <- list(...)
  res <- list()
  res$color <- ifelse(!is.null(x$color), x$color, "black")
  res$color <- ifelse(!is.null(x$colour), x$colour, res$color)
  res$linetype <- ifelse(!is.null(x$linetype), x$linetype, "solid")
  res$size <- ifelse(!is.null(x$size), x$size, 1)
  res$fill <- ifelse(!is.null(x$fill), x$fill, "black")
  res$shape <- ifelse(!is.null(x$shape), x$shape, 19)
  res
}


#%%%%%%%%%%%%%%%%%%%%%%%%%%%%%%%%%
# Graphical parameters
#%%%%%%%%%%%%%%%%%%%%%%%%%%%%%%%%%

# Set plot orientation
# %%%%%%%%%%%%%%%%%%%%%%%%%%%%%
.set_orientation <-
  function(p, orientation = c("vertical", "horizontal", "reverse")) {
    ori <- match.arg(orientation)
    if (ori == "horizontal") p + coord_flip()
    else if (ori == "reverse")
      p + scale_y_reverse()
    else p
  }


# Change title and labels
# %%%%%%%%%%%%%%%%%%%%%%%%%%%%%
.labs <- function(p, main = NULL, xlab = NULL, ylab = NULL,
                  font.main = NULL, font.x = NULL, font.y = NULL,
                  submain = NULL, caption = NULL,
                  font.submain = NULL, font.caption = NULL)
{

  font.main <- .parse_font(font.main)
  font.x <- .parse_font(font.x)
  font.y <- .parse_font(font.y)
  font.submain <- .parse_font(font.submain)
  font.caption <- .parse_font(font.caption)

  if(is.logical(main)){
    if(!main) main <- NULL
  }

  if(is.logical(submain)){
    if(!submain) submain <- NULL
  }

  if(is.logical(caption)){
    if(!caption) caption <- NULL
  }


  if (!is.null(main)) {
      p <- p + labs(title = main)
  }

  if (!is.null(submain)) {
      p <- p + labs(subtitle = submain)
  }

  if (!is.null(caption)) {
      p <- p + labs(caption = caption)
  }

  if (!is.null(xlab)) {
    if (xlab == FALSE)
      p <- p + theme(axis.title.x = element_blank())
    else
      p <- p + labs(x = xlab)
  }

  if (!is.null(ylab)) {
    if (ylab == FALSE)
      p <- p + theme(axis.title.y = element_blank())
    else
      p <- p + labs(y = ylab)
  }

  if (!is.null(font.main))
    p <-
    p + theme(
      plot.title = element_text(
        size = font.main$size,
        lineheight = 1.0, face = font.main$face, colour = font.main$color
      )
    )
  if (!is.null(font.submain))
    p <-
    p + theme(
      plot.subtitle = element_text(
        size = font.submain$size,
        lineheight = 1.0, face = font.submain$face, colour = font.submain$color
      )
    )
  if (!is.null(font.caption))
    p <-
    p + theme(
      plot.caption = element_text(
        size = font.caption$size,
        lineheight = 1.0, face = font.caption$face, colour = font.caption$color
      )
    )
  if (!is.null(font.x))
    p <-
    p + theme(axis.title.x = element_text(
      size = font.x$size,
      face = font.x$face, colour = font.x$color
    ))
  if (!is.null(font.y))
    p <-
    p + theme(axis.title.y = element_text(
      size = font.y$size,
      face = font.y$face, colour = font.y$color
    ))


  p
}


# ticks
# %%%%%%%%%%%%%%%%%%%%%%%%%%%%%
.set_ticks <-
  function(ticks = TRUE, tickslab = TRUE, font.tickslab = NULL,
           xtickslab.rt = NULL, ytickslab.rt = NULL,
           font.xtickslab = font.tickslab, font.ytickslab = font.tickslab)
  {

    . <- xhjust <- xvjust <- NULL
   if(!is.null(xtickslab.rt)) {
     if(xtickslab.rt > 5) xhjust <- 1
     if(xtickslab.rt == 90) xvjust <- 0.5
     }
    else xhjust <- NULL

    if (ticks)
      ticks <-
        element_line(colour = "black")
    else
      ticks <- element_blank()

    if (is.null(font.xtickslab)) font.x <- list()
    else font.x <- .parse_font(font.xtickslab)
    if (is.null(font.ytickslab)) font.y <- list()
    else font.y <- .parse_font(font.ytickslab)

    if (tickslab) {
      xtickslab <- font.x %>% .add_item(hjust = xhjust, vjust = xvjust, angle = xtickslab.rt) %>%
        do.call(element_text, .)
      ytickslab <- font.y %>% .add_item(angle = ytickslab.rt) %>% do.call(element_text, .)
    }
    else {
      xtickslab <- element_blank()
      ytickslab <- element_blank()
    }
    theme(
      axis.ticks = ticks, axis.text.x = xtickslab, axis.text.y = ytickslab
    )
  }


# Change Axis limits
# %%%%%%%%%%%%%%%%%%%%%%%%%%%%%
.set_axis_limits <- function(xlim = NULL, ylim = NULL){
  if(!is.null(xlim) | !is.null(ylim)) coord_cartesian(xlim, ylim)
}


# Axis scales
# %%%%%%%%%%%%%%%%%%%%%%%%%%%%%
.set_scale <- function (p, xscale = c("none", "log2", "log10", "sqrt"),
                        yscale = c("none", "log2", "log10", "sqrt"),
                        format.scale = FALSE)
{

  xscale <- match.arg(xscale)
  yscale <- match.arg(yscale)
  .x <- ".x"

  if(format.scale){
    if(!requireNamespace("scales")) stop("The R package 'scales' is required.")

      if(yscale == "log2"){
        p <- p + scale_y_continuous(trans = scales::log2_trans(),
                           breaks = scales::trans_breaks("log2", function(x) 2^x),
                           labels = scales::trans_format("log2", scales::math_format(2^.x)))
      }
    else if(yscale == "log10"){
      p <- p + scale_y_continuous(trans = scales::log10_trans(),
                                  breaks = scales::trans_breaks("log10", function(x) 10^x),
                                  labels = scales::trans_format("log10", scales::math_format(10^.x)))
    }

    if(xscale == "log2"){
      p <- p + scale_x_continuous(trans = scales::log2_trans(),
                                  breaks = scales::trans_breaks("log2", function(x) 2^x),
                                  labels = scales::trans_format("log2", scales::math_format(2^.x)))
    }
    else if(xscale == "log10"){
      p <- p + scale_x_continuous(trans = scales::log10_trans(),
                                  breaks = scales::trans_breaks("log10", function(x) 10^x),
                                  labels = scales::trans_format("log10", scales::math_format(10^.x)))
    }

  }

  else{
    if(xscale != "none")  p <- p + scale_x_continuous(trans = xscale)
    if(yscale != "none") p <- p + scale_y_continuous(trans = yscale)
  }
p
}

# Legends
# %%%%%%%%%%%%%%%%%%%%%%%%%%%%%
.set_legend <- function(p, legend = NULL,
                        legend.title = NULL, font.legend = NULL)
{
  if(is.null(legend.title)) legend.title = waiver()
  font <- .parse_font(font.legend)

  if(!is.null(legend)) p <- p + theme(legend.position = legend)

  if(!.is_empty(legend.title)){

    if(.is_list(legend.title)) p <- p + do.call(ggplot2::labs, legend.title)
    else p <- p +
       labs(color = legend.title, fill = legend.title, linetype = legend.title, shape = legend.title)
  }

   if(!is.null(font)){
     p <- p + theme(
       legend.text = element_text(size = font$size,
                                  face = font$face, colour = font$color),
       legend.title = element_text(size = font$size,
                                   face = font$face, colour = font$color)
     )
   }

   p
}


# Set ticks by
# %%%%%%%%%%%%%%%%%%%%%%%%%%%%%
.set_ticksby <- function(p, xticks.by = NULL, yticks.by = NULL)
{
  if(!is.null(yticks.by)) {
    # Forcing ymin to start at 0 when distribution plot
    gg_mapping <- .get_gg_xy_variables(p)
    is_density_plot <-  gg_mapping["y"] %in% c("..count..", "..density..", "..ecdf..")
    ymin <- NULL
    if(is_density_plot) ymin <- 0
    p <- p + scale_y_continuous(breaks = get_breaks(by = yticks.by, from = ymin))
  }
  else if(!is.null(xticks.by)) {
    p <- p + scale_x_continuous(breaks = get_breaks(by = xticks.by))
  }
  p
}


# %%%%%%%%%%%%%%%%%%%%%%%%%%%%%
# Add stat
# %%%%%%%%%%%%%%%%%%%%%%%%%%%%%

.check_add.params <- function(add, add.params, error.plot, data, color, fill,  ...){
  if(color %in% names(data) & is.null(add.params$color))  add.params$color <- color
  if(fill %in% names(data) & is.null(add.params$fill))  add.params$fill <- fill
  if(is.null(add.params$color)) add.params$color <- color
  if(is.null(add.params$fill) & ("crossbar" %in% error.plot | "boxplot" %in% add | "violin" %in% add)) add.params$fill <- fill
  if(is.null(add.params$fill)) add.params$fill <- add.params$color
  #else add.params$fill <- add.params$color
  if(!is.null(list(...)$shape) & is.null(add.params$shape)) add.params$shape <- list(...)$shape
  add.params
}


# Calculate the mean and the SD in each group
#+++++++++++++++++++++++++
# data : a data frame
# varname : the name of the variable to be summariezed
# grps : column names to be used as grouping variables
# .mean_sd <- function(data, varname, grps){
#   summary_func <- function(x, col){
#     c(mean = base::mean(x[[col]], na.rm=TRUE),
#       sd = stats::sd(x[[col]], na.rm=TRUE))
#   }
#   data_sum <- plyr::ddply(data, grps, .fun=summary_func, varname)
#   data_sum$ymin <- data_sum$mean-data_sum$sd
#   data_sum$ymax <- data_sum$mean+data_sum$sd
#   names(data_sum)[ncol(data_sum)-3] <- varname
#   # data_sum <- plyr::rename(data_sum, c("mean" = varname))
#   return(data_sum)
# }


# Summary functions
.summary_functions <- function(){
  c("mean", "mean_se", "mean_se_", "mean_sd", "mean_ci",
    "mean_range", "median", "median_iqr", "median_hilow", "median_hilow_",
    "median_q1q3", "median_mad", "median_range")
}
.errorbar_functions <- function(){
  setdiff(.summary_functions(), c("mean", "median"))
}


# parse font
# %%%%%%%%%%%%%%%%%%%%%%%%%%%%%
.parse_font <- function(font){
  if(is.null(font)) res <- NULL
  else if(inherits(font, "list")) res <- font
  else{
    # matching size and face
    size <- grep("^[0-9]+$", font, perl = TRUE)
    face <- grep("plain|bold|italic|bold.italic", font, perl = TRUE)
    if(length(size) == 0) size <- NULL else size <- as.numeric(font[size])
    if(length(face) == 0) face <- NULL else face <- font[face]
    color <- setdiff(font, c(size, face))
    if(length(color) == 0) color <- NULL
    res <- list(size=size, face = face, color = color)
  }
  res
}


# Add annotation to a plot
# label: text to be added to a plot
# size: text size
# coord: x and coordinates
.ggannotate <- function (label, size = 12, coord = c(NULL, NULL)){
  if(is.null(unique(coord))){
    grob <- grid::grobTree(grid::textGrob(label, x = 0.3,  y = 0.80, hjust=0,
                                    gp = grid::gpar(col = "black", fontsize = size, fontface = "plain")))
    ggplot2::annotation_custom(grob)
  }
  else{
    ggplot2::annotate("text", x = coord[1], y = coord[2],
                      label = label, size = size/3)
  }
}

#:::::::::::::::::::::::::::::::::::::::::
# Check the data provided by user
#:::::::::::::::::::::::::::::::::::::::::
# combine: if TRUE, gather y variables
# return a list(data, x, y)
.check_data <- function(data, x, y, combine = FALSE)
  {

  if(missing(x) & missing(y)){
    if(!is.numeric(data))
      stop("x and y are missing. In this case data should be a numeric vector.")
    else{
      data <- data.frame(y = data, x = rep(1, length(data)))
      x <- "x"
      y <- "y"
    }
  }
  else if(missing(x)) {
    x <- "x"
    if(is.numeric(data)) data <- data.frame(x = data)
    else data$x <- rep("1", nrow(data))
  }
  # A list of y elements to plot
  else if(length(y) > 1){
    if(!all(y %in% colnames(data))){
      not_found <- setdiff(y , colnames(data))
      y <- intersect(y, colnames(data))

      if(.is_empty(y))
        stop("Can't find the y elements in the data.")

      else if(!.is_empty(not_found))
        warning("Can't find the following element in the data: ",
              .collapse(not_found))
    }
  }

  if(inherits(data, c("tbl_df", "tbl")))
    data <- as.data.frame(data)

  # Combining y variables
  #......................................................
  if(is.null(y)) y <- ""
  if(combine & length(y) > 1){
    data <- data %>%
      df_gather(cols = y, names_to = ".y.", values_to = ".value.") %>%
      dplyr::mutate(.y. = factor(.data$.y., levels = unique(.data$.y.)))
    y <- ".value."
  }
  # Combining x variables: Case of density plot or histograms
  #......................................................
  else if(combine & length(x) > 1 & y[1] %in% c("..density..", "..count..", "..ecdf..", "..qq..")){
    data <- data %>%
      df_gather(cols = x, names_to = ".y.", values_to = ".value.") %>%
      dplyr::mutate(.y. = factor(.data$.y., levels = unique(.data$.y.)))
    x <- ".value."
  }

  # If not factor, x elements on the plot should
  # appear in the same order as in the data
  if(is.character(data[[x]]))
    data[[x]] <- factor(data[[x]], levels = unique(data[[x]]))

  y <- unique(y)
  names(y) <- y
  x <- unique(x)
  names(x) <- x

  if(y[1] %in% c("..density..", "..count..", "..ecdf..", "..qq.."))
    list(x = x, data = data, y = y)    # The name of plots are x variables
  else
    list(y = y, data = data, x = x)   # The name of plots will be y variables
}


# Adjust shape when ngroups > 6, to avoid ggplot warnings
.scale_point_shape <- function(p, data, shape){
  if(shape %in% colnames(data)){
    grp <- data[[shape]]
    if(!inherits(grp, "factor")) grp <- as.factor(grp)
    ngroups <- length(levels(data[[shape]]))
    if(ngroups > 6) p <- p + scale_shape_manual(values=1:ngroups, labels = levels(data[[shape]]))
  }
  p
}

# Get not numeric columns in a data.frame
.get_not_numeric_vars <- function(data_frame){
  is_numeric <- sapply(data_frame, is.numeric)
  if(sum(!is_numeric) == 0) res = NULL
  else res <- colnames(data_frame[, !is_numeric, drop = FALSE])
  res
}


# Get the current color used in ggplot
.get_ggplot_ncolors <- function(p){
  g <- ggplot_build(p)
  gdata <- g$data[[1]]
  cols <- fills <- 1
  if("colour" %in% names(gdata)) cols <- unique(unlist(gdata["colour"]))
  if("fills" %in% names(gdata)) fills <- unique(unlist(gdata["fill"]))
  max(length(cols), length(fills))
}

# Check if character string is a valid color representation
.is_color <- function(x) {
  sapply(x, function(X) {
    tryCatch(is.matrix(grDevices::col2rgb(X)),
             error = function(e) FALSE)
  })
}


# Collapse one or two vectors
#:::::::::::::::::::::::::::::::::::::::::::::::::::::::::::::::::::::
.collapse <- function(x, y = NULL, sep = "."){
  if(missing(y))
    paste(x, collapse = sep)
  else if(is.null(x) & is.null(y))
    return(NULL)
  else if(is.null(x))
    return (as.character(y))
  else if(is.null(y))
    return(as.character(x))
  else
    paste0(x, sep, y)
}

# Check if en object is empty
#:::::::::::::::::::::::::::::::::::::::::::::::::::::::::::::::::::::
.is_empty <- function(x){
  length(x) == 0
}

# Remove NULL items in a vector or list
#
# x a vector or list
.compact <- function(x){Filter(Negate(is.null), x)}

# Check if is a list
#:::::::::::::::::::::::::::::::::::::::::::::::::::::::::::::::::::::
.is_list <- function(x){
  inherits(x, "list")
}

# Returns the levels of a factor variable
#:::::::::::::::::::::::::::::::::::::::::::::::::::::::::::::::::::::
.levels <- function(x){
  if(!is.factor(x)) x <- as.factor(x)
  levels(x)
}

# Remove items from a list
#:::::::::::::::::::::::::::::::::::::::::::::::::::::::::::::::::::::
.remove_item <- function(.list, items){
  for(item in items)
    .list[[item]] <- NULL
  .list
}

# Additems in a list
#:::::::::::::::::::::::::::::::::::::::::::::::::::::::::::::::::::::
.add_item <- function(.list, ...){
  pms <- list(...)
  for(pms.names in names(pms)){
    .list[[pms.names]] <- pms[[pms.names]]
  }
  .list
}


# Select a colun as vector from tiblle data frame
.select_vec <- function(df, column){
  dplyr::pull(df, column)
}

# Select the top up or down rows of a data frame sorted by variables
#:::::::::::::::::::::::::::::::::::::::::::::::::::::::::::::::::::::
# - df: data frame
# - x: x axis variables (grouping variables)
# - y: y axis variables (sorting variables)
# - n the number of rows
# - grps: other grouping variables
.top_up <- function(df, x, y, n, grouping.vars = NULL){
  . <- NULL
  grouping.vars <- c(x, grouping.vars) %>%
    unique()
  df %>%
    df_arrange(vars = c(grouping.vars, y)) %>%
    df_group_by(vars = grouping.vars) %>%
    do(utils::tail(., n))
}


.top_down <- function(df, x, y, n, grouping.vars = NULL){
  . <- NULL
  grouping.vars <- c(x, grouping.vars) %>%
    unique()
  df %>%
    df_arrange(vars = c(grouping.vars, y)) %>%
    df_group_by(vars = grouping.vars) %>%
    do(utils::head(., n))
}


#::::::::::::::::::::::::::::::::::::::::::::::::::::::::::
# Apply ggpubr functions on a data
#::::::::::::::::::::::::::::::::::::::::::::::::::::::::::
# fun: function, can be ggboxplot, ggdotplot, ggstripchart, ...
.plotter <- function(fun, data, x, y, combine = FALSE, merge = FALSE,
                     color = "black", fill = "white",
                     title = NULL, xlab = NULL, ylab = NULL,
                     legend = NULL, legend.title = NULL,
                     facet.by = NULL,
                     select = NULL, remove = NULL, order = NULL,
                     add = "none", add.params = list(),
                     label = NULL, font.label = list(size = 11, color = "black"),
                     label.select = NULL, repel = FALSE, label.rectangle = FALSE,
                     font.family = "", parse = FALSE,
                     ggtheme = theme_pubr(),
                     fun_name = "", group = 1, # used only by ggline
                     show.legend.text = NA,
                     ...)
  {

  if(is.logical(merge)){
    if(merge) merge = "asis"
    else merge = "none"
  }
  if(combine & merge != "none")
    stop("You should use either combine = TRUE or merge = TRUE, but not both together.")

  font.label <- .parse_font(font.label)
  if(is.null(label) & fun_name == "barplot") label  <- FALSE
  .lab <- label
  if(fun_name != "barplot") .lab <- NULL

  if(!missing(x) & !missing(y)){
    if(length(y) == 1 & length(x) == 1){
      combine <- FALSE
      merge <- "none"
    }
  }

  # Check data
  #:::::::::::::::::::::::::::::::::::::::::::::::::::::::::
  # - returns a list of updated main options:
  #       list(y, data,  x)
  opts <- .check_data(data, x, y, combine = combine | merge != "none")
  data <- opts$data
  x <- opts$x
  y <- opts$y


  is_density_plot <- y[1] %in% c("..count..", "..density..", "..ecdf..", "..qq..")

  if(combine) facet.by <- ".y." # Faceting by y variables
  if(merge != "none"){
    if(!is_density_plot) facet.by <- NULL
    if(is.null(legend.title)) legend.title <- "" # remove .y. in the legend
  }


  # Updating parameters after merging
  #:::::::::::::::::::::::::::::::::::::::::::::::::::::::::

  # Special case for density and histograms:
      # x are variables and y is ..count.. or ..density..
      # after merging ggpubr add a new column .y. which hold x variables
      # User might want to color by x variables as follow color = ".x." and
      # he aren't aware that the column is ".y." --> so we should translate this (see from line 1055)

  user.add.color <- add.params$color
  geom.text.position <- "identity"

  if(merge == "asis" ){
    .grouping.var <- ".y."  # y variables become grouping variable
  }
  else if(merge == "flip"){
    .grouping.var <- opts$x  # x variable becomes grouping variable
     opts$x <- ".y."  # y variables become x tick labels
    if(is.null(xlab)) xlab <- FALSE
  }

  if(merge == "asis" | merge == "flip"){

    if(is_density_plot){
      color <- ifelse(color == ".x.", ".y.", color)
      fill <- ifelse(fill == ".x.", ".y.", fill)
    }

    if(any(c(color, fill) %in% names(data))){
      add.params$color <- font.label$color <- ifelse(color %in% names(data), color, fill)
    }
    else if(!all(c(color, fill) %in% names(data))){
      color <- add.params$color <- font.label$color <-  .grouping.var
      #fill <- "white"
    }
    group <- .grouping.var
    geom.text.position <- position_dodge(0.8)
  }

  if(!combine & merge == "none" & length(opts$y) > 1 & is.null(title))
    title <- opts$y

  if(!combine & merge == "none" & is.null(title)){
    if(length(opts$y) > 1) title <- opts$y
    else if (length(opts$x) > 1 & is_density_plot)  # case of density plot
      title <- opts$x
  }

  # Item to display
  x <- opts$data[[opts$x]] %>% as.vector()
  if(!is.null(select))
    opts$data <- subset(opts$data, x %in% select)
  if(!is.null(remove))
    opts$data <- subset(opts$data, !(x %in% remove))
  if(!is.null(order)) opts$data[[opts$x]] <- factor(opts$data[[opts$x]], levels = order)

  # Add additional options, which can be potentially vectorized
  # when multiple plots
  opts <- opts %>% c(list(title = title, xlab = xlab, ylab = ylab)) %>%
    .compact()
  data <- opts$data
  opts$data <- list(opts$data)
 if(fun_name %in% c("ggline", "ggdotchart")) opts$group <- group
  # Plotting
  #::::::::::::::::::::::::::::::::::::::::::::::::::::::::::::::::::::
  # Apply function to each y variables
  p <- purrr::pmap(opts, fun, color = color, fill = fill, legend = legend,
                   legend.title = legend.title, ggtheme = ggtheme, facet.by = facet.by,
                   add = add, add.params  = add.params ,
                   # group = group, # for line plot
                   user.add.color = user.add.color,
                   label = .lab, # used only in ggbarplot
                   font.label = font.label, repel = repel, label.rectangle = label.rectangle,
                   ...)
  # Faceting
  if(!is.null(facet.by))
    p <-purrr::map(p, facet, facet.by = facet.by, ...)


  # Add labels
  if(!is.null(label) & fun_name != "barplot"){

    if(is.logical(label)){
      if(label) label <- opts$y
    }

    grouping.vars <- intersect(c(facet.by, color, fill), colnames(data))

    label.opts <- font.label %>%
      .add_item(data = data, x = opts$x, y = opts$y,
                label = label, label.select = label.select,
                repel = repel, label.rectangle = label.rectangle,
                family = font.family, parse = parse,
                ggtheme = NULL,
                grouping.vars = grouping.vars, facet.by = facet.by, position = geom.text.position,
                show.legend = show.legend.text)
    p <- purrr::map(p,
                   function(p, label.opts){
                     . <- NULL
                     label.opts %>% .add_item(ggp = p) %>%
                       do.call(ggtext, .)
                   },
                   label.opts
                   )
  }

  # Take into account the legend argument, when the main plot has no legend and ggtext has legend
  p <-purrr::map(p, ggpar, legend = legend, legend.title = legend.title)

  if(.is_list(p) & length(p) == 1) p <- p[[1]]
  p

}


# get the geometry of the first layer
#:::::::::::::::::::::::::::::::::::::::::::::::::::::::::
.geom <- function(p, .layer = 1){

  . <- NULL
  if(is.null(p) | .is_empty(p$layers)) return("")
  class(p$layers[[.layer]]$geom)[1] %>%
    tolower() %>%
    gsub("geom", "", .)
}


# Get the mapping variables of the first layer
#:::::::::::::::::::::::::::::::::::::::::::::::::
.mapping <- function(p){

  if(is.null(p)) return(list())

  . <- NULL

  layer0.mapping <- as.character(p$mapping) %>% gsub("~", "", .)
  layer0.mapping.labels <- p$mapping %>% names()
  names(layer0.mapping) <- layer0.mapping.labels

  layer1.mapping <- NULL

  if(!.is_empty(p$layers)){
    layer1.mapping <- p$layers[[1]]$mapping %>%
      as.character() %>% gsub("~", "", .)
    layer1.mapping.labels <- p$layers[[1]]$mapping %>%
      names()
    names(layer1.mapping) <- layer1.mapping.labels
  }

  c(layer0.mapping, layer1.mapping) %>%
    as.list()
}

# Call geom_exec function to update a plot
#:::::::::::::::::::::::::::::::::::::::::::::::::::::::::
.update_plot <- function(opts, p){
  p + do.call(geom_exec, opts)
}


# Get ggplot2 x and y variable
# ::::::::::::::::::::::::::::::::::::::::::::::::::::::::::
.get_gg_xy_variables <- function(p){
  . <- NULL
  x <- p$mapping['x'] %>% as.character() %>% gsub("~", "", .)
  y <- p$mapping['y'] %>% as.character() %>% gsub("~", "", .)
  xy <- c(x, y)
  names(xy) <- c("x", "y")
  return(xy)
}

# Add mean or median line
# used by ggdensity and gghistogram
#:::::::::::::::::::::::::::::::::::::::::::::::::::::::::::::::::::::::
# p: main plot
# data: data frame
# x: measure variables
# add: center to add
# grouping.vars: grouping variables
.add_center_line <- function(p, add = c("none", "mean", "median"), grouping.vars = NULL,
                             color = "black", linetype = "dashed", size = NULL)
{

  add <- match.arg(add)
  data <- p$data
  # x <- .mapping(p)$x
  .mapping <- .get_gg_xy_variables(p)
  x <- .mapping["x"]

  if(!(add %in% c("mean", "median")))
    return(p)
  compute_center <- switch(add, mean = mean, median = stats::median)

  # NO grouping variable
  if(.is_empty(grouping.vars)) {
    m <- ifelse(add == "mean",
                mean(data[[x]], na.rm = TRUE),
                stats::median(data[[x]], na.rm = TRUE))
    p <- p + geom_exec(geom_vline, data = data,
                       xintercept = m, color = color,
                       linetype = linetype, size = size)
  }
  # Case of grouping variable
  else {
    data_sum <- data %>%
      group_by(!!!syms(grouping.vars)) %>%
      summarise(.center = compute_center(!!sym(x), na.rm = TRUE))
    p <- p + geom_exec(geom_vline, data = data_sum,
                       xintercept = ".center", color = color,
                       linetype = linetype, size = size)
  }

  p
}


# Check legend argument
.check_legend <- function(legend){

  allowed.values <- c("top", "bottom", "left", "right", "none")

  if(is.null(legend) | is.numeric(legend))
    return(legend)
  else if(is.logical(legend)){
    if(legend) legend <- "top"
    else legend <- "none"
  }
  else if(is.character(legend)){
    legend <- legend[1]
    if(!legend %in% allowed.values)
      stop("Argument legend should be one of ", .collapse(allowed.values, sep = ", "))
  }
  return (legend)
}
```
