## Supplementary material for "Water content, transition temperature and fragility influence protection and anhydrobiotic capacity": File S6: utilities_label.R

```
#::::::::::::::::::::::::::::::::::::::::::::::::::::::::
# Helper function for adding annotation to a ggplot
#::::::::::::::::::::::::::::::::::::::::::::::::::::::::


# Get label parameters for each group
#:::::::::::::::::::::::::::::::::::::::::::::::::::::::::
# Returns a data frame with x, y, hjust, vjust
# group.id is the index position of the group in a boxplot for example
.label_params <- function(data, scales, label.x.npc = "left", label.y.npc = "right",
                          label.x = NULL, label.y = NULL, .by = c("group", "panel"),
                          group.id = NULL, ...)
  {

  .by <- match.arg(.by)
  # Check label coordinates for each group
  #:::::::::::::::::::::::::::::::::::::::::::::::::::::::::::
  if(is.null(group.id))
    group.id <- group.id <- abs(data$group[1])
  label.x.npc <- .group_coord(label.x.npc, group.id)
  label.y.npc <- .group_coord(label.y.npc, group.id)
  label.x <- .group_coord(label.x, group.id)
  label.y <- .group_coord(label.y, group.id)

  .check_npc_coord(label.x.npc, axis = "x")
  .check_npc_coord(label.y.npc, axis = "y")


  if (length(label.x) > 0) {
    x <- label.x
    hjust <- 0.5
  } else if (length(label.x.npc) > 0) {
    if (is.numeric(label.x.npc)) {
      x <- scales$x$dimension()[1] + label.x.npc *
        diff(scales$x$dimension())
      hjust <- 0.5
    } else if (is.character(label.x.npc)) {
      if (label.x.npc == "right") {
        x <- scales$x$dimension()[2]
        hjust <- 1
      } else if (label.x.npc %in% c("center", "centre", "middle")) {
        x <- mean(scales$x$dimension())
        hjust <- 0.5
      } else if (label.x.npc == "left") {
        x <- scales$x$dimension()[1]
        hjust <- 0
      }
    }
  }

  if (length(label.y) > 0) {
    y <- label.y
    vjust <- 0.5
  } else if (length(label.y.npc) > 0) {
    if (is.numeric(label.y.npc)) {
      y <- scales$y$dimension()[1] + label.y.npc *
        diff(scales$y$dimension())
      vjust <- 1.4 * group.id - (0.7 * length(group.id))
    } else if (is.character(label.y.npc)) {
      if (label.y.npc == "bottom") {
        y <- scales$y$dimension()[1]
        vjust <- -1.4 * group.id
      } else if (label.y.npc %in% c("center", "centre", "middle")) {
        y <- mean(scales$y$dimension())
        vjust <- 1.4 * group.id - (0.7 * length(group.id))
      } else if (label.y.npc == "top") {
        y <- scales$y$dimension()[2]
        vjust <- 1.4 * group.id
      }
    }
  }
  if(.by == "panel"){
    hjust <- 0.5
    vjust = 0.5
  }

  data.frame(x = x, y = y, hjust = hjust, vjust = vjust)
}


# Get label parameters by group
# Useful in boxplot, where group.ids is the index of the group: 1, 2, 3, etc
# Useful only when computation is done by panel
.label_params_by_group <- function(..., group.ids){
  purrr::map(group.ids,
             function(group.id, ...){.label_params(..., group.id = group.id)},
             ...) %>%
    dplyr::bind_rows() #%>%
    #dplyr::mutate(x = group.ids)

}


# Check label coordinates for each group
#:::::::::::::::::::::::::::::::::::::::::
# coord.values: label coordinate for each group. If too short, they are recycled.
# group.id the id of groups as returned by ggplot_build()
.group_coord <- function(coord.values, group.id){
  if(!.is_empty(coord.values)){
    coord.values <- ifelse(length(coord.values) >= group.id,
                           coord.values[group.id], coord.values[1])
  }
  coord.values
}


# Check NPC coord
#:::::::::::::::::::::::::::::::::::::::::
# npc: Normalised Parent Coordinates.
#   The origin of the viewport is (0, 0) and the viewport has a width and height of 1 unit.
#   For example, (0.5, 0.5) is the centre of the viewport.
# coord: should be between 0 and 1
# axis: should be "x" or "y"
.check_npc_coord <- function(.coord, axis = c("x", "y")){

  axis <- match.arg(axis)
  if(axis == "x")
    allowed.values <- c('right', 'left', 'center', 'centre', 'middle')
  else if(axis == "y")
    allowed.values <- c( 'bottom', 'top', 'center', 'centre', 'middle')

  .message <- paste0("'*.npc coord for ", axis, " axis should be either a numeric value in [0-1] ",
                     "or a character strings including one of ",
                     .collapse(allowed.values, sep = ", "))

  if(!is.null(.coord)){

    if(is.numeric(.coord)){
      if (any(.coord < 0 | .coord > 1)) {
        stop(.message)
      }
    }
    else if(is.character(.coord)){
      if(!(.coord %in% allowed.values))
        stop(.message)

    }
    else
      stop(.message)
  }
}
```
