## Supplementary material for "Water content, transition temperature and fragility influence protection and anhydrobiotic capacity": File S6: S2a_PerwaterVsTgmid_Malt.R

```
setwd("c:/R/Glass")

library(ggplot2) #load in libraries
library(readxl)
library(ggsignif)
library(polynom)
library(hrbrthemes)
library(dplyr)
source("stat_cor.R")
source("utilities.R")
source("utilities_label.R")

#Treh <- read_excel("Treh_glyc.xlsx", sheet = "Trehalose") 
#reading excel sheet into var Treh

#Sucr <- read_excel("Sucr_glyc.xlsx", sheet = "Sucrose")
#reading excel sheet into var Sucr

Malt <- read_excel("Malt_glyc2.xlsx", sheet = "Maltose")
#reading excel sheet into var Sucr

#df1 <- data.frame(y = Treh$Perprot, x = Treh$Perwater, Disaccharide = Treh$Sugar)
#read x and y variables into dataframe df1

#df2 <- data.frame(y = Sucr$Perprot, x = Sucr$Perwater, Disaccharide = Sucr$Sugar)
#read x and y variables into dataframe df1

df3 <- data.frame(y = Malt$Tg_midpt, x = Malt$Perwater, Disaccharide = Malt$Sugar)
#read x and y variables into dataframe df1

#df <- rbind(df1, df2, df3)

p1 <- 
  ggplot(df3, aes(x = x, y = y)) +
  geom_point() +
  geom_smooth(method = lm, fill = "grey", se = TRUE, color = 'red') +
  
  xlab("Water content (%)") +   
  ylab("Glass transition midpoint (°C)") + 
  ggtitle("Maltose water content versus Glass transition midpoint (°C)") +
  theme_bw() +
  theme(plot.title = element_text(hjust = 0.5), strip.text.x = element_blank(), legend.position = "none") 

p1 <- p1 + stat_cor(aes(label = paste(..rr.label.., ..p.label.., sep = "~','~")), 
                    label.x = 1, label.y = 90)
p1

#ggsave(file = "Treh_glyc_TgoffVsPerprot.eps")
#ggsave(file = "Treh_glyc_TgoffVsPerprot.tiff")
ggsave(file = "Treh_glyc_PerwaterVsTgmid_Malt_supp.jpeg")
ggsave("Treh_glyc_PerwaterVsTgmid_Malt_supp.pdf", width = 8, height = 6, units = "in")
```
