## Supplementary material for "Water content, transition temperature and fragility influence protection and anhydrobiotic capacity": File S6: S2c_PerwaterVsTgmid_Sucr.R

```
setwd("c:/R/Glass")

library(ggplot2) #load in libraries
library(readxl)
library(ggsignif)
library(polynom)
library(hrbrthemes)
library(dplyr)
source("stat_cor.R")
source("utilities.R")
source("utilities_label.R")

#Treh <- read_excel("Treh_glyc.xlsx", sheet = "Trehalose") 
#reading excel sheet into var Treh

Sucr <- read_excel("Sucr_glyc.xlsx", sheet = "Sucrose")
#reading excel sheet into var Sucr

#Malt <- read_excel("Malt_glyc.xlsx", sheet = "Maltose")
#reading excel sheet into var Sucr

#df1 <- data.frame(y = Treh$Perprot, x = Treh$Perwater, Disaccharide = Treh$Sugar)
#read x and y variables into dataframe df1

df2 <- data.frame(y = Sucr$Tg_midpt, x = Sucr$Perwater, Disaccharide = Sucr$Sugar)
#read x and y variables into dataframe df1

#df3 <- data.frame(y = Malt$Perprot, x = Malt$Perwater, Disaccharide = Malt$Sugar)
#read x and y variables into dataframe df1

#df <- rbind(df1, df2, df3)

p1 <- 
  ggplot(df2, aes(x = x, y = y)) +
  geom_point() +
  geom_smooth(method = lm, fill = "grey", se = TRUE, color = 'green') +
  
  xlab("Water content (%)") +   
  ylab("Glass transition midpoint (°C)") + 
  ggtitle("Sucrose water content versus Glass transition midpoint (°C)") +
  coord_cartesian(ylim = c(0, 75)) +
  theme_bw() +
  theme(plot.title = element_text(hjust = 0.5), strip.text.x = element_blank(), legend.position = "none") 

p1 <- p1 + stat_cor(aes(label = paste(..rr.label.., ..p.label.., sep = "~','~")), 
                    label.x = 4, label.y = 60)
p1

#ggsave(file = "Treh_glyc_TgoffVsPerprot.eps")
#ggsave(file = "Treh_glyc_TgoffVsPerprot.tiff")
ggsave(file = "Treh_glyc_PerwaterVsTgmid_Sucr_supp.jpeg")
ggsave("Treh_glyc_PerwaterVsTgmid_Sucr_supp.pdf", width = 8, height = 6, units = "in")
```
