## Supplementary material for "Water content, transition temperature and fragility influence protection and anhydrobiotic capacity": File S6: S4_TgallVsPerprot.R

```
setwd("c:/R/Glass")

library(ggplot2) #load in libraries
library(readxl)
library(ggsignif)
library(polynom)
library(hrbrthemes)
library(dplyr)
source("stat_cor.R")
source("utilities.R")
source("utilities_label.R")

Treh <- read_excel("Treh_glyc2.xlsx", sheet = "Trehalose") 
#reading excel sheet into var Treh

Sucr <- read_excel("Sucr_glyc.xlsx", sheet = "Sucrose")
#reading excel sheet into var Sucr

Malt <- read_excel("Malt_glyc2.xlsx", sheet = "Maltose")
#reading excel sheet into var Sucr

df1 <- data.frame(y = Treh$Perprot, x = Treh$Tg_onset, Disaccharide = Treh$Sugar)
df11 <- data.frame(y = Treh$Perprot, x = Treh$Tg_onset)
#read x and y variables into dataframe df1

df2 <- data.frame(y = Sucr$Perprot, x = Sucr$Tg_onset, Disaccharide = Sucr$Sugar)
df12 <- data.frame(y = Sucr$Perprot, x = Sucr$Tg_onset)
#read x and y variables into dataframe df2

df3 <- data.frame(y = Malt$Perprot, x = Malt$Tg_onset, Disaccharide = Malt$Sugar)
df13 <- data.frame(y = Malt$Perprot, x = Malt$Tg_onset)
#read x and y variables into dataframe df3

dfa <- rbind(df1, df2, df3)
#stack Onset data into single dataframe and then add column to identify as onset

Param_a <- c("Onset", "Onset", "Onset", "Onset", "Onset", "Onset", "Onset", 
             "Onset", "Onset", "Onset", "Onset", "Onset", "Onset", "Onset", 
             "Onset", "Onset", "Onset", "Onset")
#create a column Param_a that has values Onset

dfa$Param_a <- Param_a
#bind the column Param_a to the datafram dfa

colnames(dfa) <- c('y', 'x', 'Disaccharide', 'Parameter')
#rename columns so they will match later

df4 <- data.frame(y = Treh$Perprot, x = Treh$Tg_offset, Disaccharide = Treh$Sugar)
df14 <- data.frame(y = Treh$Perprot, x = Treh$Tg_offset)
#read x and y variables into dataframe df4

df5 <- data.frame(y = Sucr$Perprot, x = Sucr$Tg_offset, Disaccharide = Sucr$Sugar)
df15 <- data.frame(y = Sucr$Perprot, x = Sucr$Tg_offset)
#read x and y variables into dataframe df5

df6 <- data.frame(y = Malt$Perprot, x = Malt$Tg_offset, Disaccharide = Malt$Sugar)
df16 <- data.frame(y = Malt$Perprot, x = Malt$Tg_offset)
#read x and y variables into dataframe df6

dfb <- rbind(df4, df5, df6)
#stack Offset data into single dataframe and then add column to identify as Offset

Param_b <- c("Offset", "Offset", "Offset", "Offset", "Offset", "Offset", "Offset", 
             "Offset", "Offset", "Offset", "Offset", "Offset", "Offset", "Offset", 
             "Offset", "Offset", "Offset", "Offset")
#create a column Param_a that has values Offset

dfb$Param_b <- Param_b
#bind the column Param_a to the datafram dfa

colnames(dfb) <- c('y', 'x', 'Disaccharide', 'Parameter')
#rename columns so they will match later

df7 <- data.frame(y = Treh$Perprot, x = Treh$Tg_midpt, Disaccharide = Treh$Sugar)
df17 <- data.frame(y = Treh$Perprot, x = Treh$Tg_midpt)
#read x and y variables into dataframe df7

df8 <- data.frame(y = Sucr$Perprot, x = Sucr$Tg_midpt, Disaccharide = Sucr$Sugar)
df18 <- data.frame(y = Sucr$Perprot, x = Sucr$Tg_midpt)
#read x and y variables into dataframe df8

df9 <- data.frame(y = Malt$Perprot, x = Malt$Tg_midpt, Disaccharide = Malt$Sugar)
df19 <- data.frame(y = Malt$Perprot, x = Malt$Tg_midpt)
#read x and y variables into dataframe df9

dfc <- rbind(df7, df8, df9)
#stack Midpoint data into single dataframe and then add column to identify as midpt

Param_c <- c("Midpoint", "Midpoint", "Midpoint", "Midpoint", "Midpoint", "Midpoint", "Midpoint", 
             "Midpoint", "Midpoint", "Midpoint", "Midpoint", "Midpoint", "Midpoint", "Midpoint", 
             "Midpoint", "Midpoint", "Midpoint", "Midpoint")
#create a column Param_a that has values Midpoint

dfc$Param_c <- Param_c
#bind the column Param_a to the datafram dfa

colnames(dfc) <- c('y', 'x', 'Disaccharide', 'Parameter')
#rename columns so they will match later

df <- rbind(dfa, dfb, dfc)
#Stack dfa, dfb, and dfc into single dataframe with onset, Offset, and midpt
#being grouping parameters

p <- ggplot(df, aes(x = x, y = y, color = Disaccharide, shape = Parameter)) +
  geom_point(size = 2) +
  geom_smooth(method = lm, fill = "grey", se = TRUE) +

  xlab("Glass transition (°C)") +   
  ylab("LDH protection (%)") + 
  ylim(0, 80) +
  ggtitle("Glass transition (°C) versus L-Lactate Dehydrogenase (LDH) protection") +
  theme_bw() +
  theme(plot.title = element_text(hjust = 0.5), strip.text.x = element_blank()) 

p <- p + stat_cor(aes(label = paste(..rr.label.., ..p.label.., sep = "~','~"), 
                        color = Disaccharide), label.x = 70)
p
#ggsave(file = "Treh_glyc_TgonVsPerprot.eps")
#ggsave(file = "Treh_glyc_TgonVsPerprot.tiff")
ggsave(file = "Revised_Treh_glyc_TgallVsPerprot.jpeg")
ggsave("Revised_Treh_glyc_TgallVsPerprot.pdf", width = 8, height = 6, units = "in")
```
