## Supplementary material for "Water content, transition temperature and fragility influence protection and anhydrobiotic capacity": File S6: S6_TgdeltaVsPerprot.R

```
setwd("c:/R/Glass")

library(ggplot2) #load in libraries
library(readxl)
library(ggsignif)
library(polynom)
library(hrbrthemes)
library(dplyr)
source("stat_cor.R")
source("utilities.R")
source("utilities_label.R")

Treh <- read_excel("Treh_glyc2.xlsx", sheet = "Trehalose") 
#reading excel sheet into var Treh

Sucr <- read_excel("Sucr_glyc.xlsx", sheet = "Sucrose")
#reading excel sheet into var Sucr

Malt <- read_excel("Malt_glyc2.xlsx", sheet = "Maltose")
#reading excel sheet into var Sucr

df1 <- data.frame(y = Treh$Perprot, x = Treh$Tg_delta, Disaccharide = Treh$Sugar)
#read x and y variables into dataframe df1

df2 <- data.frame(y = Sucr$Perprot, x = Sucr$Tg_delta, Disaccharide = Sucr$Sugar)
#read x and y variables into dataframe df1

df3 <- data.frame(y = Malt$Perprot, x = Malt$Tg_delta, Disaccharide = Malt$Sugar)
#read x and y variables into dataframe df1

df <- rbind(df1, df2, df3)

p1 <- 
  ggplot(df, aes(x = x, y = y, color = Disaccharide)) +
  geom_point() +
  geom_smooth(method = lm, fill = "grey", se = TRUE) +

  xlab("Enzyme protection assay termperature difference from glass transition temperature (°C)") +   
  ylab("LDH protection (%)") + 
  #ggtitle("Glass transition offset (°C) versus L-Lactate Dehydrogenase (LDH) protection") +
  theme_bw()

p1 <- p1 + stat_cor(aes(label = paste(..rr.label.., ..p.label.., sep = "~','~"), 
                        color = Disaccharide), label.x = -50)
p1

#ggsave(file = "Treh_glyc_TgonVsPerprot.eps")
#ggsave(file = "Treh_glyc_TgdeltaVsPerprot.tiff")
ggsave(file = "Revised_Treh_glyc_TgdeltaVsPerprot.jpeg")
ggsave(file = "Revised_Treh_glyc_TgdeltaVsPerprot.pdf", width = 8, height = 6, units = "in")
```
