## Supplementary material for "Water content, transition temperature and fragility influence protection and anhydrobiotic capacity": File S6: S8b_mVsPerprot_mixed_sucr.R

```
setwd("c:/R/Glass")

library(ggplot2) #load in libraries
library(readxl)
library(ggsignif)
library(polynom)
library(hrbrthemes)
library(dplyr)
source("stat_cor.R")
source("utilities.R")
source("utilities_label.R")

#Treh <- read_excel("Treh_glyc2.xlsx", sheet = "Sheet2") 
#reading excel sheet into var Treh

Sucr <- read_excel("Sucr_glyc.xlsx", sheet = "Sheet2")
#reading excel sheet into var Sucr

#Malt <- read_excel("Malt_glyc2.xlsx", sheet = "Sheet2")
#reading excel sheet into var Sucr

#df1 <- data.frame(y = Treh$Perprot, x = Treh$m, Disaccharide = Treh$Sugar)
#read x and y variables into dataframe df1

df <- data.frame(y = Sucr$Perprot, x = Sucr$m, Disaccharide = Sucr$Sugar)
#read x and y variables into dataframe df1

#df <- data.frame(y = Malt$Perprot, x = Malt$m, Disaccharide = Malt$Sugar)
#read x and y variables into dataframe df1

#df <- rbind(df1, df2, df3)

p1 <- 
    ggplot(df, aes(x = x, y = y)) +
    geom_point() +
    geom_smooth(method = lm, fill = "grey",  se = TRUE, color = 'green') +
    
    coord_cartesian(ylim = c(0, 45)) +
    xlab("Fragility (m-index)") +   
    ylab("LDH protection (%)") + 
    ggtitle("Glass fragility (m-index) versus L-Lactate Dehydrogenase (LDH) protection") +
    theme_bw() +
    theme(plot.title = element_text(hjust = 0.5), strip.text.x = element_blank(), legend.position = "none") 

p1 <- p1 + stat_cor(aes(label = paste(..rr.label.., ..p.label.., sep = "~','~")), 
                    label.x = 80, label.y = 35)
p1

#ggsave(file = "Treh_glyc_TgonVsPerprot.eps")
#ggsave(file = "Treh_glyc_TgonVsPerprot.tiff")
ggsave(file = "Revised_Treh_glyc_mVsPerprot_mixed_sucr.jpeg")
ggsave("Revised_Treh_glyc_mVsPerprot_mixed_sucr.pdf", width = 8, height = 6, units = "in")
```
